# Free fatty acid receptor 4 is a nutrient sensor that resolves inflammation to maintain cardiac homeostasis

**DOI:** 10.1101/776294

**Authors:** Katherine A. Murphy, Brian A. Harsch, Chastity L. Healy, Sonal S. Joshi, Shue Huang, Rachel E. Walker, Brandon M. Wagner, Katherine M. Ernste, Wei Huang, Robert Block, Casey D. Wright, Nathan Tintle, Brian C. Jensen, Gregory C Shearer, Timothy D. O’Connell

**Author notes:** Correspondence to: Timothy D. O’Connell, PhD, Department of Integrative Biology and Physiology, University of Minnesota School of Medicine, 3--141 CCRB, 2231 6th Street SE, Minneapolis, MN 55414, Gregory C Shearer, PhD, Department of Nutritional Sciences, 110 Chandlee Laboratory, University Park, PA 16802.

## Abstract

**Background:** Non-­resolving activation of immune responses is central to the pathogenesis of heart failure (HF). Free fatty acid receptor 4 (Ffar4) is a G-protein coupled receptor (GPR) for medium-and long-chain fatty acids (FA) that regulates metabolism and attenuates inflammation in diabetes and obesity. Here, we tested the hypothesis that Ffar4 functions as a cardioprotective nutrient sensor that resolves inflammation to maintain cardiac homeostasis.

**Methods:** Mice with systemic deletion of Ffar4 (Ffar4KO) were subjected to pressure overload by transverse aortic constriction (TAC). Transcriptome analysis of cardiac myocytes was performed three days post-TAC. Additionally, Ffar4-mediated effects on inflammatory oxylipin production in cardiac myocytes and oxylipin composition in plasma lipoproteins were evaluated.

**Results:** In Ffar4KO mice, TAC induced more severe remodeling, identifying an entirely novel cardioprotective role for Ffar4 in the heart. Transcriptome analysis 3-days post-TAC indicated a failure to induce cell death and inflammatory genes in Ffar4KO cardiac myocytes, as well as a specific failure to induce cytoplasmic phospholipase A_2_α (cPLA_2_α) signaling genes. In cardiac myocytes, Ffar4 signaling through cPLA_2_α-cytochrome p450 ω/ω-1 hydroxylase induced production of the EPA-derived anti-inflammatory oxylipin 18-hydroxyeicosapentaenoic acid (18-HEPE). Systemically, loss of Ffar4 altered oxylipin content in circulating plasma lipoproteins consistent with a loss of anti-inflammatory oxylipins at baseline, and inability to produce both pro-inflammatory and pro-resolving oxylipins following TAC. Finally, we confirmed that Ffar4 is expressed in human heart and down-regulated in HF.

**Conclusions:** Our results identify a novel function for Ffar4 in the heart as a FA nutrient sensor that resolves inflammation to maintain cardiac homeostasis.

## Introduction

Heart failure (HF) is a complex and heterogeneous clinical syndrome, and in the last 10 years, two broadly defined phenotypes have emerged: HF with reduced ejection fraction (HFrEF) and HF with preserved ejection fraction (HFpEF). It is now clear that failure to resolve immune activation is central to the pathogenesis of HF, both HFrEF and HFpEF. In HFrEF, typically observed secondary to ischemic injury, cardiac cell death drives an initial immune response to induce scar formation, but failure to resolve inflammation can worsen long-term ventricular remodeling (Review: (1)). In HFpEF, it has been proposed that comorbidities associated with metabolic syndrome induce peripheral inflammation that drives remodeling (Reviews (2–4)). Clinically, targeting the innate immune response has shown some success. The Canakinumab Anti-inflammatory Thrombosis Outcome Study (CANTOS) trial demonstrated a reduction in recurring cardiovascular events by targeting interleukin-1β (IL-1β) in patients with a prior myocardial infarction and high C-reactive protein (5). While the results of CANTOS are promising, successes targeting immune modulators have not been universal (6), suggesting the need for better understanding of the immune response in HF and the identification new therapeutic targets.

Free fatty acid receptor 4 (Ffar4, GPR120) is a G-protein coupled receptor (GPR) that functions as a nutrient sensor for fatty acids (FA) to regulate metabolism and attenuate inflammation (7, 8). The endogenous ligands for Ffar4 include medium and long chain (C10-C22), saturated (SFA), mono-unsaturated (MUFA), and poly-unsaturated fatty acids (PUFA), which bind the receptor with affinities in the low μM range. In terms of agonist efficacy, generally, PUFAs are full agonists, whereas SFAs are partial agonists (9–11). Consistent with a primary role in regulating metabolism, Ffar4 is expressed in enteroendocrine cells in the GI tract (11), α, β, and δ-cells in the pancreas (11–13), and both white and brown adipose (14, 15). However, Ffar4 is found in many additional tissues with high levels of expression in the lung, and lower levels of expression in brain, heart, taste buds, and immune cells, including macrophages (10, 16–18). Ffar4 signals through both G_q/11_ and βarrestin-2 (βArr2) mediated pathways, which appears to be cell-type dependent (10, 17, 19, 20). Interestingly, humans express two isoforms of Ffar4, short and long (Ffar4S, Ffar4L), differentiated by a 16 amino acid insertion in the third intracellular loop of the long-isoform, whereas other species express only one isoform, homologous to Ffar4S in humans (20). The Ffar4L isoform only signals through β-Arr2 and is unable to activate G_q/11_-mediated signaling (20).

Currently, nothing is known about Ffar4 function in the heart. Previous studies have indicated that ω3-polyunsaturated fatty acids (ω3-PUFAs) signaling through Ffar4 attenuate inflammation and obesity (10). Clinical studies indicate that ω3-PUFAs, eicosapentaenoic acid (EPA) and docosahexaenoic acid (DHA), improve outcomes in coronary heart disease (21–26) and HF (27–30), but the mechanisms underlying this benefit remain unclear. In mice, we were the first to demonstrate that EPA prevents fibrosis and contractile dysfunction in response to pathologic stress in the heart, but EPA was not incorporated into cardiac myocytes or fibroblasts, the traditional mechanism of action for EPA (16, 31). However, we found that Ffar4 is expressed in cardiac myocytes and fibroblasts, and that in cardiac fibroblasts, Ffar4 was sufficient and necessary to prevent TGFβ1-induced fibrosis (16). These finding suggest that Ffar4 might mediate ω3-PUFA cardioprotection. However, to date, no studies have directly addressed the function of Ffar4 in the heart, nor any link to ω3-mediated cardioprotection.

Based on these findings, we hypothesized that Ffar4 is a cardioprotective nutrient sensor for medium and long chain fatty acids that resolves inflammation to maintain cardiac homeostasis. To test this hypothesis, we employed mice with systemic deletion of Ffar4 (Ffar4KO mice) to determine if Ffar4 is necessary for an adaptive response to pathologic pressure overload induced by transverse aortic constriction (TAC). Here, we report that TAC induced more severe remodeling in Ffar4KO mice, identifying, for the first time, an entirely novel cardioprotective role for Ffar4 in the heart. In cardiac myocytes, Ffar4 signaling through the cPLA_2_α-Cytochrome P450_*hydroxylase*_ (CYP_*hydroxylase*_) axis specifically induced production of the EPA-derived oxylipin 18-hydroxyeicosapentaenoic acid (18-HEPE), the precursor to E-resolvins, a class of inflammation resolving oxylipins. Ffar4-mediated production of 18-HEPE was absent in Ffar4KO cardiac myocytes, providing a potential mechanism to explain the worse remodeling in the Ffar4KO post-TAC. Systemically, loss of Ffar4 altered oxylipin content in circulating plasma lipoproteins characterized by a loss of an anti-inflammatory oxylipins at baseline, and inability to produce both pro-inflammatory and pro-resolving oxylipins following TAC. Finally, we confirmed that Ffar4 is expressed in human heart and down-regulated in heart failure. In summary, our data suggest an entirely novel paradigm whereby FAs function as signaling molecules that activate GPR signaling through Ffar4 to resolve inflammation and maintain cardiac homeostasis.

## Methods

### Mice

For this study, all experimental mice were placed on a control diet (described below) at 8 weeks of age. At 12 weeks, male and female, WT and Ffar4KO mice were randomized and enrolled into the study. For all experimental analyses, data collection was done with investigator blinded to genotype and treatment.

All procedures on animals conformed to the Public Health Service (PHS) Policy on Humane Care and Use of Laboratory Animals and were reviewed and approved by the Institutional Animal Care and Use Committee at the University of Minnesota.

Ffar4KO mice were generated from cryopreserved sperm from C57Bl/6N-*Ffar4*^tm1(KOMP)Vlcg^ (Design ID: 15078;; Project ID: VG15078) purchased from The KOMP Repository, UC-Davis (Davis, CA, USA). Cryo-recovery of the mouse line was performed at the UMN Mouse Genetics Laboratory through *in vitro* fertilization (IVF) using C57Bl/6J female recipients (#000664; The Jackson Laboratory, Bar Harbor ME, USA). The line was maintained by hemizygous breeding to C57Bl/6J for backcrossing. Homozygous mice were crossed to produce wild type and knock out offspring. Genotype was determined by PCR using the primer design provided by KOMP Repository:

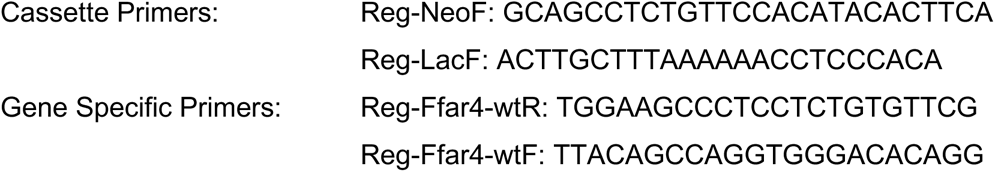

### Diet

Since fatty acids are known agonists of Ffar4, we sought to control the fatty acid profile of the diet by feeding mice a custom chow from Dyets, Inc (#180539;; Bethlehem, PA, USA) beginning at 8 weeks of age. This chow is a modified version of the AIN-93M purified rodent diet used in our previous studies, with corn oil replacing soybean oil (diet composition is listed in Supplemental Table 1) (16, 31).

### Transverse Aortic Constriction (TAC)

Transverse aortic constriction (TAC) surgery was performed as previously described (16, 31–34). Baseline measurements of cardiac function by echocardiography were collected prior to surgery. For surgery, mice were anesthetized with 3% isoflurane and maintained at 1.5% isoflurane without intubation. A small incision was made slightly left of the midline and above the left clavicle without entering the pleural cavity. The muscle tissue and lobes of the thymus were retracted to expose the aortic arch. A 7-0 surgical suture was threaded under the aortic arch and tied-off against a small piece of a blunt 28-gauge needle followed by immediate removal of the needle. To close the original incision, the muscle layer and skin were secured separately with 4-0 continuous sutures. Buprenorphine (0.1 mg/kg IP) was administered for pain management during the first 24 hours post-surgery and as needed thereafter. Sham surgery was identical except for ligation of the aorta. Pulsed-wave Doppler by echocardiography was used to confirm pressure gradients by evaluating aortic flow velocity (AoV, Supplemental Tables 3B and 4B) 7-days post-surgery at the site of constriction.

### Measurement of cardiac function by echocardiography

Echocardiography was performed before TAC (baseline), 7-days post-TAC to measure aortic velocity (AoV) to validate the TAC surgery, and 4 weeks post-TAC using the Vevo 2100 (FujiFilm VisualSonics Inc. Toronto, ON, Canada) with a MS550 transducer. For all measurements, mice were anesthetized with isoflurane, gently restrained in the supine position on the prewarmed monitoring pad, and echocardiographic images were captured as mice were recovering from anesthesia to achieve a target heart rate (HR) of 450 – 500 bpm. Parasternal long axis M-mode images of the left ventricle were captured to measure left ventricular wall thicknesses (LVPWs: systolic left ventricular posterior wall; LVPWd: diastolic left ventricular posterior wall), left ventricular internal diameters (LVIDs: systolic left ventricular internal diameter; LVIDd: diastolic left ventricular internal diameter), left ventricular volumes (ESV: end systolic volume, (7.0/(2.4 + LVIDs))*LVIDs^3^; EDV: end diastolic volume, (7.0/(2.4 + LVIDd))*LVIDd^3^), fractional shortening (FS: 100*((LVIDd – LVIDs)/LVIDd)), ejection fraction (EF: 100*((EDV – ESV)/EDV)), stroke volume (SV: EDV – ESV), and cardiac output (CO: SV*HR). Pulsed-wave Doppler images of the aortic arch were recorded at the site of constriction to measure peak aortic velocity (AoV) and pressure gradient (PG: (4*AoV^2^)/1000). Pulsed-wave Doppler images of the apical four-chamber view were taken to measure mitral flow velocities (E wave and A wave to calculate E/A ratio) as well as mitral annulus tissue velocity (E/E’: peak early transmitral flow velocity/peak early diastolic mitral annular velocity).

### Isolation and culture of adult cardiac myocytes

We previously described procedures for the isolation and culture of adult mouse cardiac myocytes (35). Here, myocytes were isolated 3-days post-TAC from male WT and Ffar4KO, sham and TAC operated mice for analysis of myocyte transcriptomes, or were cultured from WT and Ffar4KO hearts for analysis of Ffar4 signaling. Briefly, mice were anesthetized with isoflurane (3% for induction, 1.5% for maintenance), injected with heparin (100 IU/mL), the pleural cavity was opened and the heart removed, cannulated on a retrograde perfusion apparatus, and perfused with collagenase type II to dissociate ventricular myocytes. Isolated cardiac myocytes were plated at a density of 50 rod-shaped myocytes per square millimeter on laminin-coated culture dishes. Myocytes were cultured in MEM with Hank’s Balanced Salt Solution, 1 mg/ml bovine serum albumin, 10 mM 2,3-butanedione monoxime, and 100 U/ml Penicillin in a 4.5% CO_2_ incubator (%CO_2_ determined empirically to maintain culture medium at pH 7.0) at 37°C. All reagents were purchased from Millipore Sigma (Burlington, MA, USA) unless otherwise specified. Full details of the buffers, enzymes, cell culture medium, all procedures for isolation and culturing of adult mouse ventricular myocytes (AMVM) and a detailed diagram of the perfusion apparatus were described in detail previously (35). Based on our previous experience, myocyte isolation are ~95% pure, but we cannot exclude the possibility of minimal contamination with fibroblasts and endothelial cells (35). Following plating, myocytes were counted at a magnification of 20X to determine cell viability, and myocytes were incubated in a 4.5% CO_2_ incubator at 37°C overnight. For analysis of Ffar4 signaling, cardiac myocytes were treated with the Ffar4 agonist TUG-891(Cayman Chemical, MI, USA) (50 μM) for 0-60 minutes.

### Quantification of erythrocyte FA composition

At the study endpoint, blood from the posterior vena cava was collected into EDTA tubes. Red blood cell fractions were separated and analyzed for fatty acid composition as previously described (16, 31). Briefly, 50 μL of isolated erythrocytes were methylated with 14% boron trifluoride in methanol by incubation for 10 minutes at 100°C. Fatty acid methyl esters were extracted in hexane and analyzed by a GC-2010 gas chromatography system fitted with a QP2010 mass spectrometer (Shimadzu, Japan) using a Supelco SP-2560 fused silica column (Supelco, Bellefonte, PA). Area counts were obtained using Shimadzu GCMSolution software with multiple ion counts of characteristic fatty acids ions. Each fatty acid was quantified as mass percent of total fatty acids. In general, only minor differences in red blood cell fatty acid composition were detected between WT and Ffar4KO following TAC (Supplemental Table 2).

### Tissue histology

Four weeks after TAC surgery, hearts were arrested in diastole with 60mM KCl, excised, and weighed. Extracted hearts were cannulated and perfused with PBS with 60mM KCl, followed by 4% paraformaldehyde. The atria were removed from the fixed hearts prior to embedding in paraffin. Sectioning was performed by AML Laboratories (Jacksonville, FL) providing a transverse view of the ventricles. Lungs were also collected and lung weights were recorded.

Paraffin embedded ventricular sections were deparaffinized with xylene and rehydrated in ethanol. Sections were stained in 0.1% solution of Sirius red (direct red 80, Sigma-Aldrich, St Louis, MO) and fast green (Sigma-Aldrich) in 1.2% picric acid (Ricca Chemical Company, Arlington, TX), followed by dehydration in ethanol and xylene. Sections were imaged at 4× magnification. Ventricular fibrosis (as percent of total ventricular area) was quantified using Fiji software (NIH) (16, 31). Ventricular fibrosis was quantified using images captured at 4X magnification and included both the right and left ventricle. The threshold settings were adjusted to highlight and calculate the total tissue area or picrosirius red positively stained area.

### Oxylipin analysis

Plasma lipoproteins were separated by FPLC followed by measurement of esterified oxylipin (HDL, LDL, and VLDL) or unesterified (albumin). 150 μL of mouse EDTA plasma was thawed and filtered by centrifugation at 10,000 g for 5 min using Ultrafree Durapore PVDF filter (pore-size 0.2 μM; Millipore, Bedford, MA). The filtrate was then injected onto an AKTA Purifier FPLC (Amersham Biosciences, Sweden) and run at a flow rate of 0.5 mL/min in 1mM EDTA, 0.9% NaCl saline solution, pH 7.4 using an additional 100 μL of phosphate buffer solution to fill injection loop. Elutions were monitored at a UV absorbance of 280 nm and lipoproteins were separated using a Superose-6 10/300 GL size exclusion column. Fractions were collected every 0.5 min using a Foxy 200 fraction collector and were pooled for each lipoprotein fraction and stored at −80 °C. VLDL, LDL, HDL, and albumin fractions (100 μL) were spiked with BHT/EDTA (0.2 mg/mL), four deuterated octadecanoid and eicosanoid surrogates (20 μL of 1000 nM concentration with final concentration of 50 nM after reconstitution) and subjected to liquid-liquid extraction to isolate lipid content. Samples were then hydrolyzed in 0.1 M methanolic sodium hydroxide to release ester-linked oxylipins and subjected to solid phase extraction using Chromabond HLB sorbent columns. Oxylipins were eluted with 0.5 mL of methanol with 0.1% acetic acid and 1 mL of ethyl acetate and dried under nitrogen stream and reconstituted in 200 μL methanol acetonitrile (1:1) with 100 nM of 1-cyclohexyluriedo-3-dodecanoic acid used as internal standard.

Samples were analyzed by liquid chromatography using a Waters Acquity UPLC coupled to Waters Xevo triple quadrupole mass spectrometer equipped with electrospray ionization source (Waters, Milford, MA). 5 μL of the extract was injected and separation was performed using a CORTECS UPLC C18 2.1 × 100 mm with 1.6 μM particle size column. Flow rate was set at 500 μL/min and consisted of a gradient run using water with 0.1% acetic acid (Solvent A) and acetonitrile isopropanol, 90:10 (Solvent B) for 15 minutes (0-12 min from 25% B to 95% B, 12-12.5 min 95% B, 12.5-15 min 25% B). Electrospray ionization operated in negative ion mode with capillary set at 2.7 kV, desolvation temperature set at 600 °C, and source temp set to 150°C. Optimal oxylipin MRM transitions were identified by direct injection of pure standards onto the mass spectrometer and using cone voltage and collision energy ramps to optimize detection and most prevalent daughter fragments. Calibration curves were generated prior to each run using standards for each oxylipin. Peak detection and integrations were achieved through Target Lynx (Waters, Milford, MA) and each peak inspected for accuracy and corrected when needed.

### RNA-seq

Three days after surgery, cardiac myocytes were isolated from WT sham (n=4), WT TAC (n=5), Ffar4KO sham (n=4) and Ffar4KO TAC (n=8) male mice. RNA was isolated using RNeasy Fibrous Tissue Mini Kit (Qiagen). Dual-indexed Clontech Pico Mammalian stranded RNA libraries were made. 125bp paired end sequencing was performed using the HiSeq 2500 sequencer (Illumina) by the University of Minnesota Genomics Center.

Data was analyzed by the University of Minnesota Informatics Institute. 2 × 125bp FastQ paired end reads (n=8.4 Million average per sample) were trimmed using Trimmomatic (v 0.33) enabled with the optional “-q” option; 3bp sliding-window trimming from 3’ end requiring minimum Q30. Quality control on raw sequence data for each sample was performed with FastQC. Read mapping was performed via Hisat2 (v2.1.0) using the mouse genome (mm10) as reference. Gene quantification was done via Cuffquant for FPKM values and Feature Counts for raw read counts. Differentially expressed genes were identified using the edgeR (negative binomial) feature in CLC Genomics WorkBench (CLCGWB) (Qiagen, Redwood City, CA) using raw read counts. We filtered the generated list based on a minimum 1.7× Absolute Fold Change and FDR corrected p < 0.05. Two lists were generated of differentially expressed genes from 1.) WT sham compared to WT TAC animals (2789 genes) and 2.) Ffar4KO sham compared to Ffar4KO TAC animals (1656 genes). The lists were compared to each other to identify genes that overlapped between the two lists to identify common genes (1409) and those genes that were unique to the WT mice (1380 genes) and Ffar4KO mice (247 genes). The genes were further annotated in CLCGW based on Gene Ontology (GO) terms for biological function from MGI. Genes were further sorted based on the indicated GO Terms into eight categories.

Principal component analyses used CPMs that were filtered based on gene size (excluding genes less than 200bp) and variance less than 1 in raw read counts. The actual cpm values are log2 transformed and plotted using the PCA function in R.

### RT-PCR for Ffar4 and Ffar1 expression in heart

#### Human heart samples

Ventricular myocardium was obtained from non-failing and failing human hearts through the Duke Human Heart Repository (DHHR) as previously described (36). Briefly, failing human myocardium was acquired from the left ventricular free wall of explanted hearts following cardiac transplantation. Non-failing left ventricular tissue was acquired from donors whose hearts were not utilized for transplant with permission from Carolina Donor Services. No HIPAA information was provided with any of the samples used in this study. Approximately 20-40 mg of human ventricular myocardium was homogenized in 1 mL of Trizol (Life Technologies #15596-026, Carlsbad, CA) with a TissueLyser LT (Qiagen N.V. #69980, Venlo, The Netherlands). The lysate centrifuged at 12,000g (15 min at 4°C) after the addition of chloroform (200 μL). Isopropanol (0.5 mL) was then added to the aqueous phase, centrifuged at 12,000 g (10 min at 4°C). The resulting RNA pellet was washed with 1 mL of 75% ethanol, then centrifuged at 7500 g (5 min at 4°C) and resuspended in RNase-free water.

Primers used to detect human Ffar4:

hFfar4: gctcatctggggctattcg
hFfar4: gcssstcgaaatttcctggt

(The above primers detected Ffar4S (NM_001195755.1) and Ffar4L (NM_181745.3).) hFfar4 aagagctgtcgtgactcacagt (unique for Ffar4L, position 720 - 741) hFfar4 aagagggtgcggaagagc Note: Using these primers, PCR reactions will detect Ffar4S+Ffar4L or only Ffar4L, but not Ffar4S alone.

### Statistics

Cardiac phenotyping was analyzed using an independent samples, Welch’s t-test, comparing WT TAC mice to Ffar4KO TAC mice, to test the hypothesis that the Ffar4KO would have different cardiac phenotypes when under TAC, versus the WT, and to account for the unequal variances observed between the sham and TAC groups. Where specified, principal components analysis (PCA) was used for dimension reduction of oxylipin matrices on log-transformed, standardized concentrations. Mixed models were used to account for within mouse time-dependent changes in oxylipins (Figure 4) or within mouse lipoprotein differences (Figure 5). Statistical significance is indicated as a p value less than 0.05; Tukey’s test was used to test for specified post-hoc differences using JMP version 13.2.1.

## Results

### Ffar4 is necessary to mitigate the pathologic hypertrophic response induced by TAC

To test the hypothesis that Ffar4 is cardioprotective, wild-type (WT) and Ffar4KO mice were subjected to TAC, a model of pathologic pressure overload. After four weeks, TAC induced an exaggerated hypertrophic response in male Ffar4KO mice relative to WT, indicated by an increase in heart weight (HW) and heart weight-to-body weight ratio (HW/BW), with no significant difference in BW (Figures 1A and B and Supplemental Table 3A). Based on our prior studies demonstrating that in cultured fibroblasts, Ffar4 is sufficient and necessary to prevent TGFβ1-induced fibrosis, we expected that TAC would produce more fibrosis in the Ffar4KO. However, we were surprised to find that TAC produced similar levels of fibrosis in WT and Ffar4KO hearts (Figures 1C and D). TAC also caused a trend towards increased lung weights (LW) in male Ffar4KO mice, which is consistent with a worsened response to TAC (Figure 1E). Unexpectedly, the exaggerated hypertrophic response observed in male Ffar4KO mice, was not observed in female Ffar4KO mice. In female mice, TAC induced a similar degree of hypertrophy in WT and Ffar4KO mice, with no significant differences detected in HW, fibrosis, or LW (Figures 1F-H, Supplemental Table 4A).

**Figure 1.**
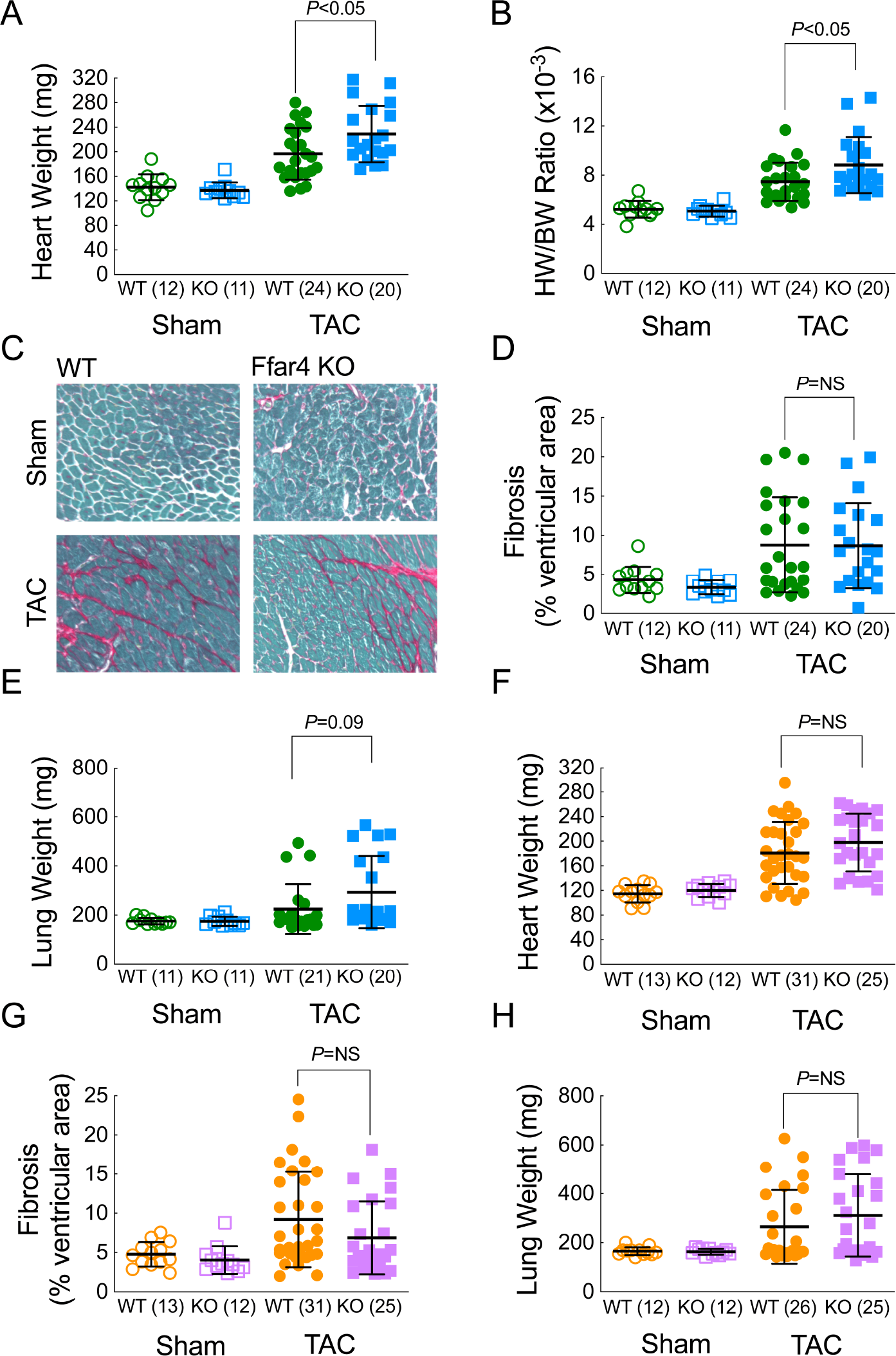
Ffar4 is necessary to mitigate the pathologic hypertrophic response induced by TAC. Four weeks following TAC or sham surgery, mice were euthanized and hearts were collected for morphological analysis. **A**, Heart weight (HW) and **B**, heart weight-to-body weight ratio (HW/BW) of male WT and Ffar4KO mice. **C**, Representative images of ventricular fibrosis quantified from ventricular cross sections from male WT and Ffar4KO mice stained with Sirius red to measure fibrosis, with Fast Green used as a counterstain. **D**, Ventricular fibrosis quantified by fibrotic area (Sirius red)/total ventricular area (Fast green) from male WT and Ffar4KO mice. **E**, Lung weight of male WT and Ffar4KO mice. **F**, Heart weight; **G**, fibrosis and **H**, lung weight for female WT and Ffar4KO mice. Data were compared by a Welch’s two sample t test. Error bars represent the mean with SD.

### Ffar4 is necessary to attenuate systolic and diastolic dysfunction induced by TAC

After four weeks, TAC induced more significant systolic and diastolic dysfunction in male Ffar4KO mice relative to WT, indicated by a more dramatic decrease in ejection fraction (EF, Figure 2A) and a greater increase in E/A ratio (Figure 2B). TAC had only a minor effect on ventricular geometry, with only a trend towards increased end-diastolic and end-systolic volumes (EDV, ESV) (Figures 2C-D, Supplemental Table 3B). However, these results do suggest that male Ffar4KO mice might be progressing to greater eccentric remodeling. In female Ffar4KO and WT hearts, TAC induced a similar degree of systolic and diastolic dysfunction. However, EDV and ESV were both significantly increased, the only significant effects observed in female Ffar4KO mice (Figures 2E-H).

**Figure 2.**
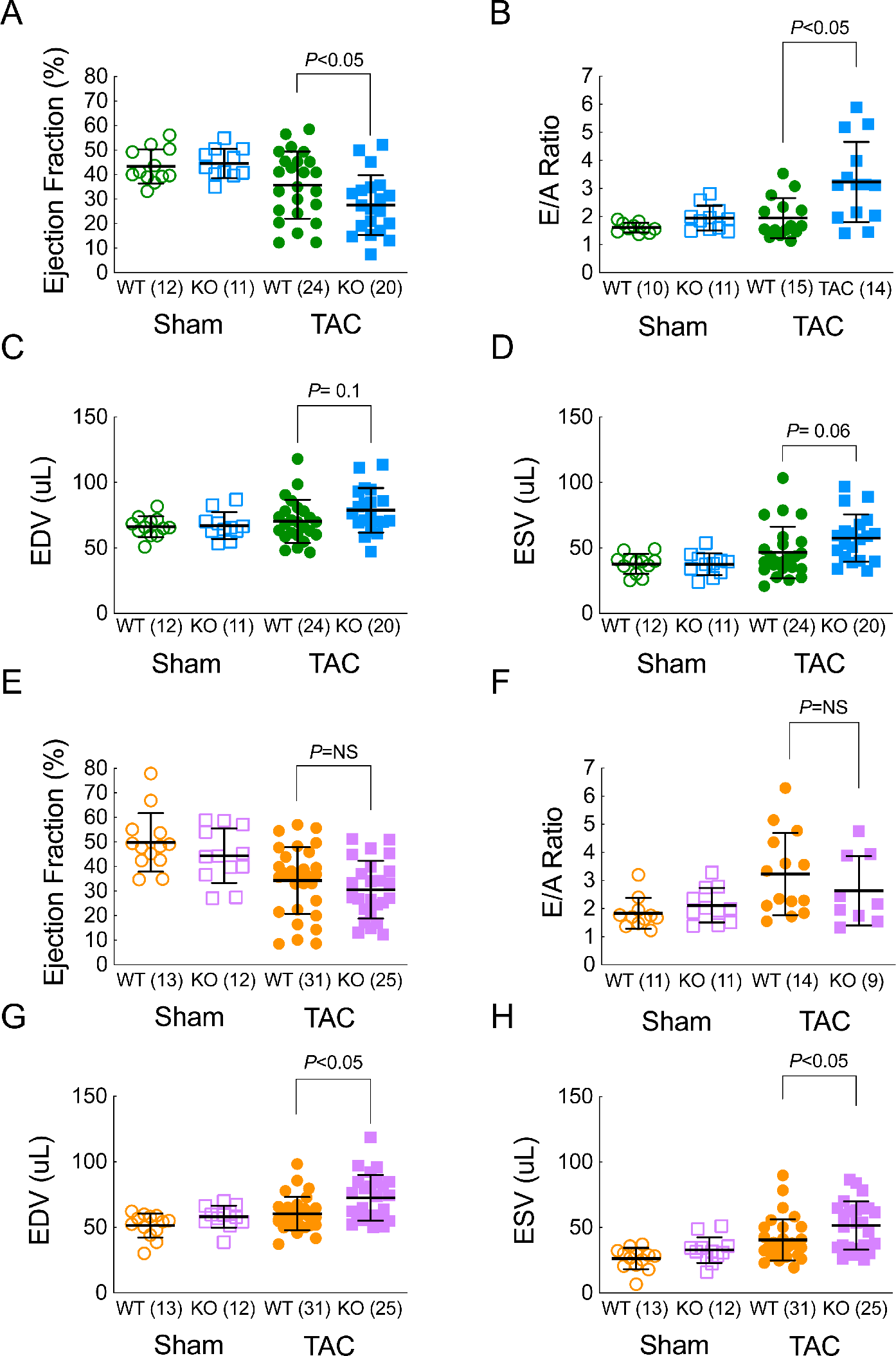
Ffar4 is necessary to attenuate systolic and diastolic dysfunction induced by TAC. Cardiac function measured by echocardiography in male **A-D**, and female **E-H**, WT and Ffar4KO mice four-weeks post-TAC or sham surgery. Males: **A**, ejection fraction (EF, %); **B**, E/A ratio; **C**, end diastolic volume (EDV, μL) and **D**, end systolic volume (ESV, μL). Females: **E**, ejection fraction (EF, %); **F**, E/A ratio; **G**, end diastolic volume (EDV, μL) and **H**. end systolic volume (ESV, μL). Data were compared by a Welch’s two sample t test. Error bars represent the mean with SD.

In summary, our results demonstrated that TAC induced a more severe remodeling response in male Ffar4KO mice, indicating for the first time that Ffar4 is necessary for an adaptive response to pathologic stress in the heart. The surprising lack of an exacerbated fibrotic response in the Ffar4KO, as we previously predicted, would indicate that the more severe remodeling observed in the Ffar4KO heart was most likely due to the lack of cardioprotective Ffar4 signaling in cardiac myocytes. Our results also indicated the potential for a significant sex-based difference in the requirement for Ffar4 in response to pathologic stress in the heart. There are two potential explanations for this sex-based difference; 1.) The response to TAC in female WT mice was greater than male WT mice (58% increase in HW, TAC versus Sham in females, 38% increase in males, despite similar pressure gradients, Figures 1A versus 1F, and Supplemental Table 3A-B and 4A-B). Given that the response to TAC is time-dependent, perhaps we would see more of a difference if female mice were examined at an earlier time point (or later); or 2.) This represents a significant sex-based difference in Ffar4 function in the heart.

### Ffar4 is necessary for induction of cardioprotective inflammatory and cell death genes, and specifically expression of genes associated with phospholipase A2 signaling

To the best of our knowledge, there are no prior reports regarding the function of Ffar4 in cardiac myocytes. Therefore, to gain insight into the potentially cardioprotective role of Ffar4 in cardiac myocytes, we used RNAseq to analyze transcriptomes of myocytes isolated from WT and Ffar4KO hearts three days post-TAC. Initially, a principal component analysis (PCA) revealed only minor differences in the transcriptomes between genotypes, whereas much larger differences were induced by surgery (Figure 3A). However, a separate analysis of genes specifically regulated by TAC in either WT or Ffar4KO cardiac myocytes identified substantial differences in the number of genes differentially expressed post-TAC depending on genotype (Figure 3B). TAC altered expression of 2,789 genes (up or down) by 1.7-fold or more in WT cardiac myocytes, whereas TAC changed expression of only 1,656 genes in Ffar4KO myocytes. Comparison of these lists revealed 1409 genes were common between the two genotypes, whereas 1380 genes were unique to WT cardiac myocytes, but only 247 genes were unique to Ffar4KO myocytes. (Figure 3B). Individual genes from each genotype were further sorted by Gene Ontology terms for biological function for eight different categories that reflect cardiac myocyte biology; cell death, inflammation, contractile function, angiogenesis, fibrosis, GPR, fatty acid metabolism, and hypertrophy (Figure 3C, Supplemental Tables 5-12). Genes categorized by biological function for cell death and inflammation had the most differentially expressed genes, with significantly more genes identified in WT cardiac myocytes (Figure 3C, Table 1). Interestingly, we also identified genes specifically associated with activation of cPLA_2_α signaling, an Ffar4 signaling pathway in macrophages (37), were induced in WT, but not Ffar4KO cardiac myocytes (Figure 3D, Table 1). In summary, transcriptome analysis three days post-TAC suggested that cardiac myocytes from Ffar4KO hearts were unable to induce the same inflammatory and cell death responses as WT animals, specifically genes associated with cPLA_2_α; signaling, likely contributing to the more severe remodeling in the Ffar4KO post-TAC.

**Figure 3.**
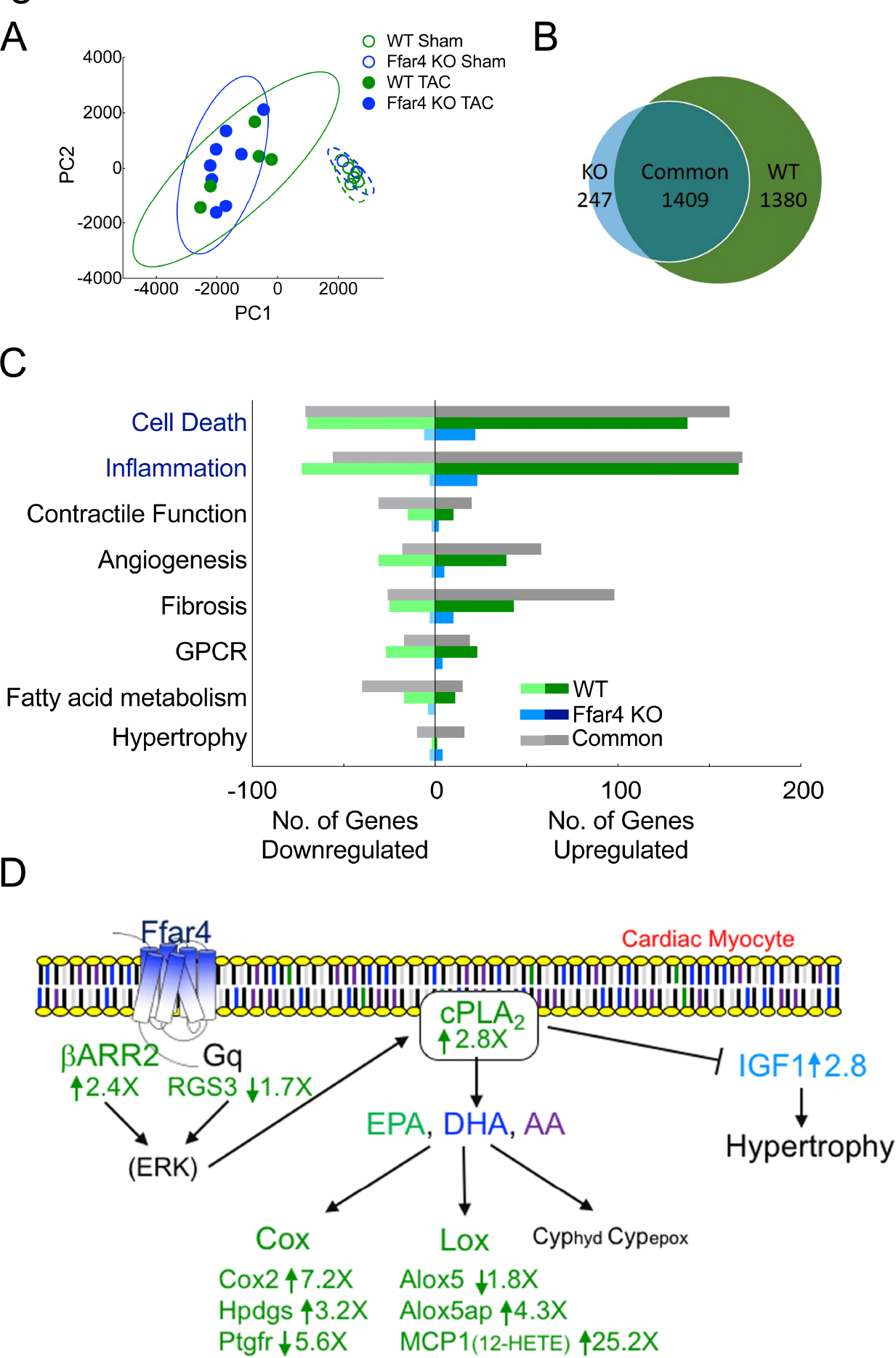
Ffar4 is necessary for induction of cardioprotective inflammatory and cell death genes, and specifically expression of genes associated with cPLA_2_ signaling. Transcriptome analysis (RNA-seq) was performed on cardiac myocytes isolated from male WT and Ffar4KO mice 3-days post-TAC or sham surgery. **A**, Principal component analysis of RNA transcriptomes from WT and Ffar4KO cardiac myocytes. **B**, Venn Diagram indicating genes changed ≥ 1.7-fold uniquely in WT cardiac myocytes post-TAC (1,380 genes), changed ≥ 1.7-fold uniquely in Ffar4KO cardiac myocytes post-TAC (247 genes), or changed ≥ 1.7-fold in both (1,409). **C**, Differentially expressed genes represented in **B** were sorted based on gene ontology (GO) terms for biological function associated with eight categories (cell death, inflammation, contractile function, angiogenesis, fibrosis, GPCR, fatty acid metabolism, and hypertrophy). Graphical representation of the number of genes up- or downregulated in each category that were unique to the WT TAC (relative to sham, green), unique to Ffar4KO TAC (relative to sham, blue), and shared between WT and Ffar4KO (grey). **D**, Known Ffar4 signaling pathways with relevant differentially expressed genes with fold changes displayed. Genes shown in green are changed in WT TAC animals relative to WT sham; genes in blue are changed in Ffar4KO TAC compared to Ffar4KO sham animals.

**Table 1:** Transcriptome analysis

| GENE | GENE NAME | FOLD $\Delta$ | FUNCTION IN HEART |
| --- | --- | --- | --- |
| **CELL DEATH AND INFLAMMATION** |  |  |  |
| *Ilrn1* | IL-1β receptor antagonist | + 220 WT | Blocks IL1β-receptor signaling<br>IL1β a new target in HF (60, 61)<br>Cantos trial shows positive results post-MI with IL-1β-antibody (5) |
| *Hmox1* | Heme-oxygenase 1 | + 24 WT | Cell survival/Cardioprotective (Review: (62)) |
| *Il33* | Interleukin 33 | + 4.0 WT | Cardioprotective (Review: (62)) |
| *Ccl2* | Chemokine (C-C motif) ligand 2 | + 25 WT | Macrophage chemotractant |
| *Ccr2* | Chemokine (C-C motif) receptor 2 | + 14 WT | Ccl2 receptor |
| *Cx3cr1* | Chemokine (C-X3-C motif) receptor 1 | + 2.8 WT |  |
| **Ffar4-cPLA2 SIGNALING** |  |  |  |
| *Pla2g4a* | Phospholipase A2, group IVA (cytosolic, calcium dependent) | + 2.8 WT | Activated by Ffar4 - cleaves AA or EPA from membrane, substrates for Cox2<br>cPLA2 inhibits hypertrophy (43).<br>Cardioprotective in ischemia (44), but controversial (44) |
| *Arrb2* | β-Aresstin2 | + 2.5 WT | Ffar4 signals through β-Arr 2 (10) |
| *RGS3* | Regulator of G-protein signaling | -1.7 WT |  |
| *Alox5ap* | Arachidonate 5-lipoxygenase activating protein | + 4.3 WT | Activation of 5-lipoxygenase – enhanced production of 5-HETEs |
| *Alox5* | Arachidonate 5-lipoxygenase | - 1.8 WT | Production of 5-HETEs |
| *Ptgs2* | Cyclooxygenase 2 | + 7.2 WT | Prostaglandin synthesis, |
| *Hpgds* | Hematopoietic prostaglandin D synthase | + 3.2 WT<br>- 5.6 WT | Cardioprotective (Review: (63)) |
| *Ptgfr* | Prostaglandin F receptor |  |  |
| *Igf1* | Insulin-like growth factor 1 | + 2.8 KO | Pro-hypertrophic, related to decreased cPLA2 activity (43) |

### Ffar4 agonist TUG-891 increases production of 18-HEPE in adult cardiac myocytes

To define the function of Ffar4 in cardiac myocytes, we focused on Ffar4 mediated activation of cPLA_2_α, based on the results from our transcriptome analysis. Upon activation, cPLA_2_α can translocate to the nuclear membrane and cleave PUFAs from the sn2-acyl bond in membrane phospholipids, traditionally arachidonic acid (AA), but also EPA, DHA, or other PUFAs (38). Cleaved FAs are subsequently metabolized to produce oxylipin products with pro-inflammatory, anti-inflammatory, or pro-resolving effects. In adult cardiac myocytes treated with the Ffar4 agonist TUG-891, we used UPLC/MS/MS to detect specific AA, EPA, and DHA derived oxylipin production to assess cPLA_2_α activity (Figure 4 A-H). Here, oxylipins produced following TUG891 treatment were considered to have one of four fates: 1) acylation into cellular membranes (esterified, myocyte); 2) free in the myocyte (unesterified, myocyte); 3) export in esterified lipids (esterified, medium); or 4) exported as a free oxylipin (unesterified, medium) (Figure 4I). We identified 57 different oxylipins in our samples, with details for MRM transitions, retention times, detection parameters, and detection limits presented in Supplemental Table 13. Results for all oxylipins at 60 minutes are reported in Supplemental Table 14. Surprisingly, we found that TUG-891 selectively increased the production of the EPA-derived oxylipin 18-HEPE, likely by means of CYP_*hydrox*_ activity in cardiac myocytes (esterified 18-HEPE, Figure 4A and unesterified 4B) and exported from myocytes (esterified 18-HEPE, Figure 4C and unesterified 4D), over time (Figures 4E-H).

**Figure 4.**
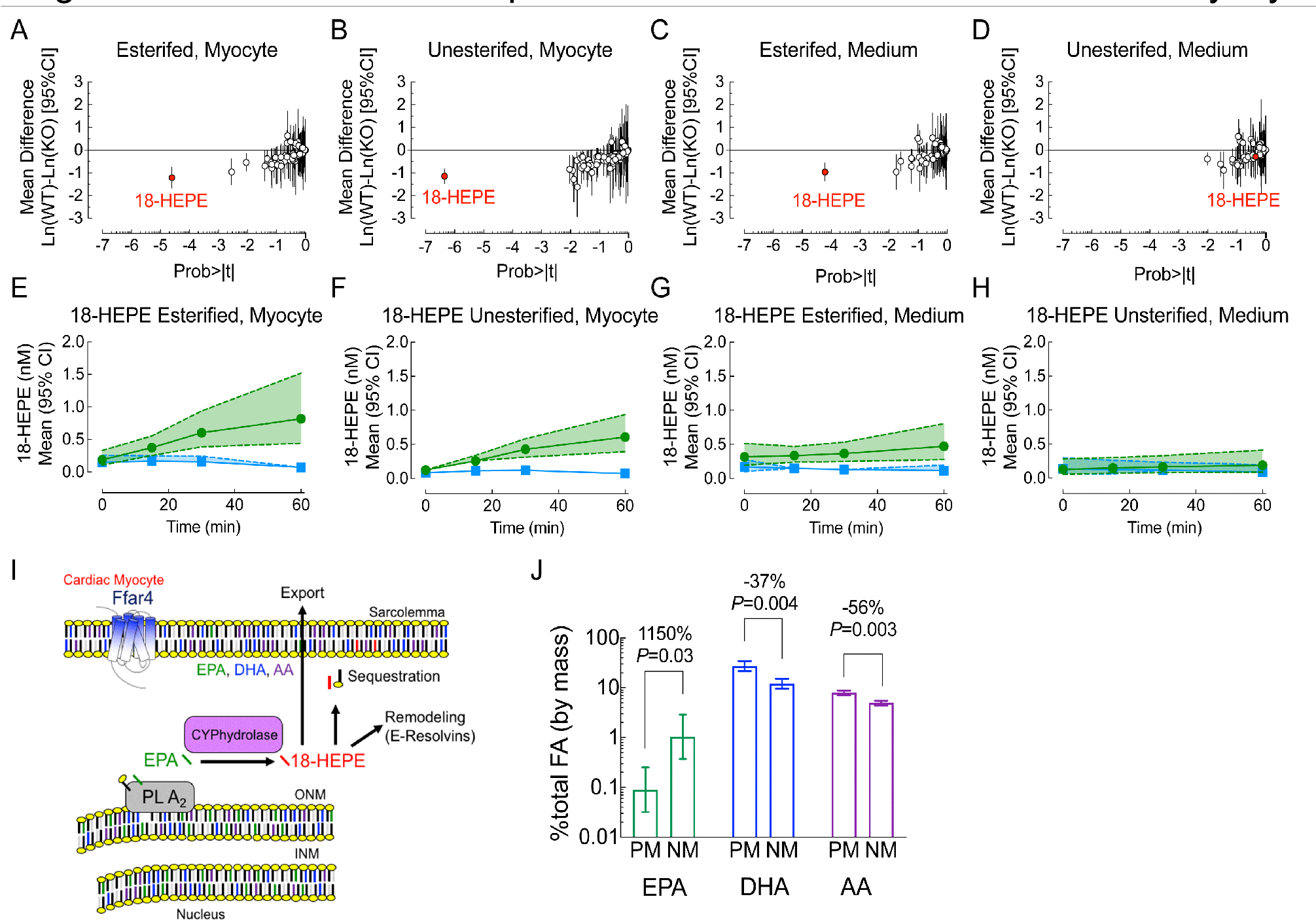
Ffar4 agonist TUG-891 increases production of 18-HEPE in adult cardiac myocytes. **A-H**, Cultured adult cardiac myocytes from WT (green) and Ffar4KO (blue) male mice were treated with the Ffar4 agonist, TUG891 (50μM) for 0, 15, 30, and 60 minutes. Oxylipins were detected by mass spectrometry from cardiac myocyte membranes (esterified) or cytosolic fractions (non-esterified, or from the culture medium in lipoproteins (esterified) or free (non-esterified). Probability plots for oxylipins detected from cardiac myocytes in the **A**, esterified or **B**, unesterified fractions after 60 minutes, or in the culture medium in the **C**, esterified or **D**, unesterifed fractions, or time course of 18-HEPE production in myocytes in the **E**, esterified or **F**, unesterifed fractions, or in the medium in the **G**, esterified or **H**, unesterified fractions (dashed lines represent the 95% CI). **I**, Cytoplasmic phospholipase A_2_α (cPLA_2_α mediated cleavage of EPA from membrane phospholipids and production of 18-HEPE by CYPhydrolase. 18-HEPE is re-acylated into membrane phospholipids (sequestration), remains free in the cell and is further metabolized (potentially E-Resolvins), or exported, esterified in a lipoprotein. **J**, EPA, DHA, and AA concentration in the nuclear membrane (NM) or sarcolemma/plasma membrane (PM). Data are Mean ± SD, n=3 separate preparations, fold differences indicated.

The Ffar4-depedent increase in 18-HEPE occurred despite AA abundance being much higher, suggesting a previously unrecognized specificity for cPLA_2_α function. One explanation for the specific increase in 18-HEPE production could be a relative enrichment of EPA at the nuclear membrane. Because cPLA_2_α can translocate to the nuclear membrane (NM) upon activation (Figure 4I), we compared the relative substrate availability in the NM to the sarcolemma/plasma membrane (PM) to determine how this affect the relative availability of PUFAs to oxylipin producing enzymes. %EPA was lower than %DHA or %AA in both PM and NM fractions, however the relative abundance of EPA was 11.5-fold greater in NM compared to PM (Figure 4J). In contrast, %DHA and %AA were lower in NM (2.3 and 1.6 fold less, respectively) compared to PM, for a total increase in the relative availability of EPA to cPLA_2_α of approximately 26-fold relative to DHA and 19-fold relative to AA.

### Loss of Ffar4 changes circulating oxylipin content consistent with a loss of anti-inflammatory oxylipins at baseline, and prevents initiation of both mediators of inflammation and of resolution following TAC

AA, EPA, and DHA oxylipins are produced intracellularly but are exported and trafficked in plasma in lipoproteins (39, 40). Therefore, we examined the consequences of disruption of Ffar4 signaling systemically on circulating oxylipins. Again, we identified 57 different oxylipins in the four plasma fractions, albumin (unesterified), HDL, LDL, and VLDL. To assess global changes in oxylipin phenotypes, we performed PCA comparing the oxylipin content in each fraction for WT and Ffar4KO, sham versus TAC and plot scores for PCA1 and PCA2 for each fraction (Figures 5A-D respectively). PCA loadings, applicable to all plots (Figure 5E), are shown with an example of the loading location of a pro-inflammatory oxylipin (12-HETE), an anti-inflammatory oxylipin (14(15)-EpETE), and a pro-resolving oxylipin (18-HEPE). Hence, a lipoprotein fraction with abundant anti-inflammatory oxylipins would load strictly rightward, a lipoprotein fraction with abundant pro-inflammatory and pro-resolving oxylipins would load downward, and hence an animal with both would load downward and to the right. Detailed loading information for all 57 oxylipins can be found in Supplemental Table 15; tests across all relevant PC scores are given in Supplemental Table 16, and tests across all individual oxylipins are given in Supplemental Table 17.

**Figure 5.**
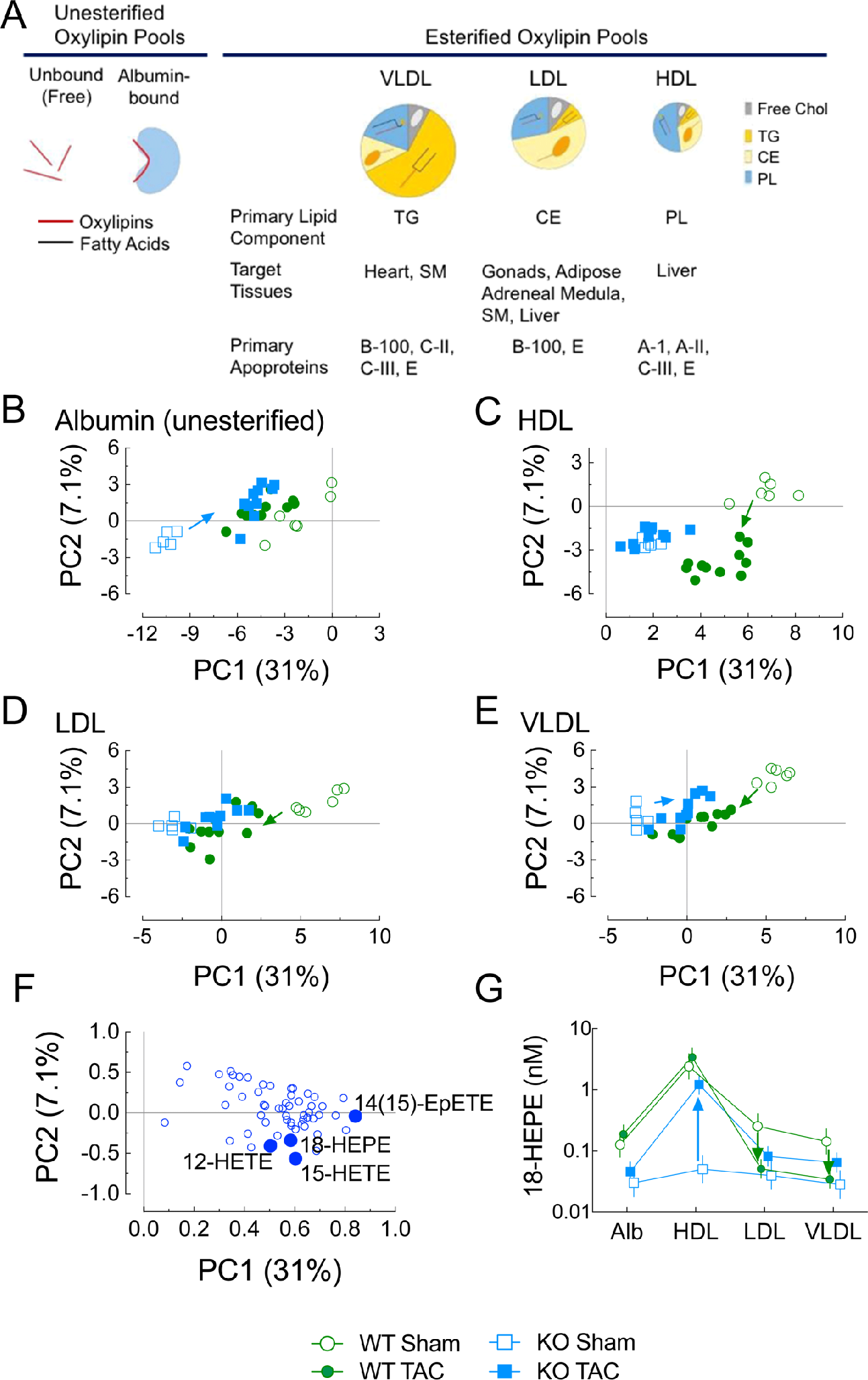
Loss of Ffar4 changes circulating oxylipin content consistent with a loss of anti-inflammatory oxylipins at baseline, and prevents initiation of both mediators of inflammation and of resolution following TAC. **A.** Description of plasma fractions containing unesterified oxylipins (Albumin fraction) and esterified oxylipins (Lipoprotein Fraction). TG, triglyceride; CE, cholesterol esters; PL, phospholipids; SM, skeletal muscle. **B-E**, Plasma was collected 4 weeks following TAC or sham surgery, and oxylipins were detected by mass spectrometry in the four plasma fractions albumin (free, unesterified fatty acids), high density lipoproteins (HDL), low-density lipoproteins (LDL) and very-low density lipoproteins (VLDL). To discriminate changes in oxylipin content between WT and Ffar4KO in sham and TAC operated mice, principle component analysis was performed for each fraction. **B**, Albumin, **C**, HDL **D**, LDL and **E**, VLDL, and **F**, PCA loadings. **G**, Specific analysis of 18-HEPE in the different plasma fractions.

All albumin fractions were located leftward, or negatively, away from all oxylipins indicating the low concentration of unesterified, freely circulating, oxylipins (Figure 5A). A small adaptive response to TAC surgery in Ffar4KO animals was evidenced by a rightward shift from the Ffar4KO sham location to co-localizing with WT sham and WT TAC.

In all lipoproteins, TAC was associated with an adaptive response in WT mice. For HDL, the oxylipin content of HDL from WT sham mice were located rightward towards 14(15)-EpETE, reflecting high levels of anti-inflammatory EPA-epoxides (Figure 5B). HDL from WT TAC mice were shifted downward and slightly left of WT sham, indicating a replacement of anti-inflammatory 14(15)-EpETE with activators of inflammation such as 12-HETE or 15-HETE, paired with production of inflammation resolving oxylipins, 18-HEPE. In contrast, HDL from Ffar4KO sham and TAC mice were both located less rightward and less downward, indicating less overall oxylipin content, less 14(15)-EpETE, and a minimal response to TAC surgery by production of 12-HETE, 15-HETE, and 18-HEPE. Therefore, in the Ffar4KO, the oxylipin profiles indicated an inability to produce oxylipins, especially anti-inflammatory signals at baseline (sham), and an inability to respond to TAC by production of either pro-inflammatory or pro-resolving oxylipins.

In a similar fashion, in WT mice, LDL oxylipin profiles indicated an adaptive response following TAC, but not in Ffar4KO mice. Relative to HDL, LDL were generally located more leftward, indicating they contained less oxylipins than HDL, consistent with our prior observations in humans (39, 41). LDL from WT sham mice (Figure 5C) were located rightward towards 14(15)-EpETE, while LDL from WT TAC mice moved leftward, away from all oxylipins, consistent with a loss of oxylipin content, co-localizing with LDL from both Ffar4KO sham and TAC mice. This suggested that the loss of Ffar4 pre-disposed towards conditions that were only present in the WT mouse following surgery.

Like LDL, VLDL were more centrally located than HDL, which is evidence of their lower overall oxylipin content relative to HDL, again consistent with prior observations in humans (39, 41). VLDL from WT sham mice (Figure 5D) were located most rightward, reflecting their high concentrations of anti-inflammatory epoxides such as 14(15)-EpETE. However, their moderately positive location upward reflected low levels of pro-inflammatory and pro-resolving oxylipins such as 12-HETE, 15-HETE, and 18-HEPE. The different location of LDL from WT TAC mice, down and to the left, indicated VLDL with less oxylipins overall, but higher levels of the pro-inflammatory and pro-resolving oxylipins. As with LDL, VLDL from Ffar4KO mice had low oxylipin regardless of their surgery status and were unable to respond to TAC with elevated pro-inflammatory/pro-resolving oxylipin content.

18-HEPE response to Ffar4 activation was reflected in plasma oxylipin pools, and most 18-HEPE was found in the HDL fraction (Figure 5F). Interestingly, 18-HEPE was dramatically lower in the Ffar4KO HDL fraction at baseline (sham), and although it increased following TAC, it never reached levels observed in the WT. In both LDL and VLDL fractions, 18-HEPE was lower in the Ffar4KO sham, and following TAC, while 18-HEPE levels decreased in the WT, 18-HEPE levels actually increased in the Ffar4KO.

### Ffar4 is expressed in the human heart and downregulated in HF, with reciprocal upregulation of Ffar1

Similar to mice, nothing is known about Ffar4 in the human heart. Therefore, we used RT-PCR to measure Ffar4 expression in samples from both healthy donors and HF patients. We found that while human hearts do express Ffar4, Ffar4 (Ffar4S + Ffar4L) expression is reduced in failing hearts, and Ffar4L represents less than 10% of total Ffar4 expression, and was not changed following TAC (Figure 6). Unexpectedly, expression of Ffar1, the only other known GPR for medium- and long-chain fatty acids, was reciprocally increased in failing hearts relative to healthy controls. This represents a significant difference from the mouse heart, which does not express detectable levels of Ffar1 (16).

**Figure 6.**
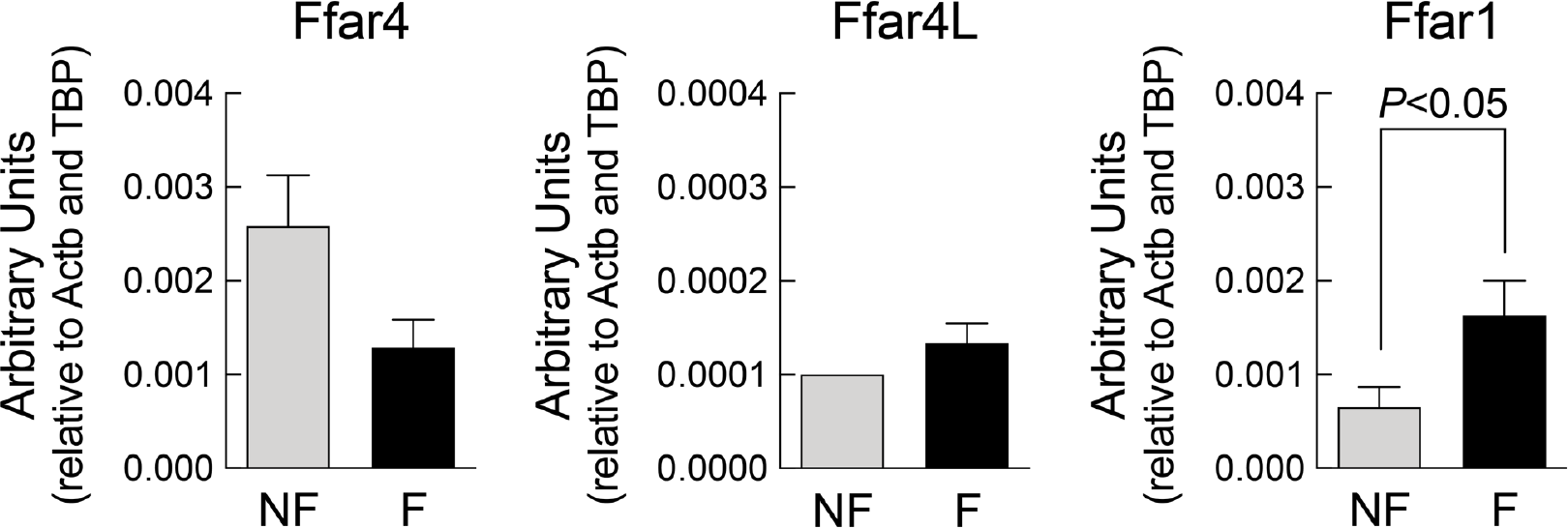
Ffar4 is expressed in the human heart and downregulated in HF, with reciprocal upregulation of Ffar1. RT-PCR to detect expression of Ffar4 (both short and long isoforms) and Ffar1 in cardiac tissue obtained from non-failing (NF) and failing (F) human hearts.

## Discussion

Here, we show for the first time that Ffar4 is cardioprotective and necessary for an adaptive response to pathologic stress in the heart. This is supported by our findings that in Ffar4KO mice, TAC induced an exaggerated hypertrophic response, although with a similar degree of fibrosis, as well as greater systolic and diastolic dysfunction, indicating worse pathologic remodeling. Furthermore, Ffar4-mediated cardioprotection occurred independent of any dietary intervention implying that Ffar4 is able to sense and respond to basal fatty acid composition to protect the heart. Mechanistically, transcriptome analysis of cardiac myocytes three days post-TAC revealed programmatic changes in gene expression associated with inflammation and cell death were lacking in Ffar4KO cardiac myocytes. More importantly, we observed specific deficits in genes associated with cPLA_2_α-mediated signaling responsible for oxylipin production in Ffar4KO cardiac myocytes. In cultured cardiac myocytes, we were surprised to find that Ffar4 preferentially increased production of one cardioprotective, EPA-derived oxylipin, 18-HEPE, despite the generally low levels of EPA detected in cardiac myocyte membranes at baseline. Furthermore, profiling oxylipin content in plasma, both esterified in circulating lipoproteins and unesterified (bound to albumin), generally indicated that loss of Ffar4 was associated with lower overall oxylipin content in all plasma fractions, and specifically less anti-inflammatory oxylipins at baseline, and inability to increase pro-inflammatory and pro-resolving oxylipins following TAC. Together, these results suggest that a key mechanism to explain the cardioprotective effects of Ffar4 is through induction of pro-resolving oxylipins in cardiac myocytes directly, and through indirect effects in balancing pro-inflammatory and pro-resolving oxylipin composition systemically, consistent with failure to activate cPLA_2_α. Additionally, we demonstrated that Ffar4 is expressed in the human heart, and downregulated in HF. In summary, our data suggest that Ffar4 functions as a nutrient sensor for medium and long chain fatty acids to resolve inflammation and maintain cardiac homeostasis.

Transcriptome analysis of cardiac myocytes performed three days post-TAC identified a deficit in cPLA_2_α signaling in Ffar4 cardiac myocytes (Figure 3) (38). There are six cPLA_2_ family members (α, β, γ, δ, ε, and ξ) which show only roughly 30% sequence homology, but have different enzymatic properties, tissue expression, and subcellular localizations (38). cPLA2α is widely expressed and has biologic functions in a multitude of cell types (38) and upon activation, it localizes to the nuclear membrane (42). Early studies suggested that cPLA_2_α functions as a negative regulator of striated muscle growth. In mice with systemic deletion of cPLA_2_α (cPLA_2_KO mice), normal developmental growth of skeletal and cardiac muscle, as well as pathologic cardiac hypertrophic growth post-TAC are exaggerated (43). Mechanistically, loss of cPLA_2_α induced sustained activation of IGF-1 signaling, a known hypertrophic stimulus, in cardiac myocytes from cPLA_2_KO hearts (43). In addition, cPLA_2_α might protect against ischemic injury in the heart, although there are conflicting reports regarding cPLA_2_α-mediated cardioprotection (44, 45). Our results demonstrate that cPLA_2_α (*Pla2g4a*) was upregulated 2.8 fold only in WT cardiac myocytes, whereas IGF-1 (*Igf1*) was upregulated 2.8 fold only in Ffar4KO cardiac myocytes, consistent with the exaggerated hypertrophic response post-TAC in hearts from Ffar4KO mice. Furthermore, the deficit in cPLA_2_α mediated signaling in Ffar4KO cardiac myocytes was associated with worse remodeling following TAC, suggesting Ffar4-cPLA_2_α signaling is cardioprotective.

Functionally, cPLA_2_α (*Pla2g4a*) cleaves PUFAs from the sn2-acyl bond in membrane phospholipids. AA is the most common substrate, but DHA and EPA are substrates, leading to the production of downstream oxylipins (38). Initially, cleaved FAs (AA, EPA, DHA) are metabolized by cyclooxygenases (COX1/2), lipoxygenases (5-LOX, 12/15-LOX), or CYP_*hydroxylase*_ and CYP_*epoxygenase*_ to generate oxygenated signaling mediators, including leukotrienes and prostaglandins, which are collectively known as oxylipins, many of which mediate pro-or anti-inflammatory responses, or initiate resolution of inflammation (46). Here, we found that in cardiac myocytes, the Ffar4 agonist TUG-891 activated cPLA_2_α and induced the production of a specific EPA-derived oxylipin, 18-HEPE (Figure 4, Supplemental Tables 14). Although, the specific source of 18-HEPE is unknown, it is likely a product of CYP_*hydroxylase*_ activity (47). Our data imply a role for Cyp26b1 which is previously identified for its role in hydroxylating retinoic acid (48). While this is the first such demonstration in cardiac myocytes, one previous study in macrophages indicated that DHA signaling through Ffar4 activates cPLA_2_α mediated oxylipin production (37). Interestingly, we found EPA levels in red blood cells were low, 0.1-0.2% of total membrane FAs, whereas DHA was 4% and AA was 13% (Supplemental Tables 2A and 2B), similar to our previous studies (16). Furthermore, we previously found that in cardiac myocytes, EPA was 0.5% of total membrane FAs, whereas DHA was 10% and AA was about 12% (16). Therefore, in cardiac myocytes, Ffar4 shows surprising degree of specificity for the production of 18-HEPE, largely to the exclusion of other EPA, DHA, or AA derived oxylipins despite the low levels of EPA in cardiac myocyte membranes. Potentially, the localization of cPLA_2_α to an EPA rich membrane enhances its capacity to mediate the specific production of EPA-derived oxylipins such as 18-HEPE.

While there are no previous studies regarding the direct effects of 18-HEPE in cardiac myocytes, macrophages from fat-1 mice, which have high endogenous levels of ω3-PUFAs, particularly EPA, due to overexpression the *c. elegans* fat-1 gene, produced high levels of 18-HEPE that were associated with cardioprotection in fat-1 mice post-TAC (49). Furthermore, direct i.p. injection of 18-HEPE attenuated remodeling post-TAC (49). Interestingly, 18-HEPE is also the precursor for E-series resolvins (RvE1, RvE2, and RvE3), a class of pro-resolving oxylipins (50). E-resolvins signal through ERV1/ChemR23, an orphan GPR that binds to and is activated by both E-resolvins and the endogenous peptide chemerin, suggesting a potential mechanism to explain EPA-mediated inflammation resolving effects (51). ERV1/ChemR23 is expressed in the heart, and preconditioning with RvE1 reduces ischemia/reperfusion injury (52), while RvE1 infusion one-week following coronary artery ligation attenuates post-MI inflammatory response (53). In our study, the production of 18-HEPE co-occurred systemically with the production of pro-inflammatory products, 12-HETE and 15-HETE. Combined with our data, these studies suggest that Ffar4-cPLA_2_α mediated production of 18-HEPE is cardioprotective by providing the capacity to resolve inflammation, either through direct effects in the heart, released from infiltrating macrophages or directly produced in cardiac myocytes, or indirect effects via production of E-resolvins and activation of ERV1/ChemR23.

Two important deficits were observed in circulating oxylipin profiles in Ffar4KO mice: first, Ffar4KO mice lacked the capacity to induce any cPLA_2_α mediated oxylipin response, and second, they lacked the capacity to modify oxylipin production in response to TAC. Surprisingly, this did not mean suppressing the activation of inflammation, but instead it meant co-producing mediators essential for the resolution of inflammation. Systemically, WT mice at baseline (sham) had the highest oxylipin content, with high levels of anti-inflammatory oxylipins such as 14(15)-EpETE. Interventions that increase the production of EpETEs, or slow their inactivation, attenuate TAC remodeling (54, 55), but we observed that TAC altered the HDL oxylipin content to include a pro-inflammatory response (12-HETE, 15-HETE). In mice, 12/15-LOX is the common source for both 12- and 15-HETE, and overexpression of 12/15-LOX is associated with increased levels of unesterified 12-HETE, 15-HETE, and heart failure (56). We found that despite increase in this pro-inflammatory insult (increased 12-HETE, 15-HETE), WT mice were protected through their capacity to co-produce the pro-resolving 18-HEPE, which was lacking in Ffar4KO mice. This suggests that the capacity to produce 12/15-LOX metabolites is beneficial, provided they can also initiate recovery.

Clinically, ω3-PUFAs are important to cardiovascular health (57, 58), and there is substantial evidence demonstrating a benefit of ω3-PUFAs in coronary heart disease (CHD) and HF. In one large clinical trial, Gruppo Italiano per lo Studio della Sopravvivenza nell’Infarto miocardico-Heart Failure (GISSI-HF) (27), and several smaller trials, ω3-PUFAs improved HF outcomes (28–30). In animal models, we have also demonstrated that EPA, in a concentration-dependent manner, prevents pathologic remodeling post-TAC (16, 31, 59). However, important questions remain regarding the mechanism underlying the benefit of ω3-supplementation in HF. Here, we found that Ffar4, a receptor for medium and long-chain fatty acids including, but not limited to, ω3-PUFAs, is cardioprotective, but whether Ffar4 is required for ω3-mediated protection is unclear at this time. Mechanistically, we found that in cardiac myocytes Ffar4 increased 18-HEPE levels, which might be directly cardioprotective or act through its downstream metabolites E-resolvins. Ultimately, this could suggest a feed-forward mechanism with multiple points for EPA mediated cardioprotection: 1. Through Ffar4-mediated production of 18-HEPE, either in cardiac myocytes (Figure 4) or potentially macrophages (37). 2. Increasing overall EPA levels, thereby increasing 18-HEPE levels, as suggested by others (49), or 3. Increased E-resolvin levels and activation of ERV1/ChemR23 either via Ffar mediated production of 18-HEPE or increased 18-HEPE levels secondary to an overall increase in EPA levels.

In conclusion, our results identify Ffar4 as a nutrient sensor for medium and long chain FAs that attenuates inflammation to maintain cardiac homeostasis. Finally, our findings suggest a novel paradigm for FAs in cardiac myocytes, whereby FA function not simply as an energy source, but as signaling molecules that activate cardioprotective GPR signaling.

## Acknowledgements

This work was supported by NIH HL130099 (TDO and GCS), Minnesota Obesity Prevention Training Program T32 NIH Grant 1T32DK083250–01A1 (KM), and a grant from Amarin Corporation. The authors also acknowledge the Genomics Core at the University of Minnesota for technical expertise on all data acquisition and analysis for transcriptome phenotypes.

## Abbreviations

12-HETE: 12-hydroxyeicosatrienoic acid
14(15)-EpETE: 14(15)-epoxytetraenoic acid
15-HETE: 15-hydroxyeicosatrienoic acid
18-HEPE: 18-hydroxyeicosapentaenoic acid
AA: arachidonic acid
AMVM: adult mouse ventricular myocytes
ATM: adipose tissue macrophage
βArr2: βarrestin-2
CANTOS: Canakinumab Anti-inflammatory Thrombosis Outcome Study
cPLA_2_α: cytoplasmic phospholipase A_2_α
COX: cyclooxygenase
CYP: cytochrome p450
DHA: docosahexaenoic acid
EPA: eicosapentaenoic acid
FA: fatty acid
Ffar: free fatty acid receptor
Ffar1: free fatty acid receptor 1
Ffar4: free fatty acid receptor 4
Ffar4KO: free fatty acid receptor 4 knock out mouse
Ffar4S: free fatty acid receptor 4 short isoform
Ffar4L: free fatty acid receptor 4 long isoform
FPLC: fast performance liquid chromatography
GISSI-HF: Gruppo Italiano per lo Studio della Sopravvivenza nell’Infarto miocardico-Heart Failure
GO: gene ontology
GPR: G-coupled protein receptor
GPR120: G-protein coupled receptor 120
HF: heart failure
HFrEF: heart failure reduced ejection fraction
HFpEF: heart failure preserved ejection fraction
HDL: high density lipoproteins
IL-1β: interleukin-1β
iPLA_2_: calcium-independent PLA_2_
KO: knock outTAC, transverse aortic constriction
KOMP: Knock Out Mouse Project
LDL: low density lipoproteins
LOX: lipoxygenase
MUFA: monounsaturated fatty acid
NM: nuclear membrane
PAF-AH: platelet-activating acetylhydrolases
PCA: principal component analysis
PM: plasma membrane
PUFA: polyunsaturated fatty acid
SFA: saturated fatty acid
TAC: transverse aortic constriction
TGFβ-1: transforming growth factor beta-1WT, wild type
VLDL: very low density lipoproteins
WT: wild type

