## Supplemental Tables 1-17 for "Free fatty acid receptor 4 is a nutrient sensor that resolves inflammation to maintain cardiac homeostasis"

### Supplemental Tables and Figures

**Supplemental Table 1.** DYETS, Inc #180539 Chow Composition Table

| Ingredient | kcal/gm | grams/kg | kcal/kg |
|---|---|---|---|
| Casein | 3.58 | 140 | 501 |
| Sucrose | 4 | 100 | 400 |
| Cornstarch | 3.6 | 466 | 1676 |
| Dyetrose | 3.8 | 155 | 589 |
| L-Cystine | 4 | 1.8 | 7.2 |
| Cellulose | 0 | 50 | 0 |
| Corn Oil | 9 | 40 | 360 |
| t-Butylhydroquinone | 0 | 0.008 | 0 |
| Mineral Mix #210050 | 0.84 | 35 | 29.4 |
| Vitamin Mix #310025 | 3.87 | 10 | 38.7 |
| Choline Bitartrate | 0 | 2.5 | 0 |
| TOTALS | | **1000.00** | **3601** |

**Supplemental Table 2A.** Male fatty acid levels 4 weeks post-TAC measured by mass spectrometry.

| Fatty Acid (abbreviation) | WT Sham (12) | KO Sham (11) | WT TAC (24) | KO TAC (20) |
|---|---|---|---|---|
| C22:6n3 (DHA) | 3.56 ± 0.76 | 3.59 ± 0.93 | 3.76 ± 0.74 | 3.71 ± 0.92 |
| C22:5n3 (n3DPA) | 0.27 ± 0.04 | 0.29 ± 0.05 | 0.31 ± 0.16 | 0.25 ± 0.08 |
| C22:5n6 (n6DPA) | 2.38 ± 0.70 | 1.92 ± 0.39 | 2.31 ± 0.57 | 2.21 ± 0.66 |
| C22:4n6 (n6DTA) | 2.17 ± 0.39 | 1.98 ± 0.43 | 2.10 ± 0.58 | 2.06 ± 0.63 |
| C20:5n3 (EPA) | 0.12 ± 0.07 | 0.13 ± 0.11 | 0.24 ± 0.25 | 0.13 ± 0.08* |
| C20:4n6 (AA) | 12.91 ± 2.00 | 12.94 ± 2.91 | 12.40 ± 2.46 | 12.74 ± 2.94 |
| C20:3n6 (dgLA) | 1.42 ± 0.28 | 1.40 ± 0.39 | 1.49 ± 0.32 | 1.42 ± 0.43 |
| C20:2n6 (EDA) | 0.24 ± 0.05 | 0.21 ± 0.08 | 0.32 ± 0.21 | 0.20 ± 0.09 |
| C18:3n3 (aLA) | 0.12 ± 0.09 | 0.13 ± 0.11 | 0.14 ± 0.17 | 0.16 ± 0.09 |
| C18:3n6 (gLA) | 0.07 ± 0.06 | 0.10 ± 0.13 | 0.16 ± 0.38 | 0.14 ± 0.22 |
| C18:2n6 (LA) | 11.96 ± 1.11 | 11.26 ± 1.98 | 11.47 ± 1.63 | 11.59 ± 2.58 |
| C20:1n9 (EA) | 0.41 ± 0.14 | 0.33 ± 0.14 | 0.50 ± 0.14 | 0.39 ± 0.21 |
| C24:0 (LgA) | 0.22 ± 0.16 | 0.26 ± 0.15 | 0.33 ± 0.32 | 0.24 ± 0.16 |
| C20:0 (ArcA) | 0.18 ± 0.08 | 0.27 ± 0.26 | 0.32 ± 0.35 | 0.26 ± 0.19 |
| C18:1n7 (VA) | 2.04 ± 0.44 | 1.89 ± 0.33 | 2.36 ± 0.35 | 2.23 ± 0.59 |
| C18:1n9 (OA) | 12.55 ± 1.00 | 12.54 ± 1.22 | 13.21 ± 1.58 | 12.33 ± 1.69 |
| C18:1n12 | 0.10 ± 0.05 | 0.10 ± 0.96 | 0.13 ± 0.12 | 0.14 ± 0.11 |
| C18:1t (ElA) | 0.24 ± 0.22 | 0.26 ± 0.27 | 0.23 ± 0.24 | 0.30 ± 0.27 |
| C18:0 (SA) | 14.58 ± 1.41 | 14.10 ± 1.66 | 13.68 ± 1.43 | 13.78 ± 1.89 |
| C17:0 | 0.31 ± 0.05 | 0.51 ± 0.58 | 0.53 ± 0.78 | 0.52 ± 0.72 |
| C16:1n7 (POA) | 0.79 ± 0.25 | 0.95 ± 0.28 | 0.94 ± 0.25 | 1.09 ± 0.51 |
| C16:0 (PA) | 32.45 ± 2.26 | 33.69 ± 2.30 | 32.04 ± 1.88 | 33.13 ± 1.84* |
| C15:0 | 0.37 ± 0.49 | 0.24 ± 0.23 | 0.23 ± 0.17 | 0.27 ± 0.20 |
| C14:0 (MA) | 0.32 ± 0.16 | 0.39 ± 0.41 | 0.39 ± 0.32 | 0.35 ± 0.22 |
| C12:0 (LA) | 0.24 ± 0.28 | 0.50 ± 1.08 | 0.43 ± 1.06 | 0.62 ± 1.48 |

Data are presented as mean ± standard deviation for the number of mice indicated in parentheses. DHA, docosahexaenoic acid; n3DPA, n-3 docosapentaenoic acid (clupanodionic acid); n6DPA, n-6 docosapentaenoic acid (osbond acid); n6DTA, adrenic acid; EPA, eicosapentaenoic acid; AA, arachidonic acid; dgLA, dihomo-gamma-linolenic acid; EDA, eicosadienoic acid; aLA, alpha-linolenic acid; gLA, gamma-linolenic acid; LA, linoleic acid; EA, eicosenoic acid; LgA, lignoceric acid; ArcA, arachidic acid; VA, cis-vaccenic acid; OA, oleic acid; ElA, elaidic acid; SA, stearic acid; POA, palmitoleic acid; PA, palmitic acid; MA, myristic acid; LA, lauric acid. * Indicates $P$<0.05 vs WT TAC.

**Supplemental Table 2B**. Female fatty acid levels 4 weeks post-TAC measured by mass spectrometry.

| **Fatty Acid (abbreviation)** | **WT Sham (12)** | **KO Sham (12)** | **WT TAC (31)** | **KO TAC (25)** |
|---|---|---|---|---|
| C22:6n3 (DHA) | 4.59 ± 0.54 | 4.69 ± 0.50 | 4.19 ± 0.36 | 4.21 ± 0.89 |
| C22:5n3 (n3DPA) | 0.21 ± 0.05 | 0.22 ± 0.06 | 0.24 ± 0.07 | 0.26 ± 0.20 |
| C22:5n6 (n6DPA) | 2.34 ± 0.33 | 2.02 ± 0.45 | 2.46 ± 0.45 | 2.15 ± 0.54* |
| C22:4n6 (n6DTA) | 2.56 ± 0.22 | 2.33 ± 0.48 | 2.42 ± 0.25 | 2.22 ± 0.49 |
| C20:5n3 (EPA) | 0.17 ± 0.14 | 0.10 ± 0.05 | 0.21 ± 0.15 | 0.13 ± 0.19 |
| C20:4n6 (AA) | 14.42 ± 1.60 | 14.73 ± 1.95 | 13.90 ± 1.41 | 13.87 ± 2.58 |
| C20:3n6 (dgLA) | 1.30 ± 0.20 | 1.21 ± 0.29 | 1.32 ± 0.19 | 1.16 ± 0.33* |
| C20:2n6 (EDA) | 0.22 ± 0.07 | 0.16 ± 0.07 | 0.27 ± 0.07 | 0.21 ± 0.15 |
| C18:3n3 (aLA) | 0.10 ± 0.14 | 0.06 ± 0.04 | 0.08 ± 0.06 | 0.11 ± 0.19 |
| C18:3n6 (gLA) | 0.05 ± 0.03 | 0.03 ± 0.03 | 0.06 ± 0.06 | 0.07 ± 0.08 |
| C18:2n6 (LA) | 10.92 ± 1.34 | 10.56 ± 1.16 | 10.37 ± 0.96 | 10.37 ± 1.69 |
| C20:1n9 (EA) | 0.29 ± 0.11 | 0.18 ± 0.13 | 0.41 ± 0.09 | 0.23 ± 0.14* |
| C24:0 (LgA) | 0.25 ± 0.12 | 0.20 ± 0.11 | 0.31 ± 0.19 | 0.27 ± 0.24 |
| C20:0 (ArcA) | 0.17 ± 0.09 | 0.12 ± 0.05 | 0.22 ± 0.12 | 0.17 ± 0.27 |
| C18:1n7 (VA) | 1.83 ± 0.28 | 1.40 ± 0.36 | 2.14 ± 0.32 | 1.86 ± 0.48* |
| C18:1n9 (OA) | 13.42 ± 1.74 | 13.07 ± 0.62 | 14.12 ± 0.80 | 13.22 ± 1.92* |
| C18:1n12 | 0.09 ± 0.05 | 0.08 ± 0.03 | 0.12 ± 0.08 | 0.18 ± 0.35 |
| C18:1t (EIA) | 0.16 ± 0.12 | 0.14 ± 0.09 | 0.20 ± 0.14 | 0.23 ± 0.30 |
| C18:0 (SA) | 15.52 ± 1.36 | 15.68 ±1.51 | 15.39 ± 1.63 | 15.60 ± 2.49 |
| C17:0 | 0.25 ± 0.05 | 0.25 ± 0.06 | 0.36 ± 0.21 | 0.29 ± 0.16 |
| C16:1n7 (POA) | 0.71 ± 0.20 | 0.50 ± 0.13 | 0.83 ± 0.46 | 0.65 ± 0.22 |
| C16:0 (PA) | 29.88 ± 1.70 | 31.84 ± 1.67 | 29.62 ± 1.78 | 31.73 ± 1.81* |
| C15:0 | 0.20 ± 0.20 | 0.12 ± 0.04 | 0.22 ± 0.18 | 0.19 ± 0.22 |
| C14:0 (MA) | 0.30 ± 0.10 | 0.22 ± 0.05 | 0.38 ± 0.28 | 0.39 ± 0.45 |
| C12:0 (LA) | 0.12 ± 0.11 | 0.10 ± 0.04 | 0.18 ± 0.16 | 0.28 ± 0.58 |

Data are presented as mean ± standard deviation for the number of mice indicated in parentheses. DHA, docosahexaenoic acid; n3DPA, n-3 docosapentaenoic acid (clupanodionic acid); n6DPA, n-6 docosapentaenoic acid (osbond acid); n6DTA, adrenic acid; EPA, eicosapentaenoic acid; AA, arachidonic acid; dgLA, dihomo-gamma-linolenic acid; EDA, eicosadienoic acid; aLA, alpha-linolenic acid; gLA, gamma-linolenic acid; LA, linoleic acid; EA, eicosenoic acid; LgA, lignoceric acid; ArcA, arachidic acid; VA, cis-vaccenic acid; OA, oleic acid; EIA, elaidic acid; SA, stearic acid; POA, palmitoleic acid; PA, palmitic acid; MA, myristic acid; LA, lauric acid. * Indicates $P<0.05$ vs WT TA

**Supplemental Table 3A. Male** morphological parameters 4 weeks post-TAC.

| MORPHOLOGICAL | WT SHAM (12) | KO SHAM (11) | WT TAC (24) | KO TAC (20) |
|---|---|---|---|---|
| HW (mg) | 142.3 ± 21.1 | 137.2 ± 12.8 | 196.6 ± 42.1 | 228.9 ± 45.8* |
| BW (g) | 27.3 ± 2.2 | 27.1 ± 1.1 | 26.4 ± 1.3 | 26.3 ± 2.1 |
| HW/BW (mg/g) | 5.2 ± 0.7 | 5.1 ± 0.5 | 7.45 ± 1.6 | 8.2 ± 2.3* |
| LW (mg) | 175.6 ± 13.1 (11) | 175.0 ± 18.4 | 224.6 ± 101.8 (21) | 293.0 ± 147.0 |
| Fibrosis (%) | 4.3 ± 1.7 | 3.3 ± 0.9 | 8.8 ± 6.1 | 8.7 ± 5.4 |

Data are presented as Mean ± standard deviation for the number of mice indicated in parentheses.
HR, heart rate; BW, body weight; LW, lung weight. * Indicates *P*<0.05 vs WT TAC

**Supplemental Table 3B. Male** cardiac function 4 weeks post-TAC measured by echocardiography.

| CARDIAC FUNCTION | WT SHAM (12) | KO SHAM (11) | WT TAC (24) | KO TAC (20) |
|---|---|---|---|---|
| HR (bpm) | 430 ± 52 | 423 ± 30 | 465 ± 67 | 473 ± 41 |
| SV (μl) | 28.5 ± 4.4 | 29.6 ± 4.4 | 23.8 ± 7.9 | 21.2 ± 9.6 |
| EF (%) | 43.4 ± 7.0 | 44.6 ± 6.0 | 35.7 ± 13.6 | 27.6 ± 12.2* |
| FS (%) | 21.2 ± 4.1 | 21.8 ± 3.4 | 17.2 ± 7.3 | 12.9 ± 6.3* |
| EDV (μl) | 66.3 ± 7.9 | 67.1 ± 10.2 | 70.4 ± 16.5 | 78.8 ± 17.0 |
| ESV (μl) | 37.7 ± 7.6 | 37.5 ± 8.3 | 46.6 ± 19.7 | 57.6 ± 18.1 |
| CO (ml/min) | 12.4 ± 2.8 | 12.5 ± 2.1 | 10.8 ± 3.4 | 10.0 ± 4.8 |
| IVS;s (mm) | 1.0 ± 0.2 | 1.0 ± 0.2 | 1.3 ± 0.2 | 1.3 ± 0.2 |
| IVS;d (mm) | 0.8 ± 0.1 | 0.8 ± 0.2 | 1.0 ± 0.2 | 1.1 ± 0.2 |
| LVID;s (mm) | 3.1 ± 0.2 | 3.1 ± 0.3 | 3.3 ± 0.6 | 3.7 ± 0.5* |
| LVID;d (mm) | 3.9 ± 0.2 | 3.9 ± 0.3 | 4.0 ± 0.4 | 4.2 ± 0.4* |
| LVPW;s (mm) | 1.1 ± 0.1 | 1.0 ± 0.1 | 1.2 ± 0.2 | 1.3 ± 0.2 |
| LVPW;d (mm) | 0.8 ± 0.1 | 0.8 ± 0.1 | 1.0 ± 0.2 | 1.1 ± 0.2 |
| E/A | 1.6 ± 0.2 (10) | 1.95 ± 0.4 | 1.95 ± 0.7 (15) | 3.23 ± 0.4 (14)* |
| E (mm/s) | 496.2 ± 77.8 (11) | 507.3 ± 75.8 | 647.8 ± 140.0 | 627.2 ± 137.5 |
| E/E’ | 27.2 ± 4.2 (5) | 27.7 ± 6.4 (4) | 39.5 ± 12.2 (8) | 47.0 ± 18.3 (5) |
| AOV (mm/s) | 798 ± 130 | 750 ± 111 | 3298 ± 638 | 3429 ± 660 |

Data are presented as Mean ± standard deviation for the number of mice indicated in parentheses. HR, heart rate; SV, stroke volume; EF, ejection fraction; FS, fractional shortening; EDV, end diastolic volume; ESV, end systolic volume; CO, cardiac output; IVS;s, interventricular septal thickness at systole; IVS;d, interventricular septal thickness at diastole; LVID;s, left ventricular internal diameter systole; LVID;d, left ventricular internal diameter diastole; LVPW;s, left ventricular posterior wall systole; LVPW;d, left ventricular posterior wall diastole; E/A, early mitral valve filling velocity/ late mitral valve filling velocity; E, early mitral valve filling velocity ; E/E’, early mitral valve filling velocity/early mitral annular tissue velocity; AoV, peak aortic velocity. * Indicates *P*<0.05 vs WT TAC.

**Supplemental Table 4A. Female** morphological parameters 4 weeks post-TAC.

| MORPHOLOGICAL | WT SHAM (13) | KO SHAM (12) | WT TAC (31) | KO TAC (25) |
|---|---|---|---|---|
| HW (mg) | 114.7 ± 14.0 | 120.2 ± 10.8 | 181.0 ± 50.1 | 198.0 ± 46.9 |
| BW (g) | 21.3 ± 1.4 | 21.6 ± 1.4 | 21.4 ± 1.6 | 21.7 ± 2.3 |
| HW/BW (mg/g) | 5.4 ± 0.5 | 5.6 ± 0.4 | 8.5 ± 2.5 | 9.3 ± 3.0 |
| LW (mg) | 165.5 ± 16.5 (12) | 163.2 ± 12.3 | 265.0 ± 150.4 (26) | 311.1 ± 167.8 |
| Fibrosis (%) | 4.8 ± 1.6 | 4.1 ± 1.8 | 9.2 ± 6.1 | 6.9 ± 4.6 |

Data are presented as Mean ± standard deviation for the number of mice indicated in parentheses.
HR, heart rate; BW, body weight; LW, lung weight. * Indicates *P*<0.05 vs WT TAC

**Supplemental Table 4B. Female** cardiac function 4 weeks post-TAC measured by echocardiography.

| CARDIAC FUNCTION | WT SHAM (13) | KO SHAM (12) | WT TAC (31) | KO TAC (25) |
|---|---|---|---|---|
| HR (bpm) | 431 ± 64 | 431 ± 39 | 474 ± 70 | 486 ± 49 |
| SV (μl) | 25.2 ± 6.1 | 25.4 ± 6.1 | 19.9 ± 8.2 | 21.0 ± 6.4 |
| EF (%) | 49.8 ± 11.9 | 44.4 ± 11.1 | 34.3 ± 13.6 | 30.6 ± 11.8 |
| FS (%) | 25.1 ± 7.9 | 21.8 ± 6.4 | 16.3 ± 7.1 | 14.4 ± 6.0 |
| EDV (μl) | 51.5 ± 9.2 | 58.2 ± 8.4 | 60.4 ± 12.7 | 72.6 ± 17.5* |
| ESV (μl) | 26.3 ± 8.2 | 32.8 ± 9.8 | 40.5 ± 15.6 | 51.6 ± 18.4* |
| CO (ml/min) | 10.9 ± 3.3 | 10.9 ± 2.7 | 9.5 ± 4.3 | 10.2 ± 3.3 |
| IVS;s (mm) | 1.0 ± 0.2 | 1.0 ± 0.3 | 1.2 ± 0.2 | 1.2 ± 0.2 |
| IVS;d (mm) | 0.8 ± 0.1 | 0.8 ± 0.1 | 1.0 ± 0.2 | 1.0 ± 0.2 |
| LVID;s (mm) | 2.7 ± 0.4 | 2.9 ± 0.4 | 3.2 ± 0.5 | 3.5 ± 0.5* |
| LVID;d (mm) | 3.5 ± 0.3 | 3.7 ± 0.2 | 3.8 ± 0.3 | 4.0 ± 0.4* |
| LVPW;s (mm) | 1.1 ± 0.2 | 1.0 ± 0.1 | 1.2 ± 0.2 | 1.2 ± 0.2 |
| LVPW;d (mm) | 0.8 ± 0.1 | 0.7 ± 0.1 | 1.1 ± 0.2 | 1.0 ± 0.2 |
| E/A | 1.8 ± 0.6 (11) | 2.1 ± 0.6 (11) | 3.2 ± 1.5 (14) | 2.6 ± 1.2 (9) |
| E (mm/s) | 473 ± 153 (12) | 473 ± 120 | 687 ± 159 (26) | 638 ± 193 |
| E/E' | 23.7 ± 3.1 (3) | 29.0 ± 10.0 (2) | 63.9 ± 30.8 (10) | 67.3 ± 30.5 (8) |
| AOV (mm/s) | 736 ± 129 | 641 ± 108 | 3071 ± 683 | 3236 ± 491 |

Data are presented as Mean ± standard deviation for the number of mice indicated in parentheses. HR, heart rate; SV, stroke volume; EF, ejection fraction; FS, fractional shortening; EDV, end diastolic volume; ESV, end systolic volume; CO, cardiac output; IVS;s, interventricular septal thickness at systole; IVS;d, interventricular septal thickness at diastole; LVID;s, left ventricular internal diameter systole; LVID;d, left ventricular internal diameter diastole; LVPW;s, left ventricular posterior wall systole; LVPW;d, left ventricular posterior wall diastole;E/A, E/A, early mitral valve filling velocity/ late mitral valve filling velocity; E, early mitral valve filling velocity ; E/E', early mitral valve filling velocity/early mitral annular tissue velocity; AoV, peak aortic velocity. * Indicates *P*<0.05 vs WT TAC.

**Supplemental Table 5. Cell Death Genes**

GO Terms: Apoptosis, apoptotic, necrosis, necrotic, cell death

Genes Unique to FFAR4 KO Sham vs FFAR4 KO TAC: 22 increased, 6 decreased

| Gene ID | Fold Change |
|---|---|
| Atf3 | 3.406 |
| Cx3cl1 | 2.987 |
| Igf1 | 2.787 |
| Dcun1d3 | 2.125 |
| Csrnp1 | 2.035 |
| Cyba | 2.019 |
| Ero1l | 1.889 |
| Errfi1 | 1.882 |
| Perp | 1.867 |
| G2e3 | 1.834 |
| Ticam1 | 1.824 |
| Mfge8 | 1.814 |
| Nqo1 | 1.799 |
| Ercc6 | 1.776 |
| Trim27 | 1.769 |
| Bcl2 | 1.768 |
| Etv6 | 1.750 |
| Gclc | 1.747 |
| Pnp | 1.740 |
| Ruvbl1 | 1.738 |
| Ace | 1.732 |
| Sqstm1 | 1.706 |
| Ndufc2 | -1.708 |
| Atg4d | -1.759 |
| Fastk | -1.790 |
| Ar | -1.853 |
| Zfp346 | -2.280 |
| Bmp10 | -8.321 |

Genes Unique to WT Sham vs WT TAC: 138 increased, 70 decreased

| Gene ID | Fold Change |
|---|---|
| Il1rn | 219.636 |
| Adam8 | 117.349 |
| Timp1 | 112.492 |
| Spp1 | 85.235 |
| Krt18 | 51.556 |
| Ankrd2 | 47.887 |
| Lrp8 | 37.091 |
| Pak3 | 32.821 |
| Sfn | 30.744 |
| Aldh1a2 | 26.354 |
| Ccl2 | 25.129 |
| Hmox1 | 24.043 |
| Birc5 | 18.964 |
| Cdk1 | 17.992 |
| Gdf15 | 17.833 |
| Hck | 16.390 |
| Ckap2 | 16.271 |
| Star | 16.221 |
| Tpx2 | 15.257 |
| Top2a | 15.100 |
| Ccr2 | 14.716 |
| Crlf1 | 14.542 |
| Lcn2 | 14.296 |
| Inhba | 13.724 |
| Lgals3 | 13.711 |
| Myc | 13.703 |
| Plk1 | 13.340 |
| Ccl7 | 12.622 |
| Cd24a | 11.770 |
| Gdf6 | 9.741 |

Genes Shared by FFAR4 KO Sham vs FFAR4 KO TAC and WT Sham vs WT TAC: 161 increased, 71 decreased

| Gene ID | Fold Change KO | Fold Change WT |
|---|---|---|
| Lox | 24.312 | 34.324 |
| Serpine1 | 16.010 | 24.941 |
| Ankrd1 | 10.962 | 6.718 |
| Cd44 | 9.335 | 9.640 |
| Tgfb2 | 8.828 | 10.745 |
| Gadd45g | 8.745 | 8.200 |
| Dbn1 | 8.621 | 8.140 |
| Fn1 | 7.686 | 8.424 |
| Ier3 | 6.883 | 7.312 |
| Thbs1 | 6.419 | 6.000 |
| Tnfrsf12a | 6.239 | 6.057 |
| Nes | 5.264 | 3.962 |
| Hspb1 | 4.956 | 5.753 |
| Ccl6 | 4.955 | 4.562 |
| Phlda1 | 4.576 | 4.771 |
| Ccl9 | 4.518 | 5.177 |
| Tgm2 | 4.504 | 4.795 |
| Emp1 | 4.430 | 5.579 |
| Tspo | 4.197 | 4.214 |
| Ptprc | 4.103 | 4.334 |
| Fcgr3 | 3.993 | 3.074 |
| Dynll1 | 3.938 | 3.725 |
| Clu | 3.867 | 4.239 |
| Prelid1 | 3.718 | 3.329 |
| Ncam1 | 3.705 | 3.728 |
| Creb3l1 | 3.704 | 4.160 |
| Adcyap1r1 | 3.613 | 4.621 |
| Prmt2 | 3.606 | 2.954 |
| Adamts12 | 3.453 | 3.580 |
| Lgals1 | 3.446 | 4.017 |

| | | | | |
|---|---|---|---|---|
| Ect2 | 9.516 | Plscr1 | 3.381 | 2.523 |
| Havcr2 | 9.472 | Mmp3 | 3.340 | 2.805 |
| Angptl4 | 9.421 | Sfrp1 | 3.307 | 3.030 |
| Bub1b | 8.822 | Otulin | 3.303 | 2.893 |
| Plaur | 7.858 | G6pdx | 3.290 | 3.119 |
| Ptgs2 | 7.153 | Ctsz | 3.283 | 3.126 |
| C5ar1 | 7.009 | Ifi204 | 3.274 | 2.803 |
| Pdpn | 6.605 | Tgfb3 | 3.271 | 2.029 |
| Brca1 | 6.587 | Dap | 3.227 | 3.250 |
| Pak1 | 6.585 | Gba | 3.173 | 2.411 |
| Sphk1 | 6.439 | Tmbim1 | 3.161 | 3.685 |
| Nlrc3 | 6.282 | Fndc1 | 3.103 | 2.449 |
| Kif14 | 6.185 | Cfl1 | 3.086 | 3.469 |
| Cit | 5.892 | Cttn | 2.980 | 3.271 |
| Cd84 | 5.723 | Ptprf | 2.946 | 3.562 |
| Fcer1g | 5.635 | Anxa1 | 2.922 | 2.855 |
| Socs3 | 5.425 | Shb | 2.914 | 2.708 |
| Tnfrsf23 | 5.322 | Slc25a24 | 2.828 | 2.192 |
| Pf4 | 5.217 | Col18a1 | 2.799 | 3.149 |
| Siglec1 | 4.901 | Mllt11 | 2.797 | 3.626 |
| Clec5a | 4.882 | Prkcd | 2.795 | 2.078 |
| Ccr5 | 4.734 | Mal | 2.792 | 3.251 |
| Cd14 | 4.651 | Adam9 | 2.775 | 2.992 |
| Ncf2 | 4.636 | Arid5a | 2.752 | 2.228 |
| Asns | 4.630 | Bak1 | 2.750 | 3.150 |
| Ncf1 | 4.390 | Lmna | 2.742 | 3.030 |
| Hells | 4.317 | Sox9 | 2.653 | 2.948 |
| Fcgr2b | 4.140 | Dab2 | 2.596 | 2.801 |
| Nckap1l | 4.056 | Syk | 2.586 | 3.022 |
| Il33 | 4.052 | Gpx1 | 2.585 | 2.557 |
| Coro1a | 3.957 | Nol12 | 2.579 | 2.751 |
| Itgb3 | 3.912 | Ptgis | 2.577 | 2.278 |
| Ripk3 | 3.867 | Cd248 | 2.557 | 2.515 |
| Phlda3 | 3.835 | Txnrd1 | 2.544 | 2.756 |
| Ptpn6 | 3.746 | P4hb | 2.540 | 2.835 |
| Gpnmb | 3.711 | Usp53 | 2.527 | 3.426 |
| Tlr2 | 3.624 | Creb3 | 2.514 | 2.240 |
| Egr3 | 3.598 | Txndc5 | 2.483 | 2.393 |
| Rad18 | 3.589 | Fam129b | 2.472 | 2.698 |
| Sdf2l1 | 3.551 | Cdkn1a | 2.459 | 2.314 |

| | | | | |
|---|---|---|---|---|
| Nek6 | 3.546 | Bin1 | 2.452 | 2.310 |
| Tox3 | 3.545 | Actn4 | 2.450 | 2.105 |
| Frzb | 3.515 | Lgmn | 2.442 | 2.530 |
| Bnc2 | 3.414 | Ilk | 2.436 | 2.613 |
| Ada | 3.300 | Eef1a1 | 2.411 | 2.414 |
| Shq1 | 3.275 | Lpar1 | 2.370 | 2.072 |
| Met | 3.273 | Flna | 2.316 | 2.414 |
| Srpx | 3.268 | Ptk2b | 2.311 | 2.958 |
| Pik3cg | 3.245 | Cryab | 2.295 | 2.313 |
| Mlkl | 3.236 | Plk2 | 2.294 | 2.215 |
| Inhbb | 3.229 | Gclm | 2.284 | 2.564 |
| Spi1 | 3.141 | Anxa5 | 2.281 | 2.446 |
| Dhcr24 | 3.043 | Hcls1 | 2.274 | 2.609 |
| Blm | 2.997 | Sulf1 | 2.265 | 2.514 |
| Casp3 | 2.931 | Itga4 | 2.244 | 2.883 |
| Cx3cr1 | 2.848 | Itgb1 | 2.223 | 2.307 |
| Rassf5 | 2.806 | Mical1 | 2.222 | 2.817 |
| Pla2g4a | 2.787 | Tnfrsf1a | 2.209 | 2.301 |
| Tmem173 | 2.787 | Plcg2 | 2.198 | 1.956 |
| Src | 2.752 | Trp53 | 2.166 | 2.165 |
| Cd300ld | 2.739 | Sigmar1 | 2.155 | 1.800 |
| Bcl3 | 2.694 | Ppp1r15a | 2.133 | 1.933 |
| Tnfrsf11a | 2.666 | Ecscr | 2.132 | 2.271 |
| Tnfrsf10b | 2.651 | Hdac6 | 2.132 | 2.373 |
| Eef1e1 | 2.600 | Tlr4 | 2.125 | 2.071 |
| Clip3 | 2.563 | Hmgb2 | 2.122 | 2.469 |
| Arrb2 | 2.484 | Csf1r | 2.121 | 1.942 |
| Robo1 | 2.443 | Emp2 | 2.111 | 2.255 |
| Pik3cd | 2.349 | Hif1a | 2.108 | 2.022 |
| Prkcb | 2.113 | Lsp1 | 2.106 | 2.371 |
| Cib1 | 2.066 | Ptprj | 2.104 | 1.777 |
| Bax | 2.038 | Cybb | 2.100 | 2.584 |
| Hax1 | 2.033 | Por | 2.086 | 2.004 |
| Katnb1 | 2.029 | Dlg5 | 2.072 | 2.097 |
| Myd88 | 2.010 | Tnfrsf1b | 2.067 | 2.151 |
| Npm1 | 1.969 | Arf4 | 2.061 | 1.837 |
| Anxa6 | 1.964 | Mt1 | 2.053 | 2.537 |
| Mapk7 | 1.950 | Sra1 | 2.030 | 1.811 |
| Fgfr1 | 1.934 | Capn2 | 2.015 | 2.061 |
| Map1s | 1.918 | Abcc1 | 2.004 | 1.837 |

| | | | | |
|---|---|---|---|---|
| Atad5 | 1.912 | Arf6 | 2.001 | 2.160 |
| Brms1 | 1.894 | Sgpl1 | 1.982 | 2.148 |
| Cpne1 | 1.885 | Rnf4 | 1.977 | 1.764 |
| Atf4 | 1.883 | Ltbr | 1.963 | 2.180 |
| Casp8 | 1.881 | Ywhaz | 1.958 | 2.058 |
| Rpl11 | 1.877 | Pde1a | 1.957 | 1.750 |
| Zmat3 | 1.868 | Hmgcr | 1.955 | 1.999 |
| Apex1 | 1.850 | Ywhah | 1.951 | 1.874 |
| Gadd45b | 1.848 | Xbp1 | 1.935 | 1.979 |
| Eif5a | 1.843 | Shisa5 | 1.935 | 2.218 |
| Map2k1 | 1.839 | Rpl10 | 1.929 | 2.058 |
| Stambp | 1.834 | Emp3 | 1.922 | 2.850 |
| Bad | 1.833 | Lrp1 | 1.916 | 1.889 |
| Pigt | 1.829 | F2r | 1.915 | 2.004 |
| Clptm1l | 1.827 | Fem1b | 1.912 | 1.836 |
| Stat3 | 1.817 | Cxcl16 | 1.906 | 2.152 |
| Ctsc | 1.815 | Inpp5d | 1.903 | 2.647 |
| Ppp1r13l | 1.799 | Pa2g4 | 1.890 | 1.892 |
| Ucp2 | 1.792 | Anp32b | 1.882 | 1.709 |
| Noc2l | 1.786 | Rassf2 | 1.881 | 1.773 |
| Nup62 | 1.780 | Mcm2 | 1.862 | 2.489 |
| Ufm1 | 1.777 | Hk1 | 1.858 | 2.066 |
| Unc5b | 1.761 | Hyal2 | 1.853 | 1.806 |
| Slc9a3r1 | 1.753 | Axl | 1.849 | 2.011 |
| Gnai2 | 1.739 | Mydgf | 1.849 | 2.087 |
| Cidea | 1.723 | Pea15a | 1.849 | 1.775 |
| Ptpn2 | 1.707 | Actr3 | 1.840 | 2.003 |
| Dnmt1 | 1.703 | Fmr1 | 1.834 | 1.908 |
| Hsf1 | -1.702 | Chst11 | 1.834 | 1.836 |
| Opa1 | -1.707 | Dapk3 | 1.833 | 1.719 |
| Aldh2 | -1.721 | Itgav | 1.831 | 1.859 |
| Asah2 | -1.724 | Tgfb1 | 1.830 | 1.963 |
| Rarg | -1.731 | Grk5 | 1.827 | 2.082 |
| Ndufs3 | -1.741 | Tgfbr1 | 1.826 | 1.858 |
| Gstp1 | -1.766 | Pak4 | 1.804 | 1.918 |
| Lims1 | -1.767 | Chmp4b | 1.803 | 1.785 |
| Rora | -1.767 | B4galt1 | 1.792 | 2.046 |
| Flcn | -1.783 | Plvap | 1.789 | 2.024 |
| Pde3a | -1.798 | Rrp1b | 1.787 | 1.710 |
| Itpkb | -1.808 | Ctsb | 1.785 | 2.093 |

| | | | | |
|---|---|---|---|---|
| Tek | -1.816 | Prnp | 1.777 | 1.799 |
| Erbb4 | -1.844 | Txn1 | 1.763 | 2.159 |
| Akt2 | -1.847 | Ralb | 1.762 | 1.903 |
| Cep63 | -1.848 | Rara | 1.750 | 1.728 |
| Txnip | -1.849 | Crip1 | 1.743 | 2.134 |
| Chchd10 | -1.867 | Gli3 | 1.738 | 2.298 |
| Ypel3 | -1.874 | Gatad2a | 1.732 | 1.816 |
| Jmy | -1.881 | Ptpn1 | 1.729 | 2.025 |
| Rapgef2 | -1.888 | Plscr3 | 1.726 | 1.934 |
| Endog | -1.889 | Pmp22 | 1.714 | 1.736 |
| Tbx3 | -1.895 | Shc1 | 1.709 | 2.106 |
| Pde5a | -1.902 | Pcnt | -1.708 | -2.008 |
| Alkbh7 | -1.918 | Slc25a4 | -1.709 | -1.835 |
| Kcnb1 | -1.924 | Sort1 | -1.710 | -1.915 |
| Hint2 | -1.935 | Gpam | -1.714 | -2.049 |
| Rnf146 | -1.940 | Mavs | -1.722 | -1.791 |
| Sh3kbp1 | -1.968 | Abcb1a | -1.728 | -2.190 |
| Prkd1 | -1.973 | Wdr92 | -1.730 | -1.893 |
| Sncaip | -1.974 | Xdh | -1.731 | -1.825 |
| Rassf3 | -1.976 | Dpep1 | -1.735 | -2.188 |
| Zbtb16 | -1.991 | Cecr2 | -1.753 | -2.072 |
| Six4 | -1.994 | Nsmaf | -1.755 | -2.085 |
| Pnpla8 | -2.032 | Optn | -1.760 | -2.252 |
| Aamdc | -2.040 | Bcl2l11 | -1.769 | -2.251 |
| Col4a3 | -2.048 | Cyfip2 | -1.769 | -2.076 |
| Map3k5 | -2.062 | Acaa2 | -1.772 | -1.909 |
| Kdr | -2.126 | Cdkn1b | -1.774 | -1.813 |
| Irf1 | -2.133 | Phlpp1 | -1.779 | -1.766 |
| Epha7 | -2.138 | Smpd2 | -1.791 | -1.964 |
| Zc3h8 | -2.141 | Msh2 | -1.794 | -1.867 |
| Rrm2b | -2.142 | Jun | -1.810 | -1.759 |
| Sept4 | -2.144 | Pdcd4 | -1.823 | -1.795 |
| Bmp6 | -2.148 | Tnfaip8 | -1.856 | -2.613 |
| Egr1 | -2.284 | Ube2b | -1.858 | -2.316 |
| Ackr3 | -2.301 | Nmnat1 | -1.862 | -2.469 |
| Tnfsf10 | -2.362 | Sod2 | -1.863 | -2.121 |
| Id1 | -2.367 | Fgf13 | -1.864 | -1.703 |
| Tmc8 | -2.456 | Agtr1a | -1.868 | -1.727 |
| Tnfrsf21 | -2.496 | Rgcc | -1.878 | -2.006 |
| Plekhf1 | -2.527 | Apbb1 | -1.896 | -1.712 |

| | |
|---|---|
| Camk2b | -2.688 |
| Rapsn | -2.692 |
| Dll1 | -2.711 |
| Wnt5a | -2.734 |
| Gata2 | -2.748 |
| Cd28 | -2.776 |
| Ceacam1 | -2.786 |
| Cradd | -2.800 |
| Tpd52l1 | -3.067 |
| Ccdc3 | -3.084 |
| Map2k6 | -3.425 |
| Gzmm | -3.537 |
| Gadd45a | -3.622 |
| Prkcq | -3.793 |
| Ung | -3.926 |
| Atcay | -5.338 |
| Ntf3 | -5.417 |
| Ptgfr | -5.664 |

| | | |
|---|---|---|
| Ago4 | -1.916 | -2.475 |
| Gm20594 | -1.936 | -2.822 |
| Lims2 | -1.938 | -1.930 |
| Ndufs1 | -1.943 | -2.387 |
| Tgfbr3 | -1.947 | -1.725 |
| Ivns1abp | -1.971 | -2.709 |
| Pink1 | -1.974 | -2.209 |
| Mapt | -1.975 | -2.059 |
| Ahr | -2.000 | -2.165 |
| Vegfb | -2.001 | -2.218 |
| Mitf | -2.008 | -1.789 |
| Lifr | -2.033 | -2.257 |
| Stat5b | -2.034 | -1.833 |
| Thrb | -2.035 | -2.350 |
| Ndnf | -2.038 | -1.973 |
| Sema6a | -2.086 | -1.958 |
| Pdk2 | -2.103 | -2.685 |
| Mtfp1 | -2.104 | -2.379 |
| Camk2a | -2.117 | -1.759 |
| Fbxo32 | -2.129 | -2.753 |
| Sox7 | -2.140 | -2.402 |
| Nqo2 | -2.199 | -2.352 |
| F3 | -2.209 | -2.642 |
| Foxo3 | -2.219 | -2.307 |
| Ccng1 | -2.229 | -2.336 |
| Ank2 | -2.232 | -2.457 |
| Egln1 | -2.240 | -2.352 |
| Rps6ka2 | -2.251 | -2.898 |
| Abcc8 | -2.273 | -2.690 |
| Slc2a4 | -2.297 | -2.415 |
| Ddt | -2.305 | -2.024 |
| Pik3r1 | -2.401 | -2.446 |
| Adamts7 | -2.457 | -2.723 |
| Ppargc1a | -2.536 | -3.388 |
| Cd274 | -2.549 | -3.672 |
| G0s2 | -2.563 | -2.086 |
| Apip | -2.620 | -2.852 |
| Aqp1 | -2.678 | -3.200 |
| Dusp1 | -2.695 | -3.057 |
| Gsn | -2.718 | -3.531 |

| | | |
|---|---|---|
| Herpud1 | -2.751 | -3.134 |
| Angpt1 | -4.454 | -4.113 |

**Supplemental Table 6. Inflammation Genes**

GO Terms: monocyte, dendritic cell, macrophage, myeloid, B cell, T cell, lymph, interferon, interleukin, inflammatory, lymphocyte, leukocyte, chemokine, cytokine, immune

Genes Unique to FFAR4 KO Sham vs FFAR4 KO TAC: 23 increased, 3 decreased

| Gene ID | Fold Change |
|---|---|
| Cx3cl1 | 2.987 |
| Igf1 | 2.787 |
| Il2rg | 2.152 |
| Klf6 | 2.127 |
| Ahcy | 2.055 |
| Cyba | 2.019 |
| Trove2 | 1.970 |
| Maf | 1.964 |
| Ssc5d | 1.963 |
| Errfi1 | 1.882 |
| Flnb | 1.843 |
| Ticam1 | 1.824 |
| Ifitm3 | 1.805 |
| Elf4 | 1.792 |
| Trim27 | 1.769 |
| Bcl2 | 1.768 |
| Smad1 | 1.757 |
| Pnp | 1.740 |
| Ace | 1.732 |
| Eif2ak4 | 1.724 |
| Tinagl1 | 1.720 |
| Prex1 | 1.716 |
| Sqstm1 | 1.706 |
| Ndufc2 | -1.708 |
| Hdac9 | -1.759 |
| Jam3 | -1.792 |

Genes Unique to WT Sham vs WT TAC: 166 increased, 73 decreased

| Gene ID | Fold Change |
|---|---|
| Ereg | 240.348 |
| Il1rn | 219.636 |
| Chil3 | 211.386 |
| Adam8 | 117.349 |
| Timp1 | 112.492 |
| Arg1 | 94.628 |
| Ankrd2 | 47.887 |
| Lrp8 | 37.091 |
| Cxcl5 | 30.522 |
| Aldh1a2 | 26.354 |
| Ccl2 | 25.129 |
| Hmox1 | 24.043 |
| Serpina3i | 22.500 |
| Anln | 22.149 |
| Ccnb2 | 19.624 |
| Birc5 | 18.964 |
| Cxcr6 | 18.350 |
| Hck | 16.390 |
| Ckap2 | 16.271 |
| Star | 16.221 |
| Ccr2 | 14.716 |
| Lcn2 | 14.296 |
| Lgals3 | 13.711 |
| Myc | 13.703 |
| Plk1 | 13.340 |
| Ccl7 | 12.622 |
| Msr1 | 12.445 |
| Cd24a | 11.770 |
| Csf2rb | 11.085 |

Genes Shared by FFAR4 KO Sham vs FFAR4 KO TAC and WT Sham vs WT TAC: 168 increased, 56 decreased

| Gene ID | Fold Change KO | Fold Change WT |
|---|---|---|
| Serpine1 | 16.010 | 24.941 |
| Postn | 15.162 | 13.826 |
| Serpina3n | 13.726 | 16.369 |
| Ankrd1 | 10.962 | 6.718 |
| Col3a1 | 10.165 | 9.226 |
| Itgam | 9.947 | 10.004 |
| Cd44 | 9.335 | 9.640 |
| Tgfb2 | 8.828 | 10.745 |
| Gadd45g | 8.745 | 8.200 |
| Nppb | 8.688 | 5.931 |
| Loxl3 | 8.357 | 14.007 |
| Fn1 | 7.686 | 8.424 |
| Ier3 | 6.883 | 7.312 |
| Itga5 | 6.523 | 7.683 |
| Thbs1 | 6.419 | 6.000 |
| Fyb | 6.236 | 5.579 |
| Prg4 | 6.235 | 6.559 |
| Ctss | 6.082 | 6.240 |
| Sbno2 | 5.549 | 5.008 |
| Capg | 5.358 | 5.617 |
| Lcp1 | 5.083 | 6.470 |
| Il4ra | 5.027 | 4.673 |
| Hspb1 | 4.956 | 5.753 |
| Ccl6 | 4.955 | 4.562 |
| Msn | 4.820 | 5.174 |
| Rhoc | 4.626 | 4.705 |
| Mmp14 | 4.608 | 5.418 |
| Ccl9 | 4.518 | 5.177 |
| Tgm2 | 4.504 | 4.795 |

| | | | | |
|---|---|---|---|---|
| Runx1 | 10.581 | Emilin1 | 4.265 | 4.364 |
| Gpr35 | 10.420 | Ptprc | 4.103 | 4.334 |
| Ccr1 | 9.871 | Litaf | 4.072 | 3.362 |
| Racgap1 | 9.775 | Vim | 4.049 | 3.986 |
| Ect2 | 9.516 | Fcgr3 | 3.993 | 3.074 |
| Havcr2 | 9.472 | Soat1 | 3.896 | 3.758 |
| Lat2 | 9.451 | Ldlr | 3.813 | 4.756 |
| Itgb2 | 9.094 | Prelid1 | 3.718 | 3.329 |
| Cd180 | 9.072 | Trim16 | 3.681 | 4.547 |
| Prc1 | 8.639 | Rnf19b | 3.642 | 4.003 |
| Kif23 | 8.412 | Tuba1b | 3.497 | 3.072 |
| Dok1 | 8.142 | Adamts12 | 3.453 | 3.580 |
| Pik3ap1 | 8.033 | Lgals1 | 3.446 | 4.017 |
| Kif4 | 7.863 | Plscr1 | 3.381 | 2.523 |
| Ptx3 | 7.589 | Ifi30 | 3.347 | 3.281 |
| Tlr13 | 7.379 | Pla2g7 | 3.332 | 4.630 |
| Myo1g | 7.226 | Sfrp1 | 3.307 | 3.030 |
| Ptgs2 | 7.153 | Otulin | 3.303 | 2.893 |
| Kif20a | 7.039 | G6pdx | 3.290 | 3.119 |
| C5ar1 | 7.009 | Osmr | 3.278 | 3.525 |
| Sele | 6.769 | Ifi204 | 3.274 | 2.803 |
| Pdpn | 6.605 | Tgfb3 | 3.271 | 2.029 |
| Sphk1 | 6.439 | Klhl6 | 3.194 | 2.011 |
| Pirb | 6.345 | Gba | 3.173 | 2.411 |
| Nlrc3 | 6.282 | Masp1 | 3.107 | 2.701 |
| C3ar1 | 6.280 | Cfl1 | 3.086 | 3.469 |
| Csf2rb2 | 6.265 | Ppp1r14b | 2.963 | 2.678 |
| Kif14 | 6.185 | Adcy7 | 2.959 | 2.815 |
| Vav1 | 5.911 | Enpp1 | 2.956 | 3.722 |
| Cit | 5.892 | Ptprf | 2.946 | 3.562 |
| Kif20b | 5.861 | Sdc1 | 2.941 | 4.236 |
| Slc11a1 | 5.774 | Anxa1 | 2.922 | 2.855 |
| Cd84 | 5.723 | Shb | 2.914 | 2.708 |
| Il17ra | 5.718 | Nme1 | 2.889 | 3.359 |
| Fcer1g | 5.635 | Zyx | 2.880 | 2.798 |
| Fcgr1 | 5.594 | Ctps | 2.861 | 3.016 |
| Tyrobp | 5.482 | C1qa | 2.831 | 2.521 |
| Socs3 | 5.425 | Cblb | 2.830 | 2.564 |
| Pf4 | 5.217 | Adgre1 | 2.830 | 3.153 |
| Siglec1 | 4.901 | Bcr | 2.803 | 2.980 |

| | | | | |
|---|---|---|---|---|
| Clec5a | 4.882 | Nppa | 2.800 | 4.948 |
| Myo1f | 4.761 | Prkcd | 2.795 | 2.078 |
| Ccr5 | 4.734 | Apbb1ip | 2.779 | 4.058 |
| Cd14 | 4.651 | Adam9 | 2.775 | 2.992 |
| Panx1 | 4.538 | Rpl3 | 2.761 | 2.997 |
| Ncf1 | 4.390 | Arid5a | 2.752 | 2.228 |
| Hells | 4.317 | Bak1 | 2.750 | 3.150 |
| Alox5ap | 4.293 | Unc93b1 | 2.747 | 2.081 |
| E2f8 | 4.272 | C1qb | 2.736 | 3.385 |
| Fcgr2b | 4.140 | Myh9 | 2.705 | 2.344 |
| Ptprn | 4.104 | C1qc | 2.665 | 2.601 |
| Nckap1l | 4.056 | Sox9 | 2.653 | 2.948 |
| Il33 | 4.052 | Mrc1 | 2.649 | 2.420 |
| Dock2 | 4.017 | Ctla2a | 2.635 | 2.344 |
| Clec12a | 3.989 | Dab2 | 2.596 | 2.801 |
| Nusap1 | 3.957 | Syk | 2.586 | 3.022 |
| Coro1a | 3.957 | Gpx1 | 2.585 | 2.557 |
| Ly86 | 3.942 | Ptgis | 2.577 | 2.278 |
| Itgb3 | 3.912 | Cd248 | 2.557 | 2.515 |
| Fermt3 | 3.897 | P4hb | 2.540 | 2.835 |
| Ripk3 | 3.867 | Actg1 | 2.531 | 2.528 |
| Blk | 3.841 | Nt5e | 2.521 | 1.947 |
| Ptpn6 | 3.746 | Anxa4 | 2.517 | 2.602 |
| Gpnmb | 3.711 | Creb3 | 2.514 | 2.240 |
| Thy1 | 3.651 | Stab1 | 2.502 | 2.184 |
| Nfam1 | 3.632 | Serpinf1 | 2.488 | 2.428 |
| Cfp | 3.626 | Iqgap1 | 2.488 | 2.517 |
| Tlr2 | 3.624 | Cdkn1a | 2.459 | 2.314 |
| Hist1h2bg | 3.605 | Lgmn | 2.442 | 2.530 |
| Egr3 | 3.598 | Rras | 2.397 | 2.153 |
| Incenp | 3.347 | Myo1c | 2.360 | 2.894 |
| Ada | 3.300 | Sdc4 | 2.343 | 2.329 |
| Met | 3.273 | Ptk2b | 2.311 | 2.958 |
| Nfkbiz | 3.249 | Vcam1 | 2.304 | 2.086 |
| Pik3cg | 3.245 | Plk2 | 2.294 | 2.215 |
| Spi1 | 3.141 | Bcar1 | 2.279 | 1.778 |
| Numbl | 3.043 | Hcls1 | 2.274 | 2.609 |
| Lcp2 | 3.022 | Itga4 | 2.244 | 2.883 |
| Blm | 2.997 | Itgb1 | 2.223 | 2.307 |
| Sept5 | 2.991 | Tnfrsf1a | 2.209 | 2.301 |

| | | | | |
|---|---|---|---|---|
| Casp3 | 2.931 | Sept11 | 2.201 | 1.849 |
| Ikzf1 | 2.902 | Plcg2 | 2.198 | 1.956 |
| Cx3cr1 | 2.848 | Fkbp1a | 2.197 | 2.456 |
| Dhx58 | 2.843 | Trp53 | 2.166 | 2.165 |
| Alcam | 2.796 | Cspg4 | 2.148 | 2.136 |
| Pla2g4a | 2.787 | Hsp90aa1 | 2.144 | 2.216 |
| Tmem173 | 2.787 | Tlr4 | 2.125 | 2.071 |
| Src | 2.752 | Hmgb2 | 2.122 | 2.469 |
| Cd300ld | 2.739 | Csf1r | 2.121 | 1.942 |
| Cdc25b | 2.733 | Ptms | 2.120 | 2.073 |
| Chst3 | 2.732 | Emp2 | 2.111 | 2.255 |
| Themis2 | 2.703 | Hif1a | 2.108 | 2.022 |
| Bcl3 | 2.694 | Lsp1 | 2.106 | 2.371 |
| Tnfrsf11a | 2.666 | Ptprj | 2.104 | 1.777 |
| Arrb2 | 2.484 | Cybb | 2.100 | 2.584 |
| Robo1 | 2.443 | Dlg5 | 2.072 | 2.097 |
| D1Ertd622e | 2.416 | Tnfrsf1b | 2.067 | 2.151 |
| Lxn | 2.378 | Ddx21 | 2.036 | 1.879 |
| Rab7b | 2.371 | Cd151 | 2.033 | 2.030 |
| Cmtm3 | 2.353 | Skap2 | 2.026 | 3.448 |
| Pik3cd | 2.349 | Lrrc32 | 2.022 | 1.833 |
| Irf8 | 2.246 | Gprc5b | 2.020 | 1.770 |
| Hist1h2be | 2.212 | Capn2 | 2.015 | 2.061 |
| Mapk11 | 2.199 | Nras | 2.013 | 2.256 |
| Prkcb | 2.113 | Arf6 | 2.001 | 2.160 |
| Cfb | 2.075 | Il1r1 | 1.999 | 1.782 |
| Bax | 2.038 | Ltbr | 1.963 | 2.180 |
| Hax1 | 2.033 | Dysf | 1.936 | 2.127 |
| Gcnt1 | 2.029 | Myo9b | 1.935 | 1.994 |
| Relb | 2.022 | Xbp1 | 1.935 | 1.979 |
| Polr3g | 2.014 | Lrp1 | 1.916 | 1.889 |
| Myd88 | 2.010 | F2r | 1.915 | 2.004 |
| Csf2ra | 2.002 | Cxcl16 | 1.906 | 2.152 |
| Susd2 | 1.987 | Inpp5d | 1.903 | 2.647 |
| Ecm1 | 1.966 | Nfil3 | 1.880 | 2.121 |
| Mapk7 | 1.950 | Mcm2 | 1.862 | 2.489 |
| Atad5 | 1.912 | Hk1 | 1.858 | 2.066 |
| Casp8 | 1.881 | Hyal2 | 1.853 | 1.806 |
| Nfkb2 | 1.866 | Axl | 1.849 | 2.011 |
| Stambp | 1.834 | Actr3 | 1.840 | 2.003 |

| | | | | |
|---|---|---|---|---|
| Bad | 1.833 | Calr | 1.837 | 1.886 |
| Stat3 | 1.817 | Dapk3 | 1.833 | 1.719 |
| Ctsc | 1.815 | Itgav | 1.831 | 1.859 |
| Rps6ka4 | 1.808 | Tgfb1 | 1.830 | 1.963 |
| Atic | 1.804 | Slc7a2 | 1.803 | 2.109 |
| Rpl39 | 1.787 | Csk | 1.803 | 1.709 |
| Noc2l | 1.786 | Chmp4b | 1.803 | 1.785 |
| Nup62 | 1.780 | Cdk6 | 1.801 | 1.781 |
| Dhx33 | 1.773 | B4galt1 | 1.792 | 2.046 |
| Apobec3 | 1.765 | Prnp | 1.777 | 1.799 |
| Map2k3 | 1.763 | Tmem176b | 1.774 | 2.331 |
| Arhgef5 | 1.746 | Dbnl | 1.754 | 2.002 |
| Adam15 | 1.726 | Rara | 1.750 | 1.728 |
| Cidea | 1.723 | Ddost | 1.744 | 1.901 |
| Ptpn2 | 1.707 | Gli3 | 1.738 | 2.298 |
| Impdh2 | 1.703 | Rpl13a | 1.722 | 1.813 |
| Zbtb7b | 1.700 | Pmp22 | 1.714 | 1.736 |
| Parp14 | -1.705 | Fam49b | 1.712 | 1.800 |
| Slc39a3 | -1.706 | Pvr | 1.708 | 1.864 |
| Rarg | -1.731 | Btnl9 | -1.701 | -2.683 |
| Flt1 | -1.746 | Gpam | -1.714 | -2.049 |
| Gstp1 | -1.766 | Mavs | -1.722 | -1.791 |
| Rora | -1.767 | Lpl | -1.737 | -2.346 |
| Alox5 | -1.779 | Rab12 | -1.760 | -1.747 |
| Flcn | -1.783 | Optn | -1.760 | -2.252 |
| Klf2 | -1.791 | Bcl2l11 | -1.769 | -2.251 |
| H2-Q4 | -1.807 | Phlpp1 | -1.779 | -1.766 |
| Itpkb | -1.808 | Msh2 | -1.794 | -1.867 |
| Tek | -1.816 | Siae | -1.799 | -1.804 |
| Mr1 | -1.834 | Jun | -1.810 | -1.759 |
| Cfh | -1.879 | Gbp5 | -1.815 | -2.163 |
| Rapgef2 | -1.888 | Pdcd4 | -1.823 | -1.795 |
| Kif16b | -1.898 | Prox1 | -1.826 | -2.173 |
| Pde5a | -1.902 | Irgm2 | -1.828 | -2.669 |
| Irf2bp2 | -1.925 | Il10rb | -1.831 | -2.017 |
| Cd300lg | -1.928 | Ephx2 | -1.832 | -1.951 |
| Gbp3 | -1.938 | Tnfaip8 | -1.856 | -2.613 |
| Il15ra | -1.954 | Wfdc1 | -1.862 | -2.771 |
| Kdm5d | -1.958 | Agtr1a | -1.868 | -1.727 |
| Rarres2 | -1.969 | Map4k2 | -1.872 | -2.778 |

| | | | | | |
|---|---|---|---|---|---|
| Prkd1 | -1.973 | | Rgcc | -1.878 | -2.006 |
| Dll4 | -1.979 | | Klhl21 | -1.884 | -1.824 |
| Tmem204 | -1.980 | | Itgb6 | -1.918 | -2.285 |
| Tspan2 | -1.987 | | Trpm4 | -1.936 | -1.708 |
| Zbtb16 | -1.991 | | Tgfbr3 | -1.947 | -1.725 |
| Kat8 | -1.993 | | Igf2 | -1.950 | -1.939 |
| C1s1 | -2.009 | | Alad | -1.970 | -2.073 |
| Map3k5 | -2.062 | | Gbp4 | -1.984 | -2.377 |
| Gbp9 | -2.071 | | Gbp7 | -1.997 | -2.989 |
| Pdgfd | -2.100 | | Ahr | -2.000 | -2.165 |
| Kdr | -2.126 | | Crhr2 | -2.026 | -1.926 |
| Irf1 | -2.133 | | Lifr | -2.033 | -2.257 |
| Zc3h8 | -2.141 | | Stat5b | -2.034 | -1.833 |
| Sept4 | -2.144 | | Mylk3 | -2.041 | -2.546 |
| Bmp6 | -2.148 | | Vtn | -2.045 | -3.620 |
| Abcd2 | -2.278 | | Igtp | -2.048 | -1.735 |
| Klhl13 | -2.282 | | Timp4 | -2.087 | -2.694 |
| Egr1 | -2.284 | | Ccbe1 | -2.092 | -2.587 |
| Adgrf5 | -2.285 | | F3 | -2.209 | -2.642 |
| Ackr3 | -2.301 | | Iigp1 | -2.287 | -2.244 |
| Fzd8 | -2.320 | | Cd200 | -2.288 | -2.106 |
| Tnfsf10 | -2.362 | | Ddt | -2.305 | -2.024 |
| Tnfrsf21 | -2.496 | | Pik3r1 | -2.401 | -2.446 |
| Ccrl2 | -2.506 | | Gbp6 | -2.453 | -4.015 |
| Slc26a6 | -2.548 | | Adamts7 | -2.457 | -2.723 |
| Prdm1 | -2.648 | | Pde4d | -2.497 | -2.374 |
| Dll1 | -2.711 | | Nr1d2 | -2.518 | -2.834 |
| Wnt5a | -2.734 | | Cd274 | -2.549 | -3.672 |
| Gata2 | -2.748 | | Ppara | -2.614 | -2.691 |
| Cd28 | -2.776 | | Dusp1 | -2.695 | -3.057 |
| Ceacam1 | -2.786 | | Gsn | -2.718 | -3.531 |
| Colec11 | -2.848 | | Lpin1 | -2.759 | -2.924 |
| P2ry14 | -2.855 | | Mlf1 | -2.995 | -3.053 |
| Enpp2 | -3.196 | | Efnb3 | -4.012 | -3.701 |
| Rps6ka5 | -3.390 | | Angpt1 | -4.454 | -4.113 |
| Cxcl9 | -3.448 | | | | |
| Gzmm | -3.537 | | | | |
| Tgtp2 | -3.550 | | | | |
| Aqp4 | -3.555 | | | | |
| Prkcq | -3.793 | | | | |

| | |
|---|---|
| Dpp4 | -3.837 |
| F830016B08Rik | -3.931 |
| Gbp10 | -4.311 |
| Tgtp1 | -5.041 |
| Mme | -5.251 |
| Ptgfr | -5.664 |
| Cdkn1c | -5.729 |
| Pla2g5 | -5.822 |
| Il15 | -5.997 |
| Cyp26b1 | -23.267 |

**Supplemental Table 7. Contractile Function Genes**

GO Terms: Heart contraction, cardiac muscle cell contraction, sarcomere, heart rate, ryanodine, sarcoplasmic reticulum, cardiac muscle cell action potential involved in contraction, cardiac contraction

Genes Unique to FFAR4 KO Sham vs FFAR4 KO TAC: 2 increased, 2 decreased

| Gene ID | Fold Change |
|---|---|
| Myh7 | 2.484 |
| Ace | 1.732 |
| Kcnj5 | -1.897 |
| Bmp10 | -8.321 |

Genes Unique to WT Sham vs WT TAC: 10 increased, 15 decreased

| Gene ID | Fold Change |
|---|---|
| Tnnt3 | 6.630 |
| Ccr5 | 4.734 |
| Ada | 3.300 |
| Met | 3.273 |
| Kcne4 | 2.899 |
| Atp1b1 | 1.927 |
| Pkp2 | 1.886 |
| Ppp1r13l | 1.799 |
| Map2k3 | 1.763 |
| Gnai2 | 1.739 |
| Gaa | -1.711 |
| Taz | -1.722 |
| Ctnna3 | -1.798 |
| Myom2 | -1.843 |
| Tnni3k | -1.889 |
| Pde5a | -1.902 |
| Hcn4 | -1.993 |
| Six4 | -1.994 |
| Epas1 | -2.109 |
| Tnnt1 | -2.132 |
| Kcna5 | -2.157 |
| Map2k6 | -3.425 |
| Cacna2d2 | -3.520 |
| Sln | -6.103 |
| Scn4b | -8.311 |

Genes Shared by FFAR4 KO Sham vs FFAR4 KO TAC and WT Sham vs WT TAC: 20 increased, 31 decreased

| Gene ID | Fold Change KO | Fold Change WT |
|---|---|---|
| Hbegf | 12.860 | 11.692 |
| Tgfb2 | 8.828 | 10.745 |
| Synpo2l | 5.543 | 4.979 |
| Xirp1 | 3.534 | 3.189 |
| Ankrd23 | 3.449 | 2.536 |
| Cav3 | 3.100 | 3.035 |
| Klhl41 | 3.026 | 2.837 |
| Nppa | 2.800 | 4.948 |
| Gpx1 | 2.585 | 2.557 |
| Wdr1 | 2.581 | 2.653 |
| Actg1 | 2.531 | 2.528 |
| Bin1 | 2.452 | 2.310 |
| Itgb1 | 2.223 | 2.307 |
| Csrp3 | 2.219 | 2.003 |
| Fkbp1a | 2.197 | 2.456 |
| Sri | 2.008 | 1.881 |
| Tmem38b | 1.900 | 2.004 |
| Gnao1 | 1.861 | 2.401 |
| Popdc2 | 1.815 | 1.864 |
| Slc8a1 | 1.727 | 1.886 |
| Trdn | -1.740 | -1.799 |
| Tmem38a | -1.815 | -2.108 |
| Prox1 | -1.826 | -2.173 |
| Tnni3 | -1.927 | -2.143 |
| Trpm4 | -1.936 | -1.708 |
| Vegfb | -2.001 | -2.218 |
| Crhr2 | -2.026 | -1.926 |
| Atp2a2 | -2.035 | -2.385 |

| | | |
|---|---|---|
| Thrb | -2.035 | -2.350 |
| Myh6 | -2.039 | -2.507 |
| Cacna1g | -2.040 | -1.762 |
| Mylk3 | -2.041 | -2.546 |
| Sp4 | -2.043 | -2.230 |
| Myl2 | -2.057 | -2.436 |
| Atp1a2 | -2.207 | -2.710 |
| Ank2 | -2.232 | -2.457 |
| Hopx | -2.233 | -3.247 |
| Rps6ka2 | -2.251 | -2.898 |
| Pik3r1 | -2.401 | -2.446 |
| Pln | -2.439 | -2.656 |
| Adra1b | -2.439 | -2.145 |
| Pde4d | -2.497 | -2.374 |
| Gstm7 | -2.611 | -2.433 |
| Kcnh2 | -2.618 | -2.107 |
| Tcap | -2.645 | -2.400 |
| Myl4 | -2.994 | -3.758 |
| Kcnj2 | -3.256 | -3.366 |
| Rgs2 | -3.313 | -3.246 |
| Hrc | -3.466 | -5.100 |
| Adra1a | -3.544 | -3.681 |
| Dhrs7c | -4.965 | -7.365 |

**Supplemental Table 8. Angiogenesis Genes**

GO Terms: Angiogenesis

Genes Unique to FFAR4 KO Sham vs FFAR4 KO TAC:
5 increased,
2 decreased

| Gene ID | Fold Change |
|---|---|
| Cx3cl1 | 2.987 |
| Mfge8 | 1.814 |
| Adamts9 | 1.760 |
| Smad1 | 1.757 |
| Ace | 1.732 |
| Hdac9 | -1.759 |
| Jam3 | -1.792 |

Genes Unique to WT Sham vs WT TAC:
39 increased,
31 decreased

| Gene ID | Fold Change |
|---|---|
| Ereg | 240.348 |
| Adam8 | 117.349 |
| Adam12 | 38.177 |
| Ccl2 | 25.129 |
| Hmox1 | 24.043 |
| Ccr2 | 14.716 |
| Lgals3 | 13.711 |
| Runx1 | 10.581 |
| Angptl4 | 9.421 |
| Itgb2 | 9.094 |
| Ptgs2 | 7.153 |
| C5ar1 | 7.009 |
| Pdpn | 6.605 |
| Brca1 | 6.587 |
| Sphk1 | 6.439 |
| C3ar1 | 6.280 |
| Mmp19 | 5.480 |
| Pf4 | 5.217 |
| Bmper | 4.639 |
| E2f8 | 4.272 |
| Itgb3 | 3.912 |
| Thy1 | 3.651 |
| Egr3 | 3.598 |
| Plau | 3.321 |
| Pik3cg | 3.245 |
| Cx3cr1 | 2.848 |
| Robo1 | 2.443 |
| Prkcb | 2.113 |

Genes Shared by FFAR4 KO Sham vs FFAR4 KO TAC and WT Sham vs WT TAC:
58 increased, 18 decreased

| Gene ID | Fold Change KO | Fold Change WT |
|---|---|---|
| Rtn4 | 17.020 | 17.496 |
| Serpine1 | 16.010 | 24.941 |
| Hbegf | 12.860 | 11.692 |
| Col8a1 | 9.567 | 9.110 |
| Tgfb2 | 8.828 | 10.745 |
| Fn1 | 7.686 | 8.424 |
| Loxl2 | 7.354 | 7.545 |
| Itga5 | 6.523 | 7.683 |
| Thbs1 | 6.419 | 6.000 |
| Tnfrsf12a | 6.239 | 6.057 |
| Anxa2 | 5.166 | 6.442 |
| Hspb1 | 4.956 | 5.753 |
| Srpx2 | 4.921 | 4.422 |
| Adamts1 | 4.522 | 4.112 |
| Emilin1 | 4.265 | 4.364 |
| Creb3l1 | 3.704 | 4.160 |
| Sparc | 3.700 | 3.853 |
| Otulin | 3.303 | 2.893 |
| Col4a1 | 3.301 | 3.032 |
| Anxa1 | 2.922 | 2.855 |
| Shb | 2.914 | 2.708 |
| Ddah1 | 2.803 | 3.495 |
| Col18a1 | 2.799 | 3.149 |
| Col4a2 | 2.797 | 2.674 |
| Myh9 | 2.705 | 2.344 |
| Syk | 2.586 | 3.022 |
| Gpx1 | 2.585 | 2.557 |
| Ptgis | 2.577 | 2.278 |

| | |
|---|---|
| Cib1 | 2.066 |
| Ecm1 | 1.966 |
| Mapk7 | 1.950 |
| Fgfr1 | 1.934 |
| Casp8 | 1.881 |
| Kctd10 | 1.863 |
| Stat3 | 1.817 |
| Hs6st1 | 1.793 |
| Unc5b | 1.761 |
| Hgs | 1.755 |
| Adam15 | 1.726 |
| Tie1 | -1.705 |
| Dcn | -1.738 |
| Flt1 | -1.746 |
| Ptprm | -1.763 |
| Synj2bp | -1.764 |
| Rora | -1.767 |
| Flcn | -1.783 |
| Klf2 | -1.791 |
| Ptprb | -1.804 |
| Foxo4 | -1.804 |
| Tek | -1.816 |
| Cd59a | -1.832 |
| Prkd1 | -1.973 |
| Dll4 | -1.979 |
| Reck | -2.041 |
| Col4a3 | -2.048 |
| Epas1 | -2.109 |
| Kdr | -2.126 |
| Plxdc1 | -2.205 |
| Ackr3 | -2.301 |
| Fzd8 | -2.320 |
| Id1 | -2.367 |
| Smoc2 | -2.392 |
| Fgf9 | -2.474 |
| Dll1 | -2.711 |
| Wnt5a | -2.734 |
| Gata2 | -2.748 |
| Ceacam1 | -2.786 |
| Enpp2 | -3.196 |

| | | |
|---|---|---|
| Stab1 | 2.502 | 2.184 |
| Serpinf1 | 2.488 | 2.428 |
| Anxa3 | 2.469 | 2.291 |
| Rras | 2.397 | 2.153 |
| Amot | 2.347 | 1.769 |
| Flna | 2.316 | 2.414 |
| Ptk2b | 2.311 | 2.958 |
| Plk2 | 2.294 | 2.215 |
| Sulf1 | 2.265 | 2.514 |
| Itgb1 | 2.223 | 2.307 |
| Tnfrsf1a | 2.209 | 2.301 |
| Cspg4 | 2.148 | 2.136 |
| Ecscr | 2.132 | 2.271 |
| Mcam | 2.131 | 1.882 |
| Emp2 | 2.111 | 2.255 |
| Hif1a | 2.108 | 2.022 |
| Cybb | 2.100 | 2.584 |
| Rnh1 | 2.067 | 2.098 |
| Parva | 2.013 | 1.962 |
| Nras | 2.013 | 2.256 |
| Glul | 1.998 | 1.937 |
| Dysf | 1.936 | 2.127 |
| Xbp1 | 1.935 | 1.979 |
| Clic4 | 1.871 | 1.906 |
| Mydgf | 1.849 | 2.087 |
| Itgav | 1.831 | 1.859 |
| Tgfbr1 | 1.826 | 1.858 |
| Pak4 | 1.804 | 1.918 |
| B4galt1 | 1.792 | 2.046 |
| Shc1 | 1.709 | 2.106 |
| Gtf2i | -1.754 | -1.711 |
| Jun | -1.810 | -1.759 |
| Prox1 | -1.826 | -2.173 |
| Agtr1a | -1.868 | -1.727 |
| Rgcc | -1.878 | -2.006 |
| Ppp1r16b | -1.902 | -2.501 |
| Igf2 | -1.950 | -1.939 |
| Vegfb | -2.001 | -2.218 |
| Crhr2 | -2.026 | -1.926 |
| Ndnf | -2.038 | -1.973 |

| | | | | |
|---|---|---|---|---|
| Egf | -4.469 | Col4a4 | -2.072 | -2.081 |
| Ephb1 | -5.173 | Sema6a | -2.086 | -1.958 |
| | | Ccbe1 | -2.092 | -2.587 |
| | | F3 | -2.209 | -2.642 |
| | | Egln1 | -2.240 | -2.352 |
| | | Abcc8 | -2.273 | -2.690 |
| | | Aqp1 | -2.678 | -3.200 |
| | | Angpt1 | -4.454 | -4.113 |

**Supplemental Table 9. Fibrosis Genes**

GO Terms: Collagen, metallopeptidase, extracellular matrix, fibroblast, matrix, fibrosis

Genes Unique to FFAR4 KO Sham vs FFAR4 KO TAC: 10 increased, 3 decreased

| Gene ID | Fold Change |
|---|---|
| Igf1 | 2.787 |
| Vwa1 | 2.072 |
| Ero1l | 1.889 |
| Errfi1 | 1.882 |
| Ccdc80 | 1.882 |
| Bcl2 | 1.768 |
| Adamts9 | 1.760 |
| Ruvbl1 | 1.738 |
| Pxdn | 1.723 |
| Lum | 1.722 |
| Adamts3 | -1.755 |
| Dnajc19 | -1.763 |
| Jam3 | -1.792 |

Genes Unique to WT Sham vs WT TAC: 43 increased, 25 decreased

| Gene ID | Fold Change |
|---|---|
| Ereg | 240.348 |
| Adam8 | 117.349 |
| Timp1 | 112.492 |
| Arg1 | 94.628 |
| Pak3 | 32.821 |
| Ccl2 | 25.129 |
| Col11a1 | 17.389 |
| Ccna2 | 14.460 |
| Lgals3 | 13.711 |
| Myc | 13.703 |
| Loxl4 | 9.535 |
| Itgb2 | 9.094 |
| Ubash3b | 7.652 |
| Ptx3 | 7.589 |
| Ptgs2 | 7.153 |
| Pdpn | 6.605 |
| Pak1 | 6.585 |
| Sphk1 | 6.439 |
| Nlrc3 | 6.282 |
| Il17ra | 5.718 |
| Mmp19 | 5.480 |
| Fbn2 | 5.043 |
| Tgif1 | 4.348 |
| Itgb3 | 3.912 |
| Thy1 | 3.651 |
| Egr3 | 3.598 |
| Plau | 3.321 |
| Pik3cg | 3.245 |
| Itga11 | 3.174 |

Genes Shared by FFAR4 KO Sham vs FFAR4 KO TAC and WT Sham vs WT TAC: 98 increased, 26 decreased

| Gene ID | Fold Change KO | Fold Change WT |
|---|---|---|
| Lox | 24.312 | 34.324 |
| Serpine1 | 16.010 | 24.941 |
| Postn | 15.162 | 13.826 |
| Col1a1 | 11.043 | 11.083 |
| Col12a1 | 10.283 | 11.121 |
| Col3a1 | 10.165 | 9.226 |
| Col5a2 | 9.968 | 9.928 |
| Col8a1 | 9.567 | 9.110 |
| Cd44 | 9.335 | 9.640 |
| Tgfb2 | 8.828 | 10.745 |
| Loxl3 | 8.357 | 14.007 |
| Col1a2 | 8.165 | 7.647 |
| Fn1 | 7.686 | 8.424 |
| Loxl2 | 7.354 | 7.545 |
| Col5a1 | 6.935 | 6.841 |
| Thbs1 | 6.419 | 6.000 |
| Ctss | 6.082 | 6.240 |
| Fbn1 | 5.625 | 5.364 |
| Anxa2 | 5.166 | 6.442 |
| Lcp1 | 5.083 | 6.470 |
| Mrc2 | 4.616 | 3.066 |
| Mmp14 | 4.608 | 5.418 |
| Col16a1 | 4.513 | 3.752 |
| Emilin1 | 4.265 | 4.364 |
| Vim | 4.049 | 3.986 |
| Creb3l1 | 3.704 | 4.160 |
| Adamts2 | 3.608 | 3.687 |
| Fbln2 | 3.515 | 3.839 |
| Tgfbi | 3.493 | 3.399 |

| | |
|---|---|
| Bcl3 | 2.694 |
| Col27a1 | 2.693 |
| Pdgfc | 2.636 |
| Arrb2 | 2.484 |
| Tram2 | 2.447 |
| Mmp23 | 2.431 |
| D1Ertd622e | 2.416 |
| Cib1 | 2.066 |
| Bax | 2.038 |
| Fgfr1 | 1.934 |
| Gpm6b | 1.905 |
| Nfkb2 | 1.866 |
| Sulf2 | 1.768 |
| Adam15 | 1.726 |
| Dpt | -1.722 |
| Fzd4 | -1.750 |
| Gstp1 | -1.766 |
| Lims1 | -1.767 |
| Tek | -1.816 |
| Cd59a | -1.832 |
| Gpc1 | -1.893 |
| Kif16b | -1.898 |
| Dll4 | -1.979 |
| Abi3bp | -1.980 |
| Reck | -2.041 |
| Col4a3 | -2.048 |
| Pdgfd | -2.100 |
| Dand5 | -2.293 |
| Id1 | -2.367 |
| Smoc2 | -2.392 |
| Fgf9 | -2.474 |
| Has3 | -2.573 |
| Fuz | -2.725 |
| Wnt5a | -2.734 |
| Tcf15 | -2.877 |
| Enpp2 | -3.196 |
| Dpp4 | -3.837 |
| Egf | -4.469 |
| Fgf12 | -5.785 |

| | | |
|---|---|---|
| Sh3pxd2b | 3.486 | 3.501 |
| Adamts12 | 3.453 | 3.580 |
| Cd63 | 3.435 | 3.194 |
| Ifi30 | 3.347 | 3.281 |
| Mmp3 | 3.340 | 2.805 |
| Sfrp1 | 3.307 | 3.030 |
| Col4a1 | 3.301 | 3.032 |
| Tgfb3 | 3.271 | 2.029 |
| Col15a1 | 3.191 | 2.929 |
| Aebp1 | 3.139 | 3.664 |
| Eln | 2.913 | 2.003 |
| Zyx | 2.880 | 2.798 |
| Col14a1 | 2.859 | 2.153 |
| Serpinh1 | 2.807 | 2.799 |
| Col5a3 | 2.805 | 2.260 |
| Col18a1 | 2.799 | 3.149 |
| Col4a2 | 2.797 | 2.674 |
| Prkcd | 2.795 | 2.078 |
| Fosl2 | 2.782 | 3.120 |
| Adam9 | 2.775 | 2.992 |
| Rcn3 | 2.772 | 3.552 |
| Bak1 | 2.750 | 3.150 |
| Sox9 | 2.653 | 2.948 |
| Syk | 2.586 | 3.022 |
| Fscn1 | 2.554 | 1.949 |
| Plod3 | 2.509 | 2.553 |
| P3h1 | 2.506 | 2.958 |
| Iqgap1 | 2.488 | 2.517 |
| Cdkn1a | 2.459 | 2.314 |
| Mfap4 | 2.445 | 2.813 |
| Ilk | 2.436 | 2.613 |
| Sdc4 | 2.343 | 2.329 |
| Ddr1 | 2.343 | 3.180 |
| Ptk2b | 2.311 | 2.958 |
| Vcam1 | 2.304 | 2.086 |
| Col4a5 | 2.301 | 1.958 |
| Sulf1 | 2.265 | 2.514 |
| Itga4 | 2.244 | 2.883 |
| Itgb1 | 2.223 | 2.307 |
| Tnfrsf1a | 2.209 | 2.301 |

| | | |
|---|---|---|
| Olfml2b | 2.201 | 2.322 |
| Trp53 | 2.166 | 2.165 |
| Rcc2 | 2.146 | 2.043 |
| Vps33b | 2.137 | 1.953 |
| Emp2 | 2.111 | 2.255 |
| Hif1a | 2.108 | 2.022 |
| Ptprj | 2.104 | 1.777 |
| Hsd17b12 | 2.070 | 3.062 |
| Tnfrsf1b | 2.067 | 2.151 |
| P3h3 | 2.044 | 2.019 |
| Nras | 2.013 | 2.256 |
| Lamc1 | 2.000 | 1.813 |
| Sgpl1 | 1.982 | 2.148 |
| F2r | 1.915 | 2.004 |
| Tmem38b | 1.900 | 2.004 |
| Itgbl1 | 1.869 | 1.986 |
| Hyal2 | 1.853 | 1.806 |
| Actr3 | 1.840 | 2.003 |
| Itgav | 1.831 | 1.859 |
| Tgfb1 | 1.830 | 1.963 |
| Csgalnact1 | 1.827 | 1.919 |
| Tgfbr1 | 1.826 | 1.858 |
| Cdk6 | 1.801 | 1.781 |
| Colgalt1 | 1.798 | 1.726 |
| B4galt1 | 1.792 | 2.046 |
| Ctsb | 1.785 | 2.093 |
| Coro1c | 1.759 | 1.843 |
| Slc8a1 | 1.727 | 1.886 |
| Pmp22 | 1.714 | 1.736 |
| Clasp2 | -1.723 | -1.728 |
| Clasp1 | -1.751 | -2.046 |
| Bcl2l11 | -1.769 | -2.251 |
| Pex6 | -1.773 | -2.500 |
| Jun | -1.810 | -1.759 |
| Pdcd4 | -1.823 | -1.795 |
| Sod2 | -1.863 | -2.121 |
| Rgcc | -1.878 | -2.006 |
| Ndufs4 | -1.885 | -2.218 |
| Apbb1 | -1.896 | -1.712 |
| Dach1 | -1.897 | -2.799 |

| | | |
|---|---|---|
| Itgb6 | -1.918 | -2.285 |
| Tnxb | -1.937 | -3.175 |
| Idh2 | -1.945 | -2.544 |
| Tgfbr3 | -1.947 | -1.725 |
| Ecm2 | -2.033 | -2.536 |
| Ndnf | -2.038 | -1.973 |
| Vtn | -2.045 | -3.620 |
| Col4a4 | -2.072 | -2.081 |
| Fbln1 | -2.080 | -2.386 |
| Mmp15 | -2.237 | -2.565 |
| Cd200 | -2.288 | -2.106 |
| Pik3r1 | -2.401 | -2.446 |
| Timm21 | -2.440 | -1.991 |
| Egflam | -2.554 | -2.092 |
| Aqp1 | -2.678 | -3.200 |

**Supplemental Table 10. GPR Genes**
GO Terms: G-protein

Genes Unique to FFAR4 KO Sham vs FFAR4 KO TAC:
4 increased,
0 decreased

| Gene ID | Fold Change |
|---|---|
| Cx3cl1 | 2.987 |
| Adgrd1 | 2.575 |
| Akap12 | 2.225 |
| Prex1 | 1.716 |

Genes Unique to WT Sham vs WT TAC:
23 increased,
27 decreased

| Gene ID | Fold Change |
|---|---|
| Ccl2 | 25.129 |
| Cxcr6 | 18.350 |
| Ccr2 | 14.716 |
| Ccl7 | 12.622 |
| Gpr35 | 10.420 |
| Pik3r5 | 10.221 |
| Ccr1 | 9.871 |
| C5ar1 | 7.009 |
| Plek | 6.558 |
| C3ar1 | 6.280 |
| Vav1 | 5.911 |
| Pf4 | 5.217 |
| Ccr5 | 4.734 |
| Itgb3 | 3.912 |
| P2ry6 | 3.282 |
| Pik3cg | 3.245 |
| S1pr2 | 3.129 |
| Cx3cr1 | 2.848 |
| Fzd2 | 2.615 |
| Arrb2 | 2.484 |
| Rgs10 | 2.307 |
| Adgra3 | 2.035 |
| Gnai2 | 1.739 |
| Adgrl4 | -1.714 |
| Mgrn1 | -1.726 |
| Gpsm1 | -1.735 |
| Fzd4 | -1.750 |
| Rgs3 | -1.772 |
| Rapgef2 | -1.888 |
| Pde5a | -1.902 |
| Gkap1 | -1.945 |

Genes Shared by FFAR4 KO Sham vs FFAR4 KO TAC and WT Sham vs WT TAC:
19 increased, 17 decreased

| Gene ID | Fold Change KO | Fold Change WT |
|---|---|---|
| Ccl6 | 4.955 | 4.562 |
| Ccl9 | 4.518 | 5.177 |
| Tgm2 | 4.504 | 4.795 |
| Adcyap1r1 | 3.613 | 4.621 |
| Adcy7 | 2.959 | 2.815 |
| Anxa1 | 2.922 | 2.855 |
| Adgre1 | 2.830 | 3.153 |
| Syk | 2.586 | 3.022 |
| Lpar1 | 2.370 | 2.072 |
| Bcar1 | 2.279 | 1.778 |
| Itgb1 | 2.223 | 2.307 |
| Gprc5b | 2.020 | 1.770 |
| Lrp1 | 1.916 | 1.889 |
| F2r | 1.915 | 2.004 |
| Gnao1 | 1.861 | 2.401 |
| Gpr153 | 1.852 | 2.554 |
| Grk5 | 1.827 | 2.082 |
| Usp20 | 1.795 | 2.179 |
| Fzd1 | 1.728 | 2.331 |
| Sort1 | -1.710 | -1.915 |
| Palm | -1.805 | -1.909 |
| Ano1 | -1.849 | -2.281 |
| Agtr1a | -1.868 | -1.727 |
| Gpr157 | -1.896 | -2.042 |
| Lgr6 | -1.908 | -2.249 |
| Grm1 | -1.938 | -2.100 |
| Npr3 | -1.957 | -2.357 |
| Crhr2 | -2.026 | -1.926 |
| Rgs6 | -2.082 | -1.826 |
| Ric8b | -2.130 | -2.509 |
| Rgs5 | -2.151 | -2.708 |

| | |
|---|---|
| Prex2 | -1.984 |
| Dgke | -2.003 |
| Adgrf5 | -2.285 |
| Ackr3 | -2.301 |
| Fzd8 | -2.320 |
| Kctd12b | -2.449 |
| Ccrl2 | -2.506 |
| Camk2b | -2.688 |
| Wnt5a | -2.734 |
| P2ry14 | -2.855 |
| Rgs17 | -3.355 |
| Adcy1 | -3.363 |
| Celsr2 | -3.647 |
| Rgs7bp | -3.920 |
| Akap5 | -4.109 |
| Gnb3 | -5.439 |
| Ptgfr | -5.664 |
| Mrgprh | -7.156 |
| Celsr3 | -17.699 |

| | | |
|---|---|---|
| Adra1b | -2.439 | -2.145 |
| Rgs2 | -3.313 | -3.246 |
| Adra1a | -3.544 | -3.681 |
| P2ry1 | -4.190 | -5.821 |
| Gpr22 | -6.708 | -6.137 |

**Supplemental Table 11. Fatty Acid Metabolism Genes**
GO Terms: Fatty acid

Genes Unique to FFAR4 KO Sham vs FFAR4 KO TAC:
0 increased,
4 decreased

| Gene ID | Fold Change |
|---|---|
| Prkab2 | 1.747 |
| Mlxipl | -1.737 |
| Aig1 | -1.869 |
| Scd1 | -1.872 |

Genes Unique to WT Sham vs WT TAC:
11 increased,
17 decreased

| Gene ID | Fold Change |
|---|---|
| Ptgs2 | 7.153 |
| Brca1 | 6.587 |
| Tbxas1 | 4.509 |
| Aacs | 3.760 |
| Tlr2 | 3.624 |
| Hpgds | 3.154 |
| Pdk3 | 2.571 |
| Hacd4 | 2.187 |
| Acsl4 | 2.043 |
| Lpin2 | 2.023 |
| Plin2 | 1.801 |
| Decr1 | -1.703 |
| Asah2 | -1.724 |
| Acadsb | -1.735 |
| Echdc2 | -1.741 |
| Irs1 | -1.748 |
| Acadvl | -1.760 |
| Cbr4 | -1.783 |
| Echs1 | -1.816 |
| Akt2 | -1.847 |
| Acads | -1.875 |
| Alkbh7 | -1.918 |
| Hibch | -2.002 |
| Pnpla8 | -2.032 |
| Abcd2 | -2.278 |
| Acot1 | -3.193 |
| Ucp3 | -4.553 |
| Acsm5 | -5.715 |

Genes Shared by FFAR4 KO Sham vs FFAR4 KO TAC and WT Sham vs WT TAC:
15 increased, 40 decreased

| Gene ID | Fold Change KO | Fold Change WT |
|---|---|---|
| Thbs1 | 6.419 | 6.000 |
| Ldlr | 3.813 | 4.756 |
| Myo5a | 3.745 | 2.816 |
| Ankrd23 | 3.449 | 2.536 |
| Fads3 | 2.738 | 3.250 |
| Ptgis | 2.577 | 2.278 |
| Eif6 | 2.466 | 3.325 |
| Elovl1 | 2.273 | 2.978 |
| Nucb2 | 2.250 | 2.253 |
| Tlr4 | 2.125 | 2.071 |
| Por | 2.086 | 2.004 |
| Abcc1 | 2.004 | 1.837 |
| Sgpl1 | 1.982 | 2.148 |
| Xbp1 | 1.935 | 1.979 |
| Fads1 | 1.812 | 1.754 |
| Gpam | -1.714 | -2.049 |
| Lpl | -1.737 | -2.346 |
| Lipe | -1.738 | -1.839 |
| Abcd3 | -1.744 | -1.814 |
| Tecrl | -1.746 | -1.709 |
| Hadha | -1.757 | -2.090 |
| Acat1 | -1.762 | -2.006 |
| Acot11 | -1.764 | -1.847 |
| Acaa2 | -1.772 | -1.909 |
| Phyh | -1.774 | -1.885 |
| Auh | -1.775 | -1.982 |
| Etfb | -1.791 | -1.949 |
| Aldh5a1 | -1.853 | -2.780 |
| Ivd | -1.885 | -2.559 |

| | | |
|---|---|---|
| Crat | -1.899 | -2.050 |
| Etfdh | -1.923 | -2.156 |
| Tnxb | -1.937 | -3.175 |
| Acot2 | -1.965 | -2.003 |
| Eci1 | -1.981 | -2.429 |
| Cpt2 | -2.030 | -2.238 |
| Etfa | -2.081 | -2.236 |
| Hadhb | -2.095 | -2.290 |
| Gcdh | -2.145 | -2.427 |
| Acadm | -2.161 | -2.786 |
| Ech1 | -2.223 | -2.511 |
| Acacb | -2.284 | -2.644 |
| Hadh | -2.364 | -2.757 |
| Plin5 | -2.386 | -2.429 |
| Mlycd | -2.415 | -2.520 |
| Ptgds | -2.433 | -3.263 |
| Acad11 | -2.509 | -2.647 |
| Ppargc1a | -2.536 | -3.388 |
| C1qtnf9 | -2.573 | -3.581 |
| Ppara | -2.614 | -2.691 |
| Acsl1 | -2.638 | -3.018 |
| Tecr | -2.723 | -2.728 |
| Acsl6 | -2.754 | -3.114 |
| Lpin1 | -2.759 | -2.924 |
| Dgat2 | -2.763 | -2.645 |
| Ces1d | -6.513 | -9.549 |

**Supplemental Table 11. Fatty Acid Metabolism Genes**
GO Terms: Fatty acid

Genes Unique to FFAR4 KO Sham vs FFAR4 KO TAC: 0 increased, 4 decreased

| Gene ID | Fold Change |
|---|---|
| Prkab2 | 1.747 |
| Mlxipl | -1.737 |
| Aig1 | -1.869 |
| Scd1 | -1.872 |

Genes Unique to WT Sham vs WT TAC: 11 increased, 17 decreased

| Gene ID | Fold Change |
|---|---|
| Ptgs2 | 7.153 |
| Brca1 | 6.587 |
| Tbxas1 | 4.509 |
| Aacs | 3.760 |
| Tlr2 | 3.624 |
| Hpgds | 3.154 |
| Pdk3 | 2.571 |
| Hacd4 | 2.187 |
| Acsl4 | 2.043 |
| Lpin2 | 2.023 |
| Plin2 | 1.801 |
| Decr1 | -1.703 |
| Asah2 | -1.724 |
| Acadsb | -1.735 |
| Echdc2 | -1.741 |
| Irs1 | -1.748 |
| Acadvl | -1.760 |
| Cbr4 | -1.783 |
| Echs1 | -1.816 |
| Akt2 | -1.847 |
| Acads | -1.875 |
| Alkbh7 | -1.918 |
| Hibch | -2.002 |
| Pnpla8 | -2.032 |
| Abcd2 | -2.278 |
| Acot1 | -3.193 |
| Ucp3 | -4.553 |
| Acsm5 | -5.715 |

Genes Shared by FFAR4 KO Sham vs FFAR4 KO TAC and WT Sham vs WT TAC: 15 increased, 40 decreased

| Gene ID | Fold Change KO | Fold Change WT |
|---|---|---|
| Thbs1 | 6.419 | 6.000 |
| Ldlr | 3.813 | 4.756 |
| Myo5a | 3.745 | 2.816 |
| Ankrd23 | 3.449 | 2.536 |
| Fads3 | 2.738 | 3.250 |
| Ptgis | 2.577 | 2.278 |
| Eif6 | 2.466 | 3.325 |
| Elovl1 | 2.273 | 2.978 |
| Nucb2 | 2.250 | 2.253 |
| Tlr4 | 2.125 | 2.071 |
| Por | 2.086 | 2.004 |
| Abcc1 | 2.004 | 1.837 |
| Sgpl1 | 1.982 | 2.148 |
| Xbp1 | 1.935 | 1.979 |
| Fads1 | 1.812 | 1.754 |
| Gpam | -1.714 | -2.049 |
| Lpl | -1.737 | -2.346 |
| Lipe | -1.738 | -1.839 |
| Abcd3 | -1.744 | -1.814 |
| Tecrl | -1.746 | -1.709 |
| Hadha | -1.757 | -2.090 |
| Acat1 | -1.762 | -2.006 |
| Acot11 | -1.764 | -1.847 |
| Acaa2 | -1.772 | -1.909 |
| Phyh | -1.774 | -1.885 |
| Auh | -1.775 | -1.982 |
| Etfb | -1.791 | -1.949 |
| Aldh5a1 | -1.853 | -2.780 |
| Ivd | -1.885 | -2.559 |

| | | |
|---|---|---|
| Crat | -1.899 | -2.050 |
| Etfdh | -1.923 | -2.156 |
| Tnxb | -1.937 | -3.175 |
| Acot2 | -1.965 | -2.003 |
| Eci1 | -1.981 | -2.429 |
| Cpt2 | -2.030 | -2.238 |
| Etfa | -2.081 | -2.236 |
| Hadhb | -2.095 | -2.290 |
| Gcdh | -2.145 | -2.427 |
| Acadm | -2.161 | -2.786 |
| Ech1 | -2.223 | -2.511 |
| Acacb | -2.284 | -2.644 |
| Hadh | -2.364 | -2.757 |
| Plin5 | -2.386 | -2.429 |
| Mlycd | -2.415 | -2.520 |
| Ptgds | -2.433 | -3.263 |
| Acad11 | -2.509 | -2.647 |
| Ppargc1a | -2.536 | -3.388 |
| C1qtnf9 | -2.573 | -3.581 |
| Ppara | -2.614 | -2.691 |
| Acsl1 | -2.638 | -3.018 |
| Tecr | -2.723 | -2.728 |
| Acsl6 | -2.754 | -3.114 |
| Lpin1 | -2.759 | -2.924 |
| Dgat2 | -2.763 | -2.645 |
| Ces1d | -6.513 | -9.549 |

**Supplemental Table 13. LC/MS/MS** detection and quantitation of oxylipins.

| COMPOUND | PRECURSOR ION | PRODUCT ION | DWELL TIME (SEC) | RETENTION TIME (MINS) | CONE (V) | CE (EV) | LOD (NM) | LOQ (NM) | INTERNAL STANDARD |
|---|---|---|---|---|---|---|---|---|---|
| 9-HOTRE | 293.1 | 171.1 | 0.08 | 6.37 | 36 | 16 | 0.29 | 0.89 | 9-HODE d4 |
| 9-KODE | 293.1 | 185.1 | 0.02 | 7.43 | 34 | 16 | 0.49 | 1.48 | 9-HODE d4 |
| 13-KODE | 293.1 | 195.1 | 0.02 | 7.29 | 34 | 16 | 0.39 | 1.18 | 9-HODE d4 |
| 13-HOTRE | 293.1 | 195.2 | 0.08 | 6.46 | 36 | 16 | 0.38 | 1.14 | 9-HODE d4 |
| 9-HODE | 295.1 | 171.1 | 0.02 | 7.07 | 34 | 14 | 0.14 | 0.43 | 9-HODE d4 |
| 9(10)-EPOME | 295.1 | 171.1 | 0.03 | 7.99 | 35 | 14 | 0.27 | 0.81 | 9(10)-EpOME d4 |
| 13-HODE | 295.1 | 195.1 | 0.03 | 7.03 | 34 | 14 | 0.10 | 0.31 | 9-HODE d4 |
| 12(13)-EPOME | 295.1 | 195.1 | 0.03 | 7.93 | 35 | 14 | 0.16 | 0.48 | 9(10)-EpOME d4 |
| 9-HODE D4 | 299.1 | 172.1 | 0.02 | 7.04 | 34 | 16 | 0.33 | 0.99 | - |
| 9(10)-EPOME D4 | 299.2 | 172.1 | 0.03 | 7.95 | 35 | 14 | 0.43 | 1.31 | - |
| 9(10)-DIHOME | 313.1 | 201.1 | 0.05 | 5.79 | 28 | 8 | 2.07 | 6.28 | 9(10)-EpOME d4 |
| 15-KETE | 317.1 | 113.1 | 0.02 | 7.34 | 34 | 14 | 0.27 | 0.81 | 12-HETE d8 |
| 5-HEPE | 317.1 | 115.1 | 0.04 | 6.92 | 28 | 16 | 0.60 | 1.81 | 12-HETE d8 |
| 11-HEPE | 317.1 | 121.1 | 0.05 | 6.67 | 28 | 16 | 0.46 | 1.39 | 12-HETE d8 |
| 8-HEPE | 317.1 | 127.1 | 0.05 | 6.74 | 28 | 16 | 0.79 | 2.39 | 12-HETE d8 |
| 9-HEPE | 317.1 | 149.1 | 0.04 | 6.81 | 28 | 16 | 0.34 | 1.02 | 12-HETE d8 |
| 12-HEPE | 317.1 | 179.1 | 0.05 | 6.77 | 28 | 16 | 0.27 | 0.81 | 12-HETE d8 |
| 5-KETE | 317.1 | 203.1 | 0.03 | 7.8 | 34 | 14 | 0.42 | 1.27 | 12-HETE d8 |
| 11(12)-EPETE | 317.1 | 208.1 | 0.02 | 7.45 | 32 | 10 | 0.41 | 1.23 | 14,15-EET d11 |
| 15-HEPE | 317.1 | 219.1 | 0.05 | 6.63 | 28 | 16 | 0.49 | 1.49 | 12-HETE d8 |
| 17(18)-EPETE | 317.1 | 259.4 | 0.02 | 7.2 | 32 | 10 | 1.88 | 5.71 | 14,15-EET d11 |
| 12-KETE | 317.1 | 273.2 | 0.02 | 7.54 | 34 | 14 | 0.63 | 1.90 | 12-HETE d8 |
| 8(9)-EPETE | 317.1 | 299.1 | 0.02 | 7.49 | 32 | 10 | 1.15 | 3.48 | 14,15-EET d11 |
| 18-HEPE | 317.2 | 215.2 | 0.08 | 6.46 | 34 | 12 | 0.20 | 0.60 | 12-HETE d8 |
| 14(15)-EPETE | 317.2 | 247.3 | 0.02 | 7.4 | 32 | 10 | 1.15 | 3.47 | 14,15-EET d11 |
| 12-HETE | 319.0 | 179.1 | 0.02 | 7.43 | 32 | 14 | 0.63 | 1.89 | 12-HETE d8 |
| 5-HETE | 319.1 | 114.9 | 0.08 | 7.64 | 32 | 14 | 0.29 | 0.88 | 12-HETE d8 |
| 9-HETE | 319.1 | 123.0 | 0.02 | 7.54 | 32 | 14 | 0.22 | 0.67 | 12-HETE d8 |
| 15-HETE | 319.1 | 219.0 | 0.02 | 7.17 | 32 | 14 | 0.26 | 0.79 | 12-HETE d8 |
| 8(9)-EPETRE | 319.1 | 155.1 | 0.20 | 8.23 | 30 | 10 | 0.09 | 0.28 | 14,15-EET d11 |
| 11(12)-EPETRE | 319.1 | 208.1 | 0.03 | 8.16 | 30 | 10 | 0.48 | 1.44 | 14,15-EET d11 |
| 14(15)-EPETRE | 319.1 | 219.1 | 0.03 | 7.93 | 30 | 10 | 0.43 | 1.29 | 14,15-EET d11 |
| 12-HETE D8 | 327.2 | 184.1 | 0.02 | 7.38 | 35 | 14 | 3.35 | 10.15 | - |
| 14,15-EET D11 | 330.1 | 219.0 | 0.03 | 7.88 | 30 | 10 | 0.35 | 1.05 | - |
| 12-HPETE | 335.1 | 153.1 | 0.02 | 7.54 | 20 | 8 | 0.64 | 1.94 | 12-HETE d8 |
| 15-HPETE | 335.1 | 219.1 | 0.02 | 7.33 | 20 | 6 | 0.58 | 1.77 | 12-HETE d8 |
| 11(12)-DIHETE | 335.2 | 167.2 | 0.05 | 5.53 | 25 | 8 | 2.53 | 7.66 | 14,15-EET d11 |
| 14(15)-DIHETE | 335.2 | 207.2 | 0.05 | 5.46 | 25 | 8 | 0.07 | 0.21 | 14,15-EET d11 |
| 17(18)-DIHETE | 335.2 | 247.2 | 0.11 | 5.29 | 25 | 8 | 0.54 | 1.62 | 14,15-EET d11 |
| 8(9)-DIHETE | 335.2 | 317.2 | 0.05 | 5.61 | 25 | 8 | 1.58 | 4.78 | 14,15-EET d11 |
| 8(9)-DIHETRE | 337.1 | 127.1 | 0.08 | 6.38 | 28 | 8 | 1.21 | 3.68 | 14,15-EET d11 |
| 11(12)-DIHETRE | 337.1 | 167.1 | 0.08 | 6.19 | 28 | 8 | 1.83 | 5.54 | 14,15-EET d11 |
| 14(15)-DIHETRE | 337.1 | 207.1 | 0.16 | 5.96 | 28 | 8 | 0.64 | 1.95 | 14,15-EET d11 |
| CUDA | 340.2 | 214.1 | 0.05 | 5.82 | 34 | 12 | - | - | - |
| 7(8)-EPDPE | 343.1 | 189.1 | 0.03 | 8.18 | 34 | 10 | 1.41 | 4.27 | 12-HETE d8 |
| 13(14)-EPDPE | 343.1 | 193.1 | 0.03 | 8.05 | 34 | 10 | 0.58 | 1.77 | 12-HETE d8 |
| 16(17)-EPDPE | 343.1 | 233.1 | 0.03 | 8.02 | 34 | 10 | 0.21 | 0.62 | 12-HETE d8 |
| 19(20)-EPDPE | 343.1 | 281.1 | 0.03 | 7.81 | 34 | 10 | 0.28 | 0.85 | 12-HETE d8 |
| 10(11)-EPDPE | 343.1 | 299.1 | 0.03 | 8.18 | 34 | 10 | 0.74 | 2.26 | 12-HETE d8 |
| 4-HDOHE | 343.2 | 101.1 | 0.03 | 7.81 | 35 | 10 | 0.54 | 1.65 | 12-HETE d8 |
| 8-HDOHE | 343.2 | 109.1 | 0.02 | 7.56 | 35 | 10 | 0.15 | 0.47 | 12-HETE d8 |
| 7-HDOHE | 343.2 | 141.1 | 0.02 | 7.51 | 35 | 10 | 0.59 | 1.78 | 12-HETE d8 |
| 11-HDOHE | 343.2 | 149.1 | 0.02 | 7.45 | 35 | 10 | 0.59 | 1.80 | 12-HETE d8 |
| 10-HDOHE | 343.2 | 181.1 | 0.02 | 7.37 | 35 | 10 | 0.74 | 2.24 | 12-HETE d8 |
| 14-HDOHE | 343.2 | 205.1 | 0.02 | 7.36 | 35 | 10 | 0.23 | 0.70 | 12-HETE d8 |
| 13-HDOHE | 343.2 | 221.1 | 0.02 | 7.3 | 35 | 10 | 0.45 | 1.36 | 12-HETE d8 |
| 16-HDOHE | 343.2 | 233.1 | 0.02 | 7.22 | 35 | 12 | 0.85 | 2.58 | 12-HETE d8 |
| 20-HDOHE | 343.2 | 241.1 | 0.02 | 7.06 | 35 | 12 | 0.43 | 1.30 | 12-HETE d8 |
| 17-HDOHE | 343.2 | 245.1 | 0.02 | 7.24 | 35 | 12 | 0.27 | 0.83 | 12-HETE d8 |
| 22-HDOHE | 343.2 | 281.1 | 0.04 | 6.96 | 35 | 12 | 0.66 | 2.00 | 12-HETE d8 |
| RESOLVIN D1 | 375.1 | 141.1 | 0.16 | 4.15 | 34 | 16 | 0.06 | 0.20 | 14,15-EET d11 |

**Supplemental Table 14.** Time dependent changes in myocyte oxylipins, by pool.

| Fraction | Compartment | pFA | Chemistry | Oxylipin | Time | *Myocytes* WT (CI) | KO (CI) | *Interaction* Prob>\|t\| | *Test for difference at each time* %Difference (CI) | Prob>\|t\| |
|---|---|---|---|---|---|---|---|---|---|---|
| Esterified | Cell | LA | Alcohol | 9-HODE | 0 | 15 (8, 25) | 12 (7, 21) | 0.76 | -17% (-74, 159) | >0.80 |
| Esterified | Cell | LA | Alcohol | 9-HODE | 15 | 20 (12, 34) | 16 (10, 27) | | -20% (-69, 109) | >0.80 |
| Esterified | Cell | LA | Alcohol | 9-HODE | 30 | 26 (15, 43) | 20 (12, 33) | | -23% (-69, 94) | >0.80 |
| Esterified | Cell | LA | Alcohol | 9-HODE | 60 | 28 (16, 50) | 20 (11, 36) | | -28% (-80, 155) | >0.80 |
| NEOx | Cell | LA | Alcohol | 9-HODE | 0 | 1.15 (0.63, 2.11) | 1.02 (0.56, 1.88) | >0.80 | -11% (-75, 217) | >0.80 |
| NEOx | Cell | LA | Alcohol | 9-HODE | 15 | 1.13 (0.69, 1.86) | 0.97 (0.59, 1.6) | | -14% (-68, 129) | >0.80 |
| NEOx | Cell | LA | Alcohol | 9-HODE | 30 | 1.12 (0.66, 1.88) | 0.93 (0.55, 1.56) | | -17% (-66, 107) | >0.80 |
| NEOx | Cell | LA | Alcohol | 9-HODE | 60 | 1.1 (0.57, 2.11) | 0.86 (0.45, 1.65) | | -22% (-82, 237) | >0.80 |
| Esterified | Media | LA | Alcohol | 9-HODE | 0 | 5.01 (3.09, 8.11) | 4.02 (2.48, 6.51) | >0.80 | -20% (-70, 114) | >0.80 |
| Esterified | Media | LA | Alcohol | 9-HODE | 15 | 4.85 (3.02, 7.78) | 3.91 (2.44, 6.27) | | -19% (-67, 95) | >0.80 |
| Esterified | Media | LA | Alcohol | 9-HODE | 30 | 4.9 (3.06, 7.84) | 3.97 (2.48, 6.35) | | -19% (-66, 91) | >0.80 |
| Esterified | Media | LA | Alcohol | 9-HODE | 60 | 5.65 (3.45, 9.25) | 4.63 (2.83, 7.58) | | -18% (-71, 135) | >0.80 |
| NEOx | Media | LA | Alcohol | 9-HODE | 0 | 0.6 (0.31, 1.15) | 0.52 (0.27, 1) | >0.80 | -13% (-78, 239) | >0.80 |
| NEOx | Media | LA | Alcohol | 9-HODE | 15 | 0.86 (0.48, 1.55) | 0.73 (0.41, 1.31) | | -15% (-73, 161) | >0.80 |
| NEOx | Media | LA | Alcohol | 9-HODE | 30 | 1.04 (0.58, 1.89) | 0.86 (0.47, 1.55) | | -18% (-72, 140) | >0.80 |
| NEOx | Media | LA | Alcohol | 9-HODE | 60 | 0.9 (0.45, 1.78) | 0.7 (0.35, 1.39) | | -22% (-83, 256) | >0.80 |
| Esterified | Cell | LA | Alcohol | 13-HODE | 0 | 18 (11, 31) | 15 (9, 25) | 0.53 | -19% (-74, 149) | >0.80 |
| Esterified | Cell | LA | Alcohol | 13-HODE | 15 | 23 (14, 39) | 18 (11, 29) | | -24% (-71, 96) | >0.80 |
| Esterified | Cell | LA | Alcohol | 13-HODE | 30 | 28 (17, 46) | 20 (12, 33) | | -29% (-72, 77) | >0.80 |
| Esterified | Cell | LA | Alcohol | 13-HODE | 60 | 32 (19, 57) | 20 (11, 35) | | -38% (-82, 113) | >0.80 |
| NEOx | Cell | LA | Alcohol | 13-HODE | 0 | 1.67 (0.94, 2.96) | 1.24 (0.7, 2.19) | 0.39 | -26% (-77, 141) | >0.80 |
| NEOx | Cell | LA | Alcohol | 13-HODE | 15 | 1.76 (1.19, 2.6) | 1.13 (0.77, 1.68) | | -36% (-71, 41) | 0.54 |
| NEOx | Cell | LA | Alcohol | 13-HODE | 30 | 1.93 (1.25, 2.99) | 1.08 (0.7, 1.67) | | -44% (-72, 11) | 0.13 |
| NEOx | Cell | LA | Alcohol | 13-HODE | 60 | 2.62 (1.39, 4.93) | 1.11 (0.59, 2.09) | | -58% (-90, 75) | 0.45 |
| Esterified | Media | LA | Alcohol | 13-HODE | 0 | 6.14 (3.53, 10.69) | 5.25 (3.02, 9.14) | 0.61 | -15% (-71, 148) | >0.80 |
| Esterified | Media | LA | Alcohol | 13-HODE | 15 | 5.57 (3.18, 9.76) | 4.9 (2.8, 8.59) | | -12% (-68, 144) | >0.80 |
| Esterified | Media | LA | Alcohol | 13-HODE | 30 | 5.46 (3.12, 9.53) | 4.94 (2.83, 8.64) | | -9% (-67, 149) | >0.80 |
| Esterified | Media | LA | Alcohol | 13-HODE | 60 | 6.6 (3.79, 11.48) | 6.34 (3.64, 11.03) | | -4% (-68, 188) | >0.80 |
| NEOx | Media | LA | Alcohol | 13-HODE | 0 | 0.45 (0.25, 0.83) | 0.92 (0.5, 1.69) | 0.35 | 105% (-38, 572) | 0.46 |
| NEOx | Media | LA | Alcohol | 13-HODE | 15 | 0.94 (0.69, 1.28) | 1.6 (1.17, 2.18) | | 71% (-5, 206) | 0.08 |
| NEOx | Media | LA | Alcohol | 13-HODE | 30 | 1.43 (0.95, 2.15) | 2.04 (1.36, 3.06) | | 43% (0, 103) | 0.05 |
| NEOx | Media | LA | Alcohol | 13-HODE | 60 | 1.33 (0.67, 2.65) | 1.32 (0.66, 2.63) | | -1% (-78, 349) | >0.80 |
| Esterified | Cell | aLA | Alcohol | 9-HOTrE | 0 | 0.48 (0.22, 1.04) | 0.15 (0.07, 0.33) | 0.18 | -68% (-93, 51) | 0.24 |
| Esterified | Cell | aLA | Alcohol | 9-HOTrE | 15 | 0.66 (0.42, 1.04) | 0.29 (0.19, 0.46) | | -56% (-82, 9) | 0.09 |
| Esterified | Cell | aLA | Alcohol | 9-HOTrE | 30 | 0.73 (0.42, 1.27) | 0.45 (0.26, 0.77) | | -39% (-70, 23) | 0.28 |
| Esterified | Cell | aLA | Alcohol | 9-HOTrE | 60 | 0.46 (0.19, 1.09) | 0.53 (0.22, 1.26) | | 16% (-83, 692) | >0.80 |
| NEOx | Cell | aLA | Alcohol | 9-HOTrE | 0 | 0.17 (0.07, 0.44) | 0.12 (0.05, 0.31) | >0.80 | -29% (-90, 410) | >0.80 |
| NEOx | Cell | aLA | Alcohol | 9-HOTrE | 15 | 0.15 (0.07, 0.34) | 0.11 (0.05, 0.25) | | -26% (-85, 259) | >0.80 |
| NEOx | Cell | aLA | Alcohol | 9-HOTrE | 30 | 0.15 (0.07, 0.34) | 0.12 (0.05, 0.27) | | -22% (-82, 243) | >0.80 |
| NEOx | Cell | aLA | Alcohol | 9-HOTrE | 60 | 0.2 (0.07, 0.54) | 0.17 (0.06, 0.46) | | -14% (-91, 706) | >0.80 |
| Esterified | Media | aLA | Alcohol | 9-HOTrE | 0 | 0.5 (0.23, 1.1) | 0.21 (0.1, 0.47) | 0.34 | -57% (-91, 114) | 0.60 |
| Esterified | Media | aLA | Alcohol | 9-HOTrE | 15 | 0.52 (0.31, 0.88) | 0.28 (0.17, 0.47) | | -47% (-81, 51) | 0.45 |
| Esterified | Media | aLA | Alcohol | 9-HOTrE | 30 | 0.54 (0.3, 0.98) | 0.36 (0.2, 0.65) | | -34% (-73, 61) | 0.74 |
| Esterified | Media | aLA | Alcohol | 9-HOTrE | 60 | 0.57 (0.24, 1.37) | 0.59 (0.25, 1.41) | | 3% (-85, 627) | >0.80 |
| NEOx | Media | aLA | Alcohol | 9-HOTrE | 0 | 0.26 (0.14, 0.51) | 0.2 (0.1, 0.38) | 0.56 | -25% (-81, 189) | >0.80 |
| NEOx | Media | aLA | Alcohol | 9-HOTrE | 15 | 0.21 (0.13, 0.35) | 0.17 (0.11, 0.29) | | -18% (-69, 121) | >0.80 |
| NEOx | Media | aLA | Alcohol | 9-HOTrE | 30 | 0.18 (0.11, 0.31) | 0.17 (0.1, 0.28) | | -9% (-63, 123) | >0.80 |
| NEOx | Media | aLA | Alcohol | 9-HOTrE | 60 | 0.18 (0.09, 0.37) | 0.2 (0.1, 0.41) | | 12% (-77, 446) | >0.80 |
| Esterified | Cell | aLA | Alcohol | 13-HOTrE | 0 | 0.6 (0.22, 1.63) | 0.39 (0.14, 1.05) | >0.80 | -35% (-92, 428) | >0.80 |
| Esterified | Cell | aLA | Alcohol | 13-HOTrE | 15 | 0.64 (0.29, 1.41) | 0.41 (0.19, 0.91) | | -36% (-87, 208) | >0.80 |
| Esterified | Cell | aLA | Alcohol | 13-HOTrE | 30 | 0.69 (0.3, 1.59) | 0.44 (0.19, 1.02) | | -36% (-85, 169) | >0.80 |
| Esterified | Cell | aLA | Alcohol | 13-HOTrE | 60 | 0.8 (0.27, 2.38) | 0.51 (0.17, 1.51) | | -36% (-94, 633) | >0.80 |
| NEOx | Cell | aLA | Alcohol | 13-HOTrE | 0 | 0.56 (0.23, 1.34) | 0.39 (0.16, 0.94) | 0.15 | -29% (-89, 342) | >0.80 |
| NEOx | Cell | aLA | Alcohol | 13-HOTrE | 15 | 0.41 (0.2, 0.86) | 0.21 (0.1, 0.44) | | -48% (-88, 119) | 0.75 |
| NEOx | Cell | aLA | Alcohol | 13-HOTrE | 30 | 0.4 (0.19, 0.86) | 0.15 (0.07, 0.33) | | -62% (-90, 46) | 0.26 |
| NEOx | Cell | aLA | Alcohol | 13-HOTrE | 60 | 0.91 (0.36, 2.32) | 0.18 (0.07, 0.47) | | -80% (-98, 64) | 0.21 |
| Esterified | Media | aLA | Alcohol | 13-HOTrE | 0 | 0.35 (0.16, 0.72) | 0.2 (0.09, 0.41) | 0.33 | -43% (-88, 168) | >0.80 |
| Esterified | Media | aLA | Alcohol | 13-HOTrE | 15 | 0.46 (0.25, 0.87) | 0.31 (0.17, 0.58) | | -33% (-80, 128) | >0.80 |
| Esterified | Media | aLA | Alcohol | 13-HOTrE | 30 | 0.53 (0.28, 1.02) | 0.43 (0.22, 0.82) | | -20% (-74, 151) | >0.80 |
| Esterified | Media | aLA | Alcohol | 13-HOTrE | 60 | 0.44 (0.2, 0.96) | 0.5 (0.22, 1.09) | | 14% (-81, 571) | >0.80 |
| NEOx | Media | aLA | Alcohol | 13-HOTrE | 0 | 0.14 (0.05, 0.36) | 0.38 (0.14, 1.01) | 0.29 | 179% (-64, 2042) | 0.65 |
| NEOx | Media | aLA | Alcohol | 13-HOTrE | 15 | 0.15 (0.07, 0.35) | 0.33 (0.15, 0.76) | | 117% (-57, 986) | 0.70 |
| NEOx | Media | aLA | Alcohol | 13-HOTrE | 30 | 0.18 (0.08, 0.42) | 0.31 (0.13, 0.72) | | 69% (-63, 665) | >0.80 |
| NEOx | Media | aLA | Alcohol | 13-HOTrE | 60 | 0.28 (0.1, 0.79) | 0.29 (0.1, 0.81) | | 3% (-90, 951) | >0.80 |
| Esterified | Cell | AA | Alcohol | 5-HETE | 0 | 15 (7, 32) | 11 (5, 23) | 0.57 | -27% (-85, 253) | >0.80 |
| Esterified | Cell | AA | Alcohol | 5-HETE | 15 | 20 (9, 42) | 14 (6, 29) | | -31% (-83, 185) | >0.80 |
| Esterified | Cell | AA | Alcohol | 5-HETE | 30 | 25 (12, 53) | 16 (7, 34) | | -36% (-84, 157) | >0.80 |
| Esterified | Cell | AA | Alcohol | 5-HETE | 60 | 31 (14, 69) | 17 (8, 38) | | -44% (-90, 197) | >0.80 |
| NEOx | Cell | AA | Alcohol | 5-HETE | 0 | 0.15 (0.1, 0.24) | 0.13 (0.08, 0.2) | 0.46 | -16% (-67, 119) | >0.80 |
| NEOx | Cell | AA | Alcohol | 5-HETE | 15 | 0.19 (0.13, 0.29) | 0.15 (0.1, 0.23) | | -22% (-64, 72) | >0.80 |
| NEOx | Cell | AA | Alcohol | 5-HETE | 30 | 0.22 (0.15, 0.34) | 0.16 (0.11, 0.24) | | -27% (-65, 54) | >0.80 |
| NEOx | Cell | AA | Alcohol | 5-HETE | 60 | 0.22 (0.14, 0.36) | 0.14 (0.09, 0.23) | | -37% (-78, 83) | 0.79 |
| Esterified | Media | AA | Alcohol | 5-HETE | 0 | 0.42 (0.2, 0.88) | 0.33 (0.16, 0.69) | >0.80 | -21% (-83, 256) | >0.80 |
| Esterified | Media | AA | Alcohol | 5-HETE | 15 | 0.4 (0.2, 0.81) | 0.31 (0.15, 0.63) | | -23% (-79, 190) | >0.80 |
| Esterified | Media | AA | Alcohol | 5-HETE | 30 | 0.41 (0.2, 0.83) | 0.31 (0.15, 0.63) | | -24% (-79, 174) | >0.80 |
| Esterified | Media | AA | Alcohol | 5-HETE | 60 | 0.51 (0.24, 1.09) | 0.38 (0.18, 0.81) | | -26% (-86, 285) | >0.80 |
| NEOx | Media | AA | Alcohol | 5-HETE | 0 | 0.16 (0.09, 0.26) | 0.13 (0.08, 0.22) | 0.27 | -14% (-70, 142) | >0.80 |
| NEOx | Media | AA | Alcohol | 5-HETE | 15 | 0.16 (0.1, 0.26) | 0.15 (0.09, 0.24) | | -5% (-62, 139) | >0.80 |
| NEOx | Media | AA | Alcohol | 5-HETE | 30 | 0.16 (0.1, 0.25) | 0.16 (0.1, 0.27) | | 5% (-57, 157) | >0.80 |
| NEOx | Media | AA | Alcohol | 5-HETE | 60 | 0.15 (0.09, 0.26) | 0.2 (0.12, 0.33) | | 27% (-58, 289) | >0.80 |

| | | | | | | | | | | |
|---|---|---|---|---|---|---|---|---|---|---|
| Esterified | Cell | AA | Alcohol | 9-HETE | 0 | 1.02 (0.38, 2.75) | 0.6 (0.22, 1.62) | 0.65 | -41% (-93, 362) | >0.80 |
| Esterified | Cell | AA | Alcohol | 9-HETE | 15 | 0.89 (0.42, 1.89) | 0.47 (0.22, 0.99) | | -48% (-88, 133) | 0.79 |
| Esterified | Cell | AA | Alcohol | 9-HETE | 30 | 0.96 (0.43, 2.15) | 0.45 (0.2, 1) | | -53% (-88, 80) | 0.53 |
| Esterified | Cell | AA | Alcohol | 9-HETE | 60 | 2.1 (0.72, 6.16) | 0.77 (0.26, 2.27) | | -63% (-97, 313) | >0.80 |
| NEOx | Cell | AA | Alcohol | 9-HETE | 0 | 0.32 (0.18, 0.59) | 0.3 (0.16, 0.55) | 0.62 | -6% (-73, 221) | >0.80 |
| NEOx | Cell | AA | Alcohol | 9-HETE | 15 | 0.48 (0.33, 0.71) | 0.41 (0.28, 0.61) | | -14% (-60, 86) | >0.80 |
| NEOx | Cell | AA | Alcohol | 9-HETE | 30 | 0.61 (0.39, 0.95) | 0.48 (0.31, 0.75) | | -21% (-59, 49) | >0.80 |
| NEOx | Cell | AA | Alcohol | 9-HETE | 60 | 0.6 (0.3, 1.17) | 0.39 (0.2, 0.77) | | -34% (-85, 195) | >0.80 |
| Esterified | Media | AA | Alcohol | 9-HETE | 0 | 0.4 (0.21, 0.76) | 0.41 (0.21, 0.77) | 0.07 | 1% (-73, 283) | >0.80 |
| Esterified | Media | AA | Alcohol | 9-HETE | 15 | 0.48 (0.26, 0.87) | 0.38 (0.21, 0.69) | | -21% (-75, 150) | >0.80 |
| Esterified | Media | AA | Alcohol | 9-HETE | 30 | 0.54 (0.3, 1) | 0.34 (0.19, 0.62) | | -38% (-79, 88) | >0.80 |
| Esterified | Media | AA | Alcohol | 9-HETE | 60 | 0.63 (0.32, 1.23) | 0.24 (0.12, 0.47) | | -61% (-91, 66) | 0.36 |
| NEOx | Media | AA | Alcohol | 9-HETE | 0 | 0.35 (0.16, 0.76) | 0.39 (0.18, 0.84) | >0.80 | 11% (-78, 452) | >0.80 |
| NEOx | Media | AA | Alcohol | 9-HETE | 15 | 0.4 (0.19, 0.83) | 0.44 (0.21, 0.93) | | 12% (-72, 356) | >0.80 |
| NEOx | Media | AA | Alcohol | 9-HETE | 30 | 0.39 (0.19, 0.83) | 0.45 (0.21, 0.94) | | 14% (-71, 343) | >0.80 |
| NEOx | Media | AA | Alcohol | 9-HETE | 60 | 0.27 (0.12, 0.61) | 0.32 (0.14, 0.71) | | 17% (-80, 580) | >0.80 |
| Esterified | Cell | AA | Alcohol | 12-HETE | 0 | 0.85 (0.39, 1.89) | 0.51 (0.23, 1.14) | 0.77 | -40% (-89, 218) | >0.80 |
| Esterified | Cell | AA | Alcohol | 12-HETE | 15 | 1.05 (0.53, 2.09) | 0.67 (0.34, 1.33) | | -36% (-83, 142) | >0.80 |
| Esterified | Cell | AA | Alcohol | 12-HETE | 30 | 1.2 (0.59, 2.42) | 0.8 (0.4, 1.63) | | -33% (-81, 137) | >0.80 |
| Esterified | Cell | AA | Alcohol | 12-HETE | 60 | 1.22 (0.53, 2.85) | 0.92 (0.39, 2.13) | | -25% (-89, 394) | >0.80 |
| NEOx | Cell | AA | Alcohol | 12-HETE | 0 | 0.41 (0.26, 0.65) | 0.36 (0.23, 0.57) | 0.39 | -13% (-65, 120) | >0.80 |
| NEOx | Cell | AA | Alcohol | 12-HETE | 15 | 0.47 (0.3, 0.74) | 0.43 (0.27, 0.68) | | -7% (-60, 116) | >0.80 |
| NEOx | Cell | AA | Alcohol | 12-HETE | 30 | 0.49 (0.31, 0.77) | 0.48 (0.31, 0.76) | | -2% (-57, 125) | >0.80 |
| NEOx | Cell | AA | Alcohol | 12-HETE | 60 | 0.43 (0.27, 0.68) | 0.47 (0.3, 0.75) | | 11% (-58, 195) | >0.80 |
| Esterified | Media | AA | Alcohol | 12-HETE | 0 | 0.41 (0.27, 0.61) | 0.36 (0.24, 0.53) | 0.66 | -13% (-61, 93) | >0.80 |
| Esterified | Media | AA | Alcohol | 12-HETE | 15 | 0.45 (0.3, 0.66) | 0.38 (0.25, 0.56) | | -16% (-60, 77) | >0.80 |
| Esterified | Media | AA | Alcohol | 12-HETE | 30 | 0.48 (0.32, 0.72) | 0.4 (0.27, 0.59) | | -18% (-60, 70) | >0.80 |
| Esterified | Media | AA | Alcohol | 12-HETE | 60 | 0.54 (0.36, 0.81) | 0.42 (0.28, 0.64) | | -22% (-67, 83) | >0.80 |
| NEOx | Media | AA | Alcohol | 12-HETE | 0 | 0.51 (0.32, 0.8) | 0.36 (0.23, 0.57) | 0.63 | -29% (-71, 78) | >0.80 |
| NEOx | Media | AA | Alcohol | 12-HETE | 15 | 0.49 (0.31, 0.78) | 0.36 (0.23, 0.58) | | -26% (-69, 72) | >0.80 |
| NEOx | Media | AA | Alcohol | 12-HETE | 30 | 0.48 (0.3, 0.76) | 0.36 (0.23, 0.58) | | -24% (-67, 75) | >0.80 |
| NEOx | Media | AA | Alcohol | 12-HETE | 60 | 0.47 (0.29, 0.74) | 0.38 (0.24, 0.6) | | -20% (-69, 110) | >0.80 |
| Esterified | Cell | AA | Alcohol | 15-HETE | 0 | 1.13 (0.31, 4.1) | 1.19 (0.33, 4.31) | >0.80 | 5% (-93, 1460) | >0.80 |
| Esterified | Cell | AA | Alcohol | 15-HETE | 15 | 1.21 (0.39, 3.76) | 1.29 (0.41, 4.01) | | 7% (-88, 857) | >0.80 |
| Esterified | Cell | AA | Alcohol | 15-HETE | 30 | 1.3 (0.41, 4.12) | 1.41 (0.44, 4.48) | | 9% (-86, 768) | >0.80 |
| Esterified | Cell | AA | Alcohol | 15-HETE | 60 | 1.51 (0.38, 5.89) | 1.69 (0.43, 6.62) | | 12% (-95, 2260) | >0.80 |
| NEOx | Cell | AA | Alcohol | 15-HETE | 0 | 0.33 (0.15, 0.73) | 0.3 (0.14, 0.65) | 0.61 | -11% (-82, 337) | >0.80 |
| NEOx | Cell | AA | Alcohol | 15-HETE | 15 | 0.29 (0.14, 0.61) | 0.28 (0.13, 0.58) | | -4% (-77, 289) | >0.80 |
| NEOx | Cell | AA | Alcohol | 15-HETE | 30 | 0.27 (0.13, 0.57) | 0.28 (0.13, 0.59) | | 3% (-74, 300) | >0.80 |
| NEOx | Cell | AA | Alcohol | 15-HETE | 60 | 0.31 (0.14, 0.69) | 0.37 (0.17, 0.81) | | 18% (-79, 565) | >0.80 |
| Esterified | Media | AA | Alcohol | 15-HETE | 0 | 0.22 (0.13, 0.38) | 0.18 (0.1, 0.31) | 0.42 | -19% (-74, 154) | >0.80 |
| Esterified | Media | AA | Alcohol | 15-HETE | 15 | 0.26 (0.16, 0.42) | 0.19 (0.12, 0.31) | | -27% (-71, 85) | >0.80 |
| Esterified | Media | AA | Alcohol | 15-HETE | 30 | 0.31 (0.19, 0.51) | 0.21 (0.13, 0.34) | | -34% (-73, 59) | 0.72 |
| Esterified | Media | AA | Alcohol | 15-HETE | 60 | 0.5 (0.28, 0.9) | 0.27 (0.15, 0.49) | | -46% (-85, 97) | 0.71 |
| NEOx | Media | AA | Alcohol | 15-HETE | 0 | 0.19 (0.1, 0.39) | 0.23 (0.12, 0.47) | 0.53 | 21% (-71, 404) | >0.80 |
| NEOx | Media | AA | Alcohol | 15-HETE | 15 | 0.21 (0.11, 0.39) | 0.23 (0.12, 0.44) | | 11% (-68, 278) | >0.80 |
| NEOx | Media | AA | Alcohol | 15-HETE | 30 | 0.23 (0.12, 0.43) | 0.23 (0.12, 0.44) | | 2% (-69, 232) | >0.80 |
| NEOx | Media | AA | Alcohol | 15-HETE | 60 | 0.3 (0.15, 0.62) | 0.26 (0.13, 0.54) | | -14% (-82, 314) | >0.80 |
| Esterified | Cell | EPA | Alcohol | 5-HEPE | 0 | 0.89 (0.24, 3.26) | 0.86 (0.23, 3.18) | 0.21 | -2% (-93, 1305) | >0.80 |
| Esterified | Cell | EPA | Alcohol | 5-HEPE | 15 | 1.12 (0.32, 3.99) | 0.82 (0.23, 2.91) | | -27% (-93, 680) | >0.80 |
| Esterified | Cell | EPA | Alcohol | 5-HEPE | 30 | 1.61 (0.45, 5.69) | 0.88 (0.25, 3.1) | | -46% (-95, 447) | >0.80 |
| Esterified | Cell | EPA | Alcohol | 5-HEPE | 60 | 4.71 (1.24, 17.92) | 1.43 (0.38, 5.45) | | -70% (-98, 444) | >0.80 |
| NEOx | Cell | EPA | Alcohol | 5-HEPE | 0 | 0.34 (0.21, 0.54) | 0.26 (0.16, 0.42) | >0.80 | -23% (-70, 99) | >0.80 |
| NEOx | Cell | EPA | Alcohol | 5-HEPE | 15 | 0.31 (0.2, 0.5) | 0.24 (0.15, 0.39) | | -22% (-67, 83) | >0.80 |
| NEOx | Cell | EPA | Alcohol | 5-HEPE | 30 | 0.32 (0.2, 0.51) | 0.25 (0.16, 0.4) | | -22% (-66, 81) | >0.80 |
| NEOx | Cell | EPA | Alcohol | 5-HEPE | 60 | 0.45 (0.28, 0.73) | 0.36 (0.22, 0.57) | | -21% (-71, 119) | >0.80 |
| Esterified | Media | EPA | Alcohol | 5-HEPE | 0 | 0.36 (0.23, 0.58) | 0.28 (0.17, 0.44) | 0.14 | -24% (-71, 99) | >0.80 |
| Esterified | Media | EPA | Alcohol | 5-HEPE | 15 | 0.32 (0.2, 0.5) | 0.28 (0.18, 0.43) | | -13% (-62, 102) | >0.80 |
| Esterified | Media | EPA | Alcohol | 5-HEPE | 30 | 0.29 (0.19, 0.46) | 0.29 (0.19, 0.45) | | -1% (-56, 124) | >0.80 |
| Esterified | Media | EPA | Alcohol | 5-HEPE | 60 | 0.28 (0.17, 0.45) | 0.36 (0.22, 0.58) | | 29% (-55, 269) | >0.80 |
| NEOx | Media | EPA | Alcohol | 5-HEPE | 0 | 0.39 (0.22, 0.69) | 0.2 (0.11, 0.34) | 0.001 | -50% (-84, 51) | 0.41 |
| NEOx | Media | EPA | Alcohol | 5-HEPE | 15 | 0.31 (0.18, 0.55) | 0.21 (0.12, 0.37) | | -34% (-77, 88) | >0.80 |
| NEOx | Media | EPA | Alcohol | 5-HEPE | 30 | 0.26 (0.15, 0.47) | 0.24 (0.13, 0.41) | | -11% (-68, 147) | >0.80 |
| NEOx | Media | EPA | Alcohol | 5-HEPE | 60 | 0.23 (0.13, 0.41) | 0.37 (0.21, 0.66) | | 59% (-51, 411) | >0.80 |
| Esterified | Cell | EPA | Alcohol | 8-HEPE | 0 | 1.15 (0.71, 1.86) | 0.54 (0.33, 0.87) | 0.08 | -53% (-83, 28) | 0.22 |
| Esterified | Cell | EPA | Alcohol | 8-HEPE | 15 | 0.94 (0.63, 1.4) | 0.55 (0.37, 0.82) | | -42% (-73, 28) | 0.31 |
| Esterified | Cell | EPA | Alcohol | 8-HEPE | 30 | 0.83 (0.55, 1.26) | 0.6 (0.4, 0.91) | | -28% (-65, 51) | 0.78 |
| Esterified | Cell | EPA | Alcohol | 8-HEPE | 60 | 0.83 (0.49, 1.38) | 0.93 (0.55, 1.55) | | 12% (-65, 256) | >0.80 |
| NEOx | Cell | EPA | Alcohol | 8-HEPE | 0 | 0.55 (0.29, 1.03) | 0.38 (0.2, 0.72) | 0.77 | -30% (-82, 164) | >0.80 |
| NEOx | Cell | EPA | Alcohol | 8-HEPE | 15 | 0.57 (0.32, 1.01) | 0.41 (0.23, 0.73) | | -28% (-76, 117) | >0.80 |
| NEOx | Cell | EPA | Alcohol | 8-HEPE | 30 | 0.61 (0.34, 1.08) | 0.46 (0.25, 0.81) | | -25% (-74, 115) | >0.80 |
| NEOx | Cell | EPA | Alcohol | 8-HEPE | 60 | 0.72 (0.37, 1.4) | 0.59 (0.3, 1.14) | | -18% (-82, 263) | >0.80 |
| Esterified | Media | EPA | Alcohol | 8-HEPE | 0 | 0.44 (0.22, 0.89) | 0.52 (0.26, 1.05) | 0.66 | 17% (-72, 392) | >0.80 |
| Esterified | Media | EPA | Alcohol | 8-HEPE | 15 | 0.5 (0.25, 1) | 0.56 (0.28, 1.12) | | 11% (-69, 305) | >0.80 |
| Esterified | Media | EPA | Alcohol | 8-HEPE | 30 | 0.54 (0.27, 1.07) | 0.57 (0.29, 1.13) | | 6% (-70, 273) | >0.80 |
| Esterified | Media | EPA | Alcohol | 8-HEPE | 60 | 0.51 (0.25, 1.04) | 0.48 (0.24, 0.99) | | -5% (-80, 345) | >0.80 |
| NEOx | Media | EPA | Alcohol | 8-HEPE | 0 | 0.6 (0.35, 1.03) | 0.59 (0.34, 1) | >0.80 | -2% (-68, 196) | >0.80 |
| NEOx | Media | EPA | Alcohol | 8-HEPE | 15 | 0.59 (0.36, 0.97) | 0.58 (0.35, 0.97) | | -1% (-62, 158) | >0.80 |
| NEOx | Media | EPA | Alcohol | 8-HEPE | 30 | 0.55 (0.33, 0.9) | 0.55 (0.33, 0.91) | | 1% (-60, 153) | >0.80 |
| NEOx | Media | EPA | Alcohol | 8-HEPE | 60 | 0.4 (0.23, 0.69) | 0.41 (0.24, 0.72) | | 4% (-69, 251) | >0.80 |
| Esterified | Cell | EPA | Alcohol | 11-HEPE | 0 | 0.39 (0.25, 0.6) | 0.46 (0.3, 0.72) | 0.08 | 20% (-53, 206) | >0.80 |
| Esterified | Cell | EPA | Alcohol | 11-HEPE | 15 | 0.36 (0.24, 0.53) | 0.36 (0.24, 0.52) | | -1% (-53, 112) | >0.80 |
| Esterified | Cell | EPA | Alcohol | 11-HEPE | 30 | 0.37 (0.25, 0.55) | 0.31 (0.21, 0.46) | | -17% (-60, 68) | >0.80 |
| Esterified | Cell | EPA | Alcohol | 11-HEPE | 60 | 0.58 (0.36, 0.94) | 0.33 (0.21, 0.53) | | -43% (-80, 64) | 0.59 |
| NEOx | Cell | EPA | Alcohol | 11-HEPE | 0 | 0.28 (0.15, 0.52) | 0.26 (0.14, 0.49) | 0.32 | -6% (-74, 238) | >0.80 |

| | | | | | | | | | | |
|---|---|---|---|---|---|---|---|---|---|---|
| NEOx | Cell | EPA | Alcohol | 11-HEPE | 15 | 0.32 (0.19, 0.54) | 0.26 (0.15, 0.44) | | -18% (-71, 132) | >0.80 |
| NEOx | Cell | EPA | Alcohol | 11-HEPE | 30 | 0.36 (0.21, 0.62) | 0.26 (0.15, 0.44) | | -29% (-73, 91) | >0.80 |
| NEOx | Cell | EPA | Alcohol | 11-HEPE | 60 | 0.46 (0.24, 0.89) | 0.25 (0.13, 0.48) | | -46% (-87, 130) | >0.80 |
| Esterified | Media | EPA | Alcohol | 11-HEPE | 0 | 0.29 (0.16, 0.51) | 0.23 (0.13, 0.4) | 0.7 | -22% (-76, 153) | >0.80 |
| Esterified | Media | EPA | Alcohol | 11-HEPE | 15 | 0.36 (0.22, 0.6) | 0.27 (0.16, 0.45) | | -25% (-72, 97) | >0.80 |
| Esterified | Media | EPA | Alcohol | 11-HEPE | 30 | 0.39 (0.23, 0.65) | 0.28 (0.17, 0.46) | | -29% (-72, 80) | >0.80 |
| Esterified | Media | EPA | Alcohol | 11-HEPE | 60 | 0.28 (0.15, 0.5) | 0.18 (0.1, 0.33) | | -35% (-83, 141) | >0.80 |
| NEOx | Media | EPA | Alcohol | 11-HEPE | 0 | 0.34 (0.23, 0.51) | 0.22 (0.15, 0.33) | 0.7 | -36% (-73, 50) | 0.61 |
| NEOx | Media | EPA | Alcohol | 11-HEPE | 15 | 0.32 (0.22, 0.48) | 0.21 (0.14, 0.32) | | -34% (-68, 38) | 0.54 |
| NEOx | Media | EPA | Alcohol | 11-HEPE | 30 | 0.31 (0.21, 0.46) | 0.21 (0.14, 0.31) | | -32% (-67, 39) | 0.58 |
| NEOx | Media | EPA | Alcohol | 11-HEPE | 60 | 0.3 (0.19, 0.46) | 0.22 (0.14, 0.33) | | -28% (-71, 82) | >0.80 |
| Esterified | Cell | EPA | Alcohol | 12-HEPE | 0 | 0.36 (0.16, 0.78) | 0.24 (0.11, 0.53) | >0.80 | -32% (-87, 250) | >0.80 |
| Esterified | Cell | EPA | Alcohol | 12-HEPE | 15 | 0.33 (0.17, 0.68) | 0.23 (0.12, 0.47) | | -30% (-82, 171) | >0.80 |
| Esterified | Cell | EPA | Alcohol | 12-HEPE | 30 | 0.34 (0.17, 0.7) | 0.25 (0.12, 0.51) | | -28% (-80, 162) | >0.80 |
| Esterified | Cell | EPA | Alcohol | 12-HEPE | 60 | 0.48 (0.21, 1.09) | 0.37 (0.16, 0.83) | | -23% (-88, 380) | >0.80 |
| NEOx | Cell | EPA | Alcohol | 12-HEPE | 0 | 0.37 (0.22, 0.62) | 0.19 (0.11, 0.33) | 0.34 | -48% (-82, 57) | 0.48 |
| NEOx | Cell | EPA | Alcohol | 12-HEPE | 15 | 0.33 (0.22, 0.5) | 0.2 (0.13, 0.3) | | -41% (-74, 37) | 0.42 |
| NEOx | Cell | EPA | Alcohol | 12-HEPE | 30 | 0.3 (0.19, 0.47) | 0.2 (0.13, 0.32) | | -33% (-69, 46) | 0.64 |
| NEOx | Cell | EPA | Alcohol | 12-HEPE | 60 | 0.27 (0.15, 0.47) | 0.23 (0.13, 0.41) | | -14% (-76, 206) | >0.80 |
| Esterified | Media | EPA | Alcohol | 12-HEPE | 0 | 0.33 (0.23, 0.48) | 0.24 (0.16, 0.35) | 0.04 | -28% (-67, 59) | >0.80 |
| Esterified | Media | EPA | Alcohol | 12-HEPE | 15 | 0.26 (0.19, 0.37) | 0.23 (0.17, 0.32) | | -12% (-53, 65) | >0.80 |
| Esterified | Media | EPA | Alcohol | 12-HEPE | 30 | 0.23 (0.16, 0.32) | 0.24 (0.17, 0.34) | | 7% (-41, 93) | >0.80 |
| Esterified | Media | EPA | Alcohol | 12-HEPE | 60 | 0.22 (0.14, 0.32) | 0.34 (0.23, 0.51) | | 58% (-36, 290) | 0.65 |
| NEOx | Media | EPA | Alcohol | 12-HEPE | 0 | 0.3 (0.14, 0.67) | 0.17 (0.08, 0.38) | 0.17 | -44% (-88, 162) | >0.80 |
| NEOx | Media | EPA | Alcohol | 12-HEPE | 15 | 0.26 (0.12, 0.59) | 0.17 (0.08, 0.38) | | -36% (-85, 175) | >0.80 |
| NEOx | Media | EPA | Alcohol | 12-HEPE | 30 | 0.24 (0.11, 0.53) | 0.18 (0.08, 0.39) | | -27% (-83, 208) | >0.80 |
| NEOx | Media | EPA | Alcohol | 12-HEPE | 60 | 0.23 (0.1, 0.51) | 0.22 (0.1, 0.48) | | -6% (-81, 366) | >0.80 |
| Esterified | Cell | EPA | Alcohol | 18-HEPE | 0 | 0.19 (0.11, 0.32) | 0.15 (0.09, 0.25) | 0.004 | -22% (-72, 122) | >0.80 |
| Esterified | Cell | EPA | Alcohol | 18-HEPE | 15 | 0.37 (0.29, 0.49) | 0.17 (0.13, 0.22) | | **-55% (-73, -26)** | **0.001** |
| Esterified | Cell | EPA | Alcohol | 18-HEPE | 30 | 0.6 (0.42, 0.86) | 0.16 (0.11, 0.22) | | **-74% (-81, -66)** | **<0.0001** |
| Esterified | Cell | EPA | Alcohol | 18-HEPE | 60 | 0.82 (0.45, 1.5) | 0.07 (0.04, 0.13) | | **-91% (-98, -68)** | **0.0002** |
| NEOx | Cell | EPA | Alcohol | 18-HEPE | 0 | 0.12 (0.08, 0.19) | 0.08 (0.06, 0.13) | 0.0005 | -33% (-72, 62) | 0.76 |
| NEOx | Cell | EPA | Alcohol | 18-HEPE | 15 | 0.26 (0.18, 0.37) | 0.11 (0.08, 0.16) | | **-56% (-78, -13)** | **0.01** |
| NEOx | Cell | EPA | Alcohol | 18-HEPE | 30 | 0.43 (0.3, 0.62) | 0.12 (0.08, 0.18) | | **-72% (-85, -46)** | **0.0001** |
| NEOx | Cell | EPA | Alcohol | 18-HEPE | 60 | 0.61 (0.39, 0.95) | 0.07 (0.05, 0.11) | | **-88% (-96, -67)** | **<0.0001** |
| Esterified | Media | EPA | Alcohol | 18-HEPE | 0 | 0.32 (0.18, 0.55) | 0.17 (0.1, 0.3) | 0.09 | -46% (-83, 68) | 0.57 |
| Esterified | Media | EPA | Alcohol | 18-HEPE | 15 | 0.34 (0.2, 0.56) | 0.15 (0.09, 0.25) | | -56% (-83, 16) | 0.13 |
| Esterified | Media | EPA | Alcohol | 18-HEPE | 30 | 0.37 (0.22, 0.61) | 0.13 (0.08, 0.22) | | **-64% (-86, -8)** | **0.03** |
| Esterified | Media | EPA | Alcohol | 18-HEPE | 60 | 0.47 (0.27, 0.84) | 0.12 (0.07, 0.21) | | **-75% (-93, -13)** | **0.02** |
| NEOx | Media | EPA | Alcohol | 18-HEPE | 0 | 0.14 (0.07, 0.27) | 0.14 (0.07, 0.29) | 0.14 | 6% (-75, 344) | >0.80 |
| NEOx | Media | EPA | Alcohol | 18-HEPE | 15 | 0.15 (0.08, 0.29) | 0.13 (0.07, 0.25) | | -14% (-75, 195) | >0.80 |
| NEOx | Media | EPA | Alcohol | 18-HEPE | 30 | 0.17 (0.09, 0.32) | 0.12 (0.06, 0.23) | | -30% (-79, 130) | >0.80 |
| NEOx | Media | EPA | Alcohol | 18-HEPE | 60 | 0.19 (0.09, 0.4) | 0.09 (0.04, 0.19) | | -53% (-90, 125) | 0.69 |
| Esterified | Cell | DHA | Alcohol | 4-HDoHE | 0 | 23.7 (10.5, 53.3) | 24.4 (10.9, 54.9) | 0.6 | 3% (-81, 447) | >0.80 |
| Esterified | Cell | DHA | Alcohol | 4-HDoHE | 15 | 30.9 (14.2, 67.4) | 29.5 (13.5, 64.3) | | -5% (-78, 313) | >0.80 |
| Esterified | Cell | DHA | Alcohol | 4-HDoHE | 30 | 38.1 (17.5, 83.1) | 33.7 (15.5, 73.4) | | -12% (-79, 266) | >0.80 |
| Esterified | Cell | DHA | Alcohol | 4-HDoHE | 60 | 48.6 (21.1, 112.2) | 36.8 (16, 84.8) | | -24% (-88, 365) | >0.80 |
| NEOx | Cell | DHA | Alcohol | 4-HDoHE | 0 | 0.57 (0.36, 0.89) | 0.39 (0.25, 0.62) | >0.80 | -30% (-72, 73) | >0.80 |
| NEOx | Cell | DHA | Alcohol | 4-HDoHE | 15 | 0.64 (0.4, 1) | 0.44 (0.28, 0.7) | | -30% (-70, 61) | 0.80 |
| NEOx | Cell | DHA | Alcohol | 4-HDoHE | 30 | 0.67 (0.43, 1.05) | 0.47 (0.3, 0.74) | | -30% (-69, 59) | 0.79 |
| NEOx | Cell | DHA | Alcohol | 4-HDoHE | 60 | 0.62 (0.39, 0.97) | 0.43 (0.27, 0.68) | | -30% (-73, 83) | >0.80 |
| Esterified | Media | DHA | Alcohol | 4-HDoHE | 0 | 0.5 (0.25, 0.99) | 0.59 (0.3, 1.18) | 0.39 | 19% (-72, 406) | >0.80 |
| Esterified | Media | DHA | Alcohol | 4-HDoHE | 15 | 0.76 (0.42, 1.38) | 0.79 (0.43, 1.44) | | 4% (-67, 232) | >0.80 |
| Esterified | Media | DHA | Alcohol | 4-HDoHE | 30 | 1.01 (0.55, 1.86) | 0.91 (0.49, 1.68) | | -9% (-70, 171) | >0.80 |
| Esterified | Media | DHA | Alcohol | 4-HDoHE | 60 | 1.15 (0.55, 2.39) | 0.79 (0.38, 1.64) | | -31% (-87, 253) | >0.80 |
| NEOx | Media | DHA | Alcohol | 4-HDoHE | 0 | 0.55 (0.36, 0.83) | 0.54 (0.36, 0.82) | 0.65 | -1% (-56, 123) | >0.80 |
| NEOx | Media | DHA | Alcohol | 4-HDoHE | 15 | 0.53 (0.35, 0.8) | 0.53 (0.35, 0.81) | | 1% (-52, 116) | >0.80 |
| NEOx | Media | DHA | Alcohol | 4-HDoHE | 30 | 0.5 (0.33, 0.76) | 0.52 (0.35, 0.79) | | 4% (-51, 119) | >0.80 |
| NEOx | Media | DHA | Alcohol | 4-HDoHE | 60 | 0.46 (0.3, 0.69) | 0.5 (0.33, 0.76) | | 9% (-54, 157) | >0.80 |
| Esterified | Cell | DHA | Alcohol | 7-HDoHE | 0 | 1.1 (0.37, 3.22) | 1.56 (0.53, 4.58) | 0.51 | 42% (-85, 1263) | >0.80 |
| Esterified | Cell | DHA | Alcohol | 7-HDoHE | 15 | 1.49 (0.57, 3.88) | 1.81 (0.69, 4.71) | | 21% (-81, 669) | >0.80 |
| Esterified | Cell | DHA | Alcohol | 7-HDoHE | 30 | 1.93 (0.73, 5.11) | 2 (0.75, 5.29) | | 3% (-82, 496) | >0.80 |
| Esterified | Cell | DHA | Alcohol | 7-HDoHE | 60 | 2.77 (0.89, 8.68) | 2.09 (0.67, 6.53) | | -25% (-94, 858) | >0.80 |
| NEOx | Cell | DHA | Alcohol | 7-HDoHE | 0 | 0.55 (0.37, 0.82) | 0.55 (0.37, 0.83) | 0.12 | 1% (-57, 135) | >0.80 |
| NEOx | Cell | DHA | Alcohol | 7-HDoHE | 15 | 0.64 (0.46, 0.9) | 0.56 (0.4, 0.78) | | -14% (-55, 68) | >0.80 |
| NEOx | Cell | DHA | Alcohol | 7-HDoHE | 30 | 0.73 (0.51, 1.04) | 0.54 (0.38, 0.76) | | -26% (-60, 37) | 0.68 |
| NEOx | Cell | DHA | Alcohol | 7-HDoHE | 60 | 0.83 (0.54, 1.27) | 0.45 (0.29, 0.69) | | -46% (-79, 42) | 0.39 |
| Esterified | Media | DHA | Alcohol | 7-HDoHE | 0 | 0.7 (0.37, 1.33) | 0.45 (0.24, 0.85) | 0.34 | -36% (-82, 125) | >0.80 |
| Esterified | Media | DHA | Alcohol | 7-HDoHE | 15 | 0.69 (0.37, 1.32) | 0.48 (0.25, 0.91) | | -31% (-79, 124) | >0.80 |
| Esterified | Media | DHA | Alcohol | 7-HDoHE | 30 | 0.67 (0.36, 1.28) | 0.5 (0.27, 0.95) | | -25% (-77, 138) | >0.80 |
| Esterified | Media | DHA | Alcohol | 7-HDoHE | 60 | 0.6 (0.32, 1.14) | 0.53 (0.28, 1) | | -13% (-77, 227) | >0.80 |
| NEOx | Media | DHA | Alcohol | 7-HDoHE | 0 | 0.57 (0.28, 1.16) | 0.5 (0.25, 1.01) | 0.68 | -13% (-78, 246) | >0.80 |
| NEOx | Media | DHA | Alcohol | 7-HDoHE | 15 | 0.57 (0.28, 1.15) | 0.51 (0.25, 1.04) | | -10% (-75, 232) | >0.80 |
| NEOx | Media | DHA | Alcohol | 7-HDoHE | 30 | 0.56 (0.28, 1.14) | 0.53 (0.26, 1.07) | | -7% (-74, 238) | >0.80 |
| NEOx | Media | DHA | Alcohol | 7-HDoHE | 60 | 0.57 (0.28, 1.16) | 0.57 (0.28, 1.16) | | 0% (-76, 319) | >0.80 |
| Esterified | Cell | DHA | Alcohol | 8-HDoHE | 0 | 3.92 (1.86, 8.27) | 2.98 (1.41, 6.3) | >0.80 | -24% (-84, 262) | >0.80 |
| Esterified | Cell | DHA | Alcohol | 8-HDoHE | 15 | 7.73 (4.39, 13.63) | 6.02 (3.42, 10.61) | | -22% (-75, 140) | >0.80 |
| Esterified | Cell | DHA | Alcohol | 8-HDoHE | 30 | 11.82 (6.44, 21.7) | 9.41 (5.13, 17.27) | | -20% (-71, 121) | >0.80 |
| Esterified | Cell | DHA | Alcohol | 8-HDoHE | 60 | 12.81 (5.67, 28.94) | 10.65 (4.71, 24.07) | | -17% (-87, 420) | >0.80 |
| NEOx | Cell | DHA | Alcohol | 8-HDoHE | 0 | 0.61 (0.36, 1.02) | 0.49 (0.29, 0.83) | 0.17 | -19% (-73, 138) | >0.80 |
| NEOx | Cell | DHA | Alcohol | 8-HDoHE | 15 | 0.97 (0.59, 1.59) | 0.68 (0.42, 1.12) | | -30% (-72, 78) | >0.80 |
| NEOx | Cell | DHA | Alcohol | 8-HDoHE | 30 | 1.26 (0.77, 2.07) | 0.77 (0.47, 1.26) | | -39% (-75, 50) | 0.56 |
| NEOx | Cell | DHA | Alcohol | 8-HDoHE | 60 | 1.13 (0.66, 1.94) | 0.52 (0.3, 0.89) | | -54% (-86, 50) | 0.36 |
| NEOx | Media | DHA | Alcohol | 8-HDoHE | 0 | 0.97 (0.55, 1.7) | 0.76 (0.43, 1.33) | 0.62 | -22% (-75, 144) | >0.80 |
| NEOx | Media | DHA | Alcohol | 8-HDoHE | 15 | 0.77 (0.44, 1.34) | 0.63 (0.36, 1.09) | | -18% (-71, 130) | >0.80 |

| | | | | | | | | | | |
|---|---|---|---|---|---|---|---|---|---|---|
| NEOx | Media | DHA | Alcohol | 8-HDoHE | 30 | 0.68 (0.39, 1.18) | 0.58 (0.33, 1.01) | | -15% (-69, 135) | >0.80 |
| NEOx | Media | DHA | Alcohol | 8-HDoHE | 60 | 0.75 (0.42, 1.33) | 0.7 (0.39, 1.23) | | -7% (-72, 211) | >0.80 |
| Esterified | Cell | DHA | Alcohol | 10-HDoHE | 0 | 1.29 (0.45, 3.7) | 0.99 (0.34, 2.84) | >0.80 | -23% (-91, 586) | >0.80 |
| Esterified | Cell | DHA | Alcohol | 10-HDoHE | 15 | 1.84 (0.68, 4.96) | 1.47 (0.55, 3.96) | | -20% (-88, 423) | >0.80 |
| Esterified | Cell | DHA | Alcohol | 10-HDoHE | 30 | 2.43 (0.9, 6.56) | 2.01 (0.74, 5.45) | | -17% (-86, 408) | >0.80 |
| Esterified | Cell | DHA | Alcohol | 10-HDoHE | 60 | 3.28 (1.1, 9.82) | 2.95 (0.98, 8.82) | | -10% (-92, 900) | >0.80 |
| NEOx | Cell | DHA | Alcohol | 10-HDoHE | 0 | 1.16 (0.61, 2.21) | 0.72 (0.38, 1.37) | 0.44 | -38% (-84, 139) | >0.80 |
| NEOx | Cell | DHA | Alcohol | 10-HDoHE | 15 | 0.81 (0.48, 1.39) | 0.57 (0.34, 0.98) | | -30% (-75, 100) | >0.80 |
| NEOx | Cell | DHA | Alcohol | 10-HDoHE | 30 | 0.7 (0.4, 1.22) | 0.56 (0.32, 0.97) | | -20% (-70, 111) | >0.80 |
| NEOx | Cell | DHA | Alcohol | 10-HDoHE | 60 | 0.98 (0.49, 1.96) | 1 (0.5, 1.99) | | 1% (-78, 375) | >0.80 |
| Esterified | Media | DHA | Alcohol | 10-HDoHE | 0 | 0.67 (0.36, 1.26) | 0.71 (0.38, 1.34) | 0.49 | 7% (-71, 294) | >0.80 |
| Esterified | Media | DHA | Alcohol | 10-HDoHE | 15 | 0.62 (0.34, 1.11) | 0.6 (0.33, 1.08) | | -2% (-68, 197) | >0.80 |
| Esterified | Media | DHA | Alcohol | 10-HDoHE | 30 | 0.64 (0.36, 1.16) | 0.57 (0.32, 1.04) | | -11% (-69, 161) | >0.80 |
| Esterified | Media | DHA | Alcohol | 10-HDoHE | 60 | 1.02 (0.53, 1.96) | 0.76 (0.4, 1.47) | | -25% (-82, 219) | >0.80 |
| NEOx | Media | DHA | Alcohol | 10-HDoHE | 0 | 0.65 (0.38, 1.13) | 0.72 (0.42, 1.25) | 0.4 | 11% (-64, 245) | >0.80 |
| NEOx | Media | DHA | Alcohol | 10-HDoHE | 15 | 0.64 (0.38, 1.05) | 0.64 (0.39, 1.06) | | 1% (-61, 163) | >0.80 |
| NEOx | Media | DHA | Alcohol | 10-HDoHE | 30 | 0.6 (0.36, 0.99) | 0.55 (0.33, 0.91) | | -8% (-64, 130) | >0.80 |
| NEOx | Media | DHA | Alcohol | 10-HDoHE | 60 | 0.48 (0.27, 0.84) | 0.36 (0.2, 0.64) | | -24% (-78, 166) | >0.80 |
| Esterified | Cell | DHA | Alcohol | 13-HDoHE | 0 | 0.73 (0.39, 1.35) | 0.69 (0.37, 1.28) | >0.80 | -5% (-74, 249) | >0.80 |
| Esterified | Cell | DHA | Alcohol | 13-HDoHE | 15 | 1.2 (0.71, 2.03) | 1.13 (0.67, 1.91) | | -6% (-66, 161) | >0.80 |
| Esterified | Cell | DHA | Alcohol | 13-HDoHE | 30 | 1.74 (1.02, 2.99) | 1.64 (0.95, 2.81) | | -6% (-64, 143) | >0.80 |
| Esterified | Cell | DHA | Alcohol | 13-HDoHE | 60 | 2.48 (1.28, 4.81) | 2.29 (1.18, 4.45) | | -7% (-79, 310) | >0.80 |
| NEOx | Cell | DHA | Alcohol | 13-HDoHE | 0 | 0.61 (0.36, 1.03) | 0.41 (0.25, 0.69) | 0.52 | -33% (-77, 97) | >0.80 |
| NEOx | Cell | DHA | Alcohol | 13-HDoHE | 15 | 0.61 (0.38, 1) | 0.44 (0.27, 0.72) | | -28% (-72, 82) | >0.80 |
| NEOx | Cell | DHA | Alcohol | 13-HDoHE | 30 | 0.64 (0.39, 1.04) | 0.49 (0.3, 0.8) | | -23% (-69, 88) | >0.80 |
| NEOx | Cell | DHA | Alcohol | 13-HDoHE | 60 | 0.76 (0.44, 1.3) | 0.66 (0.39, 1.13) | | -13% (-73, 183) | >0.80 |
| Esterified | Media | DHA | Alcohol | 13-HDoHE | 0 | 0.65 (0.4, 1.05) | 0.56 (0.34, 0.92) | 0.55 | -13% (-68, 134) | >0.80 |
| Esterified | Media | DHA | Alcohol | 13-HDoHE | 15 | 0.69 (0.42, 1.12) | 0.57 (0.35, 0.93) | | -17% (-66, 104) | >0.80 |
| Esterified | Media | DHA | Alcohol | 13-HDoHE | 30 | 0.73 (0.45, 1.18) | 0.58 (0.36, 0.93) | | -21% (-67, 92) | >0.80 |
| Esterified | Media | DHA | Alcohol | 13-HDoHE | 60 | 0.76 (0.46, 1.25) | 0.55 (0.33, 0.91) | | -28% (-75, 108) | >0.80 |
| NEOx | Media | DHA | Alcohol | 13-HDoHE | 0 | 0.68 (0.43, 1.06) | 0.57 (0.36, 0.88) | >0.80 | -17% (-66, 107) | >0.80 |
| NEOx | Media | DHA | Alcohol | 13-HDoHE | 15 | 0.54 (0.35, 0.83) | 0.45 (0.29, 0.69) | | -16% (-63, 88) | >0.80 |
| NEOx | Media | DHA | Alcohol | 13-HDoHE | 30 | 0.48 (0.31, 0.75) | 0.4 (0.26, 0.62) | | -16% (-62, 85) | >0.80 |
| NEOx | Media | DHA | Alcohol | 13-HDoHE | 60 | 0.56 (0.36, 0.89) | 0.47 (0.3, 0.75) | | -16% (-68, 125) | >0.80 |
| Esterified | Cell | DHA | Alcohol | 14-HDoHE | 0 | 2.2 (0.83, 5.79) | 1.29 (0.49, 3.41) | >0.80 | -41% (-92, 347) | >0.80 |
| Esterified | Cell | DHA | Alcohol | 14-HDoHE | 15 | 2.68 (1.24, 5.8) | 1.61 (0.74, 3.49) | | -40% (-87, 177) | >0.80 |
| Esterified | Cell | DHA | Alcohol | 14-HDoHE | 30 | 3.52 (1.56, 7.94) | 2.16 (0.96, 4.87) | | -39% (-85, 150) | >0.80 |
| Esterified | Cell | DHA | Alcohol | 14-HDoHE | 60 | 7.6 (2.67, 21.65) | 4.86 (1.71, 13.86) | | -36% (-94, 574) | >0.80 |
| NEOx | Cell | DHA | Alcohol | 14-HDoHE | 0 | 0.42 (0.22, 0.81) | 0.37 (0.19, 0.71) | 0.55 | -12% (-78, 248) | >0.80 |
| NEOx | Cell | DHA | Alcohol | 14-HDoHE | 15 | 0.41 (0.22, 0.76) | 0.34 (0.18, 0.62) | | -19% (-75, 160) | >0.80 |
| NEOx | Cell | DHA | Alcohol | 14-HDoHE | 30 | 0.44 (0.24, 0.81) | 0.33 (0.18, 0.61) | | -25% (-75, 129) | >0.80 |
| NEOx | Cell | DHA | Alcohol | 14-HDoHE | 60 | 0.64 (0.32, 1.27) | 0.41 (0.2, 0.81) | | -36% (-86, 192) | >0.80 |
| Esterified | Media | DHA | Alcohol | 14-HDoHE | 0 | 0.42 (0.23, 0.78) | 0.42 (0.23, 0.78) | >0.80 | -1% (-73, 259) | >0.80 |
| Esterified | Media | DHA | Alcohol | 14-HDoHE | 15 | 0.5 (0.29, 0.85) | 0.5 (0.29, 0.85) | | 0% (-65, 183) | >0.80 |
| Esterified | Media | DHA | Alcohol | 14-HDoHE | 30 | 0.54 (0.31, 0.94) | 0.55 (0.32, 0.95) | | 1% (-62, 170) | >0.80 |
| Esterified | Media | DHA | Alcohol | 14-HDoHE | 60 | 0.52 (0.27, 1) | 0.54 (0.28, 1.03) | | 2% (-76, 337) | >0.80 |
| NEOx | Media | DHA | Alcohol | 14-HDoHE | 0 | 0.57 (0.3, 1.08) | 0.49 (0.26, 0.94) | 0.27 | -13% (-76, 221) | >0.80 |
| NEOx | Media | DHA | Alcohol | 14-HDoHE | 15 | 0.41 (0.22, 0.79) | 0.4 (0.21, 0.77) | | -2% (-70, 220) | >0.80 |
| NEOx | Media | DHA | Alcohol | 14-HDoHE | 30 | 0.35 (0.19, 0.67) | 0.38 (0.2, 0.73) | | 9% (-66, 249) | >0.80 |
| NEOx | Media | DHA | Alcohol | 14-HDoHE | 60 | 0.39 (0.2, 0.75) | 0.53 (0.28, 1.02) | | 36% (-66, 445) | >0.80 |
| Esterified | Cell | DHA | Alcohol | 17-HDoHE | 0 | 3.31 (1.19, 9.18) | 5.75 (2.07, 15.96) | 0.31 | 74% (-80, 1379) | >0.80 |
| Esterified | Cell | DHA | Alcohol | 17-HDoHE | 15 | 2.93 (1.27, 6.75) | 3.92 (1.7, 9.04) | | 34% (-74, 591) | >0.80 |
| Esterified | Cell | DHA | Alcohol | 17-HDoHE | 30 | 3.5 (1.46, 8.37) | 3.61 (1.51, 8.62) | | 3% (-77, 372) | >0.80 |
| Esterified | Cell | DHA | Alcohol | 17-HDoHE | 60 | 12.28 (4.09, 36.85) | 7.5 (2.5, 22.5) | | -39% (-95, 621) | >0.80 |
| NEOx | Cell | DHA | Alcohol | 17-HDoHE | 0 | 1.18 (0.69, 2.02) | 0.73 (0.43, 1.24) | >0.80 | -38% (-79, 84) | 0.78 |
| NEOx | Cell | DHA | Alcohol | 17-HDoHE | 15 | 1.09 (0.65, 1.82) | 0.67 (0.4, 1.12) | | -39% (-77, 62) | 0.65 |
| NEOx | Cell | DHA | Alcohol | 17-HDoHE | 30 | 1.03 (0.61, 1.72) | 0.63 (0.37, 1.05) | | -39% (-76, 57) | 0.61 |
| NEOx | Cell | DHA | Alcohol | 17-HDoHE | 60 | 1 (0.58, 1.72) | 0.6 (0.35, 1.04) | | -40% (-82, 98) | >0.80 |
| Esterified | Media | DHA | Alcohol | 17-HDoHE | 0 | 0.92 (0.38, 2.24) | 1.21 (0.5, 2.94) | 0.37 | 31% (-76, 628) | >0.80 |
| Esterified | Media | DHA | Alcohol | 17-HDoHE | 15 | 0.81 (0.33, 2) | 0.98 (0.4, 2.4) | | 20% (-76, 514) | >0.80 |
| Esterified | Media | DHA | Alcohol | 17-HDoHE | 30 | 0.83 (0.34, 2.03) | 0.92 (0.38, 2.24) | | 10% (-78, 453) | >0.80 |
| Esterified | Media | DHA | Alcohol | 17-HDoHE | 60 | 1.35 (0.56, 3.28) | 1.25 (0.51, 3.03) | | -8% (-84, 446) | >0.80 |
| NEOx | Media | DHA | Alcohol | 17-HDoHE | 0 | 0.83 (0.43, 1.62) | 0.76 (0.39, 1.49) | 0.67 | -8% (-77, 262) | >0.80 |
| NEOx | Media | DHA | Alcohol | 17-HDoHE | 15 | 0.76 (0.4, 1.45) | 0.73 (0.38, 1.4) | | -3% (-71, 225) | >0.80 |
| NEOx | Media | DHA | Alcohol | 17-HDoHE | 30 | 0.72 (0.38, 1.38) | 0.73 (0.38, 1.4) | | 1% (-69, 230) | >0.80 |
| NEOx | Media | DHA | Alcohol | 17-HDoHE | 60 | 0.76 (0.38, 1.51) | 0.85 (0.43, 1.69) | | 12% (-75, 393) | >0.80 |
| Esterified | Cell | LA | Ketone | 9-KODE | 0 | 1.95 (0.68, 5.61) | 2.38 (0.83, 6.84) | 0.7 | 22% (-86, 1001) | >0.80 |
| Esterified | Cell | LA | Ketone | 9-KODE | 15 | 2.18 (0.98, 4.86) | 2.39 (1.07, 5.33) | | 10% (-78, 441) | >0.80 |
| Esterified | Cell | LA | Ketone | 9-KODE | 30 | 2.64 (1.12, 6.24) | 2.61 (1.1, 6.15) | | -1% (-77, 319) | >0.80 |
| Esterified | Cell | LA | Ketone | 9-KODE | 60 | 5 (1.59, 15.78) | 3.99 (1.26, 12.59) | | -20% (-94, 957) | >0.80 |
| NEOx | Cell | LA | Ketone | 9-KODE | 0 | 0.54 (0.09, 3.35) | 0.32 (0.05, 1.97) | 0.28 | -41% (-99, 2548) | >0.80 |
| NEOx | Cell | LA | Ketone | 9-KODE | 15 | 0.37 (0.09, 1.55) | 0.36 (0.09, 1.52) | | -2% (-94, 1595) | >0.80 |
| NEOx | Cell | LA | Ketone | 9-KODE | 30 | 0.21 (0.05, 0.96) | 0.34 (0.08, 1.58) | | 64% (-88, 2139) | >0.80 |
| NEOx | Cell | LA | Ketone | 9-KODE | 60 | 0.04 (0.01, 0.31) | 0.2 (0.03, 1.41) | | 356% (-95, 37951) | >0.80 |
| Esterified | Media | LA | Ketone | 9-KODE | 0 | 0.26 (0.09, 0.72) | 0.33 (0.12, 0.9) | 0.51 | 25% (-85, 960) | >0.80 |
| Esterified | Media | LA | Ketone | 9-KODE | 15 | 0.34 (0.14, 0.83) | 0.36 (0.15, 0.9) | | 8% (-81, 521) | >0.80 |
| Esterified | Media | LA | Ketone | 9-KODE | 30 | 0.42 (0.17, 1.07) | 0.39 (0.16, 0.99) | | -7% (-82, 389) | >0.80 |
| Esterified | Media | LA | Ketone | 9-KODE | 60 | 0.63 (0.21, 1.84) | 0.43 (0.15, 1.26) | | -32% (-94, 654) | >0.80 |
| NEOx | Media | LA | Ketone | 9-KODE | 0 | 0.08 (0.01, 0.47) | 0.09 (0.02, 0.58) | >0.80 | 23% (-97, 5197) | >0.80 |
| NEOx | Media | LA | Ketone | 9-KODE | 15 | 0.21 (0.06, 0.76) | 0.23 (0.06, 0.84) | | 10% (-92, 1358) | >0.80 |
| NEOx | Media | LA | Ketone | 9-KODE | 30 | 0.4 (0.1, 1.65) | 0.39 (0.1, 1.63) | | -1% (-90, 860) | >0.80 |
| NEOx | Media | LA | Ketone | 9-KODE | 60 | 0.5 (0.07, 3.67) | 0.39 (0.05, 2.92) | | -20% (-99, 6975) | >0.80 |
| Esterified | Cell | LA | Ketone | 13-KODE | 0 | 4.24 (1.32, 13.59) | 4.02 (1.26, 12.89) | >0.80 | -5% (-91, 870) | >0.80 |
| Esterified | Cell | LA | Ketone | 13-KODE | 15 | 6.83 (3.55, 13.15) | 6.12 (3.18, 11.77) | | -10% (-75, 226) | >0.80 |
| Esterified | Cell | LA | Ketone | 13-KODE | 30 | 8.08 (3.6, 18.14) | 6.83 (3.04, 15.32) | | -16% (-68, 121) | >0.80 |

| | | | | | | | | | | |
|---|---|---|---|---|---|---|---|---|---|---|
| Esterified | Cell | LA | Ketone | 13-KODE | 60 | 4.48 (1.21, 16.61) | 3.37 (0.91, 12.5) | | -25% (-96, 1265) | >0.80 |
| NEOx | Cell | LA | Ketone | 13-KODE | 0 | 1.6 (0.69, 3.71) | 2.41 (1.04, 5.58) | 0.06 | 50% (-72, 708) | >0.80 |
| NEOx | Cell | LA | Ketone | 13-KODE | 15 | 1.46 (0.9, 2.35) | 1.32 (0.82, 2.13) | | -9% (-65, 135) | >0.80 |
| NEOx | Cell | LA | Ketone | 13-KODE | 30 | 1.86 (1.04, 3.35) | 1.02 (0.57, 1.83) | | -45% (-73, 12) | 0.13 |
| NEOx | Cell | LA | Ketone | 13-KODE | 60 | 8.55 (3.33, 21.97) | 1.69 (0.66, 4.35) | | -80% (-98, 60) | 0.19 |
| Esterified | Media | LA | Ketone | 13-KODE | 0 | 1.12 (0.44, 2.87) | 1.04 (0.41, 2.68) | 0.06 | -7% (-87, 561) | >0.80 |
| Esterified | Media | LA | Ketone | 13-KODE | 15 | 1.86 (0.94, 3.69) | 1.04 (0.53, 2.07) | | -44% (-86, 121) | >0.80 |
| Esterified | Media | LA | Ketone | 13-KODE | 30 | 3.01 (1.42, 6.35) | 1.02 (0.48, 2.15) | | -66% (-90, 15) | 0.1 |
| Esterified | Media | LA | Ketone | 13-KODE | 60 | 7.21 (2.56, 20.29) | 0.89 (0.31, 2.49) | | -88% (-99, 26) | 0.09 |
| NEOx | Media | LA | Ketone | 13-KODE | 0 | 1.9 (0.69, 5.19) | 1.77 (0.65, 4.83) | 0.69 | -7% (-89, 659) | >0.80 |
| NEOx | Media | LA | Ketone | 13-KODE | 15 | 1.13 (0.53, 2.42) | 0.95 (0.44, 2.03) | | -16% (-82, 281) | >0.80 |
| NEOx | Media | LA | Ketone | 13-KODE | 30 | 0.98 (0.43, 2.22) | 0.74 (0.33, 1.67) | | -25% (-81, 195) | >0.80 |
| NEOx | Media | LA | Ketone | 13-KODE | 60 | 2.25 (0.75, 6.74) | 1.36 (0.46, 4.09) | | -39% (-95, 616) | >0.80 |
| Esterified | Cell | AA | Ketone | 5-KETE | 0 | 1.71 (0.58, 5.02) | 0.67 (0.23, 1.97) | 0.38 | -61% (-96, 259) | >0.80 |
| Esterified | Cell | AA | Ketone | 5-KETE | 15 | 2.6 (1.27, 5.32) | 1.34 (0.66, 2.74) | | -48% (-88, 117) | 0.74 |
| Esterified | Cell | AA | Ketone | 5-KETE | 30 | 3.52 (1.56, 7.91) | 2.38 (1.06, 5.36) | | -32% (-80, 130) | >0.80 |
| Esterified | Cell | AA | Ketone | 5-KETE | 60 | 4.46 (1.35, 14.78) | 5.22 (1.58, 17.28) | | 17% (-92, 1603) | >0.80 |
| NEOx | Cell | AA | Ketone | 5-KETE | 0 | 0.76 (0.14, 4) | 0.59 (0.11, 3.11) | 0.57 | -22% (-97, 2309) | >0.80 |
| NEOx | Cell | AA | Ketone | 5-KETE | 15 | 0.57 (0.18, 1.77) | 0.34 (0.11, 1.05) | | -41% (-94, 485) | >0.80 |
| NEOx | Cell | AA | Ketone | 5-KETE | 30 | 0.63 (0.18, 2.25) | 0.29 (0.08, 1.02) | | -55% (-94, 228) | >0.80 |
| NEOx | Cell | AA | Ketone | 5-KETE | 60 | 2.54 (0.4, 16.02) | 0.67 (0.11, 4.23) | | -74% (-100, 1538) | >0.80 |
| Esterified | Media | AA | Ketone | 5-KETE | 0 | 0.33 (0.07, 1.6) | 0.85 (0.17, 4.2) | 0.39 | 162% (-91, 7203) | >0.80 |
| Esterified | Media | AA | Ketone | 5-KETE | 15 | 0.55 (0.16, 1.9) | 1 (0.29, 3.45) | | 82% (-84, 2000) | >0.80 |
| Esterified | Media | AA | Ketone | 5-KETE | 30 | 0.89 (0.24, 3.3) | 1.12 (0.3, 4.17) | | 26% (-86, 1074) | >0.80 |
| Esterified | Media | AA | Ketone | 5-KETE | 60 | 1.95 (0.35, 11.04) | 1.19 (0.21, 6.73) | | -39% (-99, 2892) | >0.80 |
| NEOx | Media | AA | Ketone | 5-KETE | 0 | 1.26 (0.41, 3.86) | 1.57 (0.51, 4.8) | 0.65 | 24% (-88, 1133) | >0.80 |
| NEOx | Media | AA | Ketone | 5-KETE | 15 | 0.7 (0.33, 1.46) | 1 (0.48, 2.1) | | 44% (-68, 540) | >0.80 |
| NEOx | Media | AA | Ketone | 5-KETE | 30 | 0.48 (0.21, 1.11) | 0.79 (0.34, 1.84) | | 66% (-54, 494) | >0.80 |
| NEOx | Media | AA | Ketone | 5-KETE | 60 | 0.44 (0.13, 1.51) | 0.97 (0.28, 3.35) | | 122% (-86, 3467) | >0.80 |
| Esterified | Cell | AA | Ketone | 12-KETE | 0 | 0.45 (0.13, 1.52) | 0.56 (0.17, 1.9) | 0.71 | 25% (-90, 1488) | >0.80 |
| Esterified | Cell | AA | Ketone | 12-KETE | 15 | 0.55 (0.21, 1.44) | 0.62 (0.23, 1.61) | | 12% (-83, 650) | >0.80 |
| Esterified | Cell | AA | Ketone | 12-KETE | 30 | 0.82 (0.3, 2.27) | 0.82 (0.3, 2.26) | | 0% (-83, 473) | >0.80 |
| Esterified | Cell | AA | Ketone | 12-KETE | 60 | 3.32 (0.89, 12.34) | 2.64 (0.71, 9.8) | | -21% (-96, 1422) | >0.80 |
| NEOx | Cell | AA | Ketone | 12-KETE | 0 | 0.82 (0.25, 2.7) | 0.19 (0.06, 0.61) | 0.73 | -77% (-98, 160) | 0.44 |
| NEOx | Cell | AA | Ketone | 12-KETE | 15 | 0.53 (0.24, 1.2) | 0.13 (0.06, 0.3) | | -75% (-95, 30) | 0.13 |
| NEOx | Cell | AA | Ketone | 12-KETE | 30 | 0.45 (0.18, 1.12) | 0.13 (0.05, 0.32) | | -72% (-93, 18) | 0.1 |
| NEOx | Cell | AA | Ketone | 12-KETE | 60 | 0.73 (0.2, 2.71) | 0.26 (0.07, 0.97) | | -64% (-98, 578) | >0.80 |
| Esterified | Media | AA | Ketone | 12-KETE | 0 | 0.41 (0.17, 0.98) | 0.45 (0.19, 1.09) | >0.80 | 11% (-81, 558) | >0.80 |
| Esterified | Media | AA | Ketone | 12-KETE | 15 | 0.28 (0.16, 0.47) | 0.3 (0.18, 0.52) | | 10% (-63, 222) | >0.80 |
| Esterified | Media | AA | Ketone | 12-KETE | 30 | 0.23 (0.12, 0.43) | 0.25 (0.13, 0.47) | | 9% (-54, 159) | >0.80 |
| Esterified | Media | AA | Ketone | 12-KETE | 60 | 0.31 (0.11, 0.82) | 0.33 (0.12, 0.87) | | 7% (-88, 851) | >0.80 |
| NEOx | Media | AA | Ketone | 12-KETE | 0 | 0.23 (0.1, 0.57) | 0.45 (0.18, 1.1) | >0.80 | 91% (-70, 1099) | >0.80 |
| NEOx | Media | AA | Ketone | 12-KETE | 15 | 0.17 (0.09, 0.3) | 0.31 (0.17, 0.57) | | 86% (-44, 513) | 0.63 |
| NEOx | Media | AA | Ketone | 12-KETE | 30 | 0.17 (0.09, 0.33) | 0.3 (0.15, 0.59) | | 80% (-35, 400) | 0.5 |
| NEOx | Media | AA | Ketone | 12-KETE | 60 | 0.46 (0.17, 1.23) | 0.77 (0.29, 2.09) | | 70% (-82, 1467) | >0.80 |
| Esterified | Cell | AA | Ketone | 15-KETE | 0 | 0.15 (0.03, 0.65) | 0.4 (0.09, 1.78) | 0.49 | 173% (-88, 6157) | >0.80 |
| Esterified | Cell | AA | Ketone | 15-KETE | 15 | 0.21 (0.05, 0.84) | 0.46 (0.12, 1.85) | | 120% (-84, 2980) | >0.80 |
| Esterified | Cell | AA | Ketone | 15-KETE | 30 | 0.3 (0.08, 1.23) | 0.54 (0.13, 2.18) | | 78% (-86, 2125) | >0.80 |
| Esterified | Cell | AA | Ketone | 15-KETE | 60 | 0.63 (0.13, 3.04) | 0.73 (0.15, 3.51) | | 16% (-96, 3641) | >0.80 |
| NEOx | Cell | AA | Ketone | 15-KETE | 0 | 0.08 (0.02, 0.3) | 0.05 (0.01, 0.16) | 0.52 | -46% (-96, 642) | >0.80 |
| NEOx | Cell | AA | Ketone | 15-KETE | 15 | 0.08 (0.03, 0.21) | 0.03 (0.01, 0.09) | | -56% (-94, 198) | 0.79 |
| NEOx | Cell | AA | Ketone | 15-KETE | 30 | 0.09 (0.03, 0.26) | 0.03 (0.01, 0.09) | | -65% (-94, 103) | 0.47 |
| NEOx | Cell | AA | Ketone | 15-KETE | 60 | 0.26 (0.07, 1.02) | 0.06 (0.02, 0.24) | | -77% (-99, 401) | 0.71 |
| Esterified | Media | AA | Ketone | 15-KETE | 0 | 0.12 (0.04, 0.37) | 0.07 (0.02, 0.22) | 0.23 | -42% (-95, 537) | >0.80 |
| Esterified | Media | AA | Ketone | 15-KETE | 15 | 0.08 (0.03, 0.19) | 0.07 (0.03, 0.16) | | -16% (-85, 377) | >0.80 |
| Esterified | Media | AA | Ketone | 15-KETE | 30 | 0.07 (0.03, 0.17) | 0.08 (0.03, 0.21) | | 23% (-74, 488) | >0.80 |
| Esterified | Media | AA | Ketone | 15-KETE | 60 | 0.08 (0.02, 0.28) | 0.21 (0.06, 0.72) | | 160% (-85, 4252) | >0.80 |
| NEOx | Media | AA | Ketone | 15-KETE | 0 | 0.1 (0.03, 0.32) | 0.03 (0.01, 0.08) | 0.21 | -74% (-98, 185) | 0.53 |
| NEOx | Media | AA | Ketone | 15-KETE | 15 | 0.11 (0.04, 0.26) | 0.04 (0.02, 0.1) | | -62% (-94, 122) | 0.56 |
| NEOx | Media | AA | Ketone | 15-KETE | 30 | 0.11 (0.04, 0.28) | 0.06 (0.02, 0.15) | | -45% (-89, 179) | >0.80 |
| NEOx | Media | AA | Ketone | 15-KETE | 60 | 0.1 (0.03, 0.35) | 0.12 (0.03, 0.41) | | 18% (-93, 1860) | >0.80 |
| Esterified | Cell | AA | Peroxide | 12-HpETE | 0 | 0.33 (0.12, 0.89) | 0.38 (0.14, 1.04) | 0.61 | 17% (-85, 818) | >0.80 |
| Esterified | Cell | AA | Peroxide | 12-HpETE | 15 | 0.39 (0.19, 0.81) | 0.4 (0.19, 0.82) | | 2% (-76, 339) | >0.80 |
| Esterified | Cell | AA | Peroxide | 12-HpETE | 30 | 0.46 (0.21, 1.02) | 0.41 (0.19, 0.9) | | -11% (-76, 229) | >0.80 |
| Esterified | Cell | AA | Peroxide | 12-HpETE | 60 | 0.65 (0.22, 1.91) | 0.44 (0.15, 1.3) | | -32% (-94, 676) | >0.80 |
| NEOx | Cell | AA | Peroxide | 12-HpETE | 0 | 0.33 (0.16, 0.71) | 0.24 (0.11, 0.52) | 0.56 | -28% (-85, 258) | >0.80 |
| NEOx | Cell | AA | Peroxide | 12-HpETE | 15 | 0.34 (0.17, 0.66) | 0.22 (0.11, 0.43) | | -35% (-82, 139) | >0.80 |
| NEOx | Cell | AA | Peroxide | 12-HpETE | 30 | 0.38 (0.19, 0.75) | 0.22 (0.11, 0.44) | | -41% (-83, 101) | 0.8 |
| NEOx | Cell | AA | Peroxide | 12-HpETE | 60 | 0.64 (0.29, 1.45) | 0.31 (0.14, 0.7) | | -52% (-92, 194) | >0.80 |
| Esterified | Media | AA | Peroxide | 12-HpETE | 0 | 0.17 (0.07, 0.41) | 0.29 (0.12, 0.7) | 0.46 | 71% (-72, 958) | >0.80 |
| Esterified | Media | AA | Peroxide | 12-HpETE | 15 | 0.25 (0.12, 0.53) | 0.37 (0.17, 0.79) | | 47% (-66, 536) | >0.80 |
| Esterified | Media | AA | Peroxide | 12-HpETE | 30 | 0.32 (0.15, 0.69) | 0.4 (0.19, 0.87) | | 27% (-68, 404) | >0.80 |
| Esterified | Media | AA | Peroxide | 12-HpETE | 60 | 0.32 (0.13, 0.81) | 0.3 (0.12, 0.76) | | -6% (-88, 646) | >0.80 |
| NEOx | Media | AA | Peroxide | 12-HpETE | 0 | 0.23 (0.14, 0.4) | 0.35 (0.21, 0.61) | 0.7 | 52% (-50, 361) | >0.80 |
| NEOx | Media | AA | Peroxide | 12-HpETE | 15 | 0.21 (0.15, 0.3) | 0.3 (0.21, 0.43) | | 43% (-30, 192) | 0.65 |
| NEOx | Media | AA | Peroxide | 12-HpETE | 30 | 0.21 (0.14, 0.31) | 0.28 (0.19, 0.42) | | 35% (-26, 146) | 0.66 |
| NEOx | Media | AA | Peroxide | 12-HpETE | 60 | 0.25 (0.14, 0.46) | 0.3 (0.16, 0.55) | | 20% (-69, 360) | >0.80 |
| Esterified | Cell | AA | Peroxide | 15-HpETE | 0 | 3.27 (1.34, 7.97) | 3.03 (1.24, 7.4) | 0.36 | -7% (-84, 453) | >0.80 |
| Esterified | Cell | AA | Peroxide | 15-HpETE | 15 | 3.1 (1.28, 7.55) | 2.55 (1.05, 6.21) | | -18% (-84, 324) | >0.80 |
| Esterified | Cell | AA | Peroxide | 15-HpETE | 30 | 3.17 (1.31, 7.68) | 2.31 (0.96, 5.6) | | -27% (-85, 265) | >0.80 |
| Esterified | Cell | AA | Peroxide | 15-HpETE | 60 | 4.16 (1.69, 10.24) | 2.38 (0.97, 5.87) | | -43% (-91, 280) | >0.80 |
| NEOx | Cell | AA | Peroxide | 15-HpETE | 0 | 2.42 (1.22, 4.8) | 1.3 (0.66, 2.59) | >0.80 | -46% (-87, 120) | 0.78 |
| NEOx | Cell | AA | Peroxide | 15-HpETE | 15 | 2.78 (1.78, 4.32) | 1.53 (0.98, 2.38) | | -45% (-77, 34) | 0.34 |
| NEOx | Cell | AA | Peroxide | 15-HpETE | 30 | 2.83 (1.7, 4.71) | 1.59 (0.95, 2.64) | | -44% (-73, 19) | 0.2 |
| NEOx | Cell | AA | Peroxide | 15-HpETE | 60 | 2.05 (0.95, 4.39) | 1.19 (0.56, 2.56) | | -42% (-89, 221) | >0.80 |

| | | | | | | | | | | |
|---|---|---|---|---|---|---|---|---|---|---|
| Esterified | Media | AA | Peroxide | 15-HpETE | 0 | 2.75 (0.8, 9.44) | 2.45 (0.71, 8.38) | 0.6 | -11% (-93, 957) | >0.80 |
| Esterified | Media | AA | Peroxide | 15-HpETE | 15 | 1.96 (0.58, 6.67) | 1.58 (0.46, 5.38) | | -19% (-92, 677) | >0.80 |
| Esterified | Media | AA | Peroxide | 15-HpETE | 30 | 1.74 (0.51, 5.88) | 1.27 (0.38, 4.31) | | -27% (-92, 575) | >0.80 |
| Esterified | Media | AA | Peroxide | 15-HpETE | 60 | 2.64 (0.76, 9.22) | 1.6 (0.46, 5.57) | | -40% (-96, 741) | >0.80 |
| NEOx | Media | AA | Peroxide | 15-HpETE | 0 | 2.89 (1.05, 7.94) | 1.38 (0.5, 3.8) | 0.6 | -52% (-94, 279) | >0.80 |
| NEOx | Media | AA | Peroxide | 15-HpETE | 15 | 2.71 (1.01, 7.25) | 1.42 (0.53, 3.8) | | -48% (-92, 231) | >0.80 |
| NEOx | Media | AA | Peroxide | 15-HpETE | 30 | 2.55 (0.95, 6.8) | 1.46 (0.55, 3.91) | | -43% (-90, 245) | >0.80 |
| NEOx | Media | AA | Peroxide | 15-HpETE | 60 | 2.26 (0.8, 6.38) | 1.56 (0.55, 4.4) | | -31% (-93, 546) | >0.80 |
| Esterified | Cell | DHA | Triol | Resolvin D1 | 0 | 90 (31, 260) | 42 (15, 121) | 0.64 | -54% (-95, 323) | >0.80 |
| Esterified | Cell | DHA | Triol | Resolvin D1 | 15 | 106 (41, 272) | 55 (21, 141) | | -48% (-92, 218) | >0.80 |
| Esterified | Cell | DHA | Triol | Resolvin D1 | 30 | 112 (43, 291) | 64 (25, 168) | | -42% (-90, 224) | >0.80 |
| Esterified | Cell | DHA | Triol | Resolvin D1 | 60 | 89 (29, 272) | 64 (21, 194) | | -28% (-94, 757) | >0.80 |
| NEOx | Cell | DHA | Triol | Resolvin D1 | 0 | 53.47 (16.76, 170.54) | 55.79 (17.49, 177.94) | >0.80 | 4% (-91, 1070) | >0.80 |
| NEOx | Cell | DHA | Triol | Resolvin D1 | 15 | 51.01 (17.53, 148.46) | 54.05 (18.57, 157.3) | | 6% (-86, 714) | >0.80 |
| NEOx | Cell | DHA | Triol | Resolvin D1 | 30 | 45.91 (15.63, 134.88) | 49.4 (16.82, 145.13) | | 8% (-85, 660) | >0.80 |
| NEOx | Cell | DHA | Triol | Resolvin D1 | 60 | 31.24 (9.29, 105.04) | 34.67 (10.31, 116.55) | | 11% (-92, 1520) | >0.80 |
| Esterified | Media | DHA | Triol | Resolvin D1 | 0 | 48.47 (12.42, 189.16) | 55.97 (14.34, 218.42) | >0.80 | 15% (-93, 1790) | >0.80 |
| Esterified | Media | DHA | Triol | Resolvin D1 | 15 | 36.87 (9.89, 137.39) | 41.21 (11.06, 153.59) | | 12% (-91, 1220) | >0.80 |
| Esterified | Media | DHA | Triol | Resolvin D1 | 30 | 33.46 (8.99, 124.5) | 36.21 (9.73, 134.73) | | 8% (-90, 1091) | >0.80 |
| Esterified | Media | DHA | Triol | Resolvin D1 | 60 | 46.83 (11.55, 189.85) | 47.5 (11.72, 192.56) | | 1% (-95, 2002) | >0.80 |
| NEOx | Media | DHA | Triol | Resolvin D1 | 0 | 76.73 (32.62, 180.48) | 55.99 (23.8, 131.69) | 0.59 | -27% (-88, 339) | >0.80 |
| NEOx | Media | DHA | Triol | Resolvin D1 | 15 | 67.45 (32.41, 140.35) | 44.21 (21.25, 92) | | -34% (-84, 173) | >0.80 |
| NEOx | Media | DHA | Triol | Resolvin D1 | 30 | 57.66 (27.14, 122.5) | 33.96 (15.98, 72.14) | | -41% (-85, 125) | >0.80 |
| NEOx | Media | DHA | Triol | Resolvin D1 | 60 | 38.76 (15.58, 96.44) | 18.42 (7.41, 45.84) | | -52% (-94, 266) | >0.80 |
| Esterified | Cell | LA | Epoxide | 9(10)-EpOME | 0 | 2.6 (1.26, 5.35) | 2.4 (1.16, 4.94) | 0.37 | -8% (-79, 311) | >0.80 |
| Esterified | Cell | LA | Epoxide | 9(10)-EpOME | 15 | 2.79 (1.41, 5.53) | 2.27 (1.14, 4.5) | | -19% (-78, 197) | >0.80 |
| Esterified | Cell | LA | Epoxide | 9(10)-EpOME | 30 | 3.1 (1.56, 6.15) | 2.22 (1.12, 4.42) | | -28% (-79, 151) | >0.80 |
| Esterified | Cell | LA | Epoxide | 9(10)-EpOME | 60 | 4.23 (2, 8.94) | 2.36 (1.12, 4.99) | | -44% (-89, 187) | >0.80 |
| NEOx | Cell | LA | Epoxide | 9(10)-EpOME | 0 | 0.64 (0.19, 2.17) | 0.55 (0.16, 1.85) | 0.73 | -14% (-93, 891) | >0.80 |
| NEOx | Cell | LA | Epoxide | 9(10)-EpOME | 15 | 0.58 (0.17, 1.96) | 0.47 (0.14, 1.58) | | -20% (-91, 657) | >0.80 |
| NEOx | Cell | LA | Epoxide | 9(10)-EpOME | 30 | 0.58 (0.17, 1.94) | 0.44 (0.13, 1.47) | | -25% (-92, 581) | >0.80 |
| NEOx | Cell | LA | Epoxide | 9(10)-EpOME | 60 | 0.75 (0.22, 2.6) | 0.5 (0.15, 1.73) | | -33% (-95, 799) | >0.80 |
| Esterified | Media | LA | Epoxide | 9(10)-EpOME | 0 | 0.75 (0.4, 1.42) | 0.56 (0.3, 1.05) | 0.42 | -26% (-80, 172) | >0.80 |
| Esterified | Media | LA | Epoxide | 9(10)-EpOME | 15 | 0.71 (0.39, 1.31) | 0.58 (0.32, 1.07) | | -18% (-74, 156) | >0.80 |
| Esterified | Media | LA | Epoxide | 9(10)-EpOME | 30 | 0.71 (0.39, 1.31) | 0.64 (0.35, 1.17) | | -10% (-70, 171) | >0.80 |
| Esterified | Media | LA | Epoxide | 9(10)-EpOME | 60 | 0.84 (0.44, 1.6) | 0.91 (0.47, 1.74) | | 8% (-74, 344) | >0.80 |
| NEOx | Media | LA | Epoxide | 9(10)-EpOME | 0 | 0.51 (0.27, 0.97) | 0.34 (0.18, 0.65) | 0.24 | -33% (-82, 153) | >0.80 |
| NEOx | Media | LA | Epoxide | 9(10)-EpOME | 15 | 0.53 (0.32, 0.89) | 0.43 (0.26, 0.72) | | -19% (-70, 123) | >0.80 |
| NEOx | Media | LA | Epoxide | 9(10)-EpOME | 30 | 0.52 (0.31, 0.89) | 0.51 (0.3, 0.88) | | -2% (-61, 150) | >0.80 |
| NEOx | Media | LA | Epoxide | 9(10)-EpOME | 60 | 0.44 (0.22, 0.87) | 0.63 (0.32, 1.25) | | 44% (-69, 568) | >0.80 |
| Esterified | Cell | LA | Epoxide | 12(13)-EpOME | 0 | 2.14 (1.1, 4.17) | 2.71 (1.39, 5.29) | 0.05 | 27% (-67, 386) | >0.80 |
| Esterified | Cell | LA | Epoxide | 12(13)-EpOME | 15 | 2.34 (1.2, 4.54) | 2.4 (1.24, 4.65) | | 3% (-70, 250) | >0.80 |
| Esterified | Cell | LA | Epoxide | 12(13)-EpOME | 30 | 2.83 (1.46, 5.48) | 2.35 (1.21, 4.54) | | -17% (-75, 176) | >0.80 |
| Esterified | Cell | LA | Epoxide | 12(13)-EpOME | 60 | 5.62 (2.85, 11.08) | 3.05 (1.55, 6.02) | | -46% (-87, 127) | >0.80 |
| NEOx | Cell | LA | Epoxide | 12(13)-EpOME | 0 | 0.94 (0.39, 2.29) | 0.45 (0.19, 1.09) | >0.80 | -52% (-93, 206) | >0.80 |
| NEOx | Cell | LA | Epoxide | 12(13)-EpOME | 15 | 0.82 (0.39, 1.71) | 0.4 (0.19, 0.85) | | -51% (-88, 110) | 0.69 |
| NEOx | Cell | LA | Epoxide | 12(13)-EpOME | 30 | 0.77 (0.36, 1.66) | 0.4 (0.18, 0.85) | | -49% (-87, 97) | 0.67 |
| NEOx | Cell | LA | Epoxide | 12(13)-EpOME | 60 | 0.91 (0.35, 2.37) | 0.5 (0.19, 1.3) | | -45% (-93, 363) | >0.80 |
| Esterified | Media | LA | Epoxide | 12(13)-EpOME | 0 | 0.4 (0.18, 0.87) | 0.58 (0.27, 1.27) | 0.04 | 45% (-71, 619) | >0.80 |
| Esterified | Media | LA | Epoxide | 12(13)-EpOME | 15 | 0.48 (0.23, 1.03) | 0.52 (0.24, 1.1) | | 7% (-74, 340) | >0.80 |
| Esterified | Media | LA | Epoxide | 12(13)-EpOME | 30 | 0.65 (0.31, 1.39) | 0.51 (0.24, 1.09) | | -22% (-80, 210) | >0.80 |
| Esterified | Media | LA | Epoxide | 12(13)-EpOME | 60 | 1.65 (0.74, 3.66) | 0.69 (0.31, 1.54) | | -58% (-93, 137) | 0.66 |
| NEOx | Media | LA | Epoxide | 12(13)-EpOME | 0 | 0.68 (0.23, 1.95) | 0.4 (0.14, 1.16) | 0.78 | -40% (-93, 410) | >0.80 |
| NEOx | Media | LA | Epoxide | 12(13)-EpOME | 15 | 0.52 (0.18, 1.46) | 0.32 (0.11, 0.91) | | -38% (-91, 333) | >0.80 |
| NEOx | Media | LA | Epoxide | 12(13)-EpOME | 30 | 0.46 (0.16, 1.3) | 0.3 (0.11, 0.85) | | -35% (-90, 334) | >0.80 |
| NEOx | Media | LA | Epoxide | 12(13)-EpOME | 60 | 0.58 (0.2, 1.71) | 0.42 (0.14, 1.22) | | -28% (-93, 616) | >0.80 |
| Esterified | Cell | AA | Epoxide | 14(15)-EpETrE | 0 | 0.28 (0.11, 0.72) | 0.32 (0.13, 0.84) | 0.29 | 17% (-84, 751) | >0.80 |
| Esterified | Cell | AA | Epoxide | 14(15)-EpETrE | 15 | 0.3 (0.14, 0.63) | 0.27 (0.13, 0.57) | | -10% (-79, 292) | >0.80 |
| Esterified | Cell | AA | Epoxide | 14(15)-EpETrE | 30 | 0.35 (0.16, 0.77) | 0.24 (0.11, 0.54) | | -31% (-82, 166) | >0.80 |
| Esterified | Cell | AA | Epoxide | 14(15)-EpETrE | 60 | 0.62 (0.22, 1.74) | 0.25 (0.09, 0.71) | | -59% (-96, 315) | >0.80 |
| NEOx | Cell | AA | Epoxide | 14(15)-EpETrE | 0 | 0.15 (0.09, 0.23) | 0.13 (0.08, 0.21) | 0.72 | -9% (-65, 140) | >0.80 |
| NEOx | Cell | AA | Epoxide | 14(15)-EpETrE | 15 | 0.18 (0.13, 0.25) | 0.16 (0.11, 0.22) | | -13% (-56, 72) | >0.80 |
| NEOx | Cell | AA | Epoxide | 14(15)-EpETrE | 30 | 0.19 (0.13, 0.28) | 0.16 (0.11, 0.23) | | -17% (-55, 53) | >0.80 |
| NEOx | Cell | AA | Epoxide | 14(15)-EpETrE | 60 | 0.15 (0.09, 0.26) | 0.12 (0.07, 0.19) | | -24% (-76, 139) | >0.80 |
| Esterified | Media | AA | Epoxide | 14(15)-EpETrE | 0 | 0.12 (0.06, 0.22) | 0.1 (0.05, 0.18) | 0.15 | -19% (-78, 193) | >0.80 |
| Esterified | Media | AA | Epoxide | 14(15)-EpETrE | 15 | 0.18 (0.1, 0.32) | 0.12 (0.07, 0.22) | | -31% (-78, 116) | >0.80 |
| Esterified | Media | AA | Epoxide | 14(15)-EpETrE | 30 | 0.22 (0.12, 0.41) | 0.13 (0.07, 0.24) | | -41% (-81, 78) | 0.7 |
| Esterified | Media | AA | Epoxide | 14(15)-EpETrE | 60 | 0.22 (0.12, 0.42) | 0.09 (0.05, 0.18) | | -58% (-89, 70) | 0.43 |
| NEOx | Media | AA | Epoxide | 14(15)-EpETrE | 0 | 0.15 (0.09, 0.24) | 0.12 (0.08, 0.2) | 0.3 | -17% (-69, 123) | >0.80 |
| NEOx | Media | AA | Epoxide | 14(15)-EpETrE | 15 | 0.13 (0.08, 0.21) | 0.12 (0.07, 0.19) | | -9% (-62, 118) | >0.80 |
| NEOx | Media | AA | Epoxide | 14(15)-EpETrE | 30 | 0.12 (0.08, 0.19) | 0.12 (0.08, 0.19) | | -1% (-58, 133) | >0.80 |
| NEOx | Media | AA | Epoxide | 14(15)-EpETrE | 60 | 0.12 (0.07, 0.2) | 0.14 (0.09, 0.23) | | 19% (-59, 245) | >0.80 |
| Esterified | Cell | EPA | Epoxide | 8(9)-EpETE | 0 | 0.26 (0.11, 0.61) | 0.3 (0.13, 0.7) | 0.49 | 15% (-79, 519) | >0.80 |
| Esterified | Cell | EPA | Epoxide | 8(9)-EpETE | 15 | 0.3 (0.13, 0.71) | 0.32 (0.13, 0.76) | | 7% (-78, 422) | >0.80 |
| Esterified | Cell | EPA | Epoxide | 8(9)-EpETE | 30 | 0.33 (0.14, 0.77) | 0.33 (0.14, 0.77) | | 0% (-79, 376) | >0.80 |
| Esterified | Cell | EPA | Epoxide | 8(9)-EpETE | 60 | 0.33 (0.14, 0.78) | 0.29 (0.12, 0.68) | | -14% (-85, 400) | >0.80 |
| NEOx | Cell | EPA | Epoxide | 8(9)-EpETE | 0 | 0.27 (0.14, 0.52) | 0.29 (0.15, 0.55) | 0.21 | 5% (-72, 297) | >0.80 |
| NEOx | Cell | EPA | Epoxide | 8(9)-EpETE | 15 | 0.35 (0.19, 0.65) | 0.32 (0.17, 0.59) | | -9% (-72, 192) | >0.80 |
| NEOx | Cell | EPA | Epoxide | 8(9)-EpETE | 30 | 0.42 (0.23, 0.79) | 0.33 (0.18, 0.62) | | -22% (-75, 144) | >0.80 |
| NEOx | Cell | EPA | Epoxide | 8(9)-EpETE | 60 | 0.52 (0.27, 1.02) | 0.31 (0.16, 0.59) | | -42% (-86, 146) | >0.80 |
| Esterified | Media | EPA | Epoxide | 8(9)-EpETE | 0 | 0.37 (0.2, 0.68) | 0.36 (0.19, 0.65) | 0.07 | -4% (-72, 232) | >0.80 |
| Esterified | Media | EPA | Epoxide | 8(9)-EpETE | 15 | 0.39 (0.22, 0.72) | 0.31 (0.17, 0.57) | | -20% (-74, 142) | >0.80 |
| Esterified | Media | EPA | Epoxide | 8(9)-EpETE | 30 | 0.43 (0.24, 0.78) | 0.28 (0.16, 0.51) | | -34% (-78, 94) | >0.80 |
| Esterified | Media | EPA | Epoxide | 8(9)-EpETE | 60 | 0.55 (0.3, 1.02) | 0.25 (0.13, 0.46) | | -55% (-88, 70) | 0.45 |
| NEOx | Media | EPA | Epoxide | 8(9)-EpETE | 0 | 0.23 (0.15, 0.36) | 0.23 (0.15, 0.36) | >0.80 | 0% (-59, 145) | >0.80 |

| | | | | | | | | | | |
|---|---|---|---|---|---|---|---|---|---|---|
| NEOx | Media | EPA | Epoxide | 8(9)-EpETE | 15 | 0.3 (0.21, 0.43) | 0.31 (0.22, 0.44) | | 2% (-49, 102) | >0.80 |
| NEOx | Media | EPA | Epoxide | 8(9)-EpETE | 30 | 0.35 (0.24, 0.5) | 0.36 (0.25, 0.52) | | 3% (-46, 95) | >0.80 |
| NEOx | Media | EPA | Epoxide | 8(9)-EpETE | 60 | 0.32 (0.2, 0.5) | 0.33 (0.21, 0.53) | | 5% (-62, 195) | >0.80 |
| Esterified | Cell | EPA | Epoxide | 14(15)-EpETE | 0 | 2.14 (1.4, 3.3) | 2.16 (1.41, 3.33) | 0.46 | 1% (-58, 141) | >0.80 |
| Esterified | Cell | EPA | Epoxide | 14(15)-EpETE | 15 | 2.18 (1.68, 2.82) | 2 (1.54, 2.59) | | -8% (-45, 54) | >0.80 |
| Esterified | Cell | EPA | Epoxide | 14(15)-EpETE | 30 | 2.22 (1.63, 3.02) | 1.85 (1.36, 2.51) | | -17% (-45, 26) | 0.78 |
| Esterified | Cell | EPA | Epoxide | 14(15)-EpETE | 60 | 2.31 (1.42, 3.73) | 1.59 (0.98, 2.57) | | -31% (-76, 101) | >0.80 |
| NEOx | Cell | EPA | Epoxide | 14(15)-EpETE | 0 | 0.98 (0.39, 2.49) | 1.13 (0.45, 2.85) | 0.34 | 15% (-84, 701) | >0.80 |
| NEOx | Cell | EPA | Epoxide | 14(15)-EpETE | 15 | 1.54 (0.73, 3.27) | 1.41 (0.67, 3) | | -8% (-79, 302) | >0.80 |
| NEOx | Cell | EPA | Epoxide | 14(15)-EpETE | 30 | 1.85 (0.84, 4.06) | 1.36 (0.62, 2.98) | | -27% (-81, 187) | >0.80 |
| NEOx | Cell | EPA | Epoxide | 14(15)-EpETE | 60 | 1.2 (0.44, 3.25) | 0.56 (0.21, 1.52) | | -53% (-95, 343) | >0.80 |
| Esterified | Media | EPA | Epoxide | 14(15)-EpETE | 0 | 0.99 (0.35, 2.8) | 2.4 (0.85, 6.76) | 0.23 | 142% (-70, 1865) | >0.80 |
| Esterified | Media | EPA | Epoxide | 14(15)-EpETE | 15 | 1.18 (0.64, 2.19) | 1.94 (1.05, 3.61) | | 65% (-52, 466) | >0.80 |
| Esterified | Media | EPA | Epoxide | 14(15)-EpETE | 30 | 1.46 (0.7, 3.05) | 1.63 (0.78, 3.42) | | 12% (-58, 197) | >0.80 |
| Esterified | Media | EPA | Epoxide | 14(15)-EpETE | 60 | 2.46 (0.77, 7.87) | 1.28 (0.4, 4.09) | | -48% (-96, 589) | >0.80 |
| NEOx | Media | EPA | Epoxide | 14(15)-EpETE | 0 | 1.48 (0.66, 3.32) | 0.95 (0.43, 2.14) | 0.54 | -36% (-88, 245) | >0.80 |
| NEOx | Media | EPA | Epoxide | 14(15)-EpETE | 15 | 1.68 (0.91, 3.11) | 1.23 (0.67, 2.28) | | -27% (-78, 149) | >0.80 |
| NEOx | Media | EPA | Epoxide | 14(15)-EpETE | 30 | 1.73 (0.89, 3.34) | 1.44 (0.75, 2.79) | | -16% (-72, 154) | >0.80 |
| NEOx | Media | EPA | Epoxide | 14(15)-EpETE | 60 | 1.37 (0.57, 3.29) | 1.49 (0.62, 3.57) | | 9% (-85, 680) | >0.80 |
| Esterified | Cell | EPA | Epoxide | 17(18)-EpETE | 0 | 3.45 (1.25, 9.55) | 1.33 (0.48, 3.67) | 0.76 | -62% (-95, 202) | 0.73 |
| Esterified | Cell | EPA | Epoxide | 17(18)-EpETE | 15 | 4.32 (2.31, 8.08) | 1.82 (0.98, 3.41) | | -58% (-88, 48) | 0.31 |
| Esterified | Cell | EPA | Epoxide | 17(18)-EpETE | 30 | 4.49 (2.15, 9.37) | 2.08 (1, 4.34) | | -54% (-83, 28) | 0.22 |
| Esterified | Cell | EPA | Epoxide | 17(18)-EpETE | 60 | 2.77 (0.89, 8.63) | 1.55 (0.5, 4.83) | | -44% (-96, 603) | >0.80 |
| NEOx | Cell | EPA | Epoxide | 17(18)-EpETE | 0 | 5.04 (1.29, 19.76) | 2.95 (0.75, 11.57) | 0.59 | -41% (-96, 868) | >0.80 |
| NEOx | Cell | EPA | Epoxide | 17(18)-EpETE | 15 | 2.9 (0.78, 10.82) | 1.49 (0.4, 5.56) | | -49% (-96, 510) | >0.80 |
| NEOx | Cell | EPA | Epoxide | 17(18)-EpETE | 30 | 2.43 (0.65, 9.07) | 1.1 (0.29, 4.09) | | -55% (-96, 399) | >0.80 |
| NEOx | Cell | EPA | Epoxide | 17(18)-EpETE | 60 | 5.32 (1.31, 21.69) | 1.85 (0.45, 7.53) | | -65% (-98, 629) | >0.80 |
| Esterified | Media | EPA | Epoxide | 17(18)-EpETE | 0 | 3.08 (1.29, 7.36) | 1.75 (0.73, 4.18) | 0.46 | -43% (-90, 228) | >0.80 |
| Esterified | Media | EPA | Epoxide | 17(18)-EpETE | 15 | 3.02 (1.81, 5.05) | 2.08 (1.25, 3.48) | | -31% (-75, 92) | >0.80 |
| Esterified | Media | EPA | Epoxide | 17(18)-EpETE | 30 | 3.4 (1.83, 6.3) | 2.84 (1.53, 5.26) | | -16% (-63, 86) | >0.80 |
| Esterified | Media | EPA | Epoxide | 17(18)-EpETE | 60 | 6.44 (2.43, 17.1) | 7.92 (2.98, 21.03) | | 23% (-86, 973) | >0.80 |
| NEOx | Media | EPA | Epoxide | 17(18)-EpETE | 0 | 3.06 (1.37, 6.85) | 3.48 (1.55, 7.78) | 0.75 | 14% (-79, 511) | >0.80 |
| NEOx | Media | EPA | Epoxide | 17(18)-EpETE | 15 | 2.35 (1.14, 4.88) | 2.83 (1.36, 5.87) | | 20% (-70, 387) | >0.80 |
| NEOx | Media | EPA | Epoxide | 17(18)-EpETE | 30 | 2.17 (1.04, 4.55) | 2.76 (1.32, 5.78) | | 27% (-67, 383) | >0.80 |
| NEOx | Media | EPA | Epoxide | 17(18)-EpETE | 60 | 3.2 (1.37, 7.47) | 4.55 (1.95, 10.6) | | 42% (-78, 829) | >0.80 |
| Esterified | Cell | DHA | Epoxide | 10(11)-EpDPE | 0 | 1.09 (0.49, 2.4) | 1.57 (0.71, 3.45) | 0.32 | 44% (-72, 647) | >0.80 |
| Esterified | Cell | DHA | Epoxide | 10(11)-EpDPE | 15 | 2.35 (1.29, 4.27) | 2.75 (1.51, 4.99) | | 17% (-64, 284) | >0.80 |
| Esterified | Cell | DHA | Epoxide | 10(11)-EpDPE | 30 | 3.93 (2.07, 7.46) | 3.72 (1.96, 7.06) | | -5% (-68, 177) | >0.80 |
| Esterified | Cell | DHA | Epoxide | 10(11)-EpDPE | 60 | 5.1 (2.16, 12.06) | 3.17 (1.34, 7.5) | | -38% (-91, 331) | >0.80 |
| NEOx | Cell | DHA | Epoxide | 10(11)-EpDPE | 0 | 0.5 (0.23, 1.11) | 0.7 (0.32, 1.55) | 0.05 | 40% (-74, 642) | >0.80 |
| NEOx | Cell | DHA | Epoxide | 10(11)-EpDPE | 15 | 0.99 (0.49, 2) | 0.95 (0.47, 1.91) | | -4% (-75, 272) | >0.80 |
| NEOx | Cell | DHA | Epoxide | 10(11)-EpDPE | 30 | 1.6 (0.78, 3.27) | 1.04 (0.51, 2.14) | | -35% (-82, 137) | >0.80 |
| NEOx | Cell | DHA | Epoxide | 10(11)-EpDPE | 60 | 2.25 (0.97, 5.22) | 0.69 (0.3, 1.6) | | -69% (-95, 101) | 0.40 |
| Esterified | Media | DHA | Epoxide | 10(11)-EpDPE | 0 | 1.72 (0.64, 4.67) | 0.62 (0.23, 1.69) | 0.45 | -64% (-96, 193) | 0.69 |
| Esterified | Media | DHA | Epoxide | 10(11)-EpDPE | 15 | 1.42 (0.59, 3.45) | 0.61 (0.25, 1.48) | | -57% (-92, 137) | 0.67 |
| Esterified | Media | DHA | Epoxide | 10(11)-EpDPE | 30 | 1.26 (0.51, 3.1) | 0.64 (0.26, 1.58) | | -49% (-90, 157) | >0.80 |
| Esterified | Media | DHA | Epoxide | 10(11)-EpDPE | 60 | 1.22 (0.42, 3.5) | 0.87 (0.3, 2.51) | | -28% (-93, 652) | >0.80 |
| NEOx | Media | DHA | Epoxide | 10(11)-EpDPE | 0 | 1.63 (0.54, 4.97) | 0.61 (0.2, 1.85) | 0.11 | -63% (-96, 256) | 0.78 |
| NEOx | Media | DHA | Epoxide | 10(11)-EpDPE | 15 | 1.18 (0.39, 3.52) | 0.59 (0.2, 1.77) | | -50% (-93, 286) | >0.80 |
| NEOx | Media | DHA | Epoxide | 10(11)-EpDPE | 30 | 0.96 (0.32, 2.84) | 0.65 (0.22, 1.93) | | -32% (-91, 398) | >0.80 |
| NEOx | Media | DHA | Epoxide | 10(11)-EpDPE | 60 | 0.89 (0.29, 2.77) | 1.11 (0.36, 3.45) | | 24% (-89, 1301) | >0.80 |
| Esterified | Cell | DHA | Epoxide | 13(14)-EpDPE | 0 | 0.76 (0.2, 2.89) | 0.82 (0.22, 3.12) | >0.80 | 8% (-92, 1414) | >0.80 |
| Esterified | Cell | DHA | Epoxide | 13(14)-EpDPE | 15 | 0.83 (0.22, 3.16) | 0.86 (0.23, 3.29) | | 4% (-91, 1114) | >0.80 |
| Esterified | Cell | DHA | Epoxide | 13(14)-EpDPE | 30 | 0.97 (0.26, 3.67) | 0.97 (0.26, 3.67) | | 0% (-91, 1025) | >0.80 |
| Esterified | Cell | DHA | Epoxide | 13(14)-EpDPE | 60 | 1.65 (0.43, 6.33) | 1.53 (0.4, 5.88) | | -7% (-94, 1395) | >0.80 |
| NEOx | Cell | DHA | Epoxide | 13(14)-EpDPE | 0 | 0.26 (0.15, 0.44) | 0.28 (0.16, 0.49) | 0.06 | 11% (-64, 245) | >0.80 |
| NEOx | Cell | DHA | Epoxide | 13(14)-EpDPE | 15 | 0.22 (0.13, 0.38) | 0.2 (0.12, 0.35) | | -8% (-66, 150) | >0.80 |
| NEOx | Cell | DHA | Epoxide | 13(14)-EpDPE | 30 | 0.22 (0.13, 0.37) | 0.17 (0.1, 0.28) | | -24% (-72, 101) | >0.80 |
| NEOx | Cell | DHA | Epoxide | 13(14)-EpDPE | 60 | 0.3 (0.17, 0.52) | 0.15 (0.09, 0.27) | | -49% (-85, 75) | 0.57 |
| Esterified | Media | DHA | Epoxide | 13(14)-EpDPE | 0 | 0.3 (0.18, 0.5) | 0.3 (0.18, 0.5) | >0.80 | 0% (-65, 185) | >0.80 |
| Esterified | Media | DHA | Epoxide | 13(14)-EpDPE | 15 | 0.28 (0.2, 0.4) | 0.28 (0.2, 0.39) | | -3% (-52, 95) | >0.80 |
| Esterified | Media | DHA | Epoxide | 13(14)-EpDPE | 30 | 0.28 (0.19, 0.42) | 0.27 (0.18, 0.39) | | -5% (-48, 73) | >0.80 |
| Esterified | Media | DHA | Epoxide | 13(14)-EpDPE | 60 | 0.34 (0.19, 0.6) | 0.3 (0.17, 0.53) | | -11% (-75, 215) | >0.80 |
| NEOx | Media | DHA | Epoxide | 13(14)-EpDPE | 0 | 0.28 (0.16, 0.48) | 0.21 (0.12, 0.36) | >0.80 | -25% (-75, 129) | >0.80 |
| NEOx | Media | DHA | Epoxide | 13(14)-EpDPE | 15 | 0.26 (0.16, 0.43) | 0.2 (0.12, 0.33) | | -24% (-71, 100) | >0.80 |
| NEOx | Media | DHA | Epoxide | 13(14)-EpDPE | 30 | 0.25 (0.15, 0.42) | 0.19 (0.12, 0.32) | | -23% (-70, 96) | >0.80 |
| NEOx | Media | DHA | Epoxide | 13(14)-EpDPE | 60 | 0.26 (0.15, 0.46) | 0.21 (0.12, 0.36) | | -21% (-77, 167) | >0.80 |
| Esterified | Cell | DHA | Epoxide | 16(17)-EpDPE | 0 | 0.7 (0.36, 1.37) | 0.95 (0.49, 1.85) | 0.72 | 35% (-67, 446) | >0.80 |
| Esterified | Cell | DHA | Epoxide | 16(17)-EpDPE | 15 | 0.88 (0.52, 1.5) | 1.12 (0.66, 1.9) | | 27% (-56, 263) | >0.80 |
| Esterified | Cell | DHA | Epoxide | 16(17)-EpDPE | 30 | 1.07 (0.61, 1.87) | 1.27 (0.73, 2.23) | | 19% (-55, 214) | >0.80 |
| Esterified | Cell | DHA | Epoxide | 16(17)-EpDPE | 60 | 1.42 (0.69, 2.92) | 1.5 (0.73, 3.09) | | 6% (-79, 437) | >0.80 |
| NEOx | Cell | DHA | Epoxide | 16(17)-EpDPE | 0 | 0.5 (0.34, 0.76) | 0.38 (0.25, 0.56) | >0.80 | -26% (-67, 69) | >0.80 |
| NEOx | Cell | DHA | Epoxide | 16(17)-EpDPE | 15 | 0.47 (0.32, 0.71) | 0.35 (0.23, 0.52) | | -26% (-65, 56) | >0.80 |
| NEOx | Cell | DHA | Epoxide | 16(17)-EpDPE | 30 | 0.46 (0.31, 0.68) | 0.34 (0.23, 0.5) | | -26% (-64, 53) | >0.80 |
| NEOx | Cell | DHA | Epoxide | 16(17)-EpDPE | 60 | 0.47 (0.31, 0.71) | 0.34 (0.23, 0.52) | | -27% (-70, 76) | >0.80 |
| Esterified | Media | DHA | Epoxide | 16(17)-EpDPE | 0 | 0.46 (0.25, 0.88) | 0.44 (0.23, 0.82) | 0.35 | -6% (-74, 240) | >0.80 |
| Esterified | Media | DHA | Epoxide | 16(17)-EpDPE | 15 | 0.46 (0.25, 0.87) | 0.47 (0.25, 0.89) | | 3% (-68, 231) | >0.80 |
| Esterified | Media | DHA | Epoxide | 16(17)-EpDPE | 30 | 0.46 (0.25, 0.86) | 0.52 (0.28, 0.97) | | 13% (-64, 255) | >0.80 |
| Esterified | Media | DHA | Epoxide | 16(17)-EpDPE | 60 | 0.47 (0.24, 0.89) | 0.64 (0.33, 1.21) | | 36% (-66, 439) | >0.80 |
| NEOx | Media | DHA | Epoxide | 16(17)-EpDPE | 0 | 0.44 (0.28, 0.71) | 0.37 (0.23, 0.58) | 0.05 | -18% (-67, 105) | >0.80 |
| NEOx | Media | DHA | Epoxide | 16(17)-EpDPE | 15 | 0.38 (0.24, 0.61) | 0.35 (0.22, 0.56) | | -8% (-61, 118) | >0.80 |
| NEOx | Media | DHA | Epoxide | 16(17)-EpDPE | 30 | 0.35 (0.22, 0.56) | 0.36 (0.22, 0.57) | | 3% (-56, 141) | >0.80 |
| NEOx | Media | DHA | Epoxide | 16(17)-EpDPE | 60 | 0.37 (0.23, 0.59) | 0.48 (0.3, 0.76) | | 28% (-50, 231) | >0.80 |
| Esterified | Cell | DHA | Epoxide | 19(20)-EpDPE | 0 | 12.1 (5.6, 26.3) | 13.2 (6.1, 28.7) | 0.38 | 9% (-78, 431) | >0.80 |
| Esterified | Cell | DHA | Epoxide | 19(20)-EpDPE | 15 | 14.8 (7, 31.4) | 14.4 (6.8, 30.4) | | -3% (-76, 295) | >0.80 |

| | | | | | | | | | | |
|---|---|---|---|---|---|---|---|---|---|---|
| Esterified | Cell | DHA | Epoxide | 19(20)-EpDPE | 30 | 17.6 (8.3, 37.3) | 15.2 (7.2, 32.1) | | -14% (-78, 237) | >0.80 |
| Esterified | Cell | DHA | Epoxide | 19(20)-EpDPE | 60 | 23.1 (10.4, 51) | 15.6 (7.1, 34.5) | | -32% (-88, 274) | >0.80 |
| NEOx | Cell | DHA | Epoxide | 19(20)-EpDPE | 0 | 0.09 (0.05, 0.16) | 0.07 (0.04, 0.12) | >0.80 | -27% (-76, 122) | >0.80 |
| NEOx | Cell | DHA | Epoxide | 19(20)-EpDPE | 15 | 0.12 (0.08, 0.19) | 0.09 (0.06, 0.14) | | -27% (-69, 72) | >0.80 |
| NEOx | Cell | DHA | Epoxide | 19(20)-EpDPE | 30 | 0.14 (0.09, 0.22) | 0.11 (0.07, 0.17) | | -26% (-66, 63) | >0.80 |
| NEOx | Cell | DHA | Epoxide | 19(20)-EpDPE | 60 | 0.13 (0.07, 0.23) | 0.1 (0.06, 0.17) | | -25% (-79, 172) | >0.80 |
| Esterified | Media | DHA | Epoxide | 19(20)-EpDPE | 0 | 0.36 (0.19, 0.68) | 0.38 (0.2, 0.71) | 0.67 | 4% (-72, 283) | >0.80 |
| Esterified | Media | DHA | Epoxide | 19(20)-EpDPE | 15 | 0.36 (0.23, 0.55) | 0.35 (0.23, 0.53) | | -3% (-59, 127) | >0.80 |
| Esterified | Media | DHA | Epoxide | 19(20)-EpDPE | 30 | 0.38 (0.23, 0.61) | 0.34 (0.21, 0.54) | | -11% (-57, 87) | >0.80 |
| Esterified | Media | DHA | Epoxide | 19(20)-EpDPE | 60 | 0.51 (0.25, 1.02) | 0.39 (0.19, 0.78) | | -23% (-84, 267) | >0.80 |
| NEOx | Media | DHA | Epoxide | 19(20)-EpDPE | 0 | 0.12 (0.06, 0.24) | 0.08 (0.04, 0.15) | 0.69 | -38% (-85, 148) | >0.80 |
| NEOx | Media | DHA | Epoxide | 19(20)-EpDPE | 15 | 0.14 (0.07, 0.27) | 0.09 (0.04, 0.17) | | -36% (-82, 127) | >0.80 |
| NEOx | Media | DHA | Epoxide | 19(20)-EpDPE | 30 | 0.14 (0.07, 0.27) | 0.09 (0.05, 0.18) | | -33% (-80, 130) | >0.80 |
| NEOx | Media | DHA | Epoxide | 19(20)-EpDPE | 60 | 0.1 (0.05, 0.2) | 0.07 (0.04, 0.15) | | -27% (-83, 224) | >0.80 |
| Esterified | Cell | LA | Diol | 9(10)-DiHOME | 0 | 45 (14.3, 141.2) | 46.2 (14.7, 145.1) | 0.52 | 3% (-91, 1021) | >0.80 |
| Esterified | Cell | LA | Diol | 9(10)-DiHOME | 15 | 24.7 (8.7, 70.2) | 29.8 (10.5, 84.5) | | 20% (-84, 787) | >0.80 |
| Esterified | Cell | LA | Diol | 9(10)-DiHOME | 30 | 18.9 (6.6, 54.2) | 26.6 (9.3, 76.4) | | 41% (-79, 851) | >0.80 |
| Esterified | Cell | LA | Diol | 9(10)-DiHOME | 60 | 29.3 (8.8, 97.4) | 56.6 (17.1, 188.1) | | 93% (-87, 2666) | >0.80 |
| NEOx | Cell | LA | Diol | 9(10)-DiHOME | 0 | 48.26 (20.95, 111.15) | 18.98 (8.24, 43.71) | 0.72 | -61% (-93, 124) | 0.58 |
| NEOx | Cell | LA | Diol | 9(10)-DiHOME | 15 | 37.62 (20.17, 70.19) | 16.03 (8.59, 29.91) | | -57% (-88, 48) | 0.31 |
| NEOx | Cell | LA | Diol | 9(10)-DiHOME | 30 | 35.49 (18.12, 69.52) | 16.39 (8.37, 32.1) | | -54% (-85, 41) | 0.30 |
| NEOx | Cell | LA | Diol | 9(10)-DiHOME | 60 | 55.91 (22.46, 139.17) | 30.32 (12.18, 75.46) | | -46% (-93, 321) | >0.80 |
| Esterified | Media | LA | Diol | 9(10)-DiHOME | 0 | 25.2 (8.87, 71.64) | 15.51 (5.46, 44.1) | >0.80 | -38% (-93, 416) | >0.80 |
| Esterified | Media | LA | Diol | 9(10)-DiHOME | 15 | 20.46 (7.34, 56.99) | 12.63 (4.53, 35.18) | | -38% (-91, 317) | >0.80 |
| Esterified | Media | LA | Diol | 9(10)-DiHOME | 30 | 19.9 (7.17, 55.23) | 12.32 (4.44, 34.19) | | -38% (-90, 299) | >0.80 |
| Esterified | Media | LA | Diol | 9(10)-DiHOME | 60 | 32.37 (11.14, 94.1) | 20.15 (6.93, 58.59) | | -38% (-94, 511) | >0.80 |
| NEOx | Media | LA | Diol | 9(10)-DiHOME | 0 | 28.75 (8.64, 95.69) | 27.23 (8.18, 90.66) | 0.77 | -5% (-92, 997) | >0.80 |
| NEOx | Media | LA | Diol | 9(10)-DiHOME | 15 | 18.5 (5.7, 60.08) | 18.59 (5.72, 60.36) | | 0% (-89, 804) | >0.80 |
| NEOx | Media | LA | Diol | 9(10)-DiHOME | 30 | 13.94 (4.31, 45.1) | 14.86 (4.59, 48.05) | | 7% (-87, 808) | >0.80 |
| NEOx | Media | LA | Diol | 9(10)-DiHOME | 60 | 12.72 (3.72, 43.48) | 15.24 (4.46, 52.09) | | 20% (-91, 1569) | >0.80 |
| Esterified | Cell | LA | Diol | 12(13)-DiHOME | 0 | 75 (27.7, 203.1) | 61.5 (22.7, 166.5) | 0.59 | -18% (-90, 561) | >0.80 |
| Esterified | Cell | LA | Diol | 12(13)-DiHOME | 15 | 74.5 (33.3, 166.5) | 69.8 (31.2, 156.1) | | -6% (-81, 357) | >0.80 |
| Esterified | Cell | LA | Diol | 12(13)-DiHOME | 30 | 68.9 (29.6, 160) | 73.9 (31.8, 171.7) | | 7% (-75, 364) | >0.80 |
| Esterified | Cell | LA | Diol | 12(13)-DiHOME | 60 | 47.6 (16.2, 139.3) | 66.8 (22.8, 195.6) | | 40% (-87, 1471) | >0.80 |
| NEOx | Cell | LA | Diol | 12(13)-DiHOME | 0 | 42.94 (16.1, 114.48) | 30.73 (11.53, 81.94) | 0.21 | -28% (-90, 437) | >0.80 |
| NEOx | Cell | LA | Diol | 12(13)-DiHOME | 15 | 39.63 (15.4, 101.98) | 35.49 (13.79, 91.34) | | -10% (-85, 429) | >0.80 |
| NEOx | Cell | LA | Diol | 12(13)-DiHOME | 30 | 38.71 (15.05, 99.54) | 43.38 (16.87, 111.56) | | 12% (-80, 528) | >0.80 |
| NEOx | Cell | LA | Diol | 12(13)-DiHOME | 60 | 43.77 (15.96, 120.08) | 76.81 (28, 210.71) | | 75% (-80, 1466) | >0.80 |
| Esterified | Media | LA | Diol | 12(13)-DiHOME | 0 | 20.88 (6.99, 62.35) | 23.62 (7.91, 70.55) | 0.31 | 13% (-89, 1014) | >0.80 |
| Esterified | Media | LA | Diol | 12(13)-DiHOME | 15 | 29.55 (10.99, 79.48) | 26.19 (9.74, 70.43) | | -11% (-87, 492) | >0.80 |
| Esterified | Media | LA | Diol | 12(13)-DiHOME | 30 | 35.85 (13.15, 97.69) | 24.88 (9.13, 67.81) | | -31% (-89, 325) | >0.80 |
| Esterified | Media | LA | Diol | 12(13)-DiHOME | 60 | 33.22 (10.52, 104.91) | 14.14 (4.48, 44.67) | | -57% (-97, 448) | >0.80 |
| NEOx | Media | LA | Diol | 12(13)-DiHOME | 0 | 12.79 (2.84, 57.69) | 34.52 (7.65, 155.63) | 0.25 | 170% (-88, 5840) | >0.80 |
| NEOx | Media | LA | Diol | 12(13)-DiHOME | 15 | 15.27 (3.56, 65.44) | 30.12 (7.03, 129.03) | | 97% (-87, 2925) | >0.80 |
| NEOx | Media | LA | Diol | 12(13)-DiHOME | 30 | 17.17 (4.01, 73.42) | 24.74 (5.78, 105.82) | | 44% (-90, 1944) | >0.80 |
| NEOx | Media | LA | Diol | 12(13)-DiHOME | 60 | 18.09 (3.85, 85.1) | 13.93 (2.96, 65.52) | | -23% (-97, 2101) | >0.80 |
| Esterified | Cell | AA | Diol | 14(15)-DiHETrE | 0 | 4.7 (2.5, 8.7) | 6.8 (3.7, 12.7) | >0.80 | 46% (-56, 385) | >0.80 |
| Esterified | Cell | AA | Diol | 14(15)-DiHETrE | 15 | 4.8 (3.5, 6.5) | 6.8 (5, 9.2) | | 41% (-20, 148) | 0.46 |
| Esterified | Cell | AA | Diol | 14(15)-DiHETrE | 30 | 5.1 (3.4, 7.7) | 6.9 (4.6, 10.4) | | 36% (-1, 85) | 0.06 |
| Esterified | Cell | AA | Diol | 14(15)-DiHETrE | 60 | 6.4 (3.2, 12.9) | 8.1 (4, 16.3) | | 27% (-73, 486) | >0.80 |
| NEOx | Cell | AA | Diol | 14(15)-DiHETrE | 0 | 3.6 (1.33, 9.71) | 3.61 (1.34, 9.73) | >0.80 | 0% (-87, 701) | >0.80 |
| NEOx | Cell | AA | Diol | 14(15)-DiHETrE | 15 | 3.97 (1.64, 9.61) | 4.14 (1.71, 10.03) | | 4% (-81, 474) | >0.80 |
| NEOx | Cell | AA | Diol | 14(15)-DiHETrE | 30 | 4.5 (1.83, 11.08) | 4.9 (1.99, 12.06) | | 9% (-78, 450) | >0.80 |
| NEOx | Cell | AA | Diol | 14(15)-DiHETrE | 60 | 6.35 (2.23, 18.11) | 7.5 (2.63, 21.4) | | 18% (-89, 1121) | >0.80 |
| Esterified | Media | AA | Diol | 14(15)-DiHETrE | 0 | 1.63 (0.61, 4.37) | 2.18 (0.81, 5.87) | >0.80 | 34% (-82, 886) | >0.80 |
| Esterified | Media | AA | Diol | 14(15)-DiHETrE | 15 | 1.75 (0.66, 4.66) | 2.34 (0.88, 6.24) | | 34% (-78, 725) | >0.80 |
| Esterified | Media | AA | Diol | 14(15)-DiHETrE | 30 | 1.97 (0.74, 5.24) | 2.64 (1, 7.01) | | 34% (-77, 694) | >0.80 |
| Esterified | Media | AA | Diol | 14(15)-DiHETrE | 60 | 2.94 (1.08, 8.04) | 3.93 (1.44, 10.74) | | 34% (-84, 1018) | >0.80 |
| NEOx | Media | AA | Diol | 14(15)-DiHETrE | 0 | 2.85 (1.51, 5.38) | 1.86 (0.98, 3.51) | 0.11 | -35% (-82, 143) | >0.80 |
| NEOx | Media | AA | Diol | 14(15)-DiHETrE | 15 | 1.87 (1.2, 2.91) | 1.65 (1.06, 2.57) | | -12% (-64, 114) | >0.80 |
| NEOx | Media | AA | Diol | 14(15)-DiHETrE | 30 | 1.51 (0.92, 2.47) | 1.8 (1.1, 2.94) | | 19% (-45, 158) | >0.80 |
| NEOx | Media | AA | Diol | 14(15)-DiHETrE | 60 | 1.82 (0.9, 3.68) | 3.98 (1.97, 8.03) | | 118% (-55, 955) | 0.67 |
| Esterified | Cell | EPA | Diol | 8(9)-DiHETE | 0 | 44.5 (12.9, 153.3) | 54.8 (15.9, 188.8) | 0.69 | 23% (-91, 1549) | >0.80 |
| Esterified | Cell | EPA | Diol | 8(9)-DiHETE | 15 | 45.8 (16, 131.2) | 50.2 (17.6, 143.7) | | 10% (-86, 751) | >0.80 |
| Esterified | Cell | EPA | Diol | 8(9)-DiHETE | 30 | 50.8 (17.2, 150.2) | 49.5 (16.8, 146.5) | | -3% (-86, 566) | >0.80 |
| Esterified | Cell | EPA | Diol | 8(9)-DiHETE | 60 | 77.8 (20.8, 291.3) | 60 (16, 224.8) | | -23% (-96, 1388) | >0.80 |
| NEOx | Cell | EPA | Diol | 8(9)-DiHETE | 0 | 31.18 (7.38, 131.72) | 24.01 (5.68, 101.46) | 0.62 | -23% (-96, 1472) | >0.80 |
| NEOx | Cell | EPA | Diol | 8(9)-DiHETE | 15 | 29.39 (9.39, 91.95) | 18.84 (6.02, 58.96) | | -36% (-93, 511) | >0.80 |
| NEOx | Cell | EPA | Diol | 8(9)-DiHETE | 30 | 33.56 (10.07, 111.86) | 17.92 (5.38, 59.72) | | -47% (-93, 323) | >0.80 |
| NEOx | Cell | EPA | Diol | 8(9)-DiHETE | 60 | 77.93 (16.37, 370.9) | 28.83 (6.06, 137.22) | | -63% (-99, 1135) | >0.80 |
| Esterified | Media | EPA | Diol | 8(9)-DiHETE | 0 | 25.45 (6.05, 107.02) | 10.96 (2.61, 46.1) | 0.01 | -57% (-98, 657) | >0.80 |
| Esterified | Media | EPA | Diol | 8(9)-DiHETE | 15 | 19.52 (4.64, 82.05) | 15.17 (3.61, 63.79) | | -22% (-94, 997) | >0.80 |
| Esterified | Media | EPA | Diol | 8(9)-DiHETE | 30 | 15.1 (3.62, 62.96) | 21.19 (5.08, 88.35) | | 40% (-90, 1790) | >0.80 |
| Esterified | Media | EPA | Diol | 8(9)-DiHETE | 60 | 9.29 (2.17, 39.71) | 42.47 (9.94, 181.5) | | 357% (-78, 9377) | 0.66 |
| NEOx | Media | EPA | Diol | 8(9)-DiHETE | 0 | 12.82 (2.85, 57.75) | 13.06 (2.9, 58.81) | 0.57 | 2% (-95, 1968) | >0.80 |
| NEOx | Media | EPA | Diol | 8(9)-DiHETE | 15 | 15.6 (3.48, 70.05) | 14.03 (3.13, 63.01) | | -10% (-94, 1337) | >0.80 |
| NEOx | Media | EPA | Diol | 8(9)-DiHETE | 30 | 18.81 (4.23, 83.71) | 14.94 (3.36, 66.51) | | -21% (-95, 1107) | >0.80 |
| NEOx | Media | EPA | Diol | 8(9)-DiHETE | 60 | 26.56 (5.79, 121.83) | 16.46 (3.59, 75.52) | | -38% (-97, 1406) | >0.80 |
| Esterified | Cell | EPA | Diol | 11(12)-DiHETE | 0 | 15.8 (6.6, 38.1) | 6.1 (2.5, 14.7) | 0.42 | -62% (-94, 143) | 0.62 |
| Esterified | Cell | EPA | Diol | 11(12)-DiHETE | 15 | 9.3 (4.3, 20.1) | 4.2 (1.9, 9.1) | | -55% (-90, 102) | 0.60 |
| Esterified | Cell | EPA | Diol | 11(12)-DiHETE | 30 | 7.4 (3.3, 16.3) | 3.9 (1.8, 8.7) | | -47% (-87, 120) | 0.77 |
| Esterified | Cell | EPA | Diol | 11(12)-DiHETE | 60 | 11.9 (4.7, 30.2) | 8.8 (3.4, 22.3) | | -26% (-91, 491) | >0.80 |
| NEOx | Cell | EPA | Diol | 11(12)-DiHETE | 0 | 8.67 (3.97, 18.94) | 3.11 (1.42, 6.79) | 0.36 | -64% (-93, 81) | 0.40 |
| NEOx | Cell | EPA | Diol | 11(12)-DiHETE | 15 | 7.91 (4.55, 13.76) | 3.47 (1.99, 6.03) | | -56% (-86, 33) | 0.23 |
| NEOx | Cell | EPA | Diol | 11(12)-DiHETE | 30 | 7.4 (4.02, 13.62) | 3.97 (2.16, 7.3) | | -46% (-80, 42) | 0.39 |

| | | | | | | | | | | |
|---|---|---|---|---|---|---|---|---|---|---|
| NEOx | Cell | EPA | Diol | 11(12)-DiHETE | 60 | 6.95 (2.94, 16.44) | 5.57 (2.36, 13.17) | | -20% (-88, 453) | >0.80 |
| Esterified | Media | EPA | Diol | 11(12)-DiHETE | 0 | 3.19 (1.24, 8.17) | 3.78 (1.47, 9.67) | 0.55 | 18% (-81, 656) | >0.80 |
| Esterified | Media | EPA | Diol | 11(12)-DiHETE | 15 | 2.82 (1.72, 4.63) | 2.8 (1.71, 4.6) | | -1% (-62, 156) | >0.80 |
| Esterified | Media | EPA | Diol | 11(12)-DiHETE | 30 | 3.18 (1.69, 6) | 2.65 (1.4, 4.99) | | -17% (-55, 56) | >0.80 |
| Esterified | Media | EPA | Diol | 11(12)-DiHETE | 60 | 8.26 (2.85, 23.95) | 4.83 (1.66, 14) | | -42% (-94, 506) | >0.80 |
| NEOx | Media | EPA | Diol | 11(12)-DiHETE | 0 | 6.8 (3.06, 15.09) | 2.42 (1.09, 5.37) | 0.05 | -64% (-93, 89) | 0.42 |
| NEOx | Media | EPA | Diol | 11(12)-DiHETE | 15 | 5.68 (2.78, 11.64) | 2.9 (1.42, 5.94) | | -49% (-87, 102) | 0.68 |
| NEOx | Media | EPA | Diol | 11(12)-DiHETE | 30 | 5 (2.42, 10.35) | 3.66 (1.77, 7.58) | | -27% (-80, 172) | >0.80 |
| NEOx | Media | EPA | Diol | 11(12)-DiHETE | 60 | 4.51 (1.95, 10.45) | 6.8 (2.94, 15.75) | | 51% (-77, 876) | >0.80 |
| Esterified | Cell | AA | Diol | 11(12)-DiHETrE | 0 | 14.3 (6.6, 31.1) | 5.8 (2.7, 12.7) | 0.63 | -59% (-91, 96) | 0.52 |
| Esterified | Cell | AA | Diol | 11(12)-DiHETrE | 15 | 15.1 (9.5, 24.1) | 6.9 (4.4, 11) | | -54% (-82, 15) | 0.13 |
| Esterified | Cell | AA | Diol | 11(12)-DiHETrE | 30 | 14.9 (8.6, 25.9) | 7.6 (4.4, 13.3) | | -49% (-75, 6) | 0.08 |
| Esterified | Cell | AA | Diol | 11(12)-DiHETrE | 60 | 11.5 (4.8, 27.5) | 7.4 (3.1, 17.7) | | -36% (-91, 346) | >0.80 |
| NEOx | Cell | AA | Diol | 11(12)-DiHETrE | 0 | 9.9 (3.78, 25.91) | 15.29 (5.84, 40.01) | 0.01 | 54% (-78, 969) | >0.80 |
| NEOx | Cell | AA | Diol | 11(12)-DiHETrE | 15 | 8.58 (3.3, 22.33) | 8.71 (3.35, 22.66) | | 1% (-83, 496) | >0.80 |
| NEOx | Cell | AA | Diol | 11(12)-DiHETrE | 30 | 9.9 (3.82, 25.63) | 6.6 (2.55, 17.1) | | -33% (-88, 278) | >0.80 |
| NEOx | Cell | AA | Diol | 11(12)-DiHETrE | 60 | 31.09 (11.71, 82.51) | 8.96 (3.38, 23.78) | | -71% (-96, 125) | 0.45 |
| Esterified | Media | AA | Diol | 11(12)-DiHETrE | 0 | 5.69 (2.3, 14.09) | 4.7 (1.9, 11.65) | 0.7 | -17% (-87, 443) | >0.80 |
| Esterified | Media | AA | Diol | 11(12)-DiHETrE | 15 | 6.32 (2.7, 14.81) | 5.59 (2.39, 13.1) | | -12% (-82, 345) | >0.80 |
| Esterified | Media | AA | Diol | 11(12)-DiHETrE | 30 | 6.82 (2.9, 16.03) | 6.46 (2.74, 15.18) | | -5% (-80, 349) | >0.80 |
| Esterified | Media | AA | Diol | 11(12)-DiHETrE | 60 | 7.23 (2.81, 18.56) | 7.84 (3.05, 20.14) | | 9% (-86, 763) | >0.80 |
| NEOx | Media | AA | Diol | 11(12)-DiHETrE | 0 | 9.46 (4.55, 19.65) | 5.83 (2.81, 12.11) | 0.13 | -38% (-86, 175) | >0.80 |
| NEOx | Media | AA | Diol | 11(12)-DiHETrE | 15 | 5.93 (3.69, 9.52) | 5.09 (3.17, 8.18) | | -14% (-67, 123) | >0.80 |
| NEOx | Media | AA | Diol | 11(12)-DiHETrE | 30 | 4.69 (2.72, 8.07) | 5.61 (3.26, 9.67) | | 20% (-46, 167) | >0.80 |
| NEOx | Media | AA | Diol | 11(12)-DiHETrE | 60 | 5.88 (2.61, 13.26) | 13.68 (6.07, 30.85) | | 133% (-62, 1331) | 0.73 |
| Esterified | Cell | EPA | Epoxide | 11(12)-EpETE | 0 | 0.3 (0.17, 0.55) | 0.4 (0.22, 0.74) | 0.24 | 34% (-61, 364) | >0.80 |
| Esterified | Cell | EPA | Epoxide | 11(12)-EpETE | 15 | 0.29 (0.16, 0.5) | 0.33 (0.19, 0.59) | | 17% (-60, 239) | >0.80 |
| Esterified | Cell | EPA | Epoxide | 11(12)-EpETE | 30 | 0.29 (0.17, 0.51) | 0.3 (0.17, 0.52) | | 1% (-64, 183) | >0.80 |
| Esterified | Cell | EPA | Epoxide | 11(12)-EpETE | 60 | 0.37 (0.2, 0.7) | 0.29 (0.15, 0.53) | | -24% (-80, 199) | >0.80 |
| NEOx | Cell | EPA | Epoxide | 11(12)-EpETE | 0 | 0.39 (0.24, 0.64) | 0.44 (0.27, 0.71) | 0.42 | 11% (-60, 206) | >0.80 |
| NEOx | Cell | EPA | Epoxide | 11(12)-EpETE | 15 | 0.33 (0.23, 0.5) | 0.34 (0.23, 0.5) | | 1% (-54, 119) | >0.80 |
| NEOx | Cell | EPA | Epoxide | 11(12)-EpETE | 30 | 0.31 (0.21, 0.47) | 0.28 (0.19, 0.43) | | -9% (-55, 87) | >0.80 |
| NEOx | Cell | EPA | Epoxide | 11(12)-EpETE | 60 | 0.35 (0.21, 0.59) | 0.27 (0.16, 0.45) | | -25% (-77, 142) | >0.80 |
| Esterified | Media | EPA | Epoxide | 11(12)-EpETE | 0 | 0.32 (0.2, 0.52) | 0.23 (0.14, 0.37) | >0.80 | -29% (-74, 99) | >0.80 |
| Esterified | Media | EPA | Epoxide | 11(12)-EpETE | 15 | 0.5 (0.32, 0.77) | 0.35 (0.23, 0.54) | | -29% (-69, 63) | >0.80 |
| Esterified | Media | EPA | Epoxide | 11(12)-EpETE | 30 | 0.59 (0.38, 0.91) | 0.41 (0.26, 0.64) | | -30% (-68, 55) | 0.76 |
| Esterified | Media | EPA | Epoxide | 11(12)-EpETE | 60 | 0.35 (0.21, 0.59) | 0.24 (0.15, 0.41) | | -31% (-78, 118) | >0.80 |
| NEOx | Media | EPA | Epoxide | 11(12)-EpETE | 0 | 0.34 (0.2, 0.57) | 0.28 (0.16, 0.47) | 0.21 | 23% (-58, 257) | >0.80 |
| NEOx | Media | EPA | Epoxide | 11(12)-EpETE | 15 | 0.35 (0.21, 0.58) | 0.32 (0.19, 0.53) | | 9% (-58, 183) | >0.80 |
| NEOx | Media | EPA | Epoxide | 11(12)-EpETE | 30 | 0.34 (0.2, 0.56) | 0.35 (0.21, 0.58) | | -3% (-61, 146) | >0.80 |
| NEOx | Media | EPA | Epoxide | 11(12)-EpETE | 60 | 0.26 (0.15, 0.44) | 0.33 (0.19, 0.57) | | -23% (-76, 145) | >0.80 |
| Esterified | Cell | AA | Epoxide | 11(12)-EpETrE | 0 | 0.19 (0.09, 0.41) | 0.21 (0.1, 0.46) | >0.80 | 11% (-77, 446) | >0.80 |
| Esterified | Cell | AA | Epoxide | 11(12)-EpETrE | 15 | 0.14 (0.07, 0.25) | 0.15 (0.08, 0.28) | | 13% (-66, 277) | >0.80 |
| Esterified | Cell | AA | Epoxide | 11(12)-EpETrE | 30 | 0.12 (0.06, 0.23) | 0.14 (0.07, 0.26) | | 14% (-62, 248) | >0.80 |
| Esterified | Cell | AA | Epoxide | 11(12)-EpETrE | 60 | 0.17 (0.08, 0.39) | 0.2 (0.09, 0.45) | | 17% (-81, 637) | >0.80 |
| NEOx | Cell | AA | Epoxide | 11(12)-EpETrE | 0 | 0.09 (0.04, 0.18) | 0.07 (0.04, 0.15) | 0.42 | -19% (-82, 261) | >0.80 |
| NEOx | Cell | AA | Epoxide | 11(12)-EpETrE | 15 | 0.09 (0.04, 0.17) | 0.08 (0.04, 0.16) | | -9% (-75, 231) | >0.80 |
| NEOx | Cell | AA | Epoxide | 11(12)-EpETrE | 30 | 0.09 (0.05, 0.18) | 0.09 (0.05, 0.18) | | 2% (-71, 254) | >0.80 |
| NEOx | Cell | AA | Epoxide | 11(12)-EpETrE | 60 | 0.11 (0.05, 0.23) | 0.14 (0.07, 0.29) | | 27% (-75, 552) | >0.80 |
| Esterified | Media | AA | Epoxide | 11(12)-EpETrE | 0 | 0.11 (0.05, 0.22) | 0.12 (0.06, 0.25) | 0.69 | 12% (-75, 402) | >0.80 |
| Esterified | Media | AA | Epoxide | 11(12)-EpETrE | 15 | 0.1 (0.06, 0.17) | 0.1 (0.06, 0.17) | | 4% (-64, 202) | >0.80 |
| Esterified | Media | AA | Epoxide | 11(12)-EpETrE | 30 | 0.1 (0.06, 0.18) | 0.09 (0.05, 0.17) | | -4% (-63, 150) | >0.80 |
| Esterified | Media | AA | Epoxide | 11(12)-EpETrE | 60 | 0.15 (0.07, 0.32) | 0.12 (0.06, 0.27) | | -18% (-86, 380) | >0.80 |
| NEOx | Media | AA | Epoxide | 11(12)-EpETrE | 0 | 0.07 (0.04, 0.14) | 0.12 (0.06, 0.22) | 0.1 | 57% (-56, 466) | >0.80 |
| NEOx | Media | AA | Epoxide | 11(12)-EpETrE | 15 | 0.11 (0.07, 0.18) | 0.13 (0.09, 0.2) | | 16% (-51, 173) | >0.80 |
| NEOx | Media | AA | Epoxide | 11(12)-EpETrE | 30 | 0.15 (0.09, 0.24) | 0.13 (0.08, 0.21) | | -14% (-59, 80) | >0.80 |
| NEOx | Media | AA | Epoxide | 11(12)-EpETrE | 60 | 0.15 (0.08, 0.31) | 0.07 (0.04, 0.14) | | -53% (-90, 117) | 0.67 |
| Esterified | Cell | DHA | Alcohol | 11-HDoHE | 0 | 1.69 (0.76, 3.76) | 2.21 (0.99, 4.93) | 0.26 | 31% (-75, 601) | >0.80 |
| Esterified | Cell | DHA | Alcohol | 11-HDoHE | 15 | 2.13 (1.14, 3.97) | 2.2 (1.18, 4.09) | | 3% (-70, 255) | >0.80 |
| Esterified | Cell | DHA | Alcohol | 11-HDoHE | 30 | 2.77 (1.43, 5.36) | 2.25 (1.16, 4.36) | | -19% (-74, 151) | >0.80 |
| Esterified | Cell | DHA | Alcohol | 11-HDoHE | 60 | 5.11 (2.14, 12.21) | 2.57 (1.08, 6.15) | | -50% (-93, 257) | >0.80 |
| NEOx | Cell | DHA | Alcohol | 11-HDoHE | 0 | 0.24 (0.13, 0.42) | 0.17 (0.1, 0.31) | 0.79 | -27% (-78, 147) | >0.80 |
| NEOx | Cell | DHA | Alcohol | 11-HDoHE | 15 | 0.27 (0.17, 0.44) | 0.21 (0.13, 0.34) | | -24% (-70, 96) | >0.80 |
| NEOx | Cell | DHA | Alcohol | 11-HDoHE | 30 | 0.3 (0.18, 0.5) | 0.24 (0.14, 0.39) | | -21% (-67, 91) | >0.80 |
| NEOx | Cell | DHA | Alcohol | 11-HDoHE | 60 | 0.32 (0.17, 0.59) | 0.27 (0.15, 0.51) | | -15% (-79, 245) | >0.80 |
| Esterified | Media | DHA | Alcohol | 11-HDoHE | 0 | 0.3 (0.18, 0.49) | 0.3 (0.18, 0.51) | 0.66 | 3% (-64, 199) | >0.80 |
| Esterified | Media | DHA | Alcohol | 11-HDoHE | 15 | 0.21 (0.14, 0.32) | 0.2 (0.13, 0.32) | | -2% (-58, 128) | >0.80 |
| Esterified | Media | DHA | Alcohol | 11-HDoHE | 30 | 0.2 (0.13, 0.31) | 0.18 (0.12, 0.29) | | -7% (-58, 106) | >0.80 |
| Esterified | Media | DHA | Alcohol | 11-HDoHE | 60 | 0.41 (0.24, 0.7) | 0.34 (0.2, 0.59) | | -16% (-75, 180) | >0.80 |
| NEOx | Media | DHA | Alcohol | 11-HDoHE | 0 | 0.34 (0.21, 0.55) | 0.29 (0.18, 0.47) | 0.69 | -15% (-68, 125) | >0.80 |
| NEOx | Media | DHA | Alcohol | 11-HDoHE | 15 | 0.27 (0.17, 0.42) | 0.22 (0.14, 0.35) | | -18% (-65, 96) | >0.80 |
| NEOx | Media | DHA | Alcohol | 11-HDoHE | 30 | 0.23 (0.15, 0.37) | 0.19 (0.12, 0.29) | | -20% (-66, 85) | >0.80 |
| NEOx | Media | DHA | Alcohol | 11-HDoHE | 60 | 0.25 (0.16, 0.41) | 0.19 (0.12, 0.31) | | -25% (-74, 113) | >0.80 |
| Esterified | Cell | EPA | Diol | 14(15)-DiHETE | 0 | 1.5 (0.4, 6) | 4 (1, 15.9) | 0.1 | 163% (-85, 4565) | >0.80 |
| Esterified | Cell | EPA | Diol | 14(15)-DiHETE | 15 | 5.3 (1.9, 14.8) | 7.4 (2.7, 20.6) | | 39% (-82, 978) | >0.80 |
| Esterified | Cell | EPA | Diol | 14(15)-DiHETE | 30 | 12.4 (4.1, 37.4) | 9.1 (3, 27.7) | | -26% (-88, 363) | >0.80 |
| Esterified | Cell | EPA | Diol | 14(15)-DiHETE | 60 | 19.5 (4.3, 88.4) | 4.1 (0.9, 18.4) | | -79% (-99, 522) | 0.74 |
| NEOx | Cell | EPA | Diol | 14(15)-DiHETE | 0 | 7.56 (3.37, 16.95) | 5.43 (2.42, 12.18) | >0.80 | -28% (-86, 261) | >0.80 |
| NEOx | Cell | EPA | Diol | 14(15)-DiHETE | 15 | 7.26 (4.6, 11.46) | 4.93 (3.13, 7.78) | | -32% (-72, 67) | 0.8 |
| NEOx | Cell | EPA | Diol | 14(15)-DiHETE | 30 | 8.87 (5.06, 15.56) | 5.69 (3.24, 9.98) | | -36% (-67, 26) | 0.35 |
| NEOx | Cell | EPA | Diol | 14(15)-DiHETE | 60 | 27.18 (10.94, 67.48) | 15.57 (6.27, 38.66) | | -43% (-92, 328) | >0.80 |
| Esterified | Media | EPA | Diol | 14(15)-DiHETE | 0 | 1.33 (0.37, 4.81) | 2.69 (0.74, 9.71) | >0.80 | 102% (-84, 2469) | >0.80 |
| Esterified | Media | EPA | Diol | 14(15)-DiHETE | 15 | 2.75 (1.38, 5.49) | 5.06 (2.53, 10.11) | | 84% (-52, 606) | 0.75 |
| Esterified | Media | EPA | Diol | 14(15)-DiHETE | 30 | 4.64 (1.93, 11.13) | 7.8 (3.25, 18.71) | | 68% (-34, 328) | 0.55 |
| Esterified | Media | EPA | Diol | 14(15)-DiHETE | 60 | 7.26 (1.7, 30.96) | 10.15 (2.38, 43.32) | | 40% (-94, 3311) | >0.80 |

| | | | | | | | | | | |
|---|---|---|---|---|---|---|---|---|---|---|
| NEOx | Media | EPA | Diol | 14(15)-DiHETE | 0 | 7.33 (2.62, 20.51) | 7.04 (2.52, 19.7) | 0.35 | -4% (-89, 729) | >0.80 |
| NEOx | Media | EPA | Diol | 14(15)-DiHETE | 15 | 3.99 (1.62, 9.84) | 4.78 (1.94, 11.79) | | 20% (-79, 586) | >0.80 |
| NEOx | Media | EPA | Diol | 14(15)-DiHETE | 30 | 2.99 (1.19, 7.51) | 4.47 (1.78, 11.24) | | 50% (-71, 679) | >0.80 |
| NEOx | Media | EPA | Diol | 14(15)-DiHETE | 60 | 4.37 (1.47, 13.01) | 10.19 (3.42, 30.3) | | 133% (-80, 2560) | >0.80 |
| Esterified | Cell | AA | Prostanoid | 15-deoxy-PGJ2 | 0 | 26 (13, 52) | 32 (16, 64) | 0.56 | 24% (-71, 436) | >0.80 |
| Esterified | Cell | AA | Prostanoid | 15-deoxy-PGJ2 | 15 | 39 (23, 67) | 43 (25, 75) | | 11% (-62, 225) | >0.80 |
| Esterified | Cell | AA | Prostanoid | 15-deoxy-PGJ2 | 30 | 54 (30, 96) | 54 (30, 96) | | 0% (-63, 165) | >0.80 |
| Esterified | Cell | AA | Prostanoid | 15-deoxy-PGJ2 | 60 | 78 (36, 167) | 62 (29, 134) | | -20% (-86, 345) | >0.80 |
| NEOx | Cell | AA | Prostanoid | 15-deoxy-PGJ2 | 0 | 28.74 (12.44, 66.38) | 33.29 (14.41, 76.89) | 0.55 | 16% (-80, 571) | >0.80 |
| NEOx | Cell | AA | Prostanoid | 15-deoxy-PGJ2 | 15 | 22.03 (10.87, 44.64) | 28.8 (14.21, 58.36) | | 31% (-67, 419) | >0.80 |
| NEOx | Cell | AA | Prostanoid | 15-deoxy-PGJ2 | 30 | 18.07 (8.71, 37.49) | 26.66 (12.85, 55.31) | | 48% (-59, 436) | >0.80 |
| NEOx | Cell | AA | Prostanoid | 15-deoxy-PGJ2 | 60 | 14.9 (6.09, 36.47) | 28.01 (11.44, 68.55) | | 88% (-75, 1299) | >0.80 |
| Esterified | Media | AA | Prostanoid | 15-deoxy-PGJ2 | 0 | 50.61 (23.3, 109.91) | 41.49 (19.1, 90.11) | 0.45 | -18% (-84, 315) | >0.80 |
| Esterified | Media | AA | Prostanoid | 15-deoxy-PGJ2 | 15 | 60.86 (33.41, 110.84) | 42.78 (23.49, 77.92) | | -30% (-79, 131) | >0.80 |
| Esterified | Media | AA | Prostanoid | 15-deoxy-PGJ2 | 30 | 69.18 (36.54, 130.95) | 41.69 (22.03, 78.93) | | -40% (-80, 78) | 0.73 |
| Esterified | Media | AA | Prostanoid | 15-deoxy-PGJ2 | 60 | 75.48 (32.49, 175.39) | 33.45 (14.39, 77.71) | | -56% (-93, 195) | 0.80 |
| NEOx | Media | AA | Prostanoid | 15-deoxy-PGJ2 | 0 | 31.7 (12.41, 80.95) | 22.36 (8.75, 57.09) | 0.26 | -29% (-90, 400) | >0.80 |
| NEOx | Media | AA | Prostanoid | 15-deoxy-PGJ2 | 15 | 45.2 (21.9, 93.27) | 42.22 (20.46, 87.11) | | -7% (-78, 293) | >0.80 |
| NEOx | Media | AA | Prostanoid | 15-deoxy-PGJ2 | 30 | 45.66 (21.12, 98.73) | 56.48 (26.12, 122.13) | | 24% (-67, 358) | >0.80 |
| NEOx | Media | AA | Prostanoid | 15-deoxy-PGJ2 | 60 | 16.57 (5.98, 45.91) | 35.94 (12.97, 99.62) | | 117% (-78, 2048) | >0.80 |
| Esterified | Cell | EPA | Alcohol | 15-HEPE | 0 | 0.7 (0.35, 1.41) | 0.51 (0.26, 1.03) | 0.5 | -26% (-83, 214) | >0.80 |
| Esterified | Cell | EPA | Alcohol | 15-HEPE | 15 | 0.78 (0.47, 1.29) | 0.5 (0.3, 0.83) | | -35% (-76, 77) | 0.79 |
| Esterified | Cell | EPA | Alcohol | 15-HEPE | 30 | 0.79 (0.46, 1.37) | 0.45 (0.26, 0.78) | | -43% (-77, 39) | 0.40 |
| Esterified | Cell | EPA | Alcohol | 15-HEPE | 60 | 0.61 (0.28, 1.32) | 0.27 (0.12, 0.58) | | -56% (-92, 147) | 0.71 |
| NEOx | Cell | EPA | Alcohol | 15-HEPE | 0 | 0.73 (0.31, 1.73) | 0.45 (0.19, 1.06) | 0.4 | -39% (-90, 270) | >0.80 |
| NEOx | Cell | EPA | Alcohol | 15-HEPE | 15 | 0.71 (0.31, 1.59) | 0.5 (0.22, 1.13) | | -29% (-85, 230) | >0.80 |
| NEOx | Cell | EPA | Alcohol | 15-HEPE | 30 | 0.72 (0.32, 1.63) | 0.59 (0.26, 1.33) | | -18% (-81, 260) | >0.80 |
| NEOx | Cell | EPA | Alcohol | 15-HEPE | 60 | 0.86 (0.35, 2.12) | 0.95 (0.38, 2.32) | | 9% (-85, 691) | >0.80 |
| Esterified | Media | EPA | Alcohol | 15-HEPE | 0 | 0.45 (0.23, 0.87) | 0.53 (0.27, 1.04) | 0.29 | 19% (-71, 385) | >0.80 |
| Esterified | Media | EPA | Alcohol | 15-HEPE | 15 | 0.43 (0.25, 0.74) | 0.43 (0.25, 0.74) | | 0% (-66, 195) | >0.80 |
| Esterified | Media | EPA | Alcohol | 15-HEPE | 30 | 0.46 (0.26, 0.81) | 0.38 (0.22, 0.68) | | -16% (-69, 130) | >0.80 |
| Esterified | Media | EPA | Alcohol | 15-HEPE | 60 | 0.73 (0.35, 1.49) | 0.43 (0.21, 0.88) | | -41% (-88, 196) | >0.80 |
| NEOx | Media | EPA | Alcohol | 15-HEPE | 0 | 1.07 (0.55, 2.07) | 0.63 (0.33, 1.22) | 0.22 | -41% (-85, 134) | >0.80 |
| NEOx | Media | EPA | Alcohol | 15-HEPE | 15 | 0.68 (0.4, 1.15) | 0.49 (0.29, 0.84) | | -27% (-74, 105) | >0.80 |
| NEOx | Media | EPA | Alcohol | 15-HEPE | 30 | 0.5 (0.29, 0.87) | 0.45 (0.26, 0.78) | | -10% (-65, 133) | >0.80 |
| NEOx | Media | EPA | Alcohol | 15-HEPE | 60 | 0.43 (0.21, 0.88) | 0.59 (0.29, 1.2) | | 37% (-72, 576) | >0.80 |
| Esterified | Cell | DHA | Alcohol | 16-HDoHE | 0 | 1.1 (0.35, 3.43) | 0.77 (0.25, 2.41) | 0.8 | -30% (-93, 576) | >0.80 |
| Esterified | Cell | DHA | Alcohol | 16-HDoHE | 15 | 1.51 (0.8, 2.86) | 1.16 (0.61, 2.19) | | -23% (-78, 169) | >0.80 |
| Esterified | Cell | DHA | Alcohol | 16-HDoHE | 30 | 1.74 (0.79, 3.83) | 1.47 (0.67, 3.22) | | -16% (-67, 113) | >0.80 |
| Esterified | Cell | DHA | Alcohol | 16-HDoHE | 60 | 1.38 (0.38, 4.95) | 1.39 (0.39, 4.97) | | 1% (-94, 1594) | >0.80 |
| NEOx | Cell | DHA | Alcohol | 16-HDoHE | 0 | 0.12 (0.06, 0.25) | 0.05 (0.03, 0.11) | >0.80 | -55% (-90, 94) | 0.56 |
| NEOx | Cell | DHA | Alcohol | 16-HDoHE | 15 | 0.13 (0.09, 0.2) | 0.06 (0.04, 0.09) | | -56% (-81, 3) | 0.06 |
| NEOx | Cell | DHA | Alcohol | 16-HDoHE | 30 | 0.14 (0.08, 0.24) | 0.06 (0.04, 0.1) | | -58% (-78, -16) | 0.009 |
| NEOx | Cell | DHA | Alcohol | 16-HDoHE | 60 | 0.14 (0.06, 0.31) | 0.06 (0.02, 0.12) | | -60% (-93, 143) | 0.64 |
| Esterified | Media | DHA | Alcohol | 16-HDoHE | 0 | 0.11 (0.04, 0.26) | 0.06 (0.02, 0.14) | 0.2 | -46% (-91, 235) | >0.80 |
| Esterified | Media | DHA | Alcohol | 16-HDoHE | 15 | 0.08 (0.04, 0.15) | 0.06 (0.03, 0.11) | | -26% (-80, 172) | >0.80 |
| Esterified | Media | DHA | Alcohol | 16-HDoHE | 30 | 0.06 (0.03, 0.12) | 0.06 (0.03, 0.12) | | 0% (-69, 226) | >0.80 |
| Esterified | Media | DHA | Alcohol | 16-HDoHE | 60 | 0.05 (0.02, 0.12) | 0.09 (0.03, 0.22) | | 87% (-78, 1507) | >0.80 |
| NEOx | Media | DHA | Alcohol | 16-HDoHE | 0 | 0.07 (0.03, 0.15) | 0.05 (0.02, 0.11) | 0.3 | -29% (-85, 244) | >0.80 |
| NEOx | Media | DHA | Alcohol | 16-HDoHE | 15 | 0.06 (0.03, 0.13) | 0.04 (0.02, 0.08) | | -39% (-84, 129) | >0.80 |
| NEOx | Media | DHA | Alcohol | 16-HDoHE | 30 | 0.07 (0.03, 0.13) | 0.03 (0.02, 0.07) | | -48% (-86, 85) | 0.62 |
| NEOx | Media | DHA | Alcohol | 16-HDoHE | 60 | 0.11 (0.05, 0.23) | 0.04 (0.02, 0.09) | | -63% (-93, 112) | 0.52 |
| Esterified | Cell | EPA | Diol | 17(18)-DiHETE | 0 | 152 (63, 369) | 197 (81, 477) | >0.80 | 29% (-79, 680) | >0.80 |
| Esterified | Cell | EPA | Diol | 17(18)-DiHETE | 15 | 70 (40, 121) | 87 (50, 151) | | 25% (-58, 277) | >0.80 |
| Esterified | Cell | EPA | Diol | 17(18)-DiHETE | 30 | 51 (27, 97) | 62 (32, 118) | | 21% (-51, 198) | >0.80 |
| Esterified | Cell | EPA | Diol | 17(18)-DiHETE | 60 | 112 (42, 301) | 127 (47, 341) | | 13% (-87, 923) | >0.80 |
| NEOx | Cell | EPA | Diol | 17(18)-DiHETE | 0 | 74.75 (27.51, 203.09) | 40.22 (14.8, 109.26) | 0.62 | -46% (-93, 338) | >0.80 |
| NEOx | Cell | EPA | Diol | 17(18)-DiHETE | 15 | 97.79 (42.87, 223.09) | 46.57 (20.41, 106.24) | | -52% (-91, 140) | 0.74 |
| NEOx | Cell | EPA | Diol | 17(18)-DiHETE | 30 | 119.32 (50.58, 281.44) | 50.3 (21.32, 118.65) | | -58% (-91, 90) | 0.51 |
| NEOx | Cell | EPA | Diol | 17(18)-DiHETE | 60 | 144.11 (49.25, 421.67) | 47.61 (16.27, 139.3) | | -67% (-97, 268) | 0.74 |
| Esterified | Media | EPA | Diol | 17(18)-DiHETE | 0 | 29.46 (10.05, 86.41) | 11.89 (4.05, 34.88) | 0.29 | -60% (-96, 285) | >0.80 |
| Esterified | Media | EPA | Diol | 17(18)-DiHETE | 15 | 42.65 (17.61, 103.28) | 22.93 (9.47, 55.54) | | -46% (-91, 205) | >0.80 |
| Esterified | Media | EPA | Diol | 17(18)-DiHETE | 30 | 49.19 (19.57, 123.61) | 35.24 (14.02, 88.56) | | -28% (-86, 259) | >0.80 |
| Esterified | Media | EPA | Diol | 17(18)-DiHETE | 60 | 33.12 (10.42, 105.31) | 42.12 (13.25, 133.93) | | 27% (-91, 1609) | >0.80 |
| NEOx | Media | EPA | Diol | 17(18)-DiHETE | 0 | 55.5 (24.83, 124.05) | 84.09 (37.63, 187.95) | 0.74 | 52% (-71, 703) | >0.80 |
| NEOx | Media | EPA | Diol | 17(18)-DiHETE | 15 | 27.46 (15.44, 48.84) | 38.66 (21.73, 68.77) | | 41% (-56, 346) | >0.80 |
| NEOx | Media | EPA | Diol | 17(18)-DiHETE | 30 | 19.56 (10.4, 36.79) | 25.59 (13.61, 48.14) | | 31% (-53, 263) | >0.80 |
| NEOx | Media | EPA | Diol | 17(18)-DiHETE | 60 | 29.65 (12.24, 71.78) | 33.5 (13.84, 81.1) | | 13% (-84, 723) | >0.80 |
| Esterified | Cell | DHA | Alcohol | 20-HDoHE | 0 | 4.94 (2.19, 11.12) | 3.96 (1.76, 8.93) | >0.80 | -20% (-85, 319) | >0.80 |
| Esterified | Cell | DHA | Alcohol | 20-HDoHE | 15 | 6.14 (2.77, 13.61) | 5.06 (2.28, 11.23) | | -18% (-81, 264) | >0.80 |
| Esterified | Cell | DHA | Alcohol | 20-HDoHE | 30 | 7.08 (3.2, 15.65) | 6 (2.71, 13.26) | | -15% (-80, 261) | >0.80 |
| Esterified | Cell | DHA | Alcohol | 20-HDoHE | 60 | 7.5 (3.27, 17.19) | 6.71 (2.93, 15.37) | | -11% (-85, 428) | >0.80 |
| NEOx | Cell | DHA | Alcohol | 20-HDoHE | 0 | 0.16 (0.03, 0.75) | 0.04 (0.01, 0.19) | 0.77 | -75% (-99, 522) | 0.79 |
| NEOx | Cell | DHA | Alcohol | 20-HDoHE | 15 | 0.11 (0.04, 0.31) | 0.02 (0.01, 0.07) | | -78% (-97, 82) | 0.26 |
| NEOx | Cell | DHA | Alcohol | 20-HDoHE | 30 | 0.09 (0.03, 0.31) | 0.02 (0.01, 0.06) | | -81% (-97, 19) | 0.09 |
| NEOx | Cell | DHA | Alcohol | 20-HDoHE | 60 | 0.15 (0.03, 0.84) | 0.02 (0, 0.12) | | -85% (-100, 610) | 0.67 |
| Esterified | Media | DHA | Alcohol | 20-HDoHE | 0 | 0.14 (0.03, 0.6) | 0.1 (0.02, 0.43) | 0.24 | -28% (-97, 1404) | >0.80 |
| Esterified | Media | DHA | Alcohol | 20-HDoHE | 15 | 0.08 (0.02, 0.26) | 0.08 (0.02, 0.28) | | 8% (-90, 1124) | >0.80 |
| Esterified | Media | DHA | Alcohol | 20-HDoHE | 30 | 0.05 (0.01, 0.18) | 0.08 (0.02, 0.3) | | 63% (-83, 1499) | >0.80 |
| Esterified | Media | DHA | Alcohol | 20-HDoHE | 60 | 0.04 (0.01, 0.21) | 0.17 (0.04, 0.78) | | 271% (-88, 11640) | >0.80 |
| NEOx | Media | DHA | Alcohol | 20-HDoHE | 0 | 0.04 (0.01, 0.16) | 0.06 (0.01, 0.23) | 0.44 | 45% (-92, 2452) | >0.80 |
| NEOx | Media | DHA | Alcohol | 20-HDoHE | 15 | 0.07 (0.02, 0.21) | 0.08 (0.02, 0.24) | | 12% (-88, 933) | >0.80 |
| NEOx | Media | DHA | Alcohol | 20-HDoHE | 30 | 0.08 (0.02, 0.24) | 0.06 (0.02, 0.21) | | -14% (-89, 584) | >0.80 |
| NEOx | Media | DHA | Alcohol | 20-HDoHE | 60 | 0.02 (0.01, 0.11) | 0.01 (0, 0.05) | | -49% (-98, 1288) | >0.80 |
| Esterified | Cell | AA | Alcohol | 20-HETE | 0 | 0.37 (0.21, 0.64) | 0.34 (0.2, 0.6) | 0.3 | -6% (-71, 199) | >0.80 |

| | | | | | | | | | | |
|---|---|---|---|---|---|---|---|---|---|---|
| Esterified | Cell | AA | Alcohol | 20-HETE | 15 | 0.44 (0.28, 0.71) | 0.36 (0.22, 0.58) | | -19% (-67, 104) | >0.80 |
| Esterified | Cell | AA | Alcohol | 20-HETE | 30 | 0.51 (0.32, 0.84) | 0.36 (0.22, 0.59) | | -29% (-70, 68) | >0.80 |
| Esterified | Cell | AA | Alcohol | 20-HETE | 60 | 0.65 (0.36, 1.17) | 0.35 (0.19, 0.63) | | -46% (-86, 102) | 0.72 |
| NEOx | Cell | AA | Alcohol | 20-HETE | 0 | 0.33 (0.2, 0.53) | 0.27 (0.16, 0.43) | 0.61 | -18% (-70, 122) | >0.80 |
| NEOx | Cell | AA | Alcohol | 20-HETE | 15 | 0.38 (0.24, 0.6) | 0.3 (0.19, 0.47) | | -22% (-68, 86) | >0.80 |
| NEOx | Cell | AA | Alcohol | 20-HETE | 30 | 0.41 (0.26, 0.65) | 0.3 (0.19, 0.48) | | -26% (-68, 73) | >0.80 |
| NEOx | Cell | AA | Alcohol | 20-HETE | 60 | 0.39 (0.23, 0.64) | 0.26 (0.16, 0.43) | | -32% (-77, 102) | >0.80 |
| Esterified | Media | AA | Alcohol | 20-HETE | 0 | 0.36 (0.19, 0.67) | 0.32 (0.17, 0.61) | >0.80 | -10% (-75, 231) | >0.80 |
| Esterified | Media | AA | Alcohol | 20-HETE | 15 | 0.41 (0.22, 0.73) | 0.37 (0.2, 0.67) | | -9% (-70, 180) | >0.80 |
| Esterified | Media | AA | Alcohol | 20-HETE | 30 | 0.43 (0.24, 0.78) | 0.39 (0.22, 0.71) | | -8% (-69, 172) | >0.80 |
| Esterified | Media | AA | Alcohol | 20-HETE | 60 | 0.39 (0.2, 0.75) | 0.36 (0.19, 0.7) | | -7% (-78, 286) | >0.80 |
| NEOx | Media | AA | Alcohol | 20-HETE | 0 | 0.38 (0.18, 0.81) | 0.28 (0.13, 0.6) | 0.26 | -25% (-82, 211) | >0.80 |
| NEOx | Media | AA | Alcohol | 20-HETE | 15 | 0.32 (0.15, 0.68) | 0.25 (0.12, 0.55) | | -20% (-80, 220) | >0.80 |
| NEOx | Media | AA | Alcohol | 20-HETE | 30 | 0.3 (0.14, 0.64) | 0.26 (0.12, 0.55) | | -14% (-78, 241) | >0.80 |
| NEOx | Media | AA | Alcohol | 20-HETE | 60 | 0.37 (0.17, 0.79) | 0.37 (0.17, 0.78) | | -1% (-77, 327) | >0.80 |
| Esterified | Cell | DHA | Alcohol | 22-HDoHE | 0 | 0.34 (0.12, 0.97) | 0.78 (0.28, 2.23) | 0.18 | 130% (-74, 1921) | >0.80 |
| Esterified | Cell | DHA | Alcohol | 22-HDoHE | 15 | 0.59 (0.22, 1.55) | 1.01 (0.38, 2.64) | | 71% (-73, 975) | >0.80 |
| Esterified | Cell | DHA | Alcohol | 22-HDoHE | 30 | 0.99 (0.37, 2.62) | 1.25 (0.47, 3.32) | | 27% (-78, 641) | >0.80 |
| Esterified | Cell | DHA | Alcohol | 22-HDoHE | 60 | 2.58 (0.87, 7.67) | 1.8 (0.61, 5.37) | | -30% (-94, 678) | >0.80 |
| NEOx | Cell | DHA | Alcohol | 22-HDoHE | 0 | 0.15 (0.07, 0.33) | 0.1 (0.04, 0.21) | 0.6 | -36% (-88, 237) | >0.80 |
| NEOx | Cell | DHA | Alcohol | 22-HDoHE | 15 | 0.16 (0.08, 0.31) | 0.11 (0.05, 0.22) | | -29% (-82, 175) | >0.80 |
| NEOx | Cell | DHA | Alcohol | 22-HDoHE | 30 | 0.17 (0.09, 0.36) | 0.14 (0.07, 0.28) | | -22% (-79, 182) | >0.80 |
| NEOx | Cell | DHA | Alcohol | 22-HDoHE | 60 | 0.28 (0.12, 0.64) | 0.26 (0.11, 0.6) | | -7% (-85, 496) | >0.80 |
| Esterified | Media | DHA | Alcohol | 22-HDoHE | 0 | 0.14 (0.05, 0.38) | 0.14 (0.05, 0.38) | 0.8 | -1% (-88, 733) | >0.80 |
| Esterified | Media | DHA | Alcohol | 22-HDoHE | 15 | 0.21 (0.08, 0.53) | 0.19 (0.08, 0.5) | | -6% (-85, 469) | >0.80 |
| Esterified | Media | DHA | Alcohol | 22-HDoHE | 30 | 0.28 (0.11, 0.72) | 0.25 (0.09, 0.64) | | -11% (-84, 401) | >0.80 |
| Esterified | Media | DHA | Alcohol | 22-HDoHE | 60 | 0.34 (0.12, 1) | 0.27 (0.09, 0.8) | | -20% (-92, 746) | >0.80 |
| NEOx | Media | DHA | Alcohol | 22-HDoHE | 0 | 0.14 (0.06, 0.32) | 0.08 (0.04, 0.18) | >0.80 | -44% (-89, 195) | >0.80 |
| NEOx | Media | DHA | Alcohol | 22-HDoHE | 15 | 0.17 (0.08, 0.34) | 0.1 (0.05, 0.2) | | -43% (-85, 125) | >0.80 |
| NEOx | Media | DHA | Alcohol | 22-HDoHE | 30 | 0.19 (0.09, 0.39) | 0.11 (0.05, 0.23) | | -41% (-84, 115) | >0.80 |
| NEOx | Media | DHA | Alcohol | 22-HDoHE | 60 | 0.22 (0.1, 0.51) | 0.14 (0.06, 0.31) | | -38% (-90, 296) | >0.80 |
| Esterified | Cell | AA | Prostanoid | 6-keto-PGF1a | 0 | 80 (28, 227) | 24 (8, 67) | 0.22 | -71% (-97, 161) | 0.54 |
| Esterified | Cell | AA | Prostanoid | 6-keto-PGF1a | 15 | 47 (20, 110) | 19 (8, 45) | | -59% (-92, 118) | 0.59 |
| Esterified | Cell | AA | Prostanoid | 6-keto-PGF1a | 30 | 33 (14, 81) | 19 (8, 46) | | -44% (-88, 167) | >0.80 |
| Esterified | Cell | AA | Prostanoid | 6-keto-PGF1a | 60 | 30 (10, 93) | 33 (11, 100) | | 7% (-91, 1226) | >0.80 |
| NEOx | Cell | AA | Prostanoid | 6-keto-PGF1a | 0 | 22.26 (5.08, 97.54) | 9.42 (2.15, 41.27) | 0.51 | -58% (-98, 832) | >0.80 |
| NEOx | Cell | AA | Prostanoid | 6-keto-PGF1a | 15 | 13.46 (3.61, 50.25) | 7.05 (1.89, 26.32) | | -48% (-96, 563) | >0.80 |
| NEOx | Cell | AA | Prostanoid | 6-keto-PGF1a | 30 | 10.94 (2.86, 41.75) | 7.09 (1.86, 27.07) | | -35% (-94, 623) | >0.80 |
| NEOx | Cell | AA | Prostanoid | 6-keto-PGF1a | 60 | 17.5 (3.68, 83.23) | 17.4 (3.66, 82.73) | | -1% (-97, 3105) | >0.80 |
| Esterified | Media | AA | Prostanoid | 6-keto-PGF1a | 0 | 17.19 (7.67, 38.51) | 32.1 (14.32, 71.92) | 0.74 | 87% (-66, 915) | >0.80 |
| Esterified | Media | AA | Prostanoid | 6-keto-PGF1a | 15 | 18.62 (9.33, 37.15) | 32.65 (16.36, 65.14) | | 75% (-54, 573) | >0.80 |
| Esterified | Media | AA | Prostanoid | 6-keto-PGF1a | 30 | 20.71 (10.18, 42.14) | 34.09 (16.75, 69.38) | | 65% (-53, 482) | >0.80 |
| Esterified | Media | AA | Prostanoid | 6-keto-PGF1a | 60 | 27.72 (11.73, 65.49) | 40.22 (17.02, 95.04) | | 45% (-79, 896) | >0.80 |
| NEOx | Media | AA | Prostanoid | 6-keto-PGF1a | 0 | 44.55 (17.65, 112.48) | 15.81 (6.26, 39.9) | 0.25 | -65% (-95, 148) | 0.59 |
| NEOx | Media | AA | Prostanoid | 6-keto-PGF1a | 15 | 40.21 (18.27, 88.48) | 18.5 (8.41, 40.7) | | -54% (-90, 114) | 0.65 |
| NEOx | Media | AA | Prostanoid | 6-keto-PGF1a | 30 | 32.37 (14.36, 72.95) | 19.31 (8.57, 43.51) | | -40% (-86, 152) | >0.80 |
| NEOx | Media | AA | Prostanoid | 6-keto-PGF1a | 60 | 14.88 (5.54, 39.97) | 14.92 (5.55, 40.08) | | 0% (-89, 818) | >0.80 |
| Esterified | Cell | DHA | Epoxide | 7(8)-EpDPE | 0 | 1.6 (0.74, 3.48) | 1.73 (0.8, 3.76) | >0.80 | 8% (-78, 437) | >0.80 |
| Esterified | Cell | DHA | Epoxide | 7(8)-EpDPE | 15 | 1.31 (0.63, 2.73) | 1.42 (0.68, 2.96) | | 8% (-73, 334) | >0.80 |
| Esterified | Cell | DHA | Epoxide | 7(8)-EpDPE | 30 | 1.27 (0.61, 2.65) | 1.39 (0.67, 2.89) | | 9% (-71, 315) | >0.80 |
| Esterified | Cell | DHA | Epoxide | 7(8)-EpDPE | 60 | 2.01 (0.9, 4.48) | 2.21 (0.99, 4.92) | | 10% (-81, 537) | >0.80 |
| NEOx | Cell | DHA | Epoxide | 7(8)-EpDPE | 0 | 0.3 (0.21, 0.44) | 0.18 (0.12, 0.26) | 0.24 | -41% (-73, 30) | 0.33 |
| NEOx | Cell | DHA | Epoxide | 7(8)-EpDPE | 15 | 0.27 (0.2, 0.37) | 0.18 (0.13, 0.24) | | -35% (-65, 22) | 0.31 |
| NEOx | Cell | DHA | Epoxide | 7(8)-EpDPE | 30 | 0.25 (0.18, 0.35) | 0.18 (0.13, 0.25) | | -27% (-60, 30) | 0.57 |
| NEOx | Cell | DHA | Epoxide | 7(8)-EpDPE | 60 | 0.24 (0.16, 0.36) | 0.21 (0.14, 0.32) | | -10% (-64, 122) | >0.80 |
| Esterified | Media | DHA | Epoxide | 7(8)-EpDPE | 0 | 0.28 (0.15, 0.53) | 0.26 (0.13, 0.49) | >0.80 | -8% (-75, 238) | >0.80 |
| Esterified | Media | DHA | Epoxide | 7(8)-EpDPE | 15 | 0.22 (0.12, 0.42) | 0.2 (0.11, 0.38) | | -10% (-72, 193) | >0.80 |
| Esterified | Media | DHA | Epoxide | 7(8)-EpDPE | 30 | 0.22 (0.12, 0.42) | 0.2 (0.1, 0.37) | | -12% (-72, 179) | >0.80 |
| Esterified | Media | DHA | Epoxide | 7(8)-EpDPE | 60 | 0.38 (0.2, 0.74) | 0.32 (0.17, 0.62) | | -16% (-79, 237) | >0.80 |
| NEOx | Media | DHA | Epoxide | 7(8)-EpDPE | 0 | 0.27 (0.19, 0.37) | 0.22 (0.16, 0.31) | 0.46 | -16% (-59, 70) | >0.80 |
| NEOx | Media | DHA | Epoxide | 7(8)-EpDPE | 15 | 0.24 (0.18, 0.33) | 0.22 (0.16, 0.29) | | -11% (-50, 57) | >0.80 |
| NEOx | Media | DHA | Epoxide | 7(8)-EpDPE | 30 | 0.23 (0.17, 0.31) | 0.22 (0.16, 0.29) | | -6% (-45, 61) | >0.80 |
| NEOx | Media | DHA | Epoxide | 7(8)-EpDPE | 60 | 0.23 (0.16, 0.33) | 0.25 (0.17, 0.35) | | 5% (-53, 136) | >0.80 |
| Esterified | Cell | AA | Diol | 8(9)-DiHETrE | 0 | 18.9 (7.4, 47.9) | 33.7 (13.3, 85.5) | 0.18 | 79% (-74, 1147) | >0.80 |
| Esterified | Cell | AA | Diol | 8(9)-DiHETrE | 15 | 23.6 (11.5, 48.3) | 30 (14.6, 61.4) | | 27% (-69, 427) | >0.80 |
| Esterified | Cell | AA | Diol | 8(9)-DiHETrE | 30 | 28.8 (13.4, 61.9) | 26.1 (12.1, 56) | | -10% (-75, 230) | >0.80 |
| Esterified | Cell | AA | Diol | 8(9)-DiHETrE | 60 | 40.3 (14.7, 111) | 18.5 (6.7, 50.8) | | -54% (-95, 346) | >0.80 |
| NEOx | Cell | AA | Diol | 8(9)-DiHETrE | 0 | 26.41 (10.42, 66.89) | 13.13 (5.18, 33.27) | 0.2 | -50% (-93, 249) | >0.80 |
| NEOx | Cell | AA | Diol | 8(9)-DiHETrE | 15 | 31.2 (14.2, 68.55) | 20.75 (9.44, 45.59) | | -33% (-86, 209) | >0.80 |
| NEOx | Cell | AA | Diol | 8(9)-DiHETrE | 30 | 34.05 (15.11, 76.74) | 30.28 (13.44, 68.24) | | -11% (-79, 275) | >0.80 |
| NEOx | Cell | AA | Diol | 8(9)-DiHETrE | 60 | 32.02 (11.86, 86.43) | 50.91 (18.86, 137.42) | | 59% (-83, 1373) | >0.80 |
| Esterified | Media | AA | Diol | 8(9)-DiHETrE | 0 | 26.62 (11.38, 62.29) | 23.24 (9.93, 54.39) | 0.77 | -13% (-85, 420) | >0.80 |
| Esterified | Media | AA | Diol | 8(9)-DiHETrE | 15 | 23.59 (11.47, 48.49) | 19.43 (9.45, 39.95) | | -18% (-80, 236) | >0.80 |
| Esterified | Media | AA | Diol | 8(9)-DiHETrE | 30 | 22.92 (10.9, 48.2) | 17.82 (8.47, 37.47) | | -22% (-79, 190) | >0.80 |
| Esterified | Media | AA | Diol | 8(9)-DiHETrE | 60 | 28.51 (11.5, 70.7) | 19.73 (7.96, 48.94) | | -31% (-91, 430) | >0.80 |
| NEOx | Media | AA | Diol | 8(9)-DiHETrE | 0 | 21.43 (4.95, 92.75) | 18.81 (4.35, 81.43) | >0.80 | -12% (-96, 1627) | >0.80 |
| NEOx | Media | AA | Diol | 8(9)-DiHETrE | 15 | 13.36 (3.17, 56.31) | 11.57 (2.74, 48.77) | | -13% (-94, 1163) | >0.80 |
| NEOx | Media | AA | Diol | 8(9)-DiHETrE | 30 | 10.96 (2.61, 45.95) | 9.36 (2.23, 39.26) | | -15% (-94, 1069) | >0.80 |
| NEOx | Media | AA | Diol | 8(9)-DiHETrE | 60 | 16.82 (3.77, 75.07) | 13.99 (3.13, 62.42) | | -17% (-97, 1936) | >0.80 |
| Esterified | Cell | AA | Epoxide | 8(9)-EpETrE | 0 | 0.34 (0.19, 0.63) | 0.35 (0.19, 0.64) | 0.64 | 3% (-71, 268) | >0.80 |
| Esterified | Cell | AA | Epoxide | 8(9)-EpETrE | 15 | 0.44 (0.26, 0.73) | 0.42 (0.25, 0.7) | | -4% (-64, 159) | >0.80 |
| Esterified | Cell | AA | Epoxide | 8(9)-EpETrE | 30 | 0.54 (0.32, 0.91) | 0.48 (0.28, 0.81) | | -10% (-65, 126) | >0.80 |
| Esterified | Cell | AA | Epoxide | 8(9)-EpETrE | 60 | 0.68 (0.36, 1.31) | 0.53 (0.28, 1.02) | | -22% (-82, 236) | >0.80 |
| NEOx | Cell | AA | Epoxide | 8(9)-EpETrE | 0 | 0.28 (0.14, 0.53) | 0.28 (0.14, 0.53) | 0.68 | 0% (-74, 291) | >0.80 |
| NEOx | Cell | AA | Epoxide | 8(9)-EpETrE | 15 | 0.24 (0.14, 0.43) | 0.26 (0.14, 0.46) | | 7% (-65, 225) | >0.80 |

| | | | | | | | | | | |
|---|---|---|---|---|---|---|---|---|---|---|
| NEOx | Cell | AA | Epoxide | 8(9)-EpETrE | 30 | 0.22 (0.12, 0.39) | 0.25 (0.14, 0.44) | | 13% (-61, 226) | >0.80 |
| NEOx | Cell | AA | Epoxide | 8(9)-EpETrE | 60 | 0.18 (0.09, 0.36) | 0.23 (0.12, 0.46) | | 28% (-72, 488) | >0.80 |
| Esterified | Media | AA | Epoxide | 8(9)-EpETrE | 0 | 0.19 (0.09, 0.37) | 0.19 (0.1, 0.38) | 0.45 | 2% (-76, 330) | >0.80 |
| Esterified | Media | AA | Epoxide | 8(9)-EpETrE | 15 | 0.27 (0.15, 0.5) | 0.24 (0.13, 0.45) | | -10% (-72, 192) | >0.80 |
| Esterified | Media | AA | Epoxide | 8(9)-EpETrE | 30 | 0.33 (0.18, 0.62) | 0.27 (0.14, 0.5) | | -20% (-74, 143) | >0.80 |
| Esterified | Media | AA | Epoxide | 8(9)-EpETrE | 60 | 0.31 (0.15, 0.65) | 0.2 (0.1, 0.41) | | -37% (-88, 222) | >0.80 |
| NEOx | Media | AA | Epoxide | 8(9)-EpETrE | 0 | 0.26 (0.12, 0.58) | 0.27 (0.12, 0.6) | 0.71 | 3% (-80, 420) | >0.80 |
| NEOx | Media | AA | Epoxide | 8(9)-EpETrE | 15 | 0.27 (0.13, 0.57) | 0.29 (0.14, 0.62) | | 8% (-74, 346) | >0.80 |
| NEOx | Media | AA | Epoxide | 8(9)-EpETrE | 30 | 0.27 (0.13, 0.56) | 0.3 (0.14, 0.64) | | 14% (-71, 350) | >0.80 |
| NEOx | Media | AA | Epoxide | 8(9)-EpETrE | 60 | 0.24 (0.11, 0.54) | 0.3 (0.13, 0.68) | | 28% (-78, 651) | >0.80 |
| Esterified | Cell | EPA | Alcohol | 9-HEPE | 0 | 0.84 (0.52, 1.37) | 0.55 (0.34, 0.9) | 0.59 | -34% (-76, 83) | >0.80 |
| Esterified | Cell | EPA | Alcohol | 9-HEPE | 15 | 0.93 (0.63, 1.37) | 0.57 (0.39, 0.84) | | -39% (-71, 32) | 0.39 |
| Esterified | Cell | EPA | Alcohol | 9-HEPE | 30 | 0.97 (0.65, 1.46) | 0.56 (0.37, 0.84) | | -43% (-72, 16) | 0.18 |
| Esterified | Cell | EPA | Alcohol | 9-HEPE | 60 | 0.91 (0.54, 1.55) | 0.46 (0.27, 0.77) | | -50% (-85, 65) | 0.49 |
| NEOx | Cell | EPA | Alcohol | 9-HEPE | 0 | 0.63 (0.28, 1.43) | 0.44 (0.19, 1) | 0.59 | -30% (-87, 288) | >0.80 |
| NEOx | Cell | EPA | Alcohol | 9-HEPE | 15 | 0.67 (0.34, 1.29) | 0.42 (0.21, 0.81) | | -37% (-83, 130) | >0.80 |
| NEOx | Cell | EPA | Alcohol | 9-HEPE | 30 | 0.74 (0.37, 1.49) | 0.42 (0.21, 0.83) | | -44% (-83, 87) | 0.70 |
| NEOx | Cell | EPA | Alcohol | 9-HEPE | 60 | 1.08 (0.45, 2.6) | 0.48 (0.2, 1.17) | | -55% (-94, 226) | >0.80 |
| Esterified | Media | EPA | Alcohol | 9-HEPE | 0 | 0.4 (0.19, 0.84) | 0.53 (0.25, 1.1) | 0.56 | 30% (-71, 478) | >0.80 |
| Esterified | Media | EPA | Alcohol | 9-HEPE | 15 | 0.38 (0.19, 0.78) | 0.53 (0.26, 1.09) | | 39% (-64, 433) | >0.80 |
| Esterified | Media | EPA | Alcohol | 9-HEPE | 30 | 0.38 (0.18, 0.77) | 0.56 (0.27, 1.15) | | 49% (-60, 454) | >0.80 |
| Esterified | Media | EPA | Alcohol | 9-HEPE | 60 | 0.42 (0.2, 0.9) | 0.73 (0.34, 1.54) | | 72% (-65, 754) | >0.80 |
| NEOx | Media | EPA | Alcohol | 9-HEPE | 0 | 0.55 (0.35, 0.86) | 0.51 (0.33, 0.8) | 0.61 | -7% (-64, 139) | >0.80 |
| NEOx | Media | EPA | Alcohol | 9-HEPE | 15 | 0.36 (0.25, 0.53) | 0.36 (0.24, 0.52) | | -1% (-53, 106) | >0.80 |
| NEOx | Media | EPA | Alcohol | 9-HEPE | 30 | 0.3 (0.2, 0.44) | 0.31 (0.21, 0.46) | | 4% (-48, 107) | >0.80 |
| NEOx | Media | EPA | Alcohol | 9-HEPE | 60 | 0.38 (0.23, 0.61) | 0.44 (0.27, 0.71) | | 17% (-60, 242) | >0.80 |
| Esterified | Cell | DHA | Diol | Maresin 1 | 0 | 88 (44, 177) | 29 (14, 58) | 0.65 | -67% (-92, 35) | 0.18 |
| Esterified | Cell | DHA | Diol | Maresin 1 | 15 | 91 (59, 142) | 33 (21, 51) | | -64% (-85, -13) | 0.02 |
| Esterified | Cell | DHA | Diol | Maresin 1 | 30 | 96 (58, 161) | 38 (23, 63) | | -61% (-81, -18) | 0.008 |
| Esterified | Cell | DHA | Diol | Maresin 1 | 60 | 114 (52, 247) | 54 (25, 116) | | -53% (-92, 166) | 0.79 |
| NEOx | Cell | DHA | Diol | Maresin 1 | 0 | 24.19 (6.64, 88.12) | 23.42 (6.43, 85.29) | 0.7 | -3% (-94, 1355) | >0.80 |
| NEOx | Cell | DHA | Diol | Maresin 1 | 15 | 24.6 (8.54, 70.88) | 26.92 (9.34, 77.57) | | 9% (-86, 775) | >0.80 |
| NEOx | Cell | DHA | Diol | Maresin 1 | 30 | 24.7 (8.19, 74.5) | 30.56 (10.13, 92.16) | | 24% (-82, 751) | >0.80 |
| NEOx | Cell | DHA | Diol | Maresin 1 | 60 | 23.98 (5.97, 96.34) | 37.91 (9.44, 152.31) | | 58% (-93, 3496) | >0.80 |
| Esterified | Media | DHA | Diol | Maresin 1 | 0 | 31.63 (11.61, 86.21) | 25.82 (9.48, 70.37) | 0.74 | -18% (-89, 530) | >0.80 |
| Esterified | Media | DHA | Diol | Maresin 1 | 15 | 59.34 (31.43, 112.03) | 43.84 (23.22, 82.78) | | -26% (-79, 165) | >0.80 |
| Esterified | Media | DHA | Diol | Maresin 1 | 30 | 79.3 (37.95, 165.71) | 53.03 (25.38, 110.83) | | -33% (-77, 93) | >0.80 |
| Esterified | Media | DHA | Diol | Maresin 1 | 60 | 51.22 (16.75, 156.63) | 28.07 (9.18, 85.82) | | -45% (-95, 563) | >0.80 |
| NEOx | Media | DHA | Diol | Maresin 1 | 0 | 44.19 (15.63, 124.95) | 40.08 (14.18, 113.34) | 0.79 | -9% (-89, 668) | >0.80 |
| NEOx | Media | DHA | Diol | Maresin 1 | 15 | 60.13 (22.08, 163.76) | 57.32 (21.05, 156.1) | | -5% (-85, 526) | >0.80 |
| NEOx | Media | DHA | Diol | Maresin 1 | 30 | 61.97 (22.77, 168.64) | 62.08 (22.81, 168.93) | | 0% (-84, 522) | >0.80 |
| NEOx | Media | DHA | Diol | Maresin 1 | 60 | 28.59 (9.81, 83.3) | 31.62 (10.85, 92.15) | | 11% (-89, 1026) | >0.80 |
| Esterified | Cell | AA | Prostanoid | PGD2 | 0 | 3.5 (1.8, 6.8) | 4.4 (2.2, 8.5) | 0.55 | 24% (-68, 389) | >0.80 |
| Esterified | Cell | AA | Prostanoid | PGD2 | 15 | 4 (2.5, 6.3) | 4.4 (2.8, 6.9) | | 11% (-55, 175) | >0.80 |
| Esterified | Cell | AA | Prostanoid | PGD2 | 30 | 4.4 (2.6, 7.3) | 4.3 (2.6, 7.2) | | -1% (-55, 117) | >0.80 |
| Esterified | Cell | AA | Prostanoid | PGD2 | 60 | 5.1 (2.4, 10.5) | 4 (1.9, 8.3) | | -22% (-85, 308) | >0.80 |
| NEOx | Cell | AA | Prostanoid | PGD2 | 0 | 3.24 (0.85, 12.39) | 2.09 (0.55, 7.99) | 0.11 | -36% (-95, 714) | >0.80 |
| NEOx | Cell | AA | Prostanoid | PGD2 | 15 | 3.08 (0.79, 11.97) | 2.41 (0.62, 9.39) | | -22% (-93, 813) | >0.80 |
| NEOx | Cell | AA | Prostanoid | PGD2 | 30 | 2.72 (0.7, 10.51) | 2.59 (0.67, 10.02) | | -5% (-92, 992) | >0.80 |
| NEOx | Cell | AA | Prostanoid | PGD2 | 60 | 1.71 (0.45, 6.51) | 2.41 (0.63, 9.19) | | 41% (-90, 1799) | >0.80 |
| Esterified | Media | AA | Prostanoid | PGD2 | 0 | 2.9 (1.26, 6.68) | 3.64 (1.58, 8.39) | 0.51 | 25% (-78, 616) | >0.80 |
| Esterified | Media | AA | Prostanoid | PGD2 | 15 | 3.21 (1.69, 6.1) | 3.48 (1.83, 6.61) | | 8% (-70, 289) | >0.80 |
| Esterified | Media | AA | Prostanoid | PGD2 | 30 | 3.47 (1.75, 6.89) | 3.26 (1.64, 6.46) | | -6% (-71, 199) | >0.80 |
| Esterified | Media | AA | Prostanoid | PGD2 | 60 | 3.86 (1.56, 9.56) | 2.71 (1.09, 6.7) | | -30% (-91, 439) | >0.80 |
| NEOx | Media | AA | Prostanoid | PGD2 | 0 | 2.99 (1.39, 6.47) | 2.08 (0.96, 4.5) | 0.23 | -30% (-86, 249) | >0.80 |
| NEOx | Media | AA | Prostanoid | PGD2 | 15 | 2.51 (1.37, 4.6) | 2.23 (1.22, 4.1) | | -11% (-73, 196) | >0.80 |
| NEOx | Media | AA | Prostanoid | PGD2 | 30 | 2.17 (1.14, 4.13) | 2.48 (1.3, 4.7) | | 14% (-62, 242) | >0.80 |
| NEOx | Media | AA | Prostanoid | PGD2 | 60 | 1.81 (0.78, 4.17) | 3.37 (1.46, 7.78) | | 87% (-71, 1122) | >0.80 |
| Esterified | Cell | EPA | Prostanoid | PGD3 | 0 | 13.4 (5.1, 35.1) | 24.2 (9.2, 63.4) | 0.57 | 80% (-76, 1264) | >0.80 |
| Esterified | Cell | EPA | Prostanoid | PGD3 | 15 | 9.5 (4.1, 21.9) | 15 (6.5, 34.8) | | 59% (-69, 708) | >0.80 |
| Esterified | Cell | EPA | Prostanoid | PGD3 | 30 | 8.7 (3.7, 20.5) | 12.2 (5.2, 28.7) | | 40% (-70, 550) | >0.80 |
| Esterified | Cell | EPA | Prostanoid | PGD3 | 60 | 16.2 (5.8, 45.1) | 17.6 (6.3, 48.9) | | 9% (-89, 972) | >0.80 |
| NEOx | Cell | EPA | Prostanoid | PGD3 | 0 | 21.96 (5.76, 83.64) | 8.1 (2.13, 30.86) | 0.23 | -63% (-98, 493) | >0.80 |
| NEOx | Cell | EPA | Prostanoid | PGD3 | 15 | 15.16 (4.34, 52.96) | 7.74 (2.21, 27.04) | | -49% (-95, 450) | >0.80 |
| NEOx | Cell | EPA | Prostanoid | PGD3 | 30 | 11.61 (3.3, 40.83) | 8.2 (2.33, 28.85) | | -29% (-93, 596) | >0.80 |
| NEOx | Cell | EPA | Prostanoid | PGD3 | 60 | 9.31 (2.32, 37.42) | 12.6 (3.13, 50.63) | | 35% (-94, 2787) | >0.80 |
| Esterified | Media | EPA | Prostanoid | PGD3 | 0 | 6.39 (2.33, 17.55) | 10.29 (3.75, 28.25) | 0.41 | 61% (-81, 1234) | >0.80 |
| Esterified | Media | EPA | Prostanoid | PGD3 | 15 | 9.99 (4.48, 22.28) | 12.95 (5.8, 28.88) | | 30% (-73, 532) | >0.80 |
| Esterified | Media | EPA | Prostanoid | PGD3 | 30 | 13.95 (5.99, 32.51) | 14.56 (6.25, 33.93) | | 4% (-76, 348) | >0.80 |
| Esterified | Media | EPA | Prostanoid | PGD3 | 60 | 19.46 (6.52, 58.07) | 13.17 (4.42, 39.31) | | -32% (-94, 691) | >0.80 |
| NEOx | Media | EPA | Prostanoid | PGD3 | 0 | 15.42 (4.5, 52.86) | 15.67 (4.57, 53.72) | 0.63 | 2% (-92, 1234) | >0.80 |
| NEOx | Media | EPA | Prostanoid | PGD3 | 15 | 15.14 (4.98, 46.03) | 13.55 (4.46, 41.2) | | -10% (-89, 657) | >0.80 |
| NEOx | Media | EPA | Prostanoid | PGD3 | 30 | 13.13 (4.25, 40.55) | 10.36 (3.35, 31.97) | | -21% (-90, 504) | >0.80 |
| NEOx | Media | EPA | Prostanoid | PGD3 | 60 | 6.83 (1.87, 24.96) | 4.18 (1.14, 15.27) | | -39% (-97, 988) | >0.80 |
| Esterified | Cell | dgLA | Prostanoid | PGE1 | 0 | 1.66 (0.59, 4.68) | 1.46 (0.52, 4.11) | 0.6 | -12% (-90, 672) | >0.80 |
| Esterified | Cell | dgLA | Prostanoid | PGE1 | 15 | 1.26 (0.54, 2.96) | 0.97 (0.41, 2.27) | | -23% (-86, 308) | >0.80 |
| Esterified | Cell | dgLA | Prostanoid | PGE1 | 30 | 1.03 (0.42, 2.5) | 0.69 (0.28, 1.68) | | -33% (-86, 216) | >0.80 |
| Esterified | Cell | dgLA | Prostanoid | PGE1 | 60 | 0.85 (0.28, 2.6) | 0.43 (0.14, 1.33) | | -49% (-96, 524) | >0.80 |
| NEOx | Cell | dgLA | Prostanoid | PGE1 | 0 | 0.87 (0.28, 2.67) | 0.37 (0.12, 1.14) | 0.25 | -57% (-96, 351) | >0.80 |
| NEOx | Cell | dgLA | Prostanoid | PGE1 | 15 | 0.97 (0.37, 2.54) | 0.57 (0.22, 1.48) | | -42% (-91, 276) | >0.80 |
| NEOx | Cell | dgLA | Prostanoid | PGE1 | 30 | 0.99 (0.37, 2.66) | 0.79 (0.29, 2.12) | | -20% (-86, 358) | >0.80 |
| NEOx | Cell | dgLA | Prostanoid | PGE1 | 60 | 0.76 (0.23, 2.53) | 1.13 (0.34, 3.76) | | 49% (-90, 2089) | >0.80 |
| NEOx | Media | dgLA | Prostanoid | PGE1 | 0 | 2.82 (1.01, 7.87) | 1.54 (0.55, 4.28) | 0.59 | -46% (-94, 364) | >0.80 |
| NEOx | Media | dgLA | Prostanoid | PGE1 | 15 | 1.73 (0.78, 3.82) | 1.08 (0.49, 2.4) | | -37% (-87, 203) | >0.80 |
| NEOx | Media | dgLA | Prostanoid | PGE1 | 30 | 1.32 (0.57, 3.07) | 0.96 (0.41, 2.22) | | -28% (-83, 204) | >0.80 |

| | | | | | | | | | | |
|---|---|---|---|---|---|---|---|---|---|---|
| NEOx | Media | dgLA | Prostanoid | PGE1 | 60 | 1.5 (0.49, 4.56) | 1.44 (0.47, 4.4) | | -4% (-92, 1083) | >0.80 |
| Esterified | Cell | AA | Prostanoid | PGE2 | 0 | 14 (6, 32.4) | 9.8 (4.2, 22.6) | 0.5 | -30% (-88, 305) | >0.80 |
| Esterified | Cell | AA | Prostanoid | PGE2 | 15 | 15.4 (8.1, 29.4) | 9.2 (4.9, 17.6) | | -40% (-83, 116) | >0.80 |
| Esterified | Cell | AA | Prostanoid | PGE2 | 30 | 16.9 (8.5, 33.7) | 8.7 (4.4, 17.4) | | -48% (-84, 65) | 0.52 |
| Esterified | Cell | AA | Prostanoid | PGE2 | 60 | 20.1 (8.1, 50.4) | 7.7 (3.1, 19.2) | | -62% (-95, 200) | 0.72 |
| NEOx | Cell | AA | Prostanoid | PGE2 | 0 | 2.53 (0.62, 10.42) | 5.48 (1.33, 22.53) | 0.33 | 116% (-89, 4093) | >0.80 |
| NEOx | Cell | AA | Prostanoid | PGE2 | 15 | 5.05 (1.59, 16.11) | 7.74 (2.43, 24.66) | | 53% (-84, 1392) | >0.80 |
| NEOx | Cell | AA | Prostanoid | PGE2 | 30 | 8.74 (2.61, 29.28) | 9.47 (2.83, 31.74) | | 8% (-87, 796) | >0.80 |
| NEOx | Cell | AA | Prostanoid | PGE2 | 60 | 17.07 (3.73, 78.12) | 9.28 (2.03, 42.46) | | -46% (-98, 1557) | >0.80 |
| Esterified | Media | AA | Prostanoid | PGE2 | 0 | 9.12 (2.47, 33.67) | 4.02 (1.09, 14.83) | >0.80 | -56% (-97, 582) | >0.80 |
| Esterified | Media | AA | Prostanoid | PGE2 | 15 | 9.38 (3.14, 28.03) | 4.18 (1.4, 12.48) | | -55% (-95, 278) | >0.80 |
| Esterified | Media | AA | Prostanoid | PGE2 | 30 | 9.47 (3.05, 29.4) | 4.26 (1.37, 13.23) | | -55% (-94, 232) | >0.80 |
| Esterified | Media | AA | Prostanoid | PGE2 | 60 | 9.1 (2.25, 36.86) | 4.18 (1.03, 16.94) | | -54% (-98, 960) | >0.80 |
| NEOx | Media | AA | Prostanoid | PGE2 | 0 | 14.14 (3.78, 52.91) | 13.84 (3.7, 51.77) | 0.62 | -2% (-94, 1455) | >0.80 |
| NEOx | Media | AA | Prostanoid | PGE2 | 15 | 9.1 (2.87, 28.8) | 10.35 (3.27, 32.77) | | 14% (-88, 960) | >0.80 |
| NEOx | Media | AA | Prostanoid | PGE2 | 30 | 6.81 (2.09, 22.12) | 9 (2.77, 29.26) | | 32% (-84, 990) | >0.80 |
| NEOx | Media | AA | Prostanoid | PGE2 | 60 | 5.99 (1.48, 24.3) | 10.71 (2.64, 43.44) | | 79% (-92, 3982) | >0.80 |
| Esterified | Cell | EPA | Prostanoid | PGE3 | 0 | 28.7 (11.1, 74.1) | 25.8 (10, 66.5) | >0.80 | -10% (-88, 548) | >0.80 |
| Esterified | Cell | EPA | Prostanoid | PGE3 | 15 | 20.7 (10.2, 41.9) | 17.9 (8.8, 36.2) | | -14% (-79, 253) | >0.80 |
| Esterified | Cell | EPA | Prostanoid | PGE3 | 30 | 17 (7.9, 36.5) | 14.1 (6.6, 30.3) | | -17% (-76, 194) | >0.80 |
| Esterified | Cell | EPA | Prostanoid | PGE3 | 60 | 17.1 (6.1, 48.2) | 13.2 (4.7, 37.2) | | -23% (-93, 694) | >0.80 |
| NEOx | Cell | EPA | Prostanoid | PGE3 | 0 | 17.07 (3.97, 73.47) | 14.91 (3.46, 64.2) | >0.80 | -13% (-96, 1737) | >0.80 |
| NEOx | Cell | EPA | Prostanoid | PGE3 | 15 | 7.48 (1.97, 28.41) | 6.96 (1.83, 26.43) | | -7% (-93, 1095) | >0.80 |
| NEOx | Cell | EPA | Prostanoid | PGE3 | 30 | 5.57 (1.45, 21.47) | 5.52 (1.43, 21.26) | | -1% (-91, 1040) | >0.80 |
| NEOx | Cell | EPA | Prostanoid | PGE3 | 60 | 15.23 (3.3, 70.27) | 17.09 (3.7, 78.86) | | 12% (-96, 3223) | >0.80 |
| Esterified | Media | EPA | Prostanoid | PGE3 | 0 | 9.91 (3.41, 28.79) | 13 (4.48, 37.76) | 0.66 | 31% (-86, 1098) | >0.80 |
| Esterified | Media | EPA | Prostanoid | PGE3 | 15 | 11.65 (5.41, 25.09) | 13.4 (6.22, 28.87) | | 15% (-75, 434) | >0.80 |
| Esterified | Media | EPA | Prostanoid | PGE3 | 30 | 12.53 (5.41, 29.02) | 12.64 (5.46, 29.28) | | 1% (-74, 293) | >0.80 |
| Esterified | Media | EPA | Prostanoid | PGE3 | 60 | 11.09 (3.44, 35.77) | 8.6 (2.67, 27.76) | | -22% (-94, 977) | >0.80 |
| NEOx | Media | EPA | Prostanoid | PGE3 | 0 | 21.15 (7.85, 57) | 7.69 (2.85, 20.74) | 0.79 | -64% (-95, 184) | 0.68 |
| NEOx | Media | EPA | Prostanoid | PGE3 | 15 | 18.5 (9.12, 37.53) | 7.24 (3.57, 14.69) | | -61% (-91, 61) | 0.35 |
| NEOx | Media | EPA | Prostanoid | PGE3 | 30 | 15.81 (7.27, 34.39) | 6.65 (3.06, 14.47) | | -58% (-88, 47) | 0.30 |
| NEOx | Media | EPA | Prostanoid | PGE3 | 60 | 10.76 (3.61, 32.02) | 5.24 (1.76, 15.59) | | -51% (-96, 463) | >0.80 |
| Esterified | Cell | AA | Prostanoid | PGF2a | 0 | 6.3 (1.8, 22.3) | 9.5 (2.7, 33.6) | 0.18 | 51% (-89, 2009) | >0.80 |
| Esterified | Cell | AA | Prostanoid | PGF2a | 15 | 4.6 (1.5, 14.7) | 4.8 (1.5, 15.4) | | 4% (-89, 853) | >0.80 |
| Esterified | Cell | AA | Prostanoid | PGF2a | 30 | 4.9 (1.5, 15.7) | 3.5 (1.1, 11.3) | | -28% (-91, 499) | >0.80 |
| Esterified | Cell | AA | Prostanoid | PGF2a | 60 | 15.8 (4.2, 59.5) | 5.5 (1.5, 20.6) | | -65% (-98, 551) | >0.80 |
| NEOx | Cell | AA | Prostanoid | PGF2a | 0 | 10.34 (2.5, 42.76) | 5.58 (1.35, 23.08) | 0.78 | -46% (-97, 893) | >0.80 |
| NEOx | Cell | AA | Prostanoid | PGF2a | 15 | 10.65 (2.69, 42.12) | 6.16 (1.56, 24.36) | | -42% (-96, 661) | >0.80 |
| NEOx | Cell | AA | Prostanoid | PGF2a | 30 | 9.85 (2.5, 38.85) | 6.1 (1.55, 24.07) | | -38% (-95, 658) | >0.80 |
| NEOx | Cell | AA | Prostanoid | PGF2a | 60 | 6.07 (1.41, 26.08) | 4.31 (1, 18.54) | | -29% (-97, 1566) | >0.80 |
| Esterified | Media | AA | Prostanoid | PGF2a | 0 | 16.66 (5.24, 52.93) | 7.3 (2.3, 23.19) | 0.61 | -56% (-96, 395) | >0.80 |
| Esterified | Media | AA | Prostanoid | PGF2a | 15 | 12.33 (4.59, 33.16) | 6.22 (2.31, 16.72) | | -50% (-93, 246) | >0.80 |
| Esterified | Media | AA | Prostanoid | PGF2a | 30 | 10.32 (3.73, 28.54) | 5.99 (2.16, 16.56) | | -42% (-90, 254) | >0.80 |
| Esterified | Media | AA | Prostanoid | PGF2a | 60 | 10.4 (3.03, 35.68) | 7.99 (2.33, 27.41) | | -23% (-95, 1115) | >0.80 |
| NEOx | Media | AA | Prostanoid | PGF2a | 0 | 5.1 (1.49, 17.45) | 8.47 (2.48, 28.99) | 0.52 | 66% (-87, 2054) | >0.80 |
| NEOx | Media | AA | Prostanoid | PGF2a | 15 | 5.39 (1.73, 16.78) | 7.61 (2.45, 23.66) | | 41% (-84, 1128) | >0.80 |
| NEOx | Media | AA | Prostanoid | PGF2a | 30 | 6.3 (2.01, 19.79) | 7.54 (2.4, 23.67) | | 20% (-85, 854) | >0.80 |
| NEOx | Media | AA | Prostanoid | PGF2a | 60 | 11.58 (3.2, 41.88) | 9.98 (2.76, 36.1) | | -14% (-95, 1377) | >0.80 |
| Esterified | Cell | EPA | Prostanoid | PGF3a | 0 | 217 (68, 691) | 158 (50, 502) | 0.52 | -27% (-93, 676) | >0.80 |
| Esterified | Cell | EPA | Prostanoid | PGF3a | 15 | 212 (100, 449) | 124 (58, 262) | | -42% (-87, 164) | >0.80 |
| Esterified | Cell | EPA | Prostanoid | PGF3a | 30 | 214 (90, 506) | 100 (42, 237) | | -53% (-87, 67) | 0.47 |
| Esterified | Cell | EPA | Prostanoid | PGF3a | 60 | 240 (66, 868) | 72 (20, 262) | | -70% (-98, 437) | >0.80 |
| NEOx | Cell | EPA | Prostanoid | PGF3a | 0 | 210 (59.09, 746.31) | 89.4 (25.15, 317.71) | 0.33 | -57% (-97, 495) | >0.80 |
| NEOx | Cell | EPA | Prostanoid | PGF3a | 15 | 186.15 (57.24, 605.39) | 101.9 (31.33, 331.41) | | -45% (-94, 416) | >0.80 |
| NEOx | Cell | EPA | Prostanoid | PGF3a | 30 | 153.83 (46.94, 504.13) | 108.29 (33.04, 354.88) | | -30% (-92, 508) | >0.80 |
| NEOx | Cell | EPA | Prostanoid | PGF3a | 60 | 85.11 (22.7, 319.1) | 99.07 (26.42, 371.45) | | 16% (-94, 2045) | >0.80 |
| Esterified | Media | EPA | Prostanoid | PGF3a | 0 | 273.07 (82.52, 903.6) | 223.08 (67.42, 738.16) | 0.69 | -18% (-93, 860) | >0.80 |
| Esterified | Media | EPA | Prostanoid | PGF3a | 15 | 184.74 (82.01, 416.18) | 173.29 (76.92, 390.37) | | -6% (-82, 380) | >0.80 |
| Esterified | Media | EPA | Prostanoid | PGF3a | 30 | 154.63 (62.12, 384.93) | 166.54 (66.9, 414.58) | | 8% (-74, 340) | >0.80 |
| Esterified | Media | EPA | Prostanoid | PGF3a | 60 | 205.15 (54.56, 771.41) | 291.31 (77.47, 1095.38) | | 42% (-93, 2658) | >0.80 |
| NEOx | Media | EPA | Prostanoid | PGF3a | 0 | 298.54 (97.23, 916.67) | 154.68 (50.38, 474.94) | >0.80 | -48% (-95, 440) | >0.80 |
| NEOx | Media | EPA | Prostanoid | PGF3a | 15 | 188.55 (68.2, 521.3) | 96.96 (35.07, 268.07) | | -49% (-93, 262) | >0.80 |
| NEOx | Media | EPA | Prostanoid | PGF3a | 30 | 141.58 (50.56, 396.5) | 72.26 (25.8, 202.36) | | -49% (-92, 228) | >0.80 |
| NEOx | Media | EPA | Prostanoid | PGF3a | 60 | 134.16 (41.29, 435.87) | 67.44 (20.76, 219.12) | | -50% (-96, 588) | >0.80 |
| Esterified | Cell | DHA | Diol | Protectin D1 | 0 | 1.43 (0.45, 4.57) | 2.36 (0.74, 7.55) | 0.38 | 65% (-85, 1665) | >0.80 |
| Esterified | Cell | DHA | Diol | Protectin D1 | 15 | 1.44 (0.46, 4.51) | 2 (0.64, 6.28) | | 39% (-83, 1069) | >0.80 |
| Esterified | Cell | DHA | Diol | Protectin D1 | 30 | 1.62 (0.52, 5.04) | 1.89 (0.61, 5.91) | | 17% (-85, 836) | >0.80 |
| Esterified | Cell | DHA | Diol | Protectin D1 | 60 | 2.8 (0.85, 9.2) | 2.33 (0.71, 7.66) | | -17% (-93, 960) | >0.80 |
| NEOx | Cell | DHA | Diol | Protectin D1 | 0 | 2.02 (0.93, 4.39) | 0.9 (0.42, 1.97) | 0.7 | -55% (-91, 128) | 0.67 |
| NEOx | Cell | DHA | Diol | Protectin D1 | 15 | 2.1 (1.15, 3.85) | 1.02 (0.55, 1.86) | | -52% (-85, 61) | 0.45 |
| NEOx | Cell | DHA | Diol | Protectin D1 | 30 | 1.97 (1.03, 3.74) | 1.03 (0.54, 1.95) | | -48% (-83, 56) | 0.47 |
| NEOx | Cell | DHA | Diol | Protectin D1 | 60 | 1.25 (0.54, 2.91) | 0.76 (0.33, 1.78) | | -39% (-91, 307) | >0.80 |
| Esterified | Media | DHA | Diol | Protectin D1 | 0 | 2.51 (1.15, 5.48) | 1.38 (0.63, 3.01) | 0.35 | -45% (-88, 160) | >0.80 |
| Esterified | Media | DHA | Diol | Protectin D1 | 15 | 1.79 (1.16, 2.77) | 1.24 (0.8, 1.91) | | -31% (-71, 62) | 0.78 |
| Esterified | Media | DHA | Diol | Protectin D1 | 30 | 1.47 (0.86, 2.52) | 1.27 (0.74, 2.18) | | -14% (-54, 61) | >0.80 |
| Esterified | Media | DHA | Diol | Protectin D1 | 60 | 1.51 (0.63, 3.63) | 2.05 (0.85, 4.93) | | 36% (-80, 842) | >0.80 |
| NEOx | Media | DHA | Diol | Protectin D1 | 0 | 2.12 (0.87, 5.14) | 1.26 (0.52, 3.07) | >0.80 | -40% (-91, 280) | >0.80 |
| NEOx | Media | DHA | Diol | Protectin D1 | 15 | 1.83 (0.82, 4.08) | 1.09 (0.49, 2.43) | | -40% (-87, 178) | >0.80 |
| NEOx | Media | DHA | Diol | Protectin D1 | 30 | 1.79 (0.79, 4.03) | 1.07 (0.47, 2.4) | | -40% (-86, 159) | >0.80 |
| NEOx | Media | DHA | Diol | Protectin D1 | 60 | 2.48 (0.98, 6.29) | 1.48 (0.58, 3.75) | | -40% (-92, 370) | >0.80 |
| Esterified | Cell | dgLA | Prostanoid | TXB1 | 0 | 37 (15, 90) | 19 (8, 46) | 0.09 | -48% (-92, 228) | >0.80 |
| Esterified | Cell | dgLA | Prostanoid | TXB1 | 15 | 48 (25, 93) | 39 (20, 74) | | -20% (-78, 194) | >0.80 |
| Esterified | Cell | dgLA | Prostanoid | TXB1 | 30 | 47 (23, 95) | 58 (29, 118) | | 24% (-61, 296) | >0.80 |
| Esterified | Cell | dgLA | Prostanoid | TXB1 | 60 | 18 (7, 48) | 53 (20, 141) | | 196% (-67, 2538) | 0.67 |

| | | | | | | | | | | |
|---|---|---|---|---|---|---|---|---|---|---|
| NEOx | Cell | dgLA | Prostanoid | TXB1 | 0 | 17.95 (7.99, 40.33) | 6.16 (2.74, 13.84) | 0.26 | -66% (-94, 87) | 0.40 |
| NEOx | Cell | dgLA | Prostanoid | TXB1 | 15 | 14.65 (7.27, 29.54) | 6.22 (3.08, 12.54) | | -58% (-89, 66) | 0.40 |
| NEOx | Cell | dgLA | Prostanoid | TXB1 | 30 | 10.31 (5.03, 21.17) | 5.42 (2.64, 11.12) | | -47% (-85, 90) | 0.66 |
| NEOx | Cell | dgLA | Prostanoid | TXB1 | 60 | 3.28 (1.39, 7.75) | 2.64 (1.12, 6.24) | | -20% (-88, 451) | >0.80 |
| Esterified | Media | dgLA | Prostanoid | TXB1 | 0 | 36.82 (17.49, 77.51) | 27.25 (12.94, 57.37) | 0.23 | -26% (-84, 249) | >0.80 |
| Esterified | Media | dgLA | Prostanoid | TXB1 | 15 | 39.49 (22.73, 68.61) | 37.35 (21.5, 64.89) | | -5% (-69, 185) | >0.80 |
| Esterified | Media | dgLA | Prostanoid | TXB1 | 30 | 34.93 (19.22, 63.48) | 42.21 (23.23, 76.71) | | 21% (-55, 225) | >0.80 |
| Esterified | Media | dgLA | Prostanoid | TXB1 | 60 | 15.33 (6.79, 34.62) | 30.25 (13.4, 68.3) | | 97% (-68, 1131) | >0.80 |
| NEOx | Media | dgLA | Prostanoid | TXB1 | 0 | 11.33 (6.71, 19.13) | 10.5 (6.22, 17.73) | 0.48 | -7% (-68, 172) | >0.80 |
| NEOx | Media | dgLA | Prostanoid | TXB1 | 15 | 10.74 (7.54, 15.28) | 11.08 (7.78, 15.76) | | 3% (-49, 110) | >0.80 |
| NEOx | Media | dgLA | Prostanoid | TXB1 | 30 | 10.12 (6.8, 15.06) | 11.62 (7.81, 17.3) | | 15% (-38, 111) | >0.80 |
| NEOx | Media | dgLA | Prostanoid | TXB1 | 60 | 8.86 (4.96, 15.83) | 12.61 (7.06, 22.52) | | 42% (-61, 422) | >0.80 |
| Esterified | Cell | AA | Prostanoid | TXB2 | 0 | 5.2 (2.6, 10.6) | 3.7 (1.8, 7.5) | 0.68 | -30% (-84, 208) | >0.80 |
| Esterified | Cell | AA | Prostanoid | TXB2 | 15 | 4.4 (2.5, 7.9) | 3.4 (1.9, 6) | | -25% (-76, 135) | >0.80 |
| Esterified | Cell | AA | Prostanoid | TXB2 | 30 | 4.1 (2.3, 7.6) | 3.4 (1.8, 6.1) | | -19% (-72, 133) | >0.80 |
| Esterified | Cell | AA | Prostanoid | TXB2 | 60 | 4.7 (2.2, 10.1) | 4.4 (2.1, 9.5) | | -6% (-83, 416) | >0.80 |
| NEOx | Cell | AA | Prostanoid | TXB2 | 0 | 4.20 (1.9, 9.29) | 1.96 (0.89, 4.33) | 0.06 | -53% (-91, 146) | 0.74 |
| NEOx | Cell | AA | Prostanoid | TXB2 | 15 | 3.75 (1.88, 7.48) | 2.51 (1.26, 5.01) | | -33% (-82, 155) | >0.80 |
| NEOx | Cell | AA | Prostanoid | TXB2 | 30 | 3.08 (1.52, 6.25) | 2.96 (1.46, 6.01) | | -4% (-73, 240) | >0.80 |
| NEOx | Cell | AA | Prostanoid | TXB2 | 60 | 1.62 (0.70, 3.77) | 3.22 (1.39, 7.49) | | 98% (-70, 1206) | >0.80 |
| Esterified | Media | AA | Prostanoid | TXB2 | 0 | 3.86 (1.84, 8.13) | 3.7 (1.76, 7.79) | 0.64 | -4% (-79, 341) | >0.80 |
| Esterified | Media | AA | Prostanoid | TXB2 | 15 | 4.05 (2.49, 6.60) | 3.51 (2.16, 5.72) | | -13% (-67, 131) | >0.80 |
| Esterified | Media | AA | Prostanoid | TXB2 | 30 | 4.04 (2.32, 7.05) | 3.17 (1.82, 5.53) | | -22% (-66, 80) | >0.80 |
| Esterified | Media | AA | Prostanoid | TXB2 | 60 | 3.47 (1.52, 7.94) | 2.23 (0.97, 5.09) | | -36% (-90, 307) | >0.80 |
| NEOx | Media | AA | Prostanoid | TXB2 | 0 | 5.93 (3.19, 11.01) | 2.6 (1.4, 4.82) | 0.11 | -56% (-88, 56) | 0.37 |
| NEOx | Media | AA | Prostanoid | TXB2 | 15 | 3.87 (2.58, 5.81) | 2.29 (1.53, 3.44) | | -41% (-74, 34) | 0.38 |
| NEOx | Media | AA | Prostanoid | TXB2 | 30 | 2.72 (1.71, 4.33) | 2.18 (1.37, 3.47) | | -20% (-60, 60) | >0.80 |
| NEOx | Media | AA | Prostanoid | TXB2 | 60 | 1.7 (0.85, 3.37) | 2.49 (1.25, 4.95) | | 47% (-68, 583) | >0.80 |
| Esterified | Cell | EPA | Prostanoid | TXB3 | 0 | 6.9 (2.8, 17.3) | 8.5 (3.4, 21.3) | >0.80 | 23% (-80, 654) | >0.80 |
| Esterified | Cell | EPA | Prostanoid | TXB3 | 15 | 7.1 (4.4, 11.5) | 8.2 (5.1, 13.3) | | 16% (-55, 194) | >0.80 |
| Esterified | Cell | EPA | Prostanoid | TXB3 | 30 | 7.0 (3.8, 13.1) | 7.6 (4.1, 14.2) | | 9% (-42, 102) | >0.80 |
| Esterified | Cell | EPA | Prostanoid | TXB3 | 60 | 6.0 (2.1, 17.0) | 5.8 (2.0, 16.3) | | -4% (-90, 845) | >0.80 |
| NEOx | Cell | EPA | Prostanoid | TXB3 | 0 | 8.36 (3.54, 19.75) | 3.33 (1.41, 7.87) | 0.56 | -60% (-93, 122) | 0.59 |
| NEOx | Cell | EPA | Prostanoid | TXB3 | 15 | 8.24 (5.08, 13.39) | 3.83 (2.36, 6.23) | | -53% (-82, 21) | 0.17 |
| NEOx | Cell | EPA | Prostanoid | TXB3 | 30 | 8.31 (4.57, 15.11) | 4.51 (2.48, 8.20) | | -46% (-73, 11) | 0.12 |
| NEOx | Cell | EPA | Prostanoid | TXB3 | 60 | 9.03 (3.43, 23.79) | 6.68 (2.54, 17.58) | | -26% (-91, 528) | >0.80 |
| Esterified | Media | EPA | Prostanoid | TXB3 | 0 | 7.5 (3.31, 17.02) | 6.45 (2.84, 14.64) | >0.80 | -14% (-85, 379) | >0.80 |
| Esterified | Media | EPA | Prostanoid | TXB3 | 15 | 6.78 (3.29, 13.95) | 5.6 (2.72, 11.53) | | -17% (-80, 233) | >0.80 |
| Esterified | Media | EPA | Prostanoid | TXB3 | 30 | 6.5 (3.11, 13.57) | 5.16 (2.47, 10.77) | | -21% (-79, 197) | >0.80 |
| Esterified | Media | EPA | Prostanoid | TXB3 | 60 | 7.14 (3, 17) | 5.23 (2.2, 12.45) | | -27% (-89, 407) | >0.80 |
| NEOx | Media | EPA | Prostanoid | TXB3 | 0 | 11.31 (5.58, 22.92) | 9.46 (4.67, 19.18) | >0.80 | -16% (-81, 264) | >0.80 |
| NEOx | Media | EPA | Prostanoid | TXB3 | 15 | 9.25 (4.79, 17.85) | 7.52 (3.9, 14.52) | | -19% (-77, 184) | >0.80 |
| NEOx | Media | EPA | Prostanoid | TXB3 | 30 | 7.93 (4.09, 15.37) | 6.27 (3.24, 12.16) | | -21% (-76, 163) | >0.80 |
| NEOx | Media | EPA | Prostanoid | TXB3 | 60 | 6.73 (3.22, 14.06) | 5.03 (2.41, 10.5) | | -25% (-85, 279) | >0.80 |

**Supplemental Table 15.** Principal Components: Loading matrix

| | PC1 | PC2 | PC3 | PC4 | PC5 |
|---|---|---|---|---|---|
| 10(11)-EPDPE | 0.674 | -0.069 | -0.059 | -0.241 | 0.196 |
| 10-HDOHE | 0.414 | 0.437 | 0.071 | -0.006 | -0.255 |
| 11(12)-DIHETE | 0.578 | 0.087 | -0.020 | -0.155 | -0.019 |
| 11(12)-DIHETRE | 0.697 | -0.027 | 0.088 | -0.078 | 0.240 |
| 11(12)-EPETE | 0.466 | -0.031 | 0.570 | 0.134 | -0.127 |
| 11(12)-EPETRE | 0.593 | 0.302 | -0.218 | -0.195 | 0.148 |
| 11-HDOHE | 0.477 | 0.097 | -0.167 | 0.098 | 0.035 |
| 11-HEPE | 0.172 | 0.578 | 0.158 | -0.002 | 0.147 |
| 12(13)-DIHOME | 0.514 | -0.387 | 0.438 | -0.034 | -0.104 |
| 12(13)-EPOME | 0.667 | 0.084 | 0.202 | 0.249 | 0.188 |
| 12-HEPE | 0.340 | 0.325 | -0.201 | 0.200 | 0.102 |
| 12-HETE | 0.504 | -0.410 | -0.439 | 0.369 | 0.142 |
| 12-HPETE | 0.586 | 0.217 | -0.213 | 0.140 | -0.236 |
| 12-KETE | 0.570 | -0.086 | 0.281 | -0.236 | 0.223 |
| 13(14)-EPDPE | 0.381 | 0.446 | 0.009 | 0.061 | -0.106 |
| 13-HDOHE | 0.565 | 0.121 | -0.084 | 0.199 | -0.208 |
| 13-HODE | 0.633 | -0.086 | 0.492 | 0.243 | -0.055 |
| 13-HOTRE | 0.325 | 0.098 | 0.013 | 0.065 | 0.464 |
| 13-KODE | 0.485 | -0.117 | 0.222 | 0.359 | -0.167 |
| 14(15)-DIHETE | 0.691 | -0.266 | -0.173 | -0.105 | 0.015 |
| 14(15)-DIHETRE | 0.636 | -0.030 | -0.236 | -0.373 | -0.194 |
| 14(15)-EPETE | 0.841 | -0.040 | 0.152 | -0.171 | -0.132 |
| 14(15)-EPETRE | 0.483 | 0.087 | -0.230 | 0.194 | -0.450 |
| 14-HDOHE | 0.586 | 0.301 | -0.134 | 0.072 | -0.206 |
| 15-HEPE | 0.083 | -0.119 | -0.182 | -0.079 | 0.299 |
| 15-HETE | 0.603 | -0.565 | -0.326 | 0.107 | 0.136 |
| 15-HPETE | 0.710 | 0.083 | 0.028 | -0.294 | -0.064 |
| 15-KETE | 0.587 | -0.160 | 0.010 | 0.169 | 0.207 |
| 16(17)-EPDPE | 0.474 | 0.350 | -0.005 | -0.108 | 0.092 |
| 16-HDOHE | 0.456 | 0.257 | -0.101 | 0.081 | -0.013 |
| 17(18)-DIHETE | 0.566 | -0.035 | 0.165 | -0.409 | -0.217 |
| 17(18)-EPETE | 0.640 | -0.170 | 0.152 | -0.347 | 0.049 |
| 17-HDOHE | 0.651 | -0.057 | -0.255 | 0.035 | -0.161 |
| 18-HEPE | 0.584 | -0.340 | -0.271 | 0.117 | 0.417 |
| 19(20)-EPDPE | 0.391 | 0.199 | -0.345 | 0.236 | -0.348 |
| 20-HDOHE | 0.657 | 0.007 | -0.221 | 0.119 | -0.053 |
| 22-HDOHE | 0.299 | 0.476 | -0.169 | 0.094 | -0.078 |
| 4-HDOHE | 0.553 | 0.055 | -0.368 | 0.088 | 0.034 |
| 5-HEPE | 0.345 | 0.513 | 0.005 | -0.011 | 0.174 |
| 5-HETE | 0.687 | -0.470 | -0.301 | 0.179 | 0.134 |
| 5-KETE | 0.586 | -0.121 | 0.281 | 0.140 | 0.102 |
| 7(8)-EPDPE | 0.355 | 0.465 | 0.089 | -0.048 | 0.133 |
| 7-HDOHE | 0.503 | 0.335 | -0.051 | -0.059 | 0.001 |
| 8(9)-DIHETE | 0.532 | -0.282 | 0.011 | -0.586 | -0.112 |
| 8(9)-DIHETRE | 0.794 | 0.182 | 0.162 | 0.007 | -0.097 |
| 8(9)-EPETE | 0.604 | -0.203 | 0.199 | 0.202 | 0.122 |
| 8(9)-EPETRE | 0.763 | -0.033 | -0.194 | -0.163 | 0.190 |
| 8-HDOHE | 0.537 | 0.054 | -0.332 | 0.154 | -0.357 |
| 8-HEPE | 0.652 | 0.237 | 0.154 | -0.245 | 0.191 |
| 9(10)-DIHOME | 0.427 | -0.427 | 0.233 | 0.139 | -0.114 |
| 9(10)-EPOME | 0.617 | 0.142 | 0.410 | 0.378 | 0.067 |
| 9-HEPE | 0.679 | 0.198 | -0.071 | 0.182 | 0.275 |
| 9-HETE | 0.414 | -0.258 | -0.346 | 0.012 | 0.076 |
| 9-HODE | 0.804 | -0.214 | 0.285 | -0.003 | -0.161 |
| 9-HOTRE | 0.144 | 0.374 | 0.376 | 0.098 | 0.392 |
| 9-KODE | 0.342 | -0.346 | 0.480 | 0.296 | -0.298 |
| RESOLVIN D1 | 0.628 | -0.139 | 0.034 | -0.512 | -0.219 |

**Supplemental Table 16**: Lipoprotein oxylipins, Principal components analysis

| Principal Component | Plasma lipid pool | Surgery | Genotype<br>WT<br>Mean (CI) | <br>KO<br>Mean (CI) | Group difference within Surgery<br>Difference (CI) | <br><br>pval |
|---|---|---|---|---|---|---|
| PCA1 | Alb | SHAM | -2.04 (-3.08, -1.01) | -10.47 (-11.61, -9.34) | -8.43 (-11.11, -5.75) | <0.0001 |
| PCA1 | Alb | TAC | -4.4 (-5.17, -3.64) | -4.79 (-5.59, -3.99) | -0.39 (-2.33, 1.55) | >0.80 |
| | Within genotype group difference | Difference (CI) | -2.36 (-0.11, -4.61) | 5.68 (8.11, 3.25) | | |
| | | pval | 0.03 | <0.0001 | | |
| PCA1 | HDL | SHAM | 6.74 (6.01, 7.47) | 1.84 (1.05, 2.64) | -4.9 (-6.8, -3) | <0.0001 |
| PCA1 | HDL | TAC | 4.78 (4.24, 5.32) | 1.88 (1.31, 2.44) | -2.91 (-4.28, -1.54) | <0.0001 |
| . | | Difference (CI) | -1.96 (-0.37, -3.55) | 0.03 (1.75, -1.68) | . | . |
| . | . | . | 0.004 | >0.80 | . | . |
| PCA1 | LDL | SHAM | 6.17 (5.08, 7.27) | -3.03 (-4.23, -1.83) | -9.2 (-12.06, -6.34) | <0.0001 |
| PCA1 | LDL | TAC | -0.05 (-0.86, 0.77) | -0.42 (-1.27, 0.43) | -0.37 (-2.44, 1.69) | >0.80 |
| . | | Difference (CI) | -6.22 (-3.82, -8.62) | 2.61 (5.19, 0.02) | . | . |
| . | . | . | <0.0001 | 0.05 | . | . |
| PCA1 | VLDL | SHAM | 5.58 (4.62, 6.55) | -3.06 (-4.12, -2) | -8.65 (-11.16, -6.13) | <0.0001 |
| PCA1 | VLDL | TAC | 0.58 (-0.14, 1.29) | -0.18 (-0.93, 0.56) | -0.76 (-2.58, 1.05) | >0.80 |
| . | | Difference (CI) | -5.01 (-2.9, -7.12) | 2.88 (5.16, 0.6) | . | . |
| . | . | . | <0.0001 | 0.003 | . | . |
| PCA2 | Alb | SHAM | -2.04 (-3.08, -1.01) | -10.47 (-11.61, -9.34) | -1.98 (-4.8, 0.84) | 0.47 |
| PCA2 | Alb | TAC | -4.4 (-5.17, -3.64) | -4.79 (-5.59, -3.99) | 0.7 (-1.33, 2.74) | >0.80 |
| . | | Difference (CI) | 0.48 (2.85, -1.88) | 3.17 (5.72, 0.62) | . | . |
| . | . | . | >0.80 | 0.004 | . | . |
| PCA2 | HDL | SHAM | 6.74 (6.01, 7.47) | 1.84 (1.05, 2.64) | -3.56 (-5.07, -2.05) | <0.0001 |
| PCA2 | HDL | TAC | 4.78 (4.24, 5.32) | 1.88 (1.31, 2.44) | 1.82 (0.73, 2.91) | <0.0001 |
| . | | Difference (CI) | -4.87 (-3.6, -6.13) | 0.51 (1.87, -0.86) | . | . |
| . | . | . | <0.0001 | >0.80 | . | . |
| PCA2 | LDL | SHAM | 6.17 (5.08, 7.27) | -3.03 (-4.23, -1.83) | -1.94 (-4.19, 0.31) | 0.16 |
| PCA2 | LDL | TAC | -0.05 (-0.86, 0.77) | -0.42 (-1.27, 0.43) | 0.81 (-0.81, 2.43) | >0.80 |
| . | | Difference (CI) | -2.16 (-0.27, -4.04) | 0.59 (2.62, -1.44) | . | . |
| . | . | . | 0.01 | >0.80 | . | . |
| PCA2 | VLDL | SHAM | 5.58 (4.62, 6.55) | -3.06 (-4.12, -2) | -3.37 (-5.39, -1.36) | <0.0001 |
| PCA2 | VLDL | TAC | 0.58 (-0.14, 1.29) | -0.18 (-0.93, 0.56) | 1.06 (-0.4, 2.51) | 0.42 |
| . | | Difference (CI) | -3.89 (-2.2, -5.58) | 0.54 (2.36, -1.29) | . | . |
| . | . | . | <0.0001 | >0.80 | . | . |
| PCA3 | Alb | SHAM | -2.81 (-3.64, -1.98) | 2.01 (1.1, 2.92) | 4.82 (2.66, 6.99) | <0.0001 |
| PCA3 | Alb | TAC | -2.67 (-3.28, -2.05) | 0.1 (-0.55, 0.74) | 2.76 (1.2, 4.33) | <0.0001 |
| . | | Difference (CI) | 0.14 (1.96, -1.67) | -1.92 (0.04, -3.88) | . | . |
| . | . | . | >0.80 | 0.06 | . | . |
| PCA3 | HDL | SHAM | 0.46 (-0.05, 0.97) | -3.06 (-3.62, -2.5) | -3.52 (-4.85, -2.19) | <0.0001 |
| PCA3 | HDL | TAC | -0.06 (-0.44, 0.32) | -1.13 (-1.53, -0.74) | -1.07 (-2.03, -0.11) | 0.02 |
| . | | Difference (CI) | -0.52 (0.6, -1.64) | 1.93 (3.13, 0.72) | . | . |
| . | . | . | >0.80 | <0.0001 | . | . |
| PCA3 | LDL | SHAM | 0.27 (-0.51, 1.05) | 0.81 (-0.04, 1.66) | 0.54 (-1.49, 2.57) | >0.80 |
| PCA3 | LDL | TAC | 0.49 (-0.08, 1.07) | 0.29 (-0.31, 0.89) | -0.2 (-1.67, 1.26) | >0.80 |
| . | | Difference (CI) | 0.22 (1.92, -1.48) | -0.52 (1.32, -2.36) | . | . |
| . | . | . | >0.80 | >0.80 | . | . |
| PCA3 | VLDL | SHAM | 0.4 (-0.43, 1.22) | -0.18 (-1.08, 0.73) | -0.58 (-2.73, 1.58) | >0.80 |
| PCA3 | VLDL | TAC | 3.29 (2.69, 3.9) | 0.8 (0.16, 1.44) | -2.5 (-4.05, -0.94) | <0.0001 |
| . | | Difference (CI) | 2.9 (4.7, 1.09) | 0.98 (2.92, -0.97) | . | . |
| . | . | . | <0.0001 | >0.80 | . | . |
| PCA4 | Alb | SHAM | 1.76 (0.95, 2.56) | 2.93 (2.05, 3.82) | 2.02 (-0.11, 4.15) | 0.08 |
| PCA4 | Alb | TAC | -0.45 (-1.05, 0.15) | 1.51 (0.88, 2.14) | -1.11 (-2.64, 0.43) | 0.43 |
| . | | Difference (CI) | 1.73 (3.51, -0.06) | -1.4 (0.53, -3.32) | . | . |
| . | . | . | 0.07 | 0.42 | . | . |
| PCA4 | HDL | SHAM | 1.76 (1.33, 2.19) | 1.56 (1.09, 2.03) | -4.1 (-5.41, -2.79) | <0.0001 |
| PCA4 | HDL | TAC | 0.46 (0.14, 0.77) | -0.82 (-1.15, -0.49) | 3.01 (2.06, 3.95) | <0.0001 |
| . | | Difference (CI) | -1.67 (-0.57, -2.77) | 5.43 (6.62, 4.25) | . | . |
| . | . | . | 0.0001 | <0.0001 | . | . |
| PCA4 | LDL | SHAM | 0.7 (0.18, 1.23) | -1.32 (-1.89, -0.75) | -2.95 (-4.19, -1.71) | <0.0001 |
| PCA4 | LDL | TAC | -0.89 (-1.28, -0.5) | -2.1 (-2.5, -1.69) | 1.49 (0.6, 2.39) | <0.0001 |
| . | | Difference (CI) | -0.61 (0.43, -1.65) | 3.83 (4.96, 2.71) | . | . |
| . | . | . | 0.75 | <0.0001 | . | . |
| PCA4 | VLDL | SHAM | 0.55 (0.08, 1.03) | -1.74 (-2.26, -1.22) | -1.69 (-3.41, 0.04) | 0.06 |

| | | | | | | |
|---|---|---|---|---|---|---|
| PCA4 | VLDL | TAC | 1.01 (0.66, 1.36) | -2.32 (-2.68, -1.95) | 0.12 (-1.12, 1.37) | >0.80 |
| . | | Difference (CI) | -0.7 (0.75, -2.15) | 1.11 (2.67, -0.45) | . | . |
| . | . | . | >0.80 | 0.45 | . | . |
| PCA5 | Alb | SHAM | -0.61 (-1.42, 0.21) | 1.41 (0.52, 2.31) | 2.02 (-0.11, 4.15) | 0.08 |
| PCA5 | Alb | TAC | 1.12 (0.52, 1.73) | 0.02 (-0.62, 0.65) | -1.11 (-2.64, 0.43) | 0.43 |
| . | | Difference (CI) | 1.73 (3.51, -0.06) | -1.4 (0.53, -3.32) | . | . |
| . | . | . | 0.07 | 0.42 | . | . |
| PCA5 | HDL | SHAM | 1.36 (0.86, 1.86) | -2.74 (-3.29, -2.19) | -4.1 (-5.41, -2.79) | <0.0001 |
| PCA5 | HDL | TAC | -0.31 (-0.68, 0.06) | 2.7 (2.31, 3.09) | 3.01 (2.06, 3.95) | <0.0001 |
| . | | Difference (CI) | -1.67 (-0.57, -2.77) | 5.43 (6.62, 4.25) | . | . |
| . | . | . | 0.0001 | <0.0001 | . | . |
| PCA5 | LDL | SHAM | -0.03 (-0.5, 0.45) | -2.98 (-3.5, -2.46) | -2.95 (-4.19, -1.71) | <0.0001 |
| PCA5 | LDL | TAC | -0.64 (-0.99, -0.28) | 0.86 (0.49, 1.22) | 1.49 (0.6, 2.39) | <0.0001 |
| . | | Difference (CI) | -0.61 (0.43, -1.65) | 3.83 (4.96, 2.71) | . | . |
| . | . | . | 0.75 | <0.0001 | . | . |
| PCA5 | VLDL | SHAM | 0.05 (-0.62, 0.71) | -1.64 (-2.37, -0.92) | -1.69 (-3.41, 0.04) | 0.06 |
| PCA5 | VLDL | TAC | -0.66 (-1.14, -0.17) | -0.53 (-1.05, -0.02) | 0.12 (-1.12, 1.37) | >0.80 |
| . | | Difference (CI) | -0.7 (0.75, -2.15) | 1.11 (2.67, -0.45) | . | . |
| . | . | . | >0.80 | 0.45 | . | . |

**Supplemental Table 17**: Individual plasma oxylipins

| Oxylipin | Parent PUFA | Chemistry | Lipoprotein | Surgery | FFAR4 Status | Mean (CI) | Full Intx pval | Post Hoc Test | %Diff (CI) | Tukey post hoc pval |
|---|---|---|---|---|---|---|---|---|---|---|
| 9-HODE | LA | Alcohol | Alb | SHAM | WT | 11.5 (8.7, 15.2) | <0.0001 | WT SHAM vs KO SHAM | -82% (-91, -63) | <0.0001 |
| 9-HODE | LA | Alcohol | Alb | SHAM | KO | 2.1 (1.5, 2.8) | . | TAC WT vs TAC KO | 37% (-19, 131) | 0.71 |
| 9-HODE | LA | Alcohol | Alb | TAC | WT | 3.2 (2.6, 4) | . | SHAM WT vs TAC WT | -255% (-550, -94) | <0.0001 |
| 9-HODE | LA | Alcohol | Alb | TAC | KO | 4.4 (3.6, 5.5) | . | SHAM KO - TAC KO | 54% (76, 11) | 0.008 |
| 13-HODE | LA | Alcohol | Alb | SHAM | WT | 9.2 (6.6, 12.8) | 0.0003 | WT SHAM vs KO SHAM | 2% (-57, 139) | >0.80 |
| 13-HODE | LA | Alcohol | Alb | SHAM | KO | 9.4 (6.5, 13.4) | . | TAC WT vs TAC KO | 97% (6, 265) | 0.02 |
| 13-HODE | LA | Alcohol | Alb | TAC | WT | 4.1 (3.2, 5.2) | . | SHAM WT vs TAC WT | -127% (-364, -11) | 0.01 |
| 13-HODE | LA | Alcohol | Alb | TAC | KO | 8 (6.2, 10.3) | . | SHAM KO - TAC KO | -17% (46, -153) | >0.80 |
| 9-HOTrE | aLA | Alcohol | Alb | SHAM | WT | 0.24 (0.13, 0.47) | <0.0001 | WT SHAM vs KO SHAM | -27% (-87, 313) | >0.80 |
| 9-HOTrE | aLA | Alcohol | Alb | SHAM | KO | 0.18 (0.09, 0.37) | . | TAC WT vs TAC KO | 31% (-62, 356) | >0.80 |
| 9-HOTrE | aLA | Alcohol | Alb | TAC | WT | 0.14 (0.08, 0.22) | . | SHAM WT vs TAC WT | -79% (-664, 58) | >0.80 |
| 9-HOTrE | aLA | Alcohol | Alb | TAC | KO | 0.18 (0.11, 0.3) | . | SHAM KO - TAC KO | 0% (79, -379) | >0.80 |
| 13-HOTrE | aLA | Alcohol | Alb | SHAM | WT | 0.28 (0.13, 0.61) | 0.0002 | WT SHAM vs KO SHAM | 13% (-85, 770) | >0.80 |
| 13-HOTrE | aLA | Alcohol | Alb | SHAM | KO | 0.31 (0.13, 0.74) | . | TAC WT vs TAC KO | -39% (-86, 164) | >0.80 |
| 13-HOTrE | aLA | Alcohol | Alb | TAC | WT | 0.36 (0.2, 0.64) | . | SHAM WT vs TAC WT | 23% (-326, 86) | >0.80 |
| 13-HOTrE | aLA | Alcohol | Alb | TAC | KO | 0.22 (0.12, 0.4) | . | SHAM KO - TAC KO | -44% (77, -811) | >0.80 |
| 5-HETE | AA | Alcohol | Alb | SHAM | WT | 1.37 (0.62, 3.06) | <0.0001 | WT SHAM vs KO SHAM | -96% (-100, -72) | <0.0001 |
| 5-HETE | AA | Alcohol | Alb | SHAM | KO | 0.05 (0.02, 0.12) | . | TAC WT vs TAC KO | -71% (-94, 31) | 0.23 |
| 5-HETE | AA | Alcohol | Alb | TAC | WT | 0.2 (0.11, 0.36) | . | SHAM WT vs TAC WT | -598% (-3925, -21) | 0.02 |
| 5-HETE | AA | Alcohol | Alb | TAC | KO | 0.06 (0.03, 0.11) | . | SHAM KO - TAC KO | 16% (87, -459) | >0.80 |
| 9-HETE | AA | Alcohol | Alb | SHAM | WT | 1.45 (0.8, 2.64) | 0.01 | WT SHAM vs KO SHAM | -74% (-95, 22) | 0.15 |
| 9-HETE | AA | Alcohol | Alb | SHAM | KO | 0.37 (0.19, 0.72) | . | TAC WT vs TAC KO | -59% (-87, 26) | 0.27 |
| 9-HETE | AA | Alcohol | Alb | TAC | WT | 0.24 (0.15, 0.37) | . | SHAM WT vs TAC WT | -511% (-2162, -65) | 0.0008 |
| 9-HETE | AA | Alcohol | Alb | TAC | KO | 0.1 (0.06, 0.15) | . | SHAM KO - TAC KO | -283% (7, -1475) | 0.08 |
| 12-HETE | AA | Alcohol | Alb | SHAM | WT | 1.3 (0.69, 2.43) | <0.0001 | WT SHAM vs KO SHAM | -97% (-99, -82) | <0.0001 |
| 12-HETE | AA | Alcohol | Alb | SHAM | KO | 0.05 (0.02, 0.09) | . | TAC WT vs TAC KO | 0% (-70, 228) | >0.80 |
| 12-HETE | AA | Alcohol | Alb | TAC | WT | 0.33 (0.21, 0.53) | . | SHAM WT vs TAC WT | -287% (-1442, 3) | 0.06 |
| 12-HETE | AA | Alcohol | Alb | TAC | KO | 0.33 (0.21, 0.54) | . | SHAM KO - TAC KO | 87% (97, 40) | 0.001 |
| 15-HETE | AA | Alcohol | Alb | SHAM | WT | 1.29 (0.72, 2.29) | <0.0001 | WT SHAM vs KO SHAM | -94% (-99, -71) | <0.0001 |
| 15-HETE | AA | Alcohol | Alb | SHAM | KO | 0.08 (0.04, 0.15) | . | TAC WT vs TAC KO | -69% (-89, -7) | 0.03 |
| 15-HETE | AA | Alcohol | Alb | TAC | WT | 0.26 (0.17, 0.39) | . | SHAM WT vs TAC WT | -404% (-1683, -43) | 0.003 |
| 15-HETE | AA | Alcohol | Alb | TAC | KO | 0.08 (0.05, 0.12) | . | SHAM KO - TAC KO | -2% (74, -299) | >0.80 |
| 5-HEPE | EPA | Alcohol | Alb | SHAM | WT | 0.15 (0.08, 0.28) | <0.0001 | WT SHAM vs KO SHAM | -22% (-85, 313) | >0.80 |
| 5-HEPE | EPA | Alcohol | Alb | SHAM | KO | 0.11 (0.06, 0.23) | . | TAC WT vs TAC KO | -4% (-71, 218) | >0.80 |
| 5-HEPE | EPA | Alcohol | Alb | TAC | WT | 0.21 (0.13, 0.33) | . | SHAM WT vs TAC WT | 29% (-186, 82) | >0.80 |
| 5-HEPE | EPA | Alcohol | Alb | TAC | KO | 0.2 (0.12, 0.32) | . | SHAM KO - TAC KO | 42% (87, -161) | >0.80 |
| 8-HEPE | EPA | Alcohol | Alb | SHAM | WT | 0.05 (0.02, 0.12) | <0.0001 | WT SHAM vs KO SHAM | 68% (-78, 1190) | >0.80 |
| 8-HEPE | EPA | Alcohol | Alb | SHAM | KO | 0.09 (0.04, 0.21) | . | TAC WT vs TAC KO | -44% (-87, 142) | >0.80 |
| 8-HEPE | EPA | Alcohol | Alb | TAC | WT | 0.22 (0.12, 0.39) | . | SHAM WT vs TAC WT | 76% (-35, 96) | 0.22 |
| 8-HEPE | EPA | Alcohol | Alb | TAC | KO | 0.12 (0.07, 0.22) | . | SHAM KO - TAC KO | 26% (88, -367) | >0.80 |
| 9-HEPE | EPA | Alcohol | Alb | SHAM | WT | 0.34 (0.14, 0.83) | 0.1 | . | . | . |
| 9-HEPE | EPA | Alcohol | Alb | SHAM | KO | 0.03 (0.01, 0.09) | . | . | . | . |
| 9-HEPE | EPA | Alcohol | Alb | TAC | WT | 0.15 (0.08, 0.3) | . | . | . | . |
| 9-HEPE | EPA | Alcohol | Alb | TAC | KO | 0.17 (0.09, 0.35) | . | . | . | . |
| 11-HEPE | EPA | Alcohol | Alb | SHAM | WT | 0.06 (0.03, 0.14) | 0.02 | WT SHAM vs KO SHAM | -5% (-87, 615) | >0.80 |
| 11-HEPE | EPA | Alcohol | Alb | SHAM | KO | 0.06 (0.03, 0.14) | . | TAC WT vs TAC KO | 155% (-41, 989) | 0.62 |
| 11-HEPE | EPA | Alcohol | Alb | TAC | WT | 0.07 (0.04, 0.12) | . | SHAM WT vs TAC WT | 6% (-408, 83) | >0.80 |
| 11-HEPE | EPA | Alcohol | Alb | TAC | KO | 0.18 (0.1, 0.32) | . | SHAM KO - TAC KO | 65% (94, -117) | 0.78 |
| 12-HEPE | EPA | Alcohol | Alb | SHAM | WT | 0.24 (0.13, 0.43) | <0.0001 | WT SHAM vs KO SHAM | -4% (-79, 343) | >0.80 |
| 12-HEPE | EPA | Alcohol | Alb | SHAM | KO | 0.23 (0.12, 0.44) | . | TAC WT vs TAC KO | -30% (-77, 110) | >0.80 |
| 12-HEPE | EPA | Alcohol | Alb | TAC | WT | 0.15 (0.1, 0.23) | . | SHAM WT vs TAC WT | -65% (-494, 54) | >0.80 |
| 12-HEPE | EPA | Alcohol | Alb | TAC | KO | 0.1 (0.06, 0.16) | . | SHAM KO - TAC KO | -126% (43, -804) | 0.74 |
| 15-HEPE | EPA | Alcohol | Alb | SHAM | WT | 0.29 (0.16, 0.55) | 0.24 | . | . | . |
| 15-HEPE | EPA | Alcohol | Alb | SHAM | KO | 0.13 (0.06, 0.26) | . | . | . | . |
| 15-HEPE | EPA | Alcohol | Alb | TAC | WT | 0.3 (0.19, 0.48) | . | . | . | . |
| 15-HEPE | EPA | Alcohol | Alb | TAC | KO | 0.29 (0.18, 0.48) | . | . | . | . |
| 18-HEPE | EPA | Alcohol | Alb | SHAM | WT | 0.13 (0.08, 0.2) | <0.0001 | WT SHAM vs KO SHAM | -76% (-93, -22) | 0.006 |
| 18-HEPE | EPA | Alcohol | Alb | SHAM | KO | 0.03 (0.02, 0.05) | . | TAC WT vs TAC KO | -76% (-90, -43) | <0.0001 |
| 18-HEPE | EPA | Alcohol | Alb | TAC | WT | 0.19 (0.14, 0.26) | . | SHAM WT vs TAC WT | 33% (-82, 75) | >0.80 |
| 18-HEPE | EPA | Alcohol | Alb | TAC | KO | 0.05 (0.03, 0.07) | . | SHAM KO - TAC KO | 34% (77, -93) | >0.80 |
| 4-HDoHE | DHA | Alcohol | Alb | SHAM | WT | 0.13 (0.04, 0.38) | 0.01 | WT SHAM vs KO SHAM | -84% (-99, 157) | 0.57 |
| 4-HDoHE | DHA | Alcohol | Alb | SHAM | KO | 0.02 (0.01, 0.07) | . | TAC WT vs TAC KO | -13% (-88, 546) | >0.80 |
| 4-HDoHE | DHA | Alcohol | Alb | TAC | WT | 0.15 (0.07, 0.34) | . | SHAM WT vs TAC WT | 16% (-767, 92) | >0.80 |
| 4-HDoHE | DHA | Alcohol | Alb | TAC | KO | 0.13 (0.06, 0.31) | . | SHAM KO - TAC KO | 85% (99, -91) | 0.38 |
| 7-HDoHE | DHA | Alcohol | Alb | SHAM | WT | 0.22 (0.09, 0.54) | 0.001 | WT SHAM vs KO SHAM | -88% (-99, 26) | 0.12 |
| 7-HDoHE | DHA | Alcohol | Alb | SHAM | KO | 0.03 (0.01, 0.07) | . | TAC WT vs TAC KO | -14% (-84, 362) | >0.80 |
| 7-HDoHE | DHA | Alcohol | Alb | TAC | WT | 0.24 (0.13, 0.47) | . | SHAM WT vs TAC WT | 11% (-532, 87) | >0.80 |
| 7-HDoHE | DHA | Alcohol | Alb | TAC | KO | 0.21 (0.1, 0.42) | . | SHAM KO - TAC KO | 87% (98, -5) | 0.06 |
| 8-HDoHE | DHA | Alcohol | Alb | SHAM | WT | 0.36 (0.19, 0.68) | 0.0002 | WT SHAM vs KO SHAM | -82% (-97, 0) | 0.05 |
| 8-HDoHE | DHA | Alcohol | Alb | SHAM | KO | 0.06 (0.03, 0.13) | . | TAC WT vs TAC KO | -15% (-75, 190) | >0.80 |
| 8-HDoHE | DHA | Alcohol | Alb | TAC | WT | 0.32 (0.2, 0.52) | . | SHAM WT vs TAC WT | -11% (-360, 73) | >0.80 |
| 8-HDoHE | DHA | Alcohol | Alb | TAC | KO | 0.27 (0.16, 0.45) | . | SHAM KO - TAC KO | 76% (95, -10) | 0.09 |
| 10-HDoHE | DHA | Alcohol | Alb | SHAM | WT | 0.16 (0.07, 0.39) | 0.02 | WT SHAM vs KO SHAM | -77% (-98, 132) | 0.64 |
| 10-HDoHE | DHA | Alcohol | Alb | SHAM | KO | 0.04 (0.01, 0.1) | . | TAC WT vs TAC KO | 112% (-60, 1017) | >0.80 |
| 10-HDoHE | DHA | Alcohol | Alb | TAC | WT | 0.14 (0.07, 0.28) | . | SHAM WT vs TAC WT | -13% (-680, 84) | >0.80 |
| 10-HDoHE | DHA | Alcohol | Alb | TAC | KO | 0.3 (0.15, 0.6) | . | SHAM KO - TAC KO | 88% (98, 1) | 0.05 |
| 11-HDoHE | DHA | Alcohol | Alb | SHAM | WT | 0.16 (0.05, 0.46) | 0.24 | . | . | . |
| 11-HDoHE | DHA | Alcohol | Alb | SHAM | KO | 0.02 (0.01, 0.06) | . | . | . | . |
| 11-HDoHE | DHA | Alcohol | Alb | TAC | WT | 0.14 (0.06, 0.3) | . | . | . | . |
| 11-HDoHE | DHA | Alcohol | Alb | TAC | KO | 0.15 (0.06, 0.34) | . | . | . | . |
| 13-HDoHE | DHA | Alcohol | Alb | SHAM | WT | 0.23 (0.12, 0.46) | 0.0009 | WT SHAM vs KO SHAM | -86% (-98, -17) | 0.02 |
| 13-HDoHE | DHA | Alcohol | Alb | SHAM | KO | 0.03 (0.01, 0.07) | . | TAC WT vs TAC KO | 85% (-49, 572) | >0.80 |

| | | | | | | | | | | |
|---|---|---|---|---|---|---|---|---|---|---|
| 13-HDoHE | DHA | Alcohol | Alb | TAC | WT | 0.16 (0.1, 0.27) | . | SHAM WT vs TAC WT | -41% (-532, 68) | >0.80 |
| 13-HDoHE | DHA | Alcohol | Alb | TAC | KO | 0.3 (0.18, 0.51) | . | SHAM KO - TAC KO | 89% (98, 47) | 0.0007 |
| 14-HDoHE | DHA | Alcohol | Alb | SHAM | WT | 0.26 (0.1, 0.67) | <0.0001 | WT SHAM vs KO SHAM | -89% (-99, 25) | 0.11 |
| 14-HDoHE | DHA | Alcohol | Alb | SHAM | KO | 0.03 (0.01, 0.08) | . | TAC WT vs TAC KO | 41% (-76, 719) | >0.80 |
| 14-HDoHE | DHA | Alcohol | Alb | TAC | WT | 0.19 (0.1, 0.39) | . | SHAM WT vs TAC WT | -36% (-944, 82) | >0.80 |
| 14-HDoHE | DHA | Alcohol | Alb | TAC | KO | 0.27 (0.13, 0.56) | . | SHAM KO - TAC KO | 90% (99, 5) | 0.04 |
| 16-HDoHE | DHA | Alcohol | Alb | SHAM | WT | 0.1 (0.03, 0.27) | 0.13 | . | . | . |
| 16-HDoHE | DHA | Alcohol | Alb | SHAM | KO | 0.01 (0, 0.04) | . | . | . | . |
| 16-HDoHE | DHA | Alcohol | Alb | TAC | WT | 0.03 (0.02, 0.07) | . | . | . | . |
| 16-HDoHE | DHA | Alcohol | Alb | TAC | KO | 0.04 (0.02, 0.09) | . | . | . | . |
| 17-HDoHE | DHA | Alcohol | Alb | SHAM | WT | 0.32 (0.13, 0.79) | <0.0001 | WT SHAM vs KO SHAM | -90% (-99, 11) | 0.07 |
| 17-HDoHE | DHA | Alcohol | Alb | SHAM | KO | 0.03 (0.01, 0.09) | . | TAC WT vs TAC KO | -26% (-87, 311) | >0.80 |
| 17-HDoHE | DHA | Alcohol | Alb | TAC | WT | 0.33 (0.17, 0.66) | . | SHAM WT vs TAC WT | 5% (-597, 87) | >0.80 |
| 17-HDoHE | DHA | Alcohol | Alb | TAC | KO | 0.25 (0.12, 0.5) | . | SHAM KO - TAC KO | 87% (98, -13) | 0.09 |
| 20-HDoHE | DHA | Alcohol | Alb | SHAM | WT | 0.17 (0.07, 0.39) | 0.61 | . | . | . |
| 20-HDoHE | DHA | Alcohol | Alb | SHAM | KO | 0.03 (0.01, 0.07) | . | . | . | . |
| 20-HDoHE | DHA | Alcohol | Alb | TAC | WT | 0.14 (0.07, 0.25) | . | . | . | . |
| 20-HDoHE | DHA | Alcohol | Alb | TAC | KO | 0.12 (0.06, 0.22) | . | . | . | . |
| 22-HDoHE | DHA | Alcohol | Alb | SHAM | WT | 0.15 (0.07, 0.31) | 0.51 | . | . | . |
| 22-HDoHE | DHA | Alcohol | Alb | SHAM | KO | 0.04 (0.02, 0.08) | . | . | . | . |
| 22-HDoHE | DHA | Alcohol | Alb | TAC | WT | 0.18 (0.11, 0.31) | . | . | . | . |
| 22-HDoHE | DHA | Alcohol | Alb | TAC | KO | 0.14 (0.08, 0.25) | . | . | . | . |
| 9-KODE | LA | Ketone | Alb | SHAM | WT | 3.7 (2.35, 5.84) | <0.0001 | WT SHAM vs KO SHAM | 36% (-59, 346) | >0.80 |
| 9-KODE | LA | Ketone | Alb | SHAM | KO | 5.04 (3.06, 8.29) | . | TAC WT vs TAC KO | 458% (137, 1215) | <0.0001 |
| 9-KODE | LA | Ketone | Alb | TAC | WT | 0.4 (0.29, 0.56) | . | SHAM WT vs TAC WT | -828% (-2412, -243) | <0.0001 |
| 9-KODE | LA | Ketone | Alb | TAC | KO | 2.23 (1.56, 3.17) | . | SHAM KO - TAC KO | -126% (23, -563) | 0.34 |
| 13-KODE | LA | Ketone | Alb | SHAM | WT | 1.87 (1.15, 3.05) | <0.0001 | WT SHAM vs KO SHAM | -6% (-74, 239) | >0.80 |
| 13-KODE | LA | Ketone | Alb | SHAM | KO | 1.77 (1.03, 3.02) | . | TAC WT vs TAC KO | -9% (-64, 130) | >0.80 |
| 13-KODE | LA | Ketone | Alb | TAC | WT | 1.3 (0.91, 1.86) | . | SHAM WT vs TAC WT | -44% (-321, 51) | >0.80 |
| 13-KODE | LA | Ketone | Alb | TAC | KO | 1.19 (0.81, 1.73) | . | SHAM KO - TAC KO | -49% (53, -373) | >0.80 |
| 5-KETE | AA | Ketone | Alb | SHAM | WT | 0.33 (0.12, 0.92) | 0.003 | WT SHAM vs KO SHAM | 42% (-90, 1898) | >0.80 |
| 5-KETE | AA | Ketone | Alb | SHAM | KO | 0.47 (0.16, 1.44) | . | TAC WT vs TAC KO | -18% (-88, 449) | >0.80 |
| 5-KETE | AA | Ketone | Alb | TAC | WT | 0.46 (0.22, 0.96) | . | SHAM WT vs TAC WT | 27% (-572, 92) | >0.80 |
| 5-KETE | AA | Ketone | Alb | TAC | KO | 0.37 (0.17, 0.82) | . | SHAM KO - TAC KO | -28% (88, -1295) | >0.80 |
| 12-KETE | AA | Ketone | Alb | SHAM | WT | 0.38 (0.16, 0.88) | 0.02 | WT SHAM vs KO SHAM | -54% (-95, 312) | >0.80 |
| 12-KETE | AA | Ketone | Alb | SHAM | KO | 0.17 (0.07, 0.44) | . | TAC WT vs TAC KO | 41% (-71, 587) | >0.80 |
| 12-KETE | AA | Ketone | Alb | TAC | WT | 0.19 (0.1, 0.35) | . | SHAM WT vs TAC WT | -102% (-1171, 68) | >0.80 |
| 12-KETE | AA | Ketone | Alb | TAC | KO | 0.27 (0.14, 0.51) | . | SHAM KO - TAC KO | 34% (91, -379) | >0.80 |
| 15-KETE | AA | Ketone | Alb | SHAM | WT | 0.16 (0.08, 0.33) | 0.02 | WT SHAM vs KO SHAM | 7% (-83, 566) | >0.80 |
| 15-KETE | AA | Ketone | Alb | SHAM | KO | 0.17 (0.08, 0.37) | . | TAC WT vs TAC KO | -37% (-83, 136) | >0.80 |
| 15-KETE | AA | Ketone | Alb | TAC | WT | 0.17 (0.1, 0.29) | . | SHAM WT vs TAC WT | 5% (-340, 80) | >0.80 |
| 15-KETE | AA | Ketone | Alb | TAC | KO | 0.11 (0.06, 0.19) | . | SHAM KO - TAC KO | -60% (69, -740) | >0.80 |
| 12-HpETE | AA | Peroxide | Alb | SHAM | WT | 1.31 (0.63, 2.74) | <0.0001 | WT SHAM vs KO SHAM | -97% (-100, -79) | <0.0001 |
| 12-HpETE | AA | Peroxide | Alb | SHAM | KO | 0.04 (0.02, 0.09) | . | TAC WT vs TAC KO | 53% (-62, 513) | >0.80 |
| 12-HpETE | AA | Peroxide | Alb | TAC | WT | 0.15 (0.09, 0.26) | . | SHAM WT vs TAC WT | -781% (-4305, -76) | 0.001 |
| 12-HpETE | AA | Peroxide | Alb | TAC | KO | 0.23 (0.13, 0.4) | . | SHAM KO - TAC KO | 82% (97, -1) | 0.05 |
| 15-HpETE | AA | Peroxide | Alb | SHAM | WT | 1.47 (0.74, 2.92) | 0.04 | WT SHAM vs KO SHAM | -93% (-99, -56) | 0.0003 |
| 15-HpETE | AA | Peroxide | Alb | SHAM | KO | 0.11 (0.05, 0.23) | . | TAC WT vs TAC KO | -83% (-95, -37) | 0.001 |
| 15-HpETE | AA | Peroxide | Alb | TAC | WT | 0.51 (0.31, 0.84) | . | SHAM WT vs TAC WT | -190% (-1197, 35) | 0.45 |
| 15-HpETE | AA | Peroxide | Alb | TAC | KO | 0.09 (0.05, 0.15) | . | SHAM KO - TAC KO | -21% (76, -511) | >0.80 |
| Resolvin D1 | DHA | Triol | Alb | SHAM | WT | 0.36 (0.18, 0.76) | 0.0009 | WT SHAM vs KO SHAM | -90% (-98, -31) | 0.007 |
| Resolvin D1 | DHA | Triol | Alb | SHAM | KO | 0.04 (0.02, 0.08) | . | TAC WT vs TAC KO | 31% (-67, 419) | >0.80 |
| Resolvin D1 | DHA | Triol | Alb | TAC | WT | 0.25 (0.15, 0.43) | . | SHAM WT vs TAC WT | -46% (-620, 70) | >0.80 |
| Resolvin D1 | DHA | Triol | Alb | TAC | KO | 0.33 (0.19, 0.58) | . | SHAM KO - TAC KO | 89% (98, 36) | 0.003 |
| 9(10)-EpOME | LA | Epoxide | Alb | SHAM | WT | 2.86 (1.76, 4.64) | 0.0005 | WT SHAM vs KO SHAM | 19% (-66, 323) | >0.80 |
| 9(10)-EpOME | LA | Epoxide | Alb | SHAM | KO | 3.42 (2.01, 5.8) | . | TAC WT vs TAC KO | 122% (-11, 454) | 0.15 |
| 9(10)-EpOME | LA | Epoxide | Alb | TAC | WT | 1.47 (1.03, 2.1) | . | SHAM WT vs TAC WT | -95% (-462, 32) | 0.65 |
| 9(10)-EpOME | LA | Epoxide | Alb | TAC | KO | 3.26 (2.24, 4.75) | . | SHAM KO - TAC KO | -5% (67, -228) | >0.80 |
| 12(13)-EpOME | LA | Epoxide | Alb | SHAM | WT | 2.69 (1.62, 4.46) | <0.0001 | WT SHAM vs KO SHAM | 25% (-67, 368) | >0.80 |
| 12(13)-EpOME | LA | Epoxide | Alb | SHAM | KO | 3.36 (1.93, 5.85) | . | TAC WT vs TAC KO | 15% (-56, 198) | >0.80 |
| 12(13)-EpOME | LA | Epoxide | Alb | TAC | WT | 2.52 (1.73, 3.66) | . | SHAM WT vs TAC WT | -7% (-222, 65) | >0.80 |
| 12(13)-EpOME | LA | Epoxide | Alb | TAC | KO | 2.9 (1.96, 4.29) | . | SHAM KO - TAC KO | -16% (65, -282) | >0.80 |
| 8(9)-EpETrE | AA | Epoxide | Alb | SHAM | WT | 1.34 (0.65, 2.76) | 0.21 | . | . | . |
| 8(9)-EpETrE | AA | Epoxide | Alb | SHAM | KO | 0.05 (0.02, 0.1) | . | . | . | . |
| 8(9)-EpETrE | AA | Epoxide | Alb | TAC | WT | 0.35 (0.21, 0.6) | . | . | . | . |
| 8(9)-EpETrE | AA | Epoxide | Alb | TAC | KO | 0.14 (0.08, 0.25) | . | . | . | . |
| 11(12)-EpETrE | AA | Epoxide | Alb | SHAM | WT | 1.28 (0.57, 2.89) | 0.09 | . | . | . |
| 11(12)-EpETrE | AA | Epoxide | Alb | SHAM | KO | 0.02 (0.01, 0.05) | . | . | . | . |
| 11(12)-EpETrE | AA | Epoxide | Alb | TAC | WT | 0.16 (0.09, 0.29) | . | . | . | . |
| 11(12)-EpETrE | AA | Epoxide | Alb | TAC | KO | 0.25 (0.13, 0.46) | . | . | . | . |
| 14(15)-EpETrE | AA | Epoxide | Alb | SHAM | WT | 1.37 (0.64, 2.95) | <0.0001 | WT SHAM vs KO SHAM | -91% (-99, -35) | 0.005 |
| 14(15)-EpETrE | AA | Epoxide | Alb | SHAM | KO | 0.12 (0.05, 0.28) | . | TAC WT vs TAC KO | -5% (-78, 299) | >0.80 |
| 14(15)-EpETrE | AA | Epoxide | Alb | TAC | WT | 0.16 (0.09, 0.28) | . | SHAM WT vs TAC WT | -753% (-4437, -60) | 0.002 |
| 14(15)-EpETrE | AA | Epoxide | Alb | TAC | KO | 0.15 (0.08, 0.28) | . | SHAM KO - TAC KO | 20% (87, -384) | >0.80 |
| 8(9)-EpETE | EPA | Epoxide | Alb | SHAM | WT | 0.38 (0.2, 0.7) | <0.0001 | WT SHAM vs KO SHAM | 50% (-70, 658) | >0.80 |
| 8(9)-EpETE | EPA | Epoxide | Alb | SHAM | KO | 0.57 (0.29, 1.12) | . | TAC WT vs TAC KO | 100% (-38, 544) | 0.73 |
| 8(9)-EpETE | EPA | Epoxide | Alb | TAC | WT | 0.38 (0.24, 0.6) | . | SHAM WT vs TAC WT | 0% (-290, 74) | >0.80 |
| 8(9)-EpETE | EPA | Epoxide | Alb | TAC | KO | 0.75 (0.47, 1.22) | . | SHAM KO - TAC KO | 25% (83, -226) | >0.80 |
| 11(12)-EpETE | EPA | Epoxide | Alb | SHAM | WT | 0.28 (0.1, 0.77) | 0.59 | . | . | . |
| 11(12)-EpETE | EPA | Epoxide | Alb | SHAM | KO | 0.42 (0.14, 1.27) | . | . | . | . |
| 11(12)-EpETE | EPA | Epoxide | Alb | TAC | WT | 0.13 (0.06, 0.28) | . | . | . | . |
| 11(12)-EpETE | EPA | Epoxide | Alb | TAC | KO | 0.44 (0.2, 0.96) | . | . | . | . |
| 14(15)-EpETE | EPA | Epoxide | Alb | SHAM | WT | 1.39 (0.95, 2.05) | <0.0001 | WT SHAM vs KO SHAM | -96% (-98, -88) | <0.0001 |
| 14(15)-EpETE | EPA | Epoxide | Alb | SHAM | KO | 0.06 (0.04, 0.09) | . | TAC WT vs TAC KO | 217% (53, 558) | <0.0001 |
| 14(15)-EpETE | EPA | Epoxide | Alb | TAC | WT | 0.41 (0.31, 0.55) | . | SHAM WT vs TAC WT | -237% (-688, -44) | 0.0004 |
| 14(15)-EpETE | EPA | Epoxide | Alb | TAC | KO | 1.31 (0.97, 1.77) | . | SHAM KO - TAC KO | 95% (98, 89) | <0.0001 |

| | | | | | | | | | | |
|---|---|---|---|---|---|---|---|---|---|---|
| 17(18)-EpETE | EPA | Epoxide | Alb | SHAM | WT | 0.63 (0.45, 0.87) | 0.01 | WT SHAM vs KO SHAM | -33% (-72, 57) | >0.80 |
| 17(18)-EpETE | EPA | Epoxide | Alb | SHAM | KO | 0.42 (0.29, 0.6) | . | TAC WT vs TAC KO | -29% (-62, 31) | >0.80 |
| 17(18)-EpETE | EPA | Epoxide | Alb | TAC | WT | 2.92 (2.3, 3.72) | . | SHAM WT vs TAC WT | 79% (56, 90) | <0.0001 |
| 17(18)-EpETE | EPA | Epoxide | Alb | TAC | KO | 2.08 (1.61, 2.68) | . | SHAM KO - TAC KO | 80% (91, 56) | <0.0001 |
| 7(8)-EpDPE | DHA | Epoxide | Alb | SHAM | WT | 0.07 (0.02, 0.2) | 0.1 | . | . | . |
| 7(8)-EpDPE | DHA | Epoxide | Alb | SHAM | KO | 0.03 (0.01, 0.09) | . | . | . | . |
| 7(8)-EpDPE | DHA | Epoxide | Alb | TAC | WT | 0.09 (0.04, 0.2) | . | . | . | . |
| 7(8)-EpDPE | DHA | Epoxide | Alb | TAC | KO | 0.09 (0.04, 0.2) | . | . | . | . |
| 10(11)-EpDPE | DHA | Epoxide | Alb | SHAM | WT | 0.11 (0.04, 0.27) | 0.04 | WT SHAM vs KO SHAM | -55% (-96, 395) | >0.80 |
| 10(11)-EpDPE | DHA | Epoxide | Alb | SHAM | KO | 0.05 (0.02, 0.13) | . | TAC WT vs TAC KO | -15% (-85, 383) | >0.80 |
| 10(11)-EpDPE | DHA | Epoxide | Alb | TAC | WT | 0.42 (0.21, 0.84) | . | SHAM WT vs TAC WT | 75% (-87, 97) | 0.5 |
| 10(11)-EpDPE | DHA | Epoxide | Alb | TAC | KO | 0.36 (0.18, 0.74) | . | SHAM KO - TAC KO | 87% (99, -15) | 0.09 |
| 13(14)-EpDPE | DHA | Epoxide | Alb | SHAM | WT | 0.31 (0.16, 0.58) | <0.0001 | WT SHAM vs KO SHAM | -93% (-99, -62) | <0.0001 |
| 13(14)-EpDPE | DHA | Epoxide | Alb | SHAM | KO | 0.02 (0.01, 0.04) | . | TAC WT vs TAC KO | 80% (-45, 492) | >0.80 |
| 13(14)-EpDPE | DHA | Epoxide | Alb | TAC | WT | 0.11 (0.07, 0.18) | . | SHAM WT vs TAC WT | -170% (-972, 32) | 0.43 |
| 13(14)-EpDPE | DHA | Epoxide | Alb | TAC | KO | 0.21 (0.13, 0.34) | . | SHAM KO - TAC KO | 89% (98, 52) | 0.0002 |
| 16(17)-EpDPE | DHA | Epoxide | Alb | SHAM | WT | 0.1 (0.03, 0.36) | 0.02 | WT SHAM vs KO SHAM | -71% (-99, 693) | >0.80 |
| 16(17)-EpDPE | DHA | Epoxide | Alb | SHAM | KO | 0.03 (0.01, 0.12) | . | TAC WT vs TAC KO | -12% (-92, 861) | >0.80 |
| 16(17)-EpDPE | DHA | Epoxide | Alb | TAC | WT | 0.13 (0.05, 0.33) | . | SHAM WT vs TAC WT | 20% (-1186, 95) | >0.80 |
| 16(17)-EpDPE | DHA | Epoxide | Alb | TAC | KO | 0.11 (0.04, 0.3) | . | SHAM KO - TAC KO | 74% (99, -426) | >0.80 |
| 19(20)-EpDPE | DHA | Epoxide | Alb | SHAM | WT | 0.32 (0.12, 0.85) | <0.0001 | WT SHAM vs KO SHAM | -94% (-100, -25) | 0.02 |
| 19(20)-EpDPE | DHA | Epoxide | Alb | SHAM | KO | 0.02 (0.01, 0.06) | . | TAC WT vs TAC KO | -35% (-89, 302) | >0.80 |
| 19(20)-EpDPE | DHA | Epoxide | Alb | TAC | WT | 0.08 (0.04, 0.17) | . | SHAM WT vs TAC WT | -291% (-3130, 53) | 0.61 |
| 19(20)-EpDPE | DHA | Epoxide | Alb | TAC | KO | 0.05 (0.03, 0.11) | . | SHAM KO - TAC KO | 64% (96, -252) | >0.80 |
| 9(10)-DiHOME | LA | Diol | Alb | SHAM | WT | 2.5 (1.65, 3.78) | <0.0001 | WT SHAM vs KO SHAM | 31% (-55, 284) | >0.80 |
| 9(10)-DiHOME | LA | Diol | Alb | SHAM | KO | 3.27 (2.08, 5.14) | . | TAC WT vs TAC KO | -14% (-61, 86) | >0.80 |
| 9(10)-DiHOME | LA | Diol | Alb | TAC | WT | 4.17 (3.07, 5.65) | . | SHAM WT vs TAC WT | 40% (-48, 76) | 0.79 |
| 9(10)-DiHOME | LA | Diol | Alb | TAC | KO | 3.57 (2.6, 4.92) | . | SHAM KO - TAC KO | 9% (65, -142) | >0.80 |
| 12(13)-DiHOME | LA | Diol | Alb | SHAM | WT | 5.67 (3.59, 8.94) | <0.0001 | WT SHAM vs KO SHAM | 373% (44, 1453) | 0.002 |
| 12(13)-DiHOME | LA | Diol | Alb | SHAM | KO | 26.8 (16.26, 44.17) | . | TAC WT vs TAC KO | 820% (290, 2070) | <0.0001 |
| 12(13)-DiHOME | LA | Diol | Alb | TAC | WT | 1.96 (1.4, 2.74) | . | SHAM WT vs TAC WT | -190% (-685, -7) | 0.03 |
| 12(13)-DiHOME | LA | Diol | Alb | TAC | KO | 18 (12.64, 25.64) | . | SHAM KO - TAC KO | -49% (49, -336) | >0.80 |
| 8(9)-DiHETrE | AA | Diol | Alb | SHAM | WT | 0.39 (0.21, 0.74) | 0.67 | . | . | . |
| 8(9)-DiHETrE | AA | Diol | Alb | SHAM | KO | 0.04 (0.02, 0.08) | . | . | . | . |
| 8(9)-DiHETrE | AA | Diol | Alb | TAC | WT | 0.41 (0.26, 0.66) | . | . | . | . |
| 8(9)-DiHETrE | AA | Diol | Alb | TAC | KO | 0.38 (0.23, 0.62) | . | . | . | . |
| 11(12)-DiHETrE | AA | Diol | Alb | SHAM | WT | 0.24 (0.15, 0.39) | 0.07 | . | . | . |
| 11(12)-DiHETrE | AA | Diol | Alb | SHAM | KO | 0.11 (0.07, 0.19) | . | . | . | . |
| 11(12)-DiHETrE | AA | Diol | Alb | TAC | WT | 0.49 (0.35, 0.69) | . | . | . | . |
| 11(12)-DiHETrE | AA | Diol | Alb | TAC | KO | 0.42 (0.3, 0.61) | . | . | . | . |
| 14(15)-DiHETrE | AA | Diol | Alb | SHAM | WT | 0.38 (0.18, 0.79) | <0.0001 | WT SHAM vs KO SHAM | -96% (-99, -76) | <0.0001 |
| 14(15)-DiHETrE | AA | Diol | Alb | SHAM | KO | 0.01 (0.01, 0.03) | . | TAC WT vs TAC KO | -32% (-83, 171) | >0.80 |
| 14(15)-DiHETrE | AA | Diol | Alb | TAC | WT | 0.39 (0.23, 0.67) | . | SHAM WT vs TAC WT | 4% (-380, 81) | >0.80 |
| 14(15)-DiHETrE | AA | Diol | Alb | TAC | KO | 0.27 (0.15, 0.47) | . | SHAM KO - TAC KO | 95% (99, 72) | <0.0001 |
| 8(9)-DiHETE | EPA | Diol | Alb | SHAM | WT | 1.04 (0.29, 3.73) | <0.0001 | WT SHAM vs KO SHAM | -80% (-99, 468) | >0.80 |
| 8(9)-DiHETE | EPA | Diol | Alb | SHAM | KO | 0.21 (0.05, 0.86) | . | TAC WT vs TAC KO | -97% (-100, -63) | 0.0005 |
| 8(9)-DiHETE | EPA | Diol | Alb | TAC | WT | 1.88 (0.73, 4.82) | . | SHAM WT vs TAC WT | 45% (-799, 97) | >0.80 |
| 8(9)-DiHETE | EPA | Diol | Alb | TAC | KO | 0.06 (0.02, 0.17) | . | SHAM KO - TAC KO | -238% (83, -6730) | >0.80 |
| 11(12)-DiHETE | EPA | Diol | Alb | SHAM | WT | 0.24 (0.07, 0.75) | 0.007 | WT SHAM vs KO SHAM | -70% (-99, 512) | >0.80 |
| 11(12)-DiHETE | EPA | Diol | Alb | SHAM | KO | 0.07 (0.02, 0.25) | . | TAC WT vs TAC KO | -85% (-98, 28) | 0.14 |
| 11(12)-DiHETE | EPA | Diol | Alb | TAC | WT | 0.46 (0.19, 1.07) | . | SHAM WT vs TAC WT | 48% (-545, 96) | >0.80 |
| 11(12)-DiHETE | EPA | Diol | Alb | TAC | KO | 0.07 (0.03, 0.16) | . | SHAM KO - TAC KO | -7% (93, -1530) | >0.80 |
| 14(15)-DiHETE | EPA | Diol | Alb | SHAM | WT | 1.6 (0.78, 3.28) | 0.06 | . | . | . |
| 14(15)-DiHETE | EPA | Diol | Alb | SHAM | KO | 0.07 (0.03, 0.16) | . | . | . | . |
| 14(15)-DiHETE | EPA | Diol | Alb | TAC | WT | 0.52 (0.3, 0.88) | . | . | . | . |
| 14(15)-DiHETE | EPA | Diol | Alb | TAC | KO | 0.07 (0.04, 0.13) | . | . | . | . |
| 17(18)-DiHETE | EPA | Diol | Alb | SHAM | WT | 0.22 (0.08, 0.58) | 0.29 | . | . | . |
| 17(18)-DiHETE | EPA | Diol | Alb | SHAM | KO | 0.06 (0.02, 0.19) | . | . | . | . |
| 17(18)-DiHETE | EPA | Diol | Alb | TAC | WT | 0.37 (0.18, 0.76) | . | . | . | . |
| 17(18)-DiHETE | EPA | Diol | Alb | TAC | KO | 0.21 (0.1, 0.44) | . | . | . | . |
| 9-HODE | LA | Alcohol | HDL | SHAM | WT | 50.1 (42.5, 59) | <0.0001 | WT SHAM vs KO SHAM | -75% (-83, -61) | <0.0001 |
| 9-HODE | LA | Alcohol | HDL | SHAM | KO | 12.7 (10.6, 15.1) | . | TAC WT vs TAC KO | -71% (-78, -60) | <0.0001 |
| 9-HODE | LA | Alcohol | HDL | TAC | WT | 35.4 (31.4, 39.9) | . | SHAM WT vs TAC WT | -42% (-102, 1) | 0.06 |
| 9-HODE | LA | Alcohol | HDL | TAC | KO | 10.3 (9.1, 11.7) | . | SHAM KO - TAC KO | -22% (17, -80) | >0.80 |
| 13-HODE | LA | Alcohol | HDL | SHAM | WT | 47.6 (37.7, 60.2) | 0.0003 | WT SHAM vs KO SHAM | -80% (-89, -63) | <0.0001 |
| 13-HODE | LA | Alcohol | HDL | SHAM | KO | 9.5 (7.4, 12.3) | . | TAC WT vs TAC KO | -49% (-67, -21) | 0.0001 |
| 13-HODE | LA | Alcohol | HDL | TAC | WT | 19.4 (16.3, 23) | . | SHAM WT vs TAC WT | -146% (-310, -48) | <0.0001 |
| 13-HODE | LA | Alcohol | HDL | TAC | KO | 9.9 (8.2, 11.8) | . | SHAM KO - TAC KO | 3% (44, -68) | >0.80 |
| 9-HOTrE | aLA | Alcohol | HDL | SHAM | WT | 1.05 (0.61, 1.81) | <0.0001 | WT SHAM vs KO SHAM | -96% (-99, -83) | <0.0001 |
| 9-HOTrE | aLA | Alcohol | HDL | SHAM | KO | 0.04 (0.02, 0.08) | . | TAC WT vs TAC KO | 398% (79, 1286) | <0.0001 |
| 9-HOTrE | aLA | Alcohol | HDL | TAC | WT | 0.05 (0.03, 0.07) | . | SHAM WT vs TAC WT | -2183% (-7393, -595) | <0.0001 |
| 9-HOTrE | aLA | Alcohol | HDL | TAC | KO | 0.23 (0.15, 0.35) | . | SHAM KO - TAC KO | 81% (95, 32) | 0.002 |
| 13-HOTrE | aLA | Alcohol | HDL | SHAM | WT | 1.05 (0.63, 1.75) | 0.0002 | WT SHAM vs KO SHAM | -82% (-95, -33) | 0.002 |
| 13-HOTrE | aLA | Alcohol | HDL | SHAM | KO | 0.19 (0.11, 0.33) | . | TAC WT vs TAC KO | -3% (-63, 155) | >0.80 |
| 13-HOTrE | aLA | Alcohol | HDL | TAC | WT | 0.45 (0.31, 0.66) | . | SHAM WT vs TAC WT | -131% (-608, 24) | 0.36 |
| 13-HOTrE | aLA | Alcohol | HDL | TAC | KO | 0.44 (0.3, 0.66) | . | SHAM KO - TAC KO | 58% (87, -41) | 0.43 |
| 5-HETE | AA | Alcohol | HDL | SHAM | WT | 11.6 (9.8, 13.8) | <0.0001 | WT SHAM vs KO SHAM | 84% (18, 187) | 0.001 |
| 5-HETE | AA | Alcohol | HDL | SHAM | KO | 21.3 (17.7, 25.7) | . | TAC WT vs TAC KO | -27% (-47, 1) | 0.06 |
| 5-HETE | AA | Alcohol | HDL | TAC | WT | 25.3 (22.3, 28.7) | . | SHAM WT vs TAC WT | 54% (33, 68) | <0.0001 |
| 5-HETE | AA | Alcohol | HDL | TAC | KO | 18.4 (16.2, 21) | . | SHAM KO - TAC KO | -16% (23, -73) | >0.80 |
| 9-HETE | AA | Alcohol | HDL | SHAM | WT | 0.95 (0.61, 1.48) | 0.01 | WT SHAM vs KO SHAM | 337% (37, 1286) | 0.003 |
| 9-HETE | AA | Alcohol | HDL | SHAM | KO | 4.15 (2.55, 6.74) | . | TAC WT vs TAC KO | 106% (-11, 373) | 0.16 |
| 9-HETE | AA | Alcohol | HDL | TAC | WT | 0.82 (0.59, 1.13) | . | SHAM WT vs TAC WT | -16% (-207, 56) | >0.80 |
| 9-HETE | AA | Alcohol | HDL | TAC | KO | 1.68 (1.19, 2.36) | . | SHAM KO - TAC KO | -147% (13, -603) | 0.16 |
| 12-HETE | AA | Alcohol | HDL | SHAM | WT | 2.8 (2.1, 3.6) | <0.0001 | WT SHAM vs KO SHAM | 183% (42, 463) | 0.0002 |
| 12-HETE | AA | Alcohol | HDL | SHAM | KO | 7.9 (5.9, 10.5) | . | TAC WT vs TAC KO | -86% (-91, -77) | <0.0001 |

| | | | | | | | | | | |
|---|---|---|---|---|---|---|---|---|---|---|
| 12-HETE | AA | Alcohol | HDL | TAC | WT | 13.9 (11.5, 16.9) | . | SHAM WT vs TAC WT | 80% (64, 89) | <0.0001 |
| 12-HETE | AA | Alcohol | HDL | TAC | KO | 2 (1.6, 2.4) | . | SHAM KO - TAC KO | -302% (-116, -649) | <0.0001 |
| 15-HETE | AA | Alcohol | HDL | SHAM | WT | 0.84 (0.67, 1.07) | <0.0001 | WT SHAM vs KO SHAM | 657% (309, 1300) | <0.0001 |
| 15-HETE | AA | Alcohol | HDL | SHAM | KO | 6.4 (4.94, 8.28) | . | TAC WT vs TAC KO | -75% (-84, -61) | <0.0001 |
| 15-HETE | AA | Alcohol | HDL | TAC | WT | 11.27 (9.47, 13.41) | . | SHAM WT vs TAC WT | 93% (87, 96) | <0.0001 |
| 15-HETE | AA | Alcohol | HDL | TAC | KO | 2.82 (2.35, 3.39) | . | SHAM KO - TAC KO | -126% (-30, -295) | 0.0003 |
| 5-HEPE | EPA | Alcohol | HDL | SHAM | WT | 0.29 (0.19, 0.44) | <0.0001 | WT SHAM vs KO SHAM | -39% (-79, 77) | >0.80 |
| 5-HEPE | EPA | Alcohol | HDL | SHAM | KO | 0.18 (0.11, 0.28) | . | TAC WT vs TAC KO | 165% (22, 474) | 0.003 |
| 5-HEPE | EPA | Alcohol | HDL | TAC | WT | 0.1 (0.07, 0.13) | . | SHAM WT vs TAC WT | -201% (-639, -23) | 0.005 |
| 5-HEPE | EPA | Alcohol | HDL | TAC | KO | 0.26 (0.19, 0.35) | . | SHAM KO - TAC KO | 31% (74, -82) | >0.80 |
| 8-HEPE | EPA | Alcohol | HDL | SHAM | WT | 1.08 (0.75, 1.54) | <0.0001 | WT SHAM vs KO SHAM | -70% (-88, -24) | 0.002 |
| 8-HEPE | EPA | Alcohol | HDL | SHAM | KO | 0.32 (0.22, 0.48) | . | TAC WT vs TAC KO | 3% (-47, 103) | >0.80 |
| 8-HEPE | EPA | Alcohol | HDL | TAC | WT | 0.4 (0.31, 0.53) | . | SHAM WT vs TAC WT | -166% (-482, -22) | 0.004 |
| 8-HEPE | EPA | Alcohol | HDL | TAC | KO | 0.42 (0.32, 0.55) | . | SHAM KO - TAC KO | 23% (67, -79) | >0.80 |
| 9-HEPE | EPA | Alcohol | HDL | SHAM | WT | 0.89 (0.71, 1.11) | 0.1 | . | . | . |
| 9-HEPE | EPA | Alcohol | HDL | SHAM | KO | 0.15 (0.12, 0.2) | . | . | . | . |
| 9-HEPE | EPA | Alcohol | HDL | TAC | WT | 0.39 (0.33, 0.47) | . | . | . | . |
| 9-HEPE | EPA | Alcohol | HDL | TAC | KO | 0.57 (0.48, 0.68) | . | . | . | . |
| 11-HEPE | EPA | Alcohol | HDL | SHAM | WT | 0.13 (0.07, 0.26) | 0.02 | WT SHAM vs KO SHAM | -69% (-95, 74) | 0.52 |
| 11-HEPE | EPA | Alcohol | HDL | SHAM | KO | 0.04 (0.02, 0.08) | . | TAC WT vs TAC KO | 574% (93, 2248) | 0.0001 |
| 11-HEPE | EPA | Alcohol | HDL | TAC | WT | 0.03 (0.02, 0.05) | . | SHAM WT vs TAC WT | -343% (-1789, -4) | 0.04 |
| 11-HEPE | EPA | Alcohol | HDL | TAC | KO | 0.2 (0.12, 0.34) | . | SHAM KO - TAC KO | 80% (96, 3) | 0.04 |
| 12-HEPE | EPA | Alcohol | HDL | SHAM | WT | 0.29 (0.17, 0.51) | <0.0001 | WT SHAM vs KO SHAM | -55% (-89, 89) | >0.80 |
| 12-HEPE | EPA | Alcohol | HDL | SHAM | KO | 0.13 (0.07, 0.24) | . | TAC WT vs TAC KO | -45% (-81, 54) | 0.76 |
| 12-HEPE | EPA | Alcohol | HDL | TAC | WT | 0.18 (0.12, 0.28) | . | SHAM WT vs TAC WT | -61% (-435, 52) | >0.80 |
| 12-HEPE | EPA | Alcohol | HDL | TAC | KO | 0.1 (0.07, 0.15) | . | SHAM KO - TAC KO | -32% (64, -383) | >0.80 |
| 15-HEPE | EPA | Alcohol | HDL | SHAM | WT | 0.05 (0.03, 0.07) | 0.24 | . | . | . |
| 15-HEPE | EPA | Alcohol | HDL | SHAM | KO | 0.15 (0.1, 0.23) | . | . | . | . |
| 15-HEPE | EPA | Alcohol | HDL | TAC | WT | 0.25 (0.19, 0.33) | . | . | . | . |
| 15-HEPE | EPA | Alcohol | HDL | TAC | KO | 2.12 (1.57, 2.87) | . | . | . | . |
| 18-HEPE | EPA | Alcohol | HDL | SHAM | WT | 2.42 (2.01, 2.92) | <0.0001 | WT SHAM vs KO SHAM | -98% (-99, -97) | <0.0001 |
| 18-HEPE | EPA | Alcohol | HDL | SHAM | KO | 0.05 (0.04, 0.06) | . | TAC WT vs TAC KO | -64% (-75, -49) | <0.0001 |
| 18-HEPE | EPA | Alcohol | HDL | TAC | WT | 3.42 (2.97, 3.92) | . | SHAM WT vs TAC WT | 29% (-7, 53) | 0.2 |
| 18-HEPE | EPA | Alcohol | HDL | TAC | KO | 1.22 (1.06, 1.41) | . | SHAM KO - TAC KO | 96% (97, 94) | <0.0001 |
| 4-HDoHE | DHA | Alcohol | HDL | SHAM | WT | 0.86 (0.39, 1.92) | 0.01 | WT SHAM vs KO SHAM | 110% (-74, 1603) | >0.80 |
| 4-HDoHE | DHA | Alcohol | HDL | SHAM | KO | 1.81 (0.75, 4.37) | . | TAC WT vs TAC KO | 24% (-73, 461) | >0.80 |
| 4-HDoHE | DHA | Alcohol | HDL | TAC | WT | 0.28 (0.15, 0.51) | . | SHAM WT vs TAC WT | -207% (-1669, 47) | 0.62 |
| 4-HDoHE | DHA | Alcohol | HDL | TAC | KO | 0.35 (0.19, 0.65) | . | SHAM KO - TAC KO | -420% (21, -3349) | 0.15 |
| 7-HDoHE | DHA | Alcohol | HDL | SHAM | WT | 0.99 (0.44, 2.24) | 0.001 | WT SHAM vs KO SHAM | -91% (-99, -26) | 0.01 |
| 7-HDoHE | DHA | Alcohol | HDL | SHAM | KO | 0.09 (0.04, 0.22) | . | TAC WT vs TAC KO | -58% (-91, 94) | 0.8 |
| 7-HDoHE | DHA | Alcohol | HDL | TAC | WT | 0.42 (0.23, 0.77) | . | SHAM WT vs TAC WT | -134% (-1273, 60) | >0.80 |
| 7-HDoHE | DHA | Alcohol | HDL | TAC | KO | 0.18 (0.1, 0.34) | . | SHAM KO - TAC KO | 51% (93, -232) | >0.80 |
| 8-HDoHE | DHA | Alcohol | HDL | SHAM | WT | 1 (0.5, 2.01) | 0.0002 | WT SHAM vs KO SHAM | 121% (-64, 1257) | >0.80 |
| 8-HDoHE | DHA | Alcohol | HDL | SHAM | KO | 2.21 (1.03, 4.75) | . | TAC WT vs TAC KO | -74% (-93, -3) | 0.04 |
| 8-HDoHE | DHA | Alcohol | HDL | TAC | WT | 0.83 (0.5, 1.4) | . | SHAM WT vs TAC WT | -20% (-449, 74) | >0.80 |
| 8-HDoHE | DHA | Alcohol | HDL | TAC | KO | 0.22 (0.13, 0.37) | . | SHAM KO - TAC KO | -917% (-97, -5151) | 0.0005 |
| 10-HDoHE | DHA | Alcohol | HDL | SHAM | WT | 0.25 (0.11, 0.56) | 0.02 | WT SHAM vs KO SHAM | -59% (-95, 248) | >0.80 |
| 10-HDoHE | DHA | Alcohol | HDL | SHAM | KO | 0.1 (0.04, 0.25) | . | TAC WT vs TAC KO | -60% (-91, 85) | 0.72 |
| 10-HDoHE | DHA | Alcohol | HDL | TAC | WT | 0.27 (0.15, 0.5) | . | SHAM WT vs TAC WT | 9% (-442, 85) | >0.80 |
| 10-HDoHE | DHA | Alcohol | HDL | TAC | KO | 0.11 (0.06, 0.21) | . | SHAM KO - TAC KO | 5% (86, -550) | >0.80 |
| 11-HDoHE | DHA | Alcohol | HDL | SHAM | WT | 0.31 (0.18, 0.54) | 0.24 | . | . | . |
| 11-HDoHE | DHA | Alcohol | HDL | SHAM | KO | 0.21 (0.11, 0.39) | . | . | . | . |
| 11-HDoHE | DHA | Alcohol | HDL | TAC | WT | 0.3 (0.2, 0.45) | . | . | . | . |
| 11-HDoHE | DHA | Alcohol | HDL | TAC | KO | 0.28 (0.18, 0.43) | . | . | . | . |
| 13-HDoHE | DHA | Alcohol | HDL | SHAM | WT | 0.37 (0.17, 0.79) | 0.0009 | WT SHAM vs KO SHAM | 85% (-74, 1235) | >0.80 |
| 13-HDoHE | DHA | Alcohol | HDL | SHAM | KO | 0.69 (0.3, 1.58) | . | TAC WT vs TAC KO | -53% (-89, 96) | >0.80 |
| 13-HDoHE | DHA | Alcohol | HDL | TAC | WT | 0.52 (0.3, 0.92) | . | SHAM WT vs TAC WT | 29% (-271, 86) | >0.80 |
| 13-HDoHE | DHA | Alcohol | HDL | TAC | KO | 0.25 (0.14, 0.45) | . | SHAM KO - TAC KO | -179% (53, -1561) | 0.78 |
| 14-HDoHE | DHA | Alcohol | HDL | SHAM | WT | 0.39 (0.24, 0.64) | <0.0001 | WT SHAM vs KO SHAM | 25% (-66, 355) | >0.80 |
| 14-HDoHE | DHA | Alcohol | HDL | SHAM | KO | 0.49 (0.28, 0.84) | . | TAC WT vs TAC KO | -71% (-89, -27) | 0.001 |
| 14-HDoHE | DHA | Alcohol | HDL | TAC | WT | 0.62 (0.43, 0.89) | . | SHAM WT vs TAC WT | 37% (-88, 79) | >0.80 |
| 14-HDoHE | DHA | Alcohol | HDL | TAC | KO | 0.18 (0.12, 0.26) | . | SHAM KO - TAC KO | -177% (14, -794) | 0.16 |
| 16-HDoHE | DHA | Alcohol | HDL | SHAM | WT | 0.06 (0.03, 0.13) | 0.13 | . | . | . |
| 16-HDoHE | DHA | Alcohol | HDL | SHAM | KO | 0.05 (0.02, 0.11) | . | . | . | . |
| 16-HDoHE | DHA | Alcohol | HDL | TAC | WT | 0.05 (0.03, 0.1) | . | . | . | . |
| 16-HDoHE | DHA | Alcohol | HDL | TAC | KO | 0.06 (0.03, 0.1) | . | . | . | . |
| 17-HDoHE | DHA | Alcohol | HDL | SHAM | WT | 0.94 (0.53, 1.67) | <0.0001 | WT SHAM vs KO SHAM | 148% (-44, 998) | 0.69 |
| 17-HDoHE | DHA | Alcohol | HDL | SHAM | KO | 2.34 (1.25, 4.37) | . | TAC WT vs TAC KO | -53% (-84, 38) | 0.48 |
| 17-HDoHE | DHA | Alcohol | HDL | TAC | WT | 1.36 (0.89, 2.07) | . | SHAM WT vs TAC WT | 31% (-142, 80) | >0.80 |
| 17-HDoHE | DHA | Alcohol | HDL | TAC | KO | 0.64 (0.41, 1) | . | SHAM KO - TAC KO | -265% (5, -1301) | 0.07 |
| 20-HDoHE | DHA | Alcohol | HDL | SHAM | WT | 2.16 (1.44, 3.25) | 0.61 | . | . | . |
| 20-HDoHE | DHA | Alcohol | HDL | SHAM | KO | 0.74 (0.48, 1.16) | . | . | . | . |
| 20-HDoHE | DHA | Alcohol | HDL | TAC | WT | 0.43 (0.32, 0.58) | . | . | . | . |
| 20-HDoHE | DHA | Alcohol | HDL | TAC | KO | 0.27 (0.2, 0.37) | . | . | . | . |
| 22-HDoHE | DHA | Alcohol | HDL | SHAM | WT | 0.31 (0.16, 0.59) | 0.51 | . | . | . |
| 22-HDoHE | DHA | Alcohol | HDL | SHAM | KO | 0.18 (0.09, 0.37) | . | . | . | . |
| 22-HDoHE | DHA | Alcohol | HDL | TAC | WT | 0.07 (0.04, 0.11) | . | . | . | . |
| 22-HDoHE | DHA | Alcohol | HDL | TAC | KO | 0.17 (0.11, 0.29) | . | . | . | . |
| 9-KODE | LA | Ketone | HDL | SHAM | WT | 4.03 (3.09, 5.26) | <0.0001 | WT SHAM vs KO SHAM | 47% (-27, 193) | >0.80 |
| 9-KODE | LA | Ketone | HDL | SHAM | KO | 5.91 (4.42, 7.91) | . | TAC WT vs TAC KO | -88% (-93, -80) | <0.0001 |
| 9-KODE | LA | Ketone | HDL | TAC | WT | 14.28 (11.73, 17.37) | . | SHAM WT vs TAC WT | 72% (50, 84) | <0.0001 |
| 9-KODE | LA | Ketone | HDL | TAC | KO | 1.71 (1.39, 2.1) | . | SHAM KO - TAC KO | -246% (-85, -547) | <0.0001 |
| 13-KODE | LA | Ketone | HDL | SHAM | WT | 2.69 (1.74, 4.18) | <0.0001 | WT SHAM vs KO SHAM | 269% (17, 1059) | 0.01 |
| 13-KODE | LA | Ketone | HDL | SHAM | KO | 9.94 (6.14, 16.08) | . | TAC WT vs TAC KO | -30% (-69, 59) | >0.80 |
| 13-KODE | LA | Ketone | HDL | TAC | WT | 4.63 (3.35, 6.4) | . | SHAM WT vs TAC WT | 42% (-52, 78) | 0.8 |
| 13-KODE | LA | Ketone | HDL | TAC | KO | 3.23 (2.3, 4.53) | . | SHAM KO - TAC KO | -208% (-9, -768) | 0.02 |

| | | | | | | | | | | |
|---|---|---|---|---|---|---|---|---|---|---|
| 5-KETE | AA | Ketone | HDL | SHAM | WT | 1.18 (0.88, 1.58) | 0.003 | WT SHAM vs KO SHAM | -58% (-81, -11) | 0.01 |
| 5-KETE | AA | Ketone | HDL | SHAM | KO | 0.49 (0.36, 0.67) | . | TAC WT vs TAC KO | -66% (-80, -40) | <0.0001 |
| 5-KETE | AA | Ketone | HDL | TAC | WT | 3.58 (2.89, 4.44) | . | SHAM WT vs TAC WT | 67% (38, 83) | <0.0001 |
| 5-KETE | AA | Ketone | HDL | TAC | KO | 1.24 (0.99, 1.55) | . | SHAM KO - TAC KO | 60% (80, 21) | 0.001 |
| 12-KETE | AA | Ketone | HDL | SHAM | WT | 1.47 (1.08, 1.99) | 0.02 | WT SHAM vs KO SHAM | -80% (-91, -56) | <0.0001 |
| 12-KETE | AA | Ketone | HDL | SHAM | KO | 0.29 (0.21, 0.41) | . | TAC WT vs TAC KO | -40% (-66, 6) | 0.12 |
| 12-KETE | AA | Ketone | HDL | TAC | WT | 1.37 (1.1, 1.72) | . | SHAM WT vs TAC WT | -7% (-108, 45) | >0.80 |
| 12-KETE | AA | Ketone | HDL | TAC | KO | 0.82 (0.65, 1.04) | . | SHAM KO - TAC KO | 64% (83, 27) | 0.0004 |
| 15-KETE | AA | Ketone | HDL | SHAM | WT | 1.96 (1.41, 2.72) | 0.02 | WT SHAM vs KO SHAM | -78% (-91, -48) | <0.0001 |
| 15-KETE | AA | Ketone | HDL | SHAM | KO | 0.43 (0.3, 0.62) | . | TAC WT vs TAC KO | 3% (-45, 91) | >0.80 |
| 15-KETE | AA | Ketone | HDL | TAC | WT | 0.46 (0.36, 0.59) | . | SHAM WT vs TAC WT | -327% (-776, -108) | <0.0001 |
| 15-KETE | AA | Ketone | HDL | TAC | KO | 0.47 (0.37, 0.61) | . | SHAM KO - TAC KO | 9% (58, -97) | >0.80 |
| 12-HpETE | AA | Peroxide | HDL | SHAM | WT | 1.6 (0.87, 2.93) | <0.0001 | WT SHAM vs KO SHAM | -49% (-89, 147) | >0.80 |
| 12-HpETE | AA | Peroxide | HDL | SHAM | KO | 0.82 (0.42, 1.59) | . | TAC WT vs TAC KO | -22% (-75, 142) | >0.80 |
| 12-HpETE | AA | Peroxide | HDL | TAC | WT | 0.4 (0.26, 0.63) | . | SHAM WT vs TAC WT | -297% (-1387, -6) | 0.03 |
| 12-HpETE | AA | Peroxide | HDL | TAC | KO | 0.31 (0.2, 0.5) | . | SHAM KO - TAC KO | -162% (37, -988) | 0.54 |
| 15-HpETE | AA | Peroxide | HDL | SHAM | WT | 4.33 (3.12, 6.01) | 0.04 | WT SHAM vs KO SHAM | -55% (-81, 6) | 0.09 |
| 15-HpETE | AA | Peroxide | HDL | SHAM | KO | 1.95 (1.36, 2.8) | . | TAC WT vs TAC KO | 112% (14, 292) | 0.005 |
| 15-HpETE | AA | Peroxide | HDL | TAC | WT | 2.07 (1.63, 2.64) | . | SHAM WT vs TAC WT | -109% (-327, -2) | 0.04 |
| 15-HpETE | AA | Peroxide | HDL | TAC | KO | 4.39 (3.4, 5.66) | . | SHAM KO - TAC KO | 56% (79, 4) | 0.03 |
| Resolvin D1 | DHA | Triol | HDL | SHAM | WT | 0.79 (0.42, 1.49) | 0.0009 | WT SHAM vs KO SHAM | 126% (-56, 1070) | >0.80 |
| Resolvin D1 | DHA | Triol | HDL | SHAM | KO | 1.8 (0.9, 3.58) | . | TAC WT vs TAC KO | -13% (-73, 184) | >0.80 |
| Resolvin D1 | DHA | Triol | HDL | TAC | WT | 3.02 (1.9, 4.81) | . | SHAM WT vs TAC WT | 74% (-4, 93) | 0.07 |
| Resolvin D1 | DHA | Triol | HDL | TAC | KO | 2.62 (1.61, 4.27) | . | SHAM KO - TAC KO | 31% (84, -203) | >0.80 |
| 9(10)-EpOME | LA | Epoxide | HDL | SHAM | WT | 15.9 (12.45, 20.32) | 0.0005 | WT SHAM vs KO SHAM | -80% (-89, -62) | <0.0001 |
| 9(10)-EpOME | LA | Epoxide | HDL | SHAM | KO | 3.17 (2.43, 4.15) | . | TAC WT vs TAC KO | -8% (-42, 45) | >0.80 |
| 9(10)-EpOME | LA | Epoxide | HDL | TAC | WT | 5.41 (4.51, 6.48) | . | SHAM WT vs TAC WT | -194% (-402, -72) | <0.0001 |
| 9(10)-EpOME | LA | Epoxide | HDL | TAC | KO | 4.95 (4.1, 5.99) | . | SHAM KO - TAC KO | 36% (64, -14) | 0.32 |
| 12(13)-EpOME | LA | Epoxide | HDL | SHAM | WT | 14.11 (11.06, 18.01) | <0.0001 | WT SHAM vs KO SHAM | -80% (-90, -63) | <0.0001 |
| 12(13)-EpOME | LA | Epoxide | HDL | SHAM | KO | 2.76 (2.11, 3.6) | . | TAC WT vs TAC KO | -54% (-71, -27) | <0.0001 |
| 12(13)-EpOME | LA | Epoxide | HDL | TAC | WT | 9.68 (8.09, 11.6) | . | SHAM WT vs TAC WT | -46% (-148, 14) | 0.45 |
| 12(13)-EpOME | LA | Epoxide | HDL | TAC | KO | 4.5 (3.73, 5.44) | . | SHAM KO - TAC KO | 39% (65, -9) | 0.18 |
| 8(9)-EpETrE | AA | Epoxide | HDL | SHAM | WT | 9.19 (6.97, 12.11) | 0.21 | . | . | . |
| 8(9)-EpETrE | AA | Epoxide | HDL | SHAM | KO | 0.38 (0.28, 0.51) | . | . | . | . |
| 8(9)-EpETrE | AA | Epoxide | HDL | TAC | WT | 2.62 (2.14, 3.21) | . | . | . | . |
| 8(9)-EpETrE | AA | Epoxide | HDL | TAC | KO | 2.57 (2.08, 3.19) | . | . | . | . |
| 11(12)-EpETrE | AA | Epoxide | HDL | SHAM | WT | 1.03 (0.53, 2) | 0.09 | . | . | . |
| 11(12)-EpETrE | AA | Epoxide | HDL | SHAM | KO | 0.22 (0.11, 0.46) | . | . | . | . |
| 11(12)-EpETrE | AA | Epoxide | HDL | TAC | WT | 0.28 (0.17, 0.46) | . | . | . | . |
| 11(12)-EpETrE | AA | Epoxide | HDL | TAC | KO | 0.96 (0.58, 1.6) | . | . | . | . |
| 14(15)-EpETrE | AA | Epoxide | HDL | SHAM | WT | 2 (1.06, 3.76) | <0.0001 | WT SHAM vs KO SHAM | 7% (-79, 459) | >0.80 |
| 14(15)-EpETrE | AA | Epoxide | HDL | SHAM | KO | 2.15 (1.07, 4.29) | . | TAC WT vs TAC KO | -84% (-95, -47) | 0.0001 |
| 14(15)-EpETrE | AA | Epoxide | HDL | TAC | WT | 0.74 (0.46, 1.17) | . | SHAM WT vs TAC WT | -172% (-985, 32) | 0.42 |
| 14(15)-EpETrE | AA | Epoxide | HDL | TAC | KO | 0.12 (0.07, 0.19) | . | SHAM KO - TAC KO | -1716% (-308, -7981) | <0.0001 |
| 8(9)-EpETE | EPA | Epoxide | HDL | SHAM | WT | 3.37 (2.46, 4.62) | <0.0001 | WT SHAM vs KO SHAM | -9% (-60, 106) | >0.80 |
| 8(9)-EpETE | EPA | Epoxide | HDL | SHAM | KO | 3.06 (2.16, 4.32) | . | TAC WT vs TAC KO | -43% (-68, 4) | 0.09 |
| 8(9)-EpETE | EPA | Epoxide | HDL | TAC | WT | 3.32 (2.63, 4.19) | . | SHAM WT vs TAC WT | -2% (-102, 49) | >0.80 |
| 8(9)-EpETE | EPA | Epoxide | HDL | TAC | KO | 1.9 (1.49, 2.42) | . | SHAM KO - TAC KO | -61% (23, -238) | 0.62 |
| 11(12)-EpETE | EPA | Epoxide | HDL | SHAM | WT | 1.81 (1.27, 2.57) | 0.59 | . | . | . |
| 11(12)-EpETE | EPA | Epoxide | HDL | SHAM | KO | 0.52 (0.35, 0.76) | . | . | . | . |
| 11(12)-EpETE | EPA | Epoxide | HDL | TAC | WT | 1.4 (1.08, 1.82) | . | . | . | . |
| 11(12)-EpETE | EPA | Epoxide | HDL | TAC | KO | 0.44 (0.34, 0.58) | . | . | . | . |
| 14(15)-EpETE | EPA | Epoxide | HDL | SHAM | WT | 12.79 (10.59, 15.45) | <0.0001 | WT SHAM vs KO SHAM | -87% (-92, -79) | <0.0001 |
| 14(15)-EpETE | EPA | Epoxide | HDL | SHAM | KO | 1.62 (1.32, 1.99) | . | TAC WT vs TAC KO | -74% (-82, -63) | <0.0001 |
| 14(15)-EpETE | EPA | Epoxide | HDL | TAC | WT | 10.31 (8.97, 11.85) | . | SHAM WT vs TAC WT | -24% (-87, 18) | >0.80 |
| 14(15)-EpETE | EPA | Epoxide | HDL | TAC | KO | 2.65 (2.29, 3.06) | . | SHAM KO - TAC KO | 39% (61, 4) | 0.02 |
| 17(18)-EpETE | EPA | Epoxide | HDL | SHAM | WT | 11.33 (8.8, 14.58) | 0.01 | WT SHAM vs KO SHAM | -75% (-87, -52) | <0.0001 |
| 17(18)-EpETE | EPA | Epoxide | HDL | SHAM | KO | 2.79 (2.12, 3.68) | . | TAC WT vs TAC KO | -68% (-80, -48) | <0.0001 |
| 17(18)-EpETE | EPA | Epoxide | HDL | TAC | WT | 15.37 (12.75, 18.51) | . | SHAM WT vs TAC WT | 26% (-28, 58) | >0.80 |
| 17(18)-EpETE | EPA | Epoxide | HDL | TAC | KO | 4.93 (4.05, 5.99) | . | SHAM KO - TAC KO | 43% (69, -3) | 0.07 |
| 7(8)-EpDPE | DHA | Epoxide | HDL | SHAM | WT | 0.25 (0.11, 0.57) | 0.1 | . | . | . |
| 7(8)-EpDPE | DHA | Epoxide | HDL | SHAM | KO | 0.03 (0.01, 0.07) | . | . | . | . |
| 7(8)-EpDPE | DHA | Epoxide | HDL | TAC | WT | 0.1 (0.05, 0.19) | . | . | . | . |
| 7(8)-EpDPE | DHA | Epoxide | HDL | TAC | KO | 0.05 (0.03, 0.1) | . | . | . | . |
| 10(11)-EpDPE | DHA | Epoxide | HDL | SHAM | WT | 3.51 (2.51, 4.9) | 0.04 | WT SHAM vs KO SHAM | -49% (-79, 23) | 0.33 |
| 10(11)-EpDPE | DHA | Epoxide | HDL | SHAM | KO | 1.8 (1.24, 2.59) | . | TAC WT vs TAC KO | 3% (-45, 94) | >0.80 |
| 10(11)-EpDPE | DHA | Epoxide | HDL | TAC | WT | 2.04 (1.59, 2.61) | . | SHAM WT vs TAC WT | -72% (-258, 17) | 0.38 |
| 10(11)-EpDPE | DHA | Epoxide | HDL | TAC | KO | 2.1 (1.62, 2.73) | . | SHAM KO - TAC KO | 15% (61, -88) | >0.80 |
| 13(14)-EpDPE | DHA | Epoxide | HDL | SHAM | WT | 1.31 (0.66, 2.61) | <0.0001 | WT SHAM vs KO SHAM | -89% (-98, -36) | 0.004 |
| 13(14)-EpDPE | DHA | Epoxide | HDL | SHAM | KO | 0.14 (0.07, 0.3) | . | TAC WT vs TAC KO | -37% (-83, 128) | >0.80 |
| 13(14)-EpDPE | DHA | Epoxide | HDL | TAC | WT | 0.14 (0.08, 0.22) | . | SHAM WT vs TAC WT | -872% (-4257, -117) | 0.0001 |
| 13(14)-EpDPE | DHA | Epoxide | HDL | TAC | KO | 0.08 (0.05, 0.14) | . | SHAM KO - TAC KO | -66% (67, -741) | >0.80 |
| 16(17)-EpDPE | DHA | Epoxide | HDL | SHAM | WT | 1.46 (0.91, 2.34) | 0.02 | WT SHAM vs KO SHAM | -92% (-98, -74) | <0.0001 |
| 16(17)-EpDPE | DHA | Epoxide | HDL | SHAM | KO | 0.11 (0.07, 0.19) | . | TAC WT vs TAC KO | -11% (-63, 117) | >0.80 |
| 16(17)-EpDPE | DHA | Epoxide | HDL | TAC | WT | 0.15 (0.1, 0.21) | . | SHAM WT vs TAC WT | -894% (-2691, -254) | <0.0001 |
| 16(17)-EpDPE | DHA | Epoxide | HDL | TAC | KO | 0.13 (0.09, 0.19) | . | SHAM KO - TAC KO | 14% (72, -161) | >0.80 |
| 19(20)-EpDPE | DHA | Epoxide | HDL | SHAM | WT | 0.19 (0.06, 0.61) | <0.0001 | WT SHAM vs KO SHAM | 764% (-59, 18157) | 0.46 |
| 19(20)-EpDPE | DHA | Epoxide | HDL | SHAM | KO | 1.61 (0.44, 5.86) | . | TAC WT vs TAC KO | -75% (-97, 123) | 0.63 |
| 19(20)-EpDPE | DHA | Epoxide | HDL | TAC | WT | 0.1 (0.04, 0.25) | . | SHAM WT vs TAC WT | -80% (-2225, 86) | >0.80 |
| 19(20)-EpDPE | DHA | Epoxide | HDL | TAC | KO | 0.03 (0.01, 0.06) | . | SHAM KO - TAC KO | -6204% (-299, -99402) | 0.0002 |
| 9(10)-DiHOME | LA | Diol | HDL | SHAM | WT | 11.8 (9.5, 14.6) | <0.0001 | WT SHAM vs KO SHAM | -13% (-50, 52) | >0.80 |
| 9(10)-DiHOME | LA | Diol | HDL | SHAM | KO | 10.3 (8.2, 13) | . | TAC WT vs TAC KO | -39% (-59, -10) | 0.003 |
| 9(10)-DiHOME | LA | Diol | HDL | TAC | WT | 18.7 (16, 21.9) | . | SHAM WT vs TAC WT | 37% (0, 60) | 0.05 |
| 9(10)-DiHOME | LA | Diol | HDL | TAC | KO | 11.3 (9.6, 13.3) | . | SHAM KO - TAC KO | 9% (45, -50) | >0.80 |
| 12(13)-DiHOME | LA | Diol | HDL | SHAM | WT | 21.3 (16.4, 27.6) | <0.0001 | WT SHAM vs KO SHAM | 198% (51, 486) | <0.0001 |

| | | | | | | | | | | |
|---|---|---|---|---|---|---|---|---|---|---|
| 12(13)-DiHOME | LA | Diol | HDL | SHAM | KO | 63.3 (47.6, 84.2) | . | TAC WT vs TAC KO | -34% (-60, 7) | 0.17 |
| 12(13)-DiHOME | LA | Diol | HDL | TAC | WT | 100.3 (82.8, 121.5) | . | SHAM WT vs TAC WT | 79% (63, 88) | <0.0001 |
| 12(13)-DiHOME | LA | Diol | HDL | TAC | KO | 65.9 (53.9, 80.6) | . | SHAM KO - TAC KO | 4% (48, -77) | >0.80 |
| 8(9)-DiHETrE | AA | Diol | HDL | SHAM | WT | 13.08 (8.23, 20.79) | 0.67 | . | . | . |
| 8(9)-DiHETrE | AA | Diol | HDL | SHAM | KO | 1.78 (1.07, 2.95) | . | . | . | . |
| 8(9)-DiHETrE | AA | Diol | HDL | TAC | WT | 1.74 (1.24, 2.45) | . | . | . | . |
| 8(9)-DiHETrE | AA | Diol | HDL | TAC | KO | 0.98 (0.69, 1.4) | . | . | . | . |
| 11(12)-DiHETrE | AA | Diol | HDL | SHAM | WT | 2.29 (1.58, 3.33) | 0.07 | . | . | . |
| 11(12)-DiHETrE | AA | Diol | HDL | SHAM | KO | 0.46 (0.3, 0.68) | . | . | . | . |
| 11(12)-DiHETrE | AA | Diol | HDL | TAC | WT | 1.33 (1.01, 1.75) | . | . | . | . |
| 11(12)-DiHETrE | AA | Diol | HDL | TAC | KO | 1.44 (1.08, 1.92) | . | . | . | . |
| 14(15)-DiHETrE | AA | Diol | HDL | SHAM | WT | 0.51 (0.32, 0.81) | <0.0001 | WT SHAM vs KO SHAM | 161% (-23, 784) | 0.28 |
| 14(15)-DiHETrE | AA | Diol | HDL | SHAM | KO | 1.32 (0.79, 2.2) | . | TAC WT vs TAC KO | 8% (-55, 160) | >0.80 |
| 14(15)-DiHETrE | AA | Diol | HDL | TAC | WT | 1.3 (0.92, 1.84) | . | SHAM WT vs TAC WT | 61% (-8, 86) | 0.1 |
| 14(15)-DiHETrE | AA | Diol | HDL | TAC | KO | 1.4 (0.98, 2.02) | . | SHAM KO - TAC KO | 6% (69, -183) | >0.80 |
| 8(9)-DiHETE | EPA | Diol | HDL | SHAM | WT | 3.21 (2.48, 4.15) | <0.0001 | WT SHAM vs KO SHAM | 248% (78, 580) | <0.0001 |
| 8(9)-DiHETE | EPA | Diol | HDL | SHAM | KO | 11.16 (8.43, 14.79) | . | TAC WT vs TAC KO | -32% (-58, 10) | 0.26 |
| 8(9)-DiHETE | EPA | Diol | HDL | TAC | WT | 12.17 (10.07, 14.71) | . | SHAM WT vs TAC WT | 74% (54, 85) | <0.0001 |
| 8(9)-DiHETE | EPA | Diol | HDL | TAC | KO | 8.27 (6.78, 10.09) | . | SHAM KO - TAC KO | -35% (26, -147) | >0.80 |
| 11(12)-DiHETE | EPA | Diol | HDL | SHAM | WT | 0.94 (0.55, 1.6) | 0.007 | WT SHAM vs KO SHAM | 15% (-72, 362) | >0.80 |
| 11(12)-DiHETE | EPA | Diol | HDL | SHAM | KO | 1.07 (0.6, 1.93) | . | TAC WT vs TAC KO | -14% (-68, 136) | >0.80 |
| 11(12)-DiHETE | EPA | Diol | HDL | TAC | WT | 0.51 (0.34, 0.75) | . | SHAM WT vs TAC WT | -85% (-494, 43) | >0.80 |
| 11(12)-DiHETE | EPA | Diol | HDL | TAC | KO | 0.44 (0.29, 0.66) | . | SHAM KO - TAC KO | -145% (31, -765) | 0.45 |
| 14(15)-DiHETE | EPA | Diol | HDL | SHAM | WT | 3.28 (2.24, 4.8) | 0.06 | . | . | . |
| 14(15)-DiHETE | EPA | Diol | HDL | SHAM | KO | 1.26 (0.83, 1.91) | . | . | . | . |
| 14(15)-DiHETE | EPA | Diol | HDL | TAC | WT | 5.12 (3.87, 6.78) | . | . | . | . |
| 14(15)-DiHETE | EPA | Diol | HDL | TAC | KO | 3.02 (2.25, 4.06) | . | . | . | . |
| 17(18)-DiHETE | EPA | Diol | HDL | SHAM | WT | 0.98 (0.58, 1.65) | 0.29 | . | . | . |
| 17(18)-DiHETE | EPA | Diol | HDL | SHAM | KO | 0.37 (0.21, 0.66) | . | . | . | . |
| 17(18)-DiHETE | EPA | Diol | HDL | TAC | WT | 3.26 (2.22, 4.79) | . | . | . | . |
| 17(18)-DiHETE | EPA | Diol | HDL | TAC | KO | 0.59 (0.39, 0.88) | . | . | . | . |
| 9-HODE | LA | Alcohol | LDL | SHAM | WT | 17.5 (13.8, 22) | <0.0001 | WT SHAM vs KO SHAM | -32% (-63, 25) | 0.65 |
| 9-HODE | LA | Alcohol | LDL | SHAM | KO | 11.9 (9.2, 15.4) | . | TAC WT vs TAC KO | -11% (-42, 38) | >0.80 |
| 9-HODE | LA | Alcohol | LDL | TAC | WT | 11.5 (9.7, 13.7) | . | SHAM WT vs TAC WT | -51% (-152, 9) | 0.23 |
| 9-HODE | LA | Alcohol | LDL | TAC | KO | 10.3 (8.6, 12.3) | . | SHAM KO - TAC KO | -16% (33, -101) | >0.80 |
| 13-HODE | LA | Alcohol | LDL | SHAM | WT | 17.5 (14.1, 21.7) | 0.0003 | WT SHAM vs KO SHAM | -41% (-66, 4) | 0.1 |
| 13-HODE | LA | Alcohol | LDL | SHAM | KO | 10.4 (8.2, 13.2) | . | TAC WT vs TAC KO | -48% (-66, -23) | <0.0001 |
| 13-HODE | LA | Alcohol | LDL | TAC | WT | 18.3 (15.6, 21.5) | . | SHAM WT vs TAC WT | 4% (-53, 41) | >0.80 |
| 13-HODE | LA | Alcohol | LDL | TAC | KO | 9.4 (8, 11.2) | . | SHAM KO - TAC KO | -10% (34, -83) | >0.80 |
| 9-HOTrE | aLA | Alcohol | LDL | SHAM | WT | 0.27 (0.14, 0.54) | <0.0001 | WT SHAM vs KO SHAM | -71% (-95, 68) | 0.46 |
| 9-HOTrE | aLA | Alcohol | LDL | SHAM | KO | 0.08 (0.04, 0.16) | . | TAC WT vs TAC KO | 126% (-37, 714) | 0.63 |
| 9-HOTrE | aLA | Alcohol | LDL | TAC | WT | 0.19 (0.11, 0.31) | . | SHAM WT vs TAC WT | -45% (-542, 67) | >0.80 |
| 9-HOTrE | aLA | Alcohol | LDL | TAC | KO | 0.43 (0.25, 0.72) | . | SHAM KO - TAC KO | 82% (96, 9) | 0.03 |
| 13-HOTrE | aLA | Alcohol | LDL | SHAM | WT | 0.66 (0.32, 1.34) | 0.0002 | WT SHAM vs KO SHAM | -93% (-99, -58) | 0.0003 |
| 13-HOTrE | aLA | Alcohol | LDL | SHAM | KO | 0.04 (0.02, 0.09) | . | TAC WT vs TAC KO | -6% (-75, 259) | >0.80 |
| 13-HOTrE | aLA | Alcohol | LDL | TAC | WT | 0.4 (0.24, 0.68) | . | SHAM WT vs TAC WT | -63% (-675, 66) | >0.80 |
| 13-HOTrE | aLA | Alcohol | LDL | TAC | KO | 0.38 (0.22, 0.66) | . | SHAM KO - TAC KO | 89% (98, 39) | 0.002 |
| 5-HETE | AA | Alcohol | LDL | SHAM | WT | 1.29 (0.73, 2.29) | <0.0001 | WT SHAM vs KO SHAM | -82% (-96, -18) | 0.01 |
| 5-HETE | AA | Alcohol | LDL | SHAM | KO | 0.23 (0.13, 0.44) | . | TAC WT vs TAC KO | 4% (-65, 209) | >0.80 |
| 5-HETE | AA | Alcohol | LDL | TAC | WT | 0.49 (0.32, 0.75) | . | SHAM WT vs TAC WT | -163% (-826, 25) | 0.32 |
| 5-HETE | AA | Alcohol | LDL | TAC | KO | 0.51 (0.33, 0.8) | . | SHAM KO - TAC KO | 54% (88, -78) | 0.77 |
| 9-HETE | AA | Alcohol | LDL | SHAM | WT | 0.45 (0.18, 1.11) | 0.01 | WT SHAM vs KO SHAM | -68% (-97, 255) | >0.80 |
| 9-HETE | AA | Alcohol | LDL | SHAM | KO | 0.14 (0.05, 0.39) | . | TAC WT vs TAC KO | 194% (-48, 1553) | 0.66 |
| 9-HETE | AA | Alcohol | LDL | TAC | WT | 0.48 (0.25, 0.95) | . | SHAM WT vs TAC WT | 8% (-586, 88) | >0.80 |
| 9-HETE | AA | Alcohol | LDL | TAC | KO | 1.42 (0.7, 2.89) | . | SHAM KO - TAC KO | 90% (99, 11) | 0.03 |
| 12-HETE | AA | Alcohol | LDL | SHAM | WT | 0.7 (0.24, 2.05) | <0.0001 | WT SHAM vs KO SHAM | -94% (-100, -3) | 0.05 |
| 12-HETE | AA | Alcohol | LDL | SHAM | KO | 0.04 (0.01, 0.13) | . | TAC WT vs TAC KO | -53% (-94, 255) | >0.80 |
| 12-HETE | AA | Alcohol | LDL | TAC | WT | 0.14 (0.06, 0.3) | . | SHAM WT vs TAC WT | -417% (-5384, 51) | 0.48 |
| 12-HETE | AA | Alcohol | LDL | TAC | KO | 0.06 (0.03, 0.14) | . | SHAM KO - TAC KO | 35% (95, -725) | >0.80 |
| 15-HETE | AA | Alcohol | LDL | SHAM | WT | 1.01 (0.52, 1.94) | <0.0001 | WT SHAM vs KO SHAM | -96% (-99, -76) | <0.0001 |
| 15-HETE | AA | Alcohol | LDL | SHAM | KO | 0.04 (0.02, 0.09) | . | TAC WT vs TAC KO | 83% (-47, 530) | >0.80 |
| 15-HETE | AA | Alcohol | LDL | TAC | WT | 0.3 (0.19, 0.49) | . | SHAM WT vs TAC WT | -233% (-1302, 21) | 0.2 |
| 15-HETE | AA | Alcohol | LDL | TAC | KO | 0.55 (0.33, 0.92) | . | SHAM KO - TAC KO | 92% (98, 63) | <0.0001 |
| 5-HEPE | EPA | Alcohol | LDL | SHAM | WT | 1.22 (0.72, 2.06) | <0.0001 | WT SHAM vs KO SHAM | -98% (-99, -90) | <0.0001 |
| 5-HEPE | EPA | Alcohol | LDL | SHAM | KO | 0.03 (0.02, 0.05) | . | TAC WT vs TAC KO | -38% (-77, 66) | >0.80 |
| 5-HEPE | EPA | Alcohol | LDL | TAC | WT | 0.33 (0.23, 0.49) | . | SHAM WT vs TAC WT | -264% (-1043, -16) | 0.01 |
| 5-HEPE | EPA | Alcohol | LDL | TAC | KO | 0.21 (0.14, 0.31) | . | SHAM KO - TAC KO | 86% (96, 50) | <0.0001 |
| 8-HEPE | EPA | Alcohol | LDL | SHAM | WT | 1.45 (0.9, 2.33) | <0.0001 | WT SHAM vs KO SHAM | -90% (-97, -64) | <0.0001 |
| 8-HEPE | EPA | Alcohol | LDL | SHAM | KO | 0.15 (0.09, 0.25) | . | TAC WT vs TAC KO | 183% (15, 597) | 0.01 |
| 8-HEPE | EPA | Alcohol | LDL | TAC | WT | 0.17 (0.12, 0.25) | . | SHAM WT vs TAC WT | -739% (-2287, -195) | <0.0001 |
| 8-HEPE | EPA | Alcohol | LDL | TAC | KO | 0.49 (0.34, 0.71) | . | SHAM KO - TAC KO | 69% (90, 5) | 0.03 |
| 9-HEPE | EPA | Alcohol | LDL | SHAM | WT | 1.06 (0.59, 1.93) | 0.1 | . | . | . |
| 9-HEPE | EPA | Alcohol | LDL | SHAM | KO | 0.05 (0.03, 0.1) | . | . | . | . |
| 9-HEPE | EPA | Alcohol | LDL | TAC | WT | 0.18 (0.12, 0.28) | . | . | . | . |
| 9-HEPE | EPA | Alcohol | LDL | TAC | KO | 0.25 (0.16, 0.4) | . | . | . | . |
| 11-HEPE | EPA | Alcohol | LDL | SHAM | WT | 0.13 (0.05, 0.29) | 0.02 | WT SHAM vs KO SHAM | 12% (-88, 931) | >0.80 |
| 11-HEPE | EPA | Alcohol | LDL | SHAM | KO | 0.14 (0.06, 0.36) | . | TAC WT vs TAC KO | 94% (-61, 861) | >0.80 |
| 11-HEPE | EPA | Alcohol | LDL | TAC | WT | 0.07 (0.04, 0.13) | . | SHAM WT vs TAC WT | -78% (-1044, 72) | >0.80 |
| 11-HEPE | EPA | Alcohol | LDL | TAC | KO | 0.14 (0.07, 0.26) | . | SHAM KO - TAC KO | -3% (86, -666) | >0.80 |
| 12-HEPE | EPA | Alcohol | LDL | SHAM | WT | 1.16 (0.7, 1.93) | <0.0001 | WT SHAM vs KO SHAM | -97% (-99, -88) | <0.0001 |
| 12-HEPE | EPA | Alcohol | LDL | SHAM | KO | 0.04 (0.02, 0.07) | . | TAC WT vs TAC KO | 97% (-24, 408) | 0.45 |
| 12-HEPE | EPA | Alcohol | LDL | TAC | WT | 0.06 (0.04, 0.09) | . | SHAM WT vs TAC WT | -1766% (-5523, -519) | <0.0001 |
| 12-HEPE | EPA | Alcohol | LDL | TAC | KO | 0.12 (0.08, 0.18) | . | SHAM KO - TAC KO | 69% (91, -2) | 0.06 |
| 15-HEPE | EPA | Alcohol | LDL | SHAM | WT | 0.51 (0.27, 0.96) | 0.24 | . | . | . |
| 15-HEPE | EPA | Alcohol | LDL | SHAM | KO | 0.22 (0.11, 0.44) | . | . | . | . |
| 15-HEPE | EPA | Alcohol | LDL | TAC | WT | 0.33 (0.2, 0.52) | . | . | . | . |

| | | | | | | | | | | |
|---|---|---|---|---|---|---|---|---|---|---|
| 15-HEPE | EPA | Alcohol | LDL | TAC | KO | 0.21 (0.13, 0.34) | . | . | . | . |
| 18-HEPE | EPA | Alcohol | LDL | SHAM | WT | 0.25 (0.14, 0.48) | <0.0001 | WT SHAM vs KO SHAM | -84% (-97, -19) | 0.01 |
| 18-HEPE | EPA | Alcohol | LDL | SHAM | KO | 0.04 (0.02, 0.08) | . | TAC WT vs TAC KO | 60% (-51, 423) | >0.80 |
| 18-HEPE | EPA | Alcohol | LDL | TAC | WT | 0.05 (0.03, 0.08) | . | SHAM WT vs TAC WT | -397% (-1869, -25) | 0.01 |
| 18-HEPE | EPA | Alcohol | LDL | TAC | KO | 0.08 (0.05, 0.13) | . | SHAM KO - TAC KO | 51% (89, -115) | >0.80 |
| 4-HDoHE | DHA | Alcohol | LDL | SHAM | WT | 0.24 (0.1, 0.56) | 0.01 | WT SHAM vs KO SHAM | -57% (-95, 303) | >0.80 |
| 4-HDoHE | DHA | Alcohol | LDL | SHAM | KO | 0.1 (0.04, 0.26) | . | TAC WT vs TAC KO | -11% (-82, 348) | >0.80 |
| 4-HDoHE | DHA | Alcohol | LDL | TAC | WT | 0.15 (0.08, 0.27) | . | SHAM WT vs TAC WT | -62% (-966, 75) | >0.80 |
| 4-HDoHE | DHA | Alcohol | LDL | TAC | KO | 0.13 (0.07, 0.25) | . | SHAM KO - TAC KO | 22% (90, -495) | >0.80 |
| 7-HDoHE | DHA | Alcohol | LDL | SHAM | WT | 0.24 (0.1, 0.57) | 0.001 | WT SHAM vs KO SHAM | 4% (-89, 913) | >0.80 |
| 7-HDoHE | DHA | Alcohol | LDL | SHAM | KO | 0.25 (0.1, 0.64) | . | TAC WT vs TAC KO | -51% (-91, 152) | >0.80 |
| 7-HDoHE | DHA | Alcohol | LDL | TAC | WT | 0.51 (0.27, 0.98) | . | SHAM WT vs TAC WT | 54% (-211, 93) | >0.80 |
| 7-HDoHE | DHA | Alcohol | LDL | TAC | KO | 0.25 (0.13, 0.49) | . | SHAM KO - TAC KO | 1% (87, -671) | >0.80 |
| 8-HDoHE | DHA | Alcohol | LDL | SHAM | WT | 1.58 (0.73, 3.46) | 0.0002 | WT SHAM vs KO SHAM | -84% (-98, 26) | 0.14 |
| 8-HDoHE | DHA | Alcohol | LDL | SHAM | KO | 0.26 (0.11, 0.61) | . | TAC WT vs TAC KO | -47% (-88, 132) | >0.80 |
| 8-HDoHE | DHA | Alcohol | LDL | TAC | WT | 0.38 (0.21, 0.67) | . | SHAM WT vs TAC WT | -322% (-2229, 23) | 0.19 |
| 8-HDoHE | DHA | Alcohol | LDL | TAC | KO | 0.2 (0.11, 0.37) | . | SHAM KO - TAC KO | -30% (79, -721) | >0.80 |
| 10-HDoHE | DHA | Alcohol | LDL | SHAM | WT | 1.22 (0.7, 2.13) | 0.02 | WT SHAM vs KO SHAM | -77% (-95, -2) | 0.04 |
| 10-HDoHE | DHA | Alcohol | LDL | SHAM | KO | 0.28 (0.15, 0.51) | . | TAC WT vs TAC KO | -22% (-73, 123) | >0.80 |
| 10-HDoHE | DHA | Alcohol | LDL | TAC | WT | 0.37 (0.25, 0.57) | . | SHAM WT vs TAC WT | -226% (-1004, 4) | 0.07 |
| 10-HDoHE | DHA | Alcohol | LDL | TAC | KO | 0.29 (0.19, 0.45) | . | SHAM KO - TAC KO | 4% (74, -258) | >0.80 |
| 11-HDoHE | DHA | Alcohol | LDL | SHAM | WT | 0.19 (0.08, 0.47) | 0.24 | . | . | . |
| 11-HDoHE | DHA | Alcohol | LDL | SHAM | KO | 0.14 (0.05, 0.36) | . | . | . | . |
| 11-HDoHE | DHA | Alcohol | LDL | TAC | WT | 0.16 (0.08, 0.3) | . | . | . | . |
| 11-HDoHE | DHA | Alcohol | LDL | TAC | KO | 0.23 (0.12, 0.46) | . | . | . | . |
| 13-HDoHE | DHA | Alcohol | LDL | SHAM | WT | 0.75 (0.3, 1.87) | 0.0009 | WT SHAM vs KO SHAM | -72% (-97, 203) | >0.80 |
| 13-HDoHE | DHA | Alcohol | LDL | SHAM | KO | 0.21 (0.08, 0.58) | . | TAC WT vs TAC KO | -67% (-94, 84) | 0.61 |
| 13-HDoHE | DHA | Alcohol | LDL | TAC | WT | 0.48 (0.24, 0.93) | . | SHAM WT vs TAC WT | -58% (-1052, 78) | >0.80 |
| 13-HDoHE | DHA | Alcohol | LDL | TAC | KO | 0.16 (0.08, 0.32) | . | SHAM KO - TAC KO | -35% (84, -1048) | >0.80 |
| 14-HDoHE | DHA | Alcohol | LDL | SHAM | WT | 1.3 (0.56, 3.03) | <0.0001 | WT SHAM vs KO SHAM | -80% (-98, 87) | 0.43 |
| 14-HDoHE | DHA | Alcohol | LDL | SHAM | KO | 0.27 (0.11, 0.67) | . | TAC WT vs TAC KO | 144% (-51, 1102) | >0.80 |
| 14-HDoHE | DHA | Alcohol | LDL | TAC | WT | 0.21 (0.11, 0.39) | . | SHAM WT vs TAC WT | -530% (-3920, 1) | 0.05 |
| 14-HDoHE | DHA | Alcohol | LDL | TAC | KO | 0.5 (0.26, 0.97) | . | SHAM KO - TAC KO | 47% (93, -291) | >0.80 |
| 16-HDoHE | DHA | Alcohol | LDL | SHAM | WT | 0.33 (0.15, 0.72) | 0.13 | . | . | . |
| 16-HDoHE | DHA | Alcohol | LDL | SHAM | KO | 0.02 (0.01, 0.06) | . | . | . | . |
| 16-HDoHE | DHA | Alcohol | LDL | TAC | WT | 0.07 (0.04, 0.12) | . | . | . | . |
| 16-HDoHE | DHA | Alcohol | LDL | TAC | KO | 0.04 (0.02, 0.08) | . | . | . | . |
| 17-HDoHE | DHA | Alcohol | LDL | SHAM | WT | 2.09 (0.9, 4.88) | <0.0001 | WT SHAM vs KO SHAM | -94% (-99, -46) | 0.003 |
| 17-HDoHE | DHA | Alcohol | LDL | SHAM | KO | 0.12 (0.05, 0.31) | . | TAC WT vs TAC KO | -52% (-90, 135) | >0.80 |
| 17-HDoHE | DHA | Alcohol | LDL | TAC | WT | 0.56 (0.3, 1.04) | . | SHAM WT vs TAC WT | -275% (-2276, 41) | 0.44 |
| 17-HDoHE | DHA | Alcohol | LDL | TAC | KO | 0.27 (0.14, 0.52) | . | SHAM KO - TAC KO | 53% (94, -242) | >0.80 |
| 20-HDoHE | DHA | Alcohol | LDL | SHAM | WT | 0.36 (0.16, 0.82) | 0.61 | . | . | . |
| 20-HDoHE | DHA | Alcohol | LDL | SHAM | KO | 0.11 (0.05, 0.28) | . | . | . | . |
| 20-HDoHE | DHA | Alcohol | LDL | TAC | WT | 0.17 (0.09, 0.32) | . | . | . | . |
| 20-HDoHE | DHA | Alcohol | LDL | TAC | KO | 0.2 (0.11, 0.38) | . | . | . | . |
| 22-HDoHE | DHA | Alcohol | LDL | SHAM | WT | 0.33 (0.13, 0.83) | 0.51 | . | . | . |
| 22-HDoHE | DHA | Alcohol | LDL | SHAM | KO | 0.31 (0.11, 0.84) | . | . | . | . |
| 22-HDoHE | DHA | Alcohol | LDL | TAC | WT | 0.16 (0.08, 0.32) | . | . | . | . |
| 22-HDoHE | DHA | Alcohol | LDL | TAC | KO | 0.2 (0.1, 0.4) | . | . | . | . |
| 9-KODE | LA | Ketone | LDL | SHAM | WT | 3.45 (1.95, 6.09) | <0.0001 | WT SHAM vs KO SHAM | 17% (-74, 416) | >0.80 |
| 9-KODE | LA | Ketone | LDL | SHAM | KO | 4.03 (2.16, 7.52) | . | TAC WT vs TAC KO | 94% (-34, 467) | 0.67 |
| 9-KODE | LA | Ketone | LDL | TAC | WT | 1.64 (1.08, 2.5) | . | SHAM WT vs TAC WT | -110% (-628, 40) | 0.73 |
| 9-KODE | LA | Ketone | LDL | TAC | KO | 3.19 (2.05, 4.96) | . | SHAM KO - TAC KO | -26% (67, -384) | >0.80 |
| 13-KODE | LA | Ketone | LDL | SHAM | WT | 12.92 (7.6, 21.97) | <0.0001 | WT SHAM vs KO SHAM | -58% (-90, 67) | 0.65 |
| 13-KODE | LA | Ketone | LDL | SHAM | KO | 5.4 (3.02, 9.67) | . | TAC WT vs TAC KO | -54% (-83, 24) | 0.29 |
| 13-KODE | LA | Ketone | LDL | TAC | WT | 1.46 (0.99, 2.16) | . | SHAM WT vs TAC WT | -786% (-2725, -178) | <0.0001 |
| 13-KODE | LA | Ketone | LDL | TAC | KO | 0.67 (0.44, 1.01) | . | SHAM KO - TAC KO | -710% (-132, -2732) | <0.0001 |
| 5-KETE | AA | Ketone | LDL | SHAM | WT | 1.88 (1.26, 2.81) | 0.003 | WT SHAM vs KO SHAM | -78% (-92, -36) | 0.0005 |
| 5-KETE | AA | Ketone | LDL | SHAM | KO | 0.42 (0.27, 0.65) | . | TAC WT vs TAC KO | -52% (-77, 3) | 0.07 |
| 5-KETE | AA | Ketone | LDL | TAC | WT | 0.93 (0.69, 1.26) | . | SHAM WT vs TAC WT | -101% (-386, 17) | 0.27 |
| 5-KETE | AA | Ketone | LDL | TAC | KO | 0.45 (0.33, 0.62) | . | SHAM KO - TAC KO | 7% (64, -142) | >0.80 |
| 12-KETE | AA | Ketone | LDL | SHAM | WT | 0.55 (0.37, 0.84) | 0.02 | WT SHAM vs KO SHAM | -78% (-92, -35) | 0.0007 |
| 12-KETE | AA | Ketone | LDL | SHAM | KO | 0.12 (0.08, 0.19) | . | TAC WT vs TAC KO | 49% (-32, 225) | >0.80 |
| 12-KETE | AA | Ketone | LDL | TAC | WT | 0.79 (0.58, 1.07) | . | SHAM WT vs TAC WT | 30% (-74, 72) | >0.80 |
| 12-KETE | AA | Ketone | LDL | TAC | KO | 1.17 (0.85, 1.62) | . | SHAM KO - TAC KO | 90% (96, 72) | <0.0001 |
| 15-KETE | AA | Ketone | LDL | SHAM | WT | 0.57 (0.31, 1.05) | 0.02 | WT SHAM vs KO SHAM | -61% (-92, 92) | 0.74 |
| 15-KETE | AA | Ketone | LDL | SHAM | KO | 0.22 (0.11, 0.43) | . | TAC WT vs TAC KO | -51% (-84, 56) | 0.69 |
| 15-KETE | AA | Ketone | LDL | TAC | WT | 0.33 (0.21, 0.52) | . | SHAM WT vs TAC WT | -72% (-555, 55) | >0.80 |
| 15-KETE | AA | Ketone | LDL | TAC | KO | 0.16 (0.1, 0.26) | . | SHAM KO - TAC KO | -36% (68, -475) | >0.80 |
| 12-HpETE | AA | Peroxide | LDL | SHAM | WT | 0.99 (0.51, 1.89) | <0.0001 | WT SHAM vs KO SHAM | -70% (-94, 64) | 0.46 |
| 12-HpETE | AA | Peroxide | LDL | SHAM | KO | 0.3 (0.15, 0.61) | . | TAC WT vs TAC KO | -56% (-87, 50) | 0.56 |
| 12-HpETE | AA | Peroxide | LDL | TAC | WT | 0.38 (0.24, 0.62) | . | SHAM WT vs TAC WT | -157% (-956, 38) | 0.56 |
| 12-HpETE | AA | Peroxide | LDL | TAC | KO | 0.17 (0.1, 0.28) | . | SHAM KO - TAC KO | -75% (62, -705) | >0.80 |
| 15-HpETE | AA | Peroxide | LDL | SHAM | WT | 19.89 (12.72, 31.11) | 0.04 | WT SHAM vs KO SHAM | -81% (-94, -40) | 0.0004 |
| 15-HpETE | AA | Peroxide | LDL | SHAM | KO | 3.74 (2.29, 6.11) | . | TAC WT vs TAC KO | 172% (17, 530) | 0.007 |
| 15-HpETE | AA | Peroxide | LDL | TAC | WT | 0.94 (0.67, 1.31) | . | SHAM WT vs TAC WT | -2020% (-5535, -698) | <0.0001 |
| 15-HpETE | AA | Peroxide | LDL | TAC | KO | 2.55 (1.8, 3.6) | . | SHAM KO - TAC KO | -47% (49, -322) | >0.80 |
| Resolvin D1 | DHA | Triol | LDL | SHAM | WT | 2.75 (1.67, 4.51) | 0.0009 | WT SHAM vs KO SHAM | -23% (-79, 181) | >0.80 |
| Resolvin D1 | DHA | Triol | LDL | SHAM | KO | 2.11 (1.22, 3.63) | . | TAC WT vs TAC KO | 110% (-18, 435) | 0.28 |
| Resolvin D1 | DHA | Triol | LDL | TAC | WT | 1.01 (0.7, 1.45) | . | SHAM WT vs TAC WT | -173% (-710, 8) | 0.1 |
| Resolvin D1 | DHA | Triol | LDL | TAC | KO | 2.11 (1.44, 3.1) | . | SHAM KO - TAC KO | 0% (69, -222) | >0.80 |
| 9(10)-EpOME | LA | Epoxide | LDL | SHAM | WT | 23.6 (16.4, 33.9) | 0.0005 | WT SHAM vs KO SHAM | -85% (-94, -62) | <0.0001 |
| 9(10)-EpOME | LA | Epoxide | LDL | SHAM | KO | 3.5 (2.4, 5.2) | . | TAC WT vs TAC KO | -23% (-61, 52) | >0.80 |
| 9(10)-EpOME | LA | Epoxide | LDL | TAC | WT | 3.8 (2.9, 5) | . | SHAM WT vs TAC WT | -521% (-1274, -180) | <0.0001 |
| 9(10)-EpOME | LA | Epoxide | LDL | TAC | KO | 2.9 (2.2, 3.9) | . | SHAM KO - TAC KO | -20% (49, -184) | >0.80 |
| 12(13)-EpOME | LA | Epoxide | LDL | SHAM | WT | 22.8 (15.9, 32.7) | <0.0001 | WT SHAM vs KO SHAM | -90% (-96, -73) | <0.0001 |

| | | | | | | | | | | |
|---|---|---|---|---|---|---|---|---|---|---|
| 12(13)-EpOME | LA | Epoxide | LDL | SHAM | KO | 2.4 (1.6, 3.5) | . | TAC WT vs TAC KO | 24% (-37, 145) | >0.80 |
| 12(13)-EpOME | LA | Epoxide | LDL | TAC | WT | 2.9 (2.2, 3.8) | . | SHAM WT vs TAC WT | -680% (-1620, -253) | <0.0001 |
| 12(13)-EpOME | LA | Epoxide | LDL | TAC | KO | 3.6 (2.7, 4.8) | . | SHAM KO - TAC KO | 35% (72, -54) | >0.80 |
| 8(9)-EpETrE | AA | Epoxide | LDL | SHAM | WT | 4.94 (3.91, 6.24) | 0.21 | . | . | . |
| 8(9)-EpETrE | AA | Epoxide | LDL | SHAM | KO | 0.41 (0.32, 0.53) | . | . | . | . |
| 8(9)-EpETrE | AA | Epoxide | LDL | TAC | WT | 1.35 (1.14, 1.6) | . | . | . | . |
| 8(9)-EpETrE | AA | Epoxide | LDL | TAC | KO | 1.43 (1.19, 1.71) | . | . | . | . |
| 11(12)-EpETrE | AA | Epoxide | LDL | SHAM | WT | 1.59 (0.85, 2.97) | 0.09 | . | . | . |
| 11(12)-EpETrE | AA | Epoxide | LDL | SHAM | KO | 0.17 (0.08, 0.33) | . | . | . | . |
| 11(12)-EpETrE | AA | Epoxide | LDL | TAC | WT | 0.27 (0.17, 0.43) | . | . | . | . |
| 11(12)-EpETrE | AA | Epoxide | LDL | TAC | KO | 0.55 (0.34, 0.9) | . | . | . | . |
| 14(15)-EpETrE | AA | Epoxide | LDL | SHAM | WT | 1.71 (0.95, 3.1) | <0.0001 | WT SHAM vs KO SHAM | -39% (-87, 184) | >0.80 |
| 14(15)-EpETrE | AA | Epoxide | LDL | SHAM | KO | 1.04 (0.54, 1.98) | . | TAC WT vs TAC KO | -45% (-82, 68) | >0.80 |
| 14(15)-EpETrE | AA | Epoxide | LDL | TAC | WT | 0.54 (0.35, 0.83) | . | SHAM WT vs TAC WT | -218% (-1062, 13) | 0.13 |
| 14(15)-EpETrE | AA | Epoxide | LDL | TAC | KO | 0.3 (0.19, 0.47) | . | SHAM KO - TAC KO | -249% (14, -1313) | 0.13 |
| 8(9)-EpETE | EPA | Epoxide | LDL | SHAM | WT | 2.37 (1.54, 3.66) | <0.0001 | WT SHAM vs KO SHAM | -76% (-92, -26) | 0.003 |
| 8(9)-EpETE | EPA | Epoxide | LDL | SHAM | KO | 0.57 (0.35, 0.91) | . | TAC WT vs TAC KO | 387% (115, 1001) | <0.0001 |
| 8(9)-EpETE | EPA | Epoxide | LDL | TAC | WT | 0.34 (0.25, 0.47) | . | SHAM WT vs TAC WT | -601% (-1709, -171) | <0.0001 |
| 8(9)-EpETE | EPA | Epoxide | LDL | TAC | KO | 1.65 (1.18, 2.3) | . | SHAM KO - TAC KO | 66% (88, 4) | 0.03 |
| 11(12)-EpETE | EPA | Epoxide | LDL | SHAM | WT | 0.7 (0.46, 1.06) | 0.59 | . | . | . |
| 11(12)-EpETE | EPA | Epoxide | LDL | SHAM | KO | 0.52 (0.33, 0.82) | . | . | . | . |
| 11(12)-EpETE | EPA | Epoxide | LDL | TAC | WT | 1.39 (1.02, 1.9) | . | . | . | . |
| 11(12)-EpETE | EPA | Epoxide | LDL | TAC | KO | 0.73 (0.53, 1.01) | . | . | . | . |
| 14(15)-EpETE | EPA | Epoxide | LDL | SHAM | WT | 8.73 (6.49, 11.74) | <0.0001 | WT SHAM vs KO SHAM | -79% (-90, -54) | <0.0001 |
| 14(15)-EpETE | EPA | Epoxide | LDL | SHAM | KO | 1.84 (1.33, 2.55) | . | TAC WT vs TAC KO | 3% (-41, 80) | >0.80 |
| 14(15)-EpETE | EPA | Epoxide | LDL | TAC | WT | 2.26 (1.82, 2.82) | . | SHAM WT vs TAC WT | -286% (-637, -102) | <0.0001 |
| 14(15)-EpETE | EPA | Epoxide | LDL | TAC | KO | 2.33 (1.85, 2.93) | . | SHAM KO - TAC KO | 21% (61, -60) | >0.80 |
| 17(18)-EpETE | EPA | Epoxide | LDL | SHAM | WT | 5.28 (4.3, 6.48) | 0.01 | WT SHAM vs KO SHAM | -45% (-68, -7) | 0.01 |
| 17(18)-EpETE | EPA | Epoxide | LDL | SHAM | KO | 2.88 (2.3, 3.61) | . | TAC WT vs TAC KO | -43% (-62, -17) | 0.0002 |
| 17(18)-EpETE | EPA | Epoxide | LDL | TAC | WT | 15.01 (12.9, 17.47) | . | SHAM WT vs TAC WT | 65% (45, 78) | <0.0001 |
| 17(18)-EpETE | EPA | Epoxide | LDL | TAC | KO | 8.48 (7.23, 9.94) | . | SHAM KO - TAC KO | 66% (79, 45) | <0.0001 |
| 7(8)-EpDPE | DHA | Epoxide | LDL | SHAM | WT | 0.12 (0.05, 0.28) | 0.1 | . | . | . |
| 7(8)-EpDPE | DHA | Epoxide | LDL | SHAM | KO | 0.03 (0.01, 0.06) | . | . | . | . |
| 7(8)-EpDPE | DHA | Epoxide | LDL | TAC | WT | 0.09 (0.05, 0.16) | . | . | . | . |
| 7(8)-EpDPE | DHA | Epoxide | LDL | TAC | KO | 0.09 (0.05, 0.17) | . | . | . | . |
| 10(11)-EpDPE | DHA | Epoxide | LDL | SHAM | WT | 1.97 (0.87, 4.45) | 0.04 | WT SHAM vs KO SHAM | -78% (-97, 86) | 0.45 |
| 10(11)-EpDPE | DHA | Epoxide | LDL | SHAM | KO | 0.43 (0.18, 1.06) | . | TAC WT vs TAC KO | 170% (-42, 1161) | 0.6 |
| 10(11)-EpDPE | DHA | Epoxide | LDL | TAC | WT | 0.83 (0.45, 1.52) | . | SHAM WT vs TAC WT | -137% (-1317, 60) | >0.80 |
| 10(11)-EpDPE | DHA | Epoxide | LDL | TAC | KO | 2.24 (1.19, 4.23) | . | SHAM KO - TAC KO | 81% (97, -33) | 0.18 |
| 13(14)-EpDPE | DHA | Epoxide | LDL | SHAM | WT | 0.08 (0.04, 0.16) | <0.0001 | WT SHAM vs KO SHAM | 237% (-46, 1999) | 0.56 |
| 13(14)-EpDPE | DHA | Epoxide | LDL | SHAM | KO | 0.27 (0.13, 0.59) | . | TAC WT vs TAC KO | 42% (-62, 430) | >0.80 |
| 13(14)-EpDPE | DHA | Epoxide | LDL | TAC | WT | 0.1 (0.06, 0.16) | . | SHAM WT vs TAC WT | 17% (-282, 82) | >0.80 |
| 13(14)-EpDPE | DHA | Epoxide | LDL | TAC | KO | 0.14 (0.08, 0.24) | . | SHAM KO - TAC KO | -97% (62, -928) | >0.80 |
| 16(17)-EpDPE | DHA | Epoxide | LDL | SHAM | WT | 1.04 (0.51, 2.09) | 0.02 | WT SHAM vs KO SHAM | -82% (-97, 15) | 0.1 |
| 16(17)-EpDPE | DHA | Epoxide | LDL | SHAM | KO | 0.19 (0.09, 0.41) | . | TAC WT vs TAC KO | 189% (-23, 985) | 0.26 |
| 16(17)-EpDPE | DHA | Epoxide | LDL | TAC | WT | 0.09 (0.06, 0.16) | . | SHAM WT vs TAC WT | -1019% (-5110, -140) | <0.0001 |
| 16(17)-EpDPE | DHA | Epoxide | LDL | TAC | KO | 0.27 (0.15, 0.46) | . | SHAM KO - TAC KO | 29% (86, -274) | >0.80 |
| 19(20)-EpDPE | DHA | Epoxide | LDL | SHAM | WT | 0.26 (0.11, 0.64) | <0.0001 | WT SHAM vs KO SHAM | -94% (-99, -38) | 0.006 |
| 19(20)-EpDPE | DHA | Epoxide | LDL | SHAM | KO | 0.02 (0.01, 0.04) | . | TAC WT vs TAC KO | -41% (-89, 224) | >0.80 |
| 19(20)-EpDPE | DHA | Epoxide | LDL | TAC | WT | 0.08 (0.04, 0.15) | . | SHAM WT vs TAC WT | -246% (-2391, 52) | 0.65 |
| 19(20)-EpDPE | DHA | Epoxide | LDL | TAC | KO | 0.04 (0.02, 0.09) | . | SHAM KO - TAC KO | 66% (96, -188) | >0.80 |
| 9(10)-DiHOME | LA | Diol | LDL | SHAM | WT | 15.07 (11.27, 20.14) | <0.0001 | WT SHAM vs KO SHAM | -81% (-91, -60) | <0.0001 |
| 9(10)-DiHOME | LA | Diol | LDL | SHAM | KO | 2.84 (2.07, 3.91) | . | TAC WT vs TAC KO | -92% (-95, -86) | <0.0001 |
| 9(10)-DiHOME | LA | Diol | LDL | TAC | WT | 23.1 (18.65, 28.62) | . | SHAM WT vs TAC WT | 35% (-23, 65) | 0.54 |
| 9(10)-DiHOME | LA | Diol | LDL | TAC | KO | 1.92 (1.53, 2.4) | . | SHAM KO - TAC KO | -48% (25, -194) | 0.77 |
| 12(13)-DiHOME | LA | Diol | LDL | SHAM | WT | 119.09 (98.46, 144.03) | <0.0001 | WT SHAM vs KO SHAM | -84% (-90, -74) | <0.0001 |
| 12(13)-DiHOME | LA | Diol | LDL | SHAM | KO | 19.03 (15.45, 23.44) | . | TAC WT vs TAC KO | 3% (-28, 47) | >0.80 |
| 12(13)-DiHOME | LA | Diol | LDL | TAC | WT | 35.67 (31, 41.05) | . | SHAM WT vs TAC WT | -234% (-406, -120) | <0.0001 |
| 12(13)-DiHOME | LA | Diol | LDL | TAC | KO | 36.63 (31.62, 42.45) | . | SHAM KO - TAC KO | 48% (67, 19) | 0.0003 |
| 8(9)-DiHETrE | AA | Diol | LDL | SHAM | WT | 12.06 (5.58, 26.03) | 0.67 | . | . | . |
| 8(9)-DiHETrE | AA | Diol | LDL | SHAM | KO | 0.64 (0.27, 1.48) | . | . | . | . |
| 8(9)-DiHETrE | AA | Diol | LDL | TAC | WT | 1.21 (0.68, 2.13) | . | . | . | . |
| 8(9)-DiHETrE | AA | Diol | LDL | TAC | KO | 0.44 (0.24, 0.81) | . | . | . | . |
| 11(12)-DiHETrE | AA | Diol | LDL | SHAM | WT | 3.05 (1.76, 5.27) | 0.07 | . | . | . |
| 11(12)-DiHETrE | AA | Diol | LDL | SHAM | KO | 0.33 (0.18, 0.59) | . | . | . | . |
| 11(12)-DiHETrE | AA | Diol | LDL | TAC | WT | 0.54 (0.36, 0.81) | . | . | . | . |
| 11(12)-DiHETrE | AA | Diol | LDL | TAC | KO | 0.65 (0.42, 0.99) | . | . | . | . |
| 14(15)-DiHETrE | AA | Diol | LDL | SHAM | WT | 1.98 (1.18, 3.35) | <0.0001 | WT SHAM vs KO SHAM | -69% (-92, 20) | 0.16 |
| 14(15)-DiHETrE | AA | Diol | LDL | SHAM | KO | 0.61 (0.34, 1.08) | . | TAC WT vs TAC KO | -67% (-87, -11) | 0.02 |
| 14(15)-DiHETrE | AA | Diol | LDL | TAC | WT | 1.15 (0.78, 1.69) | . | SHAM WT vs TAC WT | -73% (-441, 45) | >0.80 |
| 14(15)-DiHETrE | AA | Diol | LDL | TAC | KO | 0.38 (0.26, 0.58) | . | SHAM KO - TAC KO | -58% (54, -443) | >0.80 |
| 8(9)-DiHETE | EPA | Diol | LDL | SHAM | WT | 10.01 (7.5, 13.37) | <0.0001 | WT SHAM vs KO SHAM | -41% (-72, 25) | 0.47 |
| 8(9)-DiHETE | EPA | Diol | LDL | SHAM | KO | 5.9 (4.3, 8.09) | . | TAC WT vs TAC KO | 130% (34, 296) | 0.0001 |
| 8(9)-DiHETE | EPA | Diol | LDL | TAC | WT | 5.45 (4.4, 6.74) | . | SHAM WT vs TAC WT | -84% (-245, 2) | 0.07 |
| 8(9)-DiHETE | EPA | Diol | LDL | TAC | KO | 12.55 (10.03, 15.69) | . | SHAM KO - TAC KO | 53% (76, 7) | 0.02 |
| 11(12)-DiHETE | EPA | Diol | LDL | SHAM | WT | 0.91 (0.47, 1.78) | 0.007 | WT SHAM vs KO SHAM | -72% (-95, 62) | 0.42 |
| 11(12)-DiHETE | EPA | Diol | LDL | SHAM | KO | 0.26 (0.12, 0.54) | . | TAC WT vs TAC KO | 20% (-66, 322) | >0.80 |
| 11(12)-DiHETE | EPA | Diol | LDL | TAC | WT | 0.45 (0.27, 0.73) | . | SHAM WT vs TAC WT | -104% (-782, 53) | >0.80 |
| 11(12)-DiHETE | EPA | Diol | LDL | TAC | KO | 0.54 (0.32, 0.9) | . | SHAM KO - TAC KO | 52% (90, -134) | >0.80 |
| 14(15)-DiHETE | EPA | Diol | LDL | SHAM | WT | 0.72 (0.38, 1.37) | 0.06 | . | . | . |
| 14(15)-DiHETE | EPA | Diol | LDL | SHAM | KO | 0.6 (0.3, 1.22) | . | . | . | . |
| 14(15)-DiHETE | EPA | Diol | LDL | TAC | WT | 1.78 (1.11, 2.86) | . | . | . | . |
| 14(15)-DiHETE | EPA | Diol | LDL | TAC | KO | 0.51 (0.31, 0.85) | . | . | . | . |
| 17(18)-DiHETE | EPA | Diol | LDL | SHAM | WT | 1.87 (1.05, 3.34) | 0.29 | . | . | . |
| 17(18)-DiHETE | EPA | Diol | LDL | SHAM | KO | 2.36 (1.25, 4.45) | . | . | . | . |
| 17(18)-DiHETE | EPA | Diol | LDL | TAC | WT | 0.78 (0.51, 1.19) | . | . | . | . |

| | | | | | | | | | | |
|---|---|---|---|---|---|---|---|---|---|---|
| 17(18)-DiHETE | EPA | Diol | LDL | TAC | KO | 1.01 (0.64, 1.58) | . | . | . | . |
| 9-HODE | LA | Alcohol | VLDL | SHAM | WT | 16.2 (12.4, 21.3) | <0.0001 | WT SHAM vs KO SHAM | -50% (-75, 1) | 0.05 |
| 9-HODE | LA | Alcohol | VLDL | SHAM | KO | 8.1 (6, 10.8) | . | TAC WT vs TAC KO | -25% (-55, 24) | 0.77 |
| 9-HODE | LA | Alcohol | VLDL | TAC | WT | 19.5 (16, 23.9) | . | SHAM WT vs TAC WT | 17% (-50, 54) | >0.80 |
| 9-HODE | LA | Alcohol | VLDL | TAC | KO | 14.6 (11.8, 18) | . | SHAM KO - TAC KO | 45% (71, -5) | 0.1 |
| 13-HODE | LA | Alcohol | VLDL | SHAM | WT | 16.5 (13, 21) | 0.0003 | WT SHAM vs KO SHAM | -48% (-72, -2) | 0.03 |
| 13-HODE | LA | Alcohol | VLDL | SHAM | KO | 8.6 (6.6, 11.2) | . | TAC WT vs TAC KO | -56% (-72, -31) | <0.0001 |
| 13-HODE | LA | Alcohol | VLDL | TAC | WT | 21.9 (18.3, 26.2) | . | SHAM WT vs TAC WT | 25% (-28, 55) | >0.80 |
| 13-HODE | LA | Alcohol | VLDL | TAC | KO | 9.6 (7.9, 11.5) | . | SHAM KO - TAC KO | 10% (49, -59) | >0.80 |
| 9-HOTrE | aLA | Alcohol | VLDL | SHAM | WT | 0.27 (0.14, 0.52) | <0.0001 | WT SHAM vs KO SHAM | -92% (-98, -57) | 0.0002 |
| 9-HOTrE | aLA | Alcohol | VLDL | SHAM | KO | 0.02 (0.01, 0.05) | . | TAC WT vs TAC KO | -64% (-89, 22) | 0.2 |
| 9-HOTrE | aLA | Alcohol | VLDL | TAC | WT | 0.68 (0.42, 1.09) | . | SHAM WT vs TAC WT | 60% (-64, 90) | 0.6 |
| 9-HOTrE | aLA | Alcohol | VLDL | TAC | KO | 0.25 (0.15, 0.41) | . | SHAM KO - TAC KO | 91% (98, 59) | <0.0001 |
| 13-HOTrE | aLA | Alcohol | VLDL | SHAM | WT | 0.25 (0.15, 0.4) | 0.0002 | WT SHAM vs KO SHAM | 13% (-68, 294) | >0.80 |
| 13-HOTrE | aLA | Alcohol | VLDL | SHAM | KO | 0.28 (0.17, 0.47) | . | TAC WT vs TAC KO | 16% (-53, 185) | >0.80 |
| 13-HOTrE | aLA | Alcohol | VLDL | TAC | WT | 0.37 (0.26, 0.53) | . | SHAM WT vs TAC WT | 33% (-90, 76) | >0.80 |
| 13-HOTrE | aLA | Alcohol | VLDL | TAC | KO | 0.43 (0.3, 0.62) | . | SHAM KO - TAC KO | 35% (79, -102) | >0.80 |
| 5-HETE | AA | Alcohol | VLDL | SHAM | WT | 1.82 (0.74, 4.5) | <0.0001 | WT SHAM vs KO SHAM | -96% (-100, -62) | 0.0006 |
| 5-HETE | AA | Alcohol | VLDL | SHAM | KO | 0.06 (0.02, 0.17) | . | TAC WT vs TAC KO | -75% (-96, 37) | 0.23 |
| 5-HETE | AA | Alcohol | VLDL | TAC | WT | 0.26 (0.13, 0.51) | . | SHAM WT vs TAC WT | -595% (-4951, 4) | 0.06 |
| 5-HETE | AA | Alcohol | VLDL | TAC | KO | 0.06 (0.03, 0.13) | . | SHAM KO - TAC KO | 1% (88, -742) | >0.80 |
| 9-HETE | AA | Alcohol | VLDL | SHAM | WT | 0.77 (0.37, 1.59) | 0.01 | WT SHAM vs KO SHAM | -34% (-90, 335) | >0.80 |
| 9-HETE | AA | Alcohol | VLDL | SHAM | KO | 0.51 (0.23, 1.13) | . | TAC WT vs TAC KO | -14% (-78, 234) | >0.80 |
| 9-HETE | AA | Alcohol | VLDL | TAC | WT | 0.32 (0.19, 0.54) | . | SHAM WT vs TAC WT | -143% (-1078, 50) | >0.80 |
| 9-HETE | AA | Alcohol | VLDL | TAC | KO | 0.27 (0.15, 0.48) | . | SHAM KO - TAC KO | -88% (66, -931) | >0.80 |
| 12-HETE | AA | Alcohol | VLDL | SHAM | WT | 0.36 (0.13, 0.98) | <0.0001 | WT SHAM vs KO SHAM | -91% (-99, 20) | 0.09 |
| 12-HETE | AA | Alcohol | VLDL | SHAM | KO | 0.03 (0.01, 0.09) | . | TAC WT vs TAC KO | -38% (-91, 319) | >0.80 |
| 12-HETE | AA | Alcohol | VLDL | TAC | WT | 0.08 (0.04, 0.18) | . | SHAM WT vs TAC WT | -325% (-3816, 54) | 0.59 |
| 12-HETE | AA | Alcohol | VLDL | TAC | KO | 0.05 (0.02, 0.11) | . | SHAM KO - TAC KO | 42% (95, -541) | >0.80 |
| 15-HETE | AA | Alcohol | VLDL | SHAM | WT | 0.29 (0.18, 0.48) | <0.0001 | WT SHAM vs KO SHAM | -58% (-88, 50) | 0.51 |
| 15-HETE | AA | Alcohol | VLDL | SHAM | KO | 0.12 (0.07, 0.21) | . | TAC WT vs TAC KO | -31% (-72, 73) | >0.80 |
| 15-HETE | AA | Alcohol | VLDL | TAC | WT | 0.27 (0.19, 0.39) | . | SHAM WT vs TAC WT | -7% (-212, 63) | >0.80 |
| 15-HETE | AA | Alcohol | VLDL | TAC | KO | 0.19 (0.13, 0.27) | . | SHAM KO - TAC KO | 35% (79, -106) | >0.80 |
| 5-HEPE | EPA | Alcohol | VLDL | SHAM | WT | 0.81 (0.5, 1.33) | <0.0001 | WT SHAM vs KO SHAM | -60% (-89, 45) | 0.46 |
| 5-HEPE | EPA | Alcohol | VLDL | SHAM | KO | 0.33 (0.19, 0.57) | . | TAC WT vs TAC KO | 13% (-55, 185) | >0.80 |
| 5-HEPE | EPA | Alcohol | VLDL | TAC | WT | 0.24 (0.17, 0.35) | . | SHAM WT vs TAC WT | -235% (-879, -15) | 0.01 |
| 5-HEPE | EPA | Alcohol | VLDL | TAC | KO | 0.27 (0.19, 0.4) | . | SHAM KO - TAC KO | -20% (62, -282) | >0.80 |
| 8-HEPE | EPA | Alcohol | VLDL | SHAM | WT | 1.7 (1.07, 2.71) | <0.0001 | WT SHAM vs KO SHAM | -91% (-97, -69) | <0.0001 |
| 8-HEPE | EPA | Alcohol | VLDL | SHAM | KO | 0.16 (0.09, 0.26) | . | TAC WT vs TAC KO | 161% (9, 524) | 0.02 |
| 8-HEPE | EPA | Alcohol | VLDL | TAC | WT | 0.33 (0.23, 0.46) | . | SHAM WT vs TAC WT | -418% (-1329, -88) | <0.0001 |
| 8-HEPE | EPA | Alcohol | VLDL | TAC | KO | 0.86 (0.6, 1.23) | . | SHAM KO - TAC KO | 82% (94, 45) | 0.0001 |
| 9-HEPE | EPA | Alcohol | VLDL | SHAM | WT | 1.04 (0.54, 2.02) | 0.1 | . | . | . |
| 9-HEPE | EPA | Alcohol | VLDL | SHAM | KO | 0.14 (0.07, 0.29) | . | . | . | . |
| 9-HEPE | EPA | Alcohol | VLDL | TAC | WT | 0.31 (0.19, 0.51) | . | . | . | . |
| 9-HEPE | EPA | Alcohol | VLDL | TAC | KO | 0.14 (0.09, 0.24) | . | . | . | . |
| 11-HEPE | EPA | Alcohol | VLDL | SHAM | WT | 1 (0.52, 1.91) | 0.02 | WT SHAM vs KO SHAM | -93% (-99, -63) | <0.0001 |
| 11-HEPE | EPA | Alcohol | VLDL | SHAM | KO | 0.07 (0.03, 0.14) | . | TAC WT vs TAC KO | -28% (-79, 146) | >0.80 |
| 11-HEPE | EPA | Alcohol | VLDL | TAC | WT | 0.21 (0.13, 0.35) | . | SHAM WT vs TAC WT | -365% (-1833, -12) | 0.02 |
| 11-HEPE | EPA | Alcohol | VLDL | TAC | KO | 0.15 (0.09, 0.26) | . | SHAM KO - TAC KO | 56% (91, -106) | >0.80 |
| 12-HEPE | EPA | Alcohol | VLDL | SHAM | WT | 0.43 (0.19, 0.95) | <0.0001 | WT SHAM vs KO SHAM | -48% (-94, 324) | >0.80 |
| 12-HEPE | EPA | Alcohol | VLDL | SHAM | KO | 0.22 (0.09, 0.54) | . | TAC WT vs TAC KO | 209% (-32, 1297) | 0.37 |
| 12-HEPE | EPA | Alcohol | VLDL | TAC | WT | 0.07 (0.04, 0.12) | . | SHAM WT vs TAC WT | -548% (-3643, -12) | 0.03 |
| 12-HEPE | EPA | Alcohol | VLDL | TAC | KO | 0.2 (0.11, 0.38) | . | SHAM KO - TAC KO | -10% (83, -628) | >0.80 |
| 15-HEPE | EPA | Alcohol | VLDL | SHAM | WT | 0.36 (0.19, 0.71) | 0.24 | . | . | . |
| 15-HEPE | EPA | Alcohol | VLDL | SHAM | KO | 0.18 (0.09, 0.37) | . | . | . | . |
| 15-HEPE | EPA | Alcohol | VLDL | TAC | WT | 0.29 (0.18, 0.47) | . | . | . | . |
| 15-HEPE | EPA | Alcohol | VLDL | TAC | KO | 0.09 (0.05, 0.15) | . | . | . | . |
| 18-HEPE | EPA | Alcohol | VLDL | SHAM | WT | 0.14 (0.08, 0.25) | <0.0001 | WT SHAM vs KO SHAM | -80% (-96, -12) | 0.02 |
| 18-HEPE | EPA | Alcohol | VLDL | SHAM | KO | 0.03 (0.02, 0.05) | . | TAC WT vs TAC KO | 87% (-37, 454) | 0.76 |
| 18-HEPE | EPA | Alcohol | VLDL | TAC | WT | 0.03 (0.02, 0.05) | . | SHAM WT vs TAC WT | -315% (-1362, -18) | 0.01 |
| 18-HEPE | EPA | Alcohol | VLDL | TAC | KO | 0.06 (0.04, 0.1) | . | SHAM KO - TAC KO | 56% (89, -70) | 0.69 |
| 4-HDoHE | DHA | Alcohol | VLDL | SHAM | WT | 0.45 (0.21, 0.94) | 0.01 | WT SHAM vs KO SHAM | -88% (-98, -15) | 0.02 |
| 4-HDoHE | DHA | Alcohol | VLDL | SHAM | KO | 0.05 (0.02, 0.12) | . | TAC WT vs TAC KO | 57% (-61, 536) | >0.80 |
| 4-HDoHE | DHA | Alcohol | VLDL | TAC | WT | 0.12 (0.07, 0.2) | . | SHAM WT vs TAC WT | -283% (-1843, 24) | 0.21 |
| 4-HDoHE | DHA | Alcohol | VLDL | TAC | KO | 0.18 (0.1, 0.33) | . | SHAM KO - TAC KO | 70% (95, -73) | 0.5 |
| 7-HDoHE | DHA | Alcohol | VLDL | SHAM | WT | 3.59 (1.85, 6.95) | 0.001 | WT SHAM vs KO SHAM | -95% (-99, -70) | <0.0001 |
| 7-HDoHE | DHA | Alcohol | VLDL | SHAM | KO | 0.19 (0.09, 0.4) | . | TAC WT vs TAC KO | 12% (-68, 288) | >0.80 |
| 7-HDoHE | DHA | Alcohol | VLDL | TAC | WT | 0.19 (0.12, 0.31) | . | SHAM WT vs TAC WT | -1805% (-7975, -349) | <0.0001 |
| 7-HDoHE | DHA | Alcohol | VLDL | TAC | KO | 0.21 (0.13, 0.35) | . | SHAM KO - TAC KO | 8% (81, -339) | >0.80 |
| 8-HDoHE | DHA | Alcohol | VLDL | SHAM | WT | 0.4 (0.19, 0.85) | 0.0002 | WT SHAM vs KO SHAM | -22% (-89, 461) | >0.80 |
| 8-HDoHE | DHA | Alcohol | VLDL | SHAM | KO | 0.31 (0.14, 0.71) | . | TAC WT vs TAC KO | 23% (-70, 409) | >0.80 |
| 8-HDoHE | DHA | Alcohol | VLDL | TAC | WT | 0.28 (0.16, 0.48) | . | SHAM WT vs TAC WT | -45% (-653, 72) | >0.80 |
| 8-HDoHE | DHA | Alcohol | VLDL | TAC | KO | 0.34 (0.19, 0.61) | . | SHAM KO - TAC KO | 8% (84, -446) | >0.80 |
| 10-HDoHE | DHA | Alcohol | VLDL | SHAM | WT | 0.59 (0.3, 1.17) | 0.02 | WT SHAM vs KO SHAM | -38% (-90, 272) | >0.80 |
| 10-HDoHE | DHA | Alcohol | VLDL | SHAM | KO | 0.36 (0.17, 0.77) | . | TAC WT vs TAC KO | -41% (-84, 115) | >0.80 |
| 10-HDoHE | DHA | Alcohol | VLDL | TAC | WT | 0.35 (0.21, 0.58) | . | SHAM WT vs TAC WT | -68% (-657, 63) | >0.80 |
| 10-HDoHE | DHA | Alcohol | VLDL | TAC | KO | 0.21 (0.12, 0.35) | . | SHAM KO - TAC KO | -76% (65, -794) | >0.80 |
| 11-HDoHE | DHA | Alcohol | VLDL | SHAM | WT | 0.31 (0.11, 0.91) | 0.24 | . | . | . |
| 11-HDoHE | DHA | Alcohol | VLDL | SHAM | KO | 0.03 (0.01, 0.1) | . | . | . | . |
| 11-HDoHE | DHA | Alcohol | VLDL | TAC | WT | 0.21 (0.1, 0.48) | . | . | . | . |
| 11-HDoHE | DHA | Alcohol | VLDL | TAC | KO | 0.1 (0.04, 0.23) | . | . | . | . |
| 13-HDoHE | DHA | Alcohol | VLDL | SHAM | WT | 0.71 (0.37, 1.36) | 0.0009 | WT SHAM vs KO SHAM | -85% (-97, -17) | 0.02 |
| 13-HDoHE | DHA | Alcohol | VLDL | SHAM | KO | 0.11 (0.05, 0.22) | . | TAC WT vs TAC KO | -62% (-89, 32) | 0.3 |
| 13-HDoHE | DHA | Alcohol | VLDL | TAC | WT | 0.44 (0.27, 0.71) | . | SHAM WT vs TAC WT | -62% (-579, 61) | >0.80 |
| 13-HDoHE | DHA | Alcohol | VLDL | TAC | KO | 0.17 (0.1, 0.28) | . | SHAM KO - TAC KO | 37% (87, -196) | >0.80 |
| 14-HDoHE | DHA | Alcohol | VLDL | SHAM | WT | 1.34 (0.58, 3.12) | <0.0001 | WT SHAM vs KO SHAM | -92% (-99, -25) | 0.01 |

| | | | | | | | | | | |
|---|---|---|---|---|---|---|---|---|---|---|
| 14-HDoHE | DHA | Alcohol | VLDL | SHAM | KO | 0.11 (0.04, 0.28) | . | TAC WT vs TAC KO | -17% (-83, 303) | >0.80 |
| 14-HDoHE | DHA | Alcohol | VLDL | TAC | WT | 0.42 (0.23, 0.79) | . | SHAM WT vs TAC WT | -217% (-1881, 49) | 0.65 |
| 14-HDoHE | DHA | Alcohol | VLDL | TAC | KO | 0.35 (0.18, 0.68) | . | SHAM KO - TAC KO | 68% (96, -131) | 0.77 |
| 16-HDoHE | DHA | Alcohol | VLDL | SHAM | WT | 0.23 (0.13, 0.42) | 0.13 | . | . | . |
| 16-HDoHE | DHA | Alcohol | VLDL | SHAM | KO | 0.02 (0.01, 0.04) | . | . | . | . |
| 16-HDoHE | DHA | Alcohol | VLDL | TAC | WT | 0.05 (0.04, 0.09) | . | . | . | . |
| 16-HDoHE | DHA | Alcohol | VLDL | TAC | KO | 0.06 (0.04, 0.09) | . | . | . | . |
| 17-HDoHE | DHA | Alcohol | VLDL | SHAM | WT | 0.67 (0.32, 1.41) | <0.0001 | WT SHAM vs KO SHAM | -4% (-86, 555) | >0.80 |
| 17-HDoHE | DHA | Alcohol | VLDL | SHAM | KO | 0.65 (0.29, 1.45) | . | TAC WT vs TAC KO | 4% (-74, 316) | >0.80 |
| 17-HDoHE | DHA | Alcohol | VLDL | TAC | WT | 0.46 (0.27, 0.8) | . | SHAM WT vs TAC WT | -46% (-626, 71) | >0.80 |
| 17-HDoHE | DHA | Alcohol | VLDL | TAC | KO | 0.48 (0.27, 0.85) | . | SHAM KO - TAC KO | -35% (76, -662) | >0.80 |
| 20-HDoHE | DHA | Alcohol | VLDL | SHAM | WT | 0.31 (0.14, 0.67) | 0.61 | . | . | . |
| 20-HDoHE | DHA | Alcohol | VLDL | SHAM | KO | 0.18 (0.08, 0.43) | . | . | . | . |
| 20-HDoHE | DHA | Alcohol | VLDL | TAC | WT | 0.18 (0.1, 0.32) | . | . | . | . |
| 20-HDoHE | DHA | Alcohol | VLDL | TAC | KO | 0.23 (0.12, 0.42) | . | . | . | . |
| 22-HDoHE | DHA | Alcohol | VLDL | SHAM | WT | 0.45 (0.22, 0.89) | 0.51 | . | . | . |
| 22-HDoHE | DHA | Alcohol | VLDL | SHAM | KO | 0.17 (0.08, 0.36) | . | . | . | . |
| 22-HDoHE | DHA | Alcohol | VLDL | TAC | WT | 0.17 (0.1, 0.28) | . | . | . | . |
| 22-HDoHE | DHA | Alcohol | VLDL | TAC | KO | 0.09 (0.05, 0.16) | . | . | . | . |
| 9-KODE | LA | Ketone | VLDL | SHAM | WT | 3.64 (2.46, 5.4) | <0.0001 | WT SHAM vs KO SHAM | -14% (-69, 139) | >0.80 |
| 9-KODE | LA | Ketone | VLDL | SHAM | KO | 3.12 (2.03, 4.8) | . | TAC WT vs TAC KO | -76% (-89, -51) | <0.0001 |
| 9-KODE | LA | Ketone | VLDL | TAC | WT | 12.06 (9.01, 16.12) | . | SHAM WT vs TAC WT | 70% (28, 87) | 0.0007 |
| 9-KODE | LA | Ketone | VLDL | TAC | KO | 2.83 (2.09, 3.84) | . | SHAM KO - TAC KO | -10% (57, -179) | >0.80 |
| 13-KODE | LA | Ketone | VLDL | SHAM | WT | 10.01 (7.99, 12.56) | <0.0001 | WT SHAM vs KO SHAM | -98% (-99, -96) | <0.0001 |
| 13-KODE | LA | Ketone | VLDL | SHAM | KO | 0.2 (0.16, 0.26) | . | TAC WT vs TAC KO | -82% (-88, -73) | <0.0001 |
| 13-KODE | LA | Ketone | VLDL | TAC | WT | 11.78 (9.96, 13.92) | . | SHAM WT vs TAC WT | 15% (-40, 48) | >0.80 |
| 13-KODE | LA | Ketone | VLDL | TAC | KO | 2.1 (1.76, 2.51) | . | SHAM KO - TAC KO | 90% (94, 84) | <0.0001 |
| 5-KETE | AA | Ketone | VLDL | SHAM | WT | 4.75 (2.84, 7.93) | 0.003 | WT SHAM vs KO SHAM | -91% (-98, -67) | <0.0001 |
| 5-KETE | AA | Ketone | VLDL | SHAM | KO | 0.4 (0.23, 0.71) | . | TAC WT vs TAC KO | -58% (-84, 9) | 0.12 |
| 5-KETE | AA | Ketone | VLDL | TAC | WT | 1.41 (0.97, 2.06) | . | SHAM WT vs TAC WT | -236% (-936, -9) | 0.02 |
| 5-KETE | AA | Ketone | VLDL | TAC | KO | 0.59 (0.39, 0.87) | . | SHAM KO - TAC KO | 31% (80, -132) | >0.80 |
| 12-KETE | AA | Ketone | VLDL | SHAM | WT | 1.48 (0.75, 2.92) | 0.02 | WT SHAM vs KO SHAM | -92% (-99, -55) | 0.0004 |
| 12-KETE | AA | Ketone | VLDL | SHAM | KO | 0.11 (0.05, 0.24) | . | TAC WT vs TAC KO | 62% (-55, 484) | >0.80 |
| 12-KETE | AA | Ketone | VLDL | TAC | WT | 0.89 (0.54, 1.47) | . | SHAM WT vs TAC WT | -66% (-634, 63) | >0.80 |
| 12-KETE | AA | Ketone | VLDL | TAC | KO | 1.44 (0.85, 2.45) | . | SHAM KO - TAC KO | 92% (98, 61) | <0.0001 |
| 15-KETE | AA | Ketone | VLDL | SHAM | WT | 0.42 (0.24, 0.73) | 0.02 | WT SHAM vs KO SHAM | -68% (-93, 37) | 0.29 |
| 15-KETE | AA | Ketone | VLDL | SHAM | KO | 0.13 (0.07, 0.24) | . | TAC WT vs TAC KO | -24% (-74, 119) | >0.80 |
| 15-KETE | AA | Ketone | VLDL | TAC | WT | 0.22 (0.14, 0.33) | . | SHAM WT vs TAC WT | -90% (-553, 45) | >0.80 |
| 15-KETE | AA | Ketone | VLDL | TAC | KO | 0.17 (0.11, 0.26) | . | SHAM KO - TAC KO | 21% (79, -201) | >0.80 |
| 12-HpETE | AA | Peroxide | VLDL | SHAM | WT | 2.32 (1.26, 4.28) | <0.0001 | WT SHAM vs KO SHAM | -87% (-97, -35) | 0.003 |
| 12-HpETE | AA | Peroxide | VLDL | SHAM | KO | 0.31 (0.16, 0.6) | . | TAC WT vs TAC KO | 46% (-54, 359) | >0.80 |
| 12-HpETE | AA | Peroxide | VLDL | TAC | WT | 0.22 (0.14, 0.35) | . | SHAM WT vs TAC WT | -953% (-3903, -177) | <0.0001 |
| 12-HpETE | AA | Peroxide | VLDL | TAC | KO | 0.32 (0.2, 0.52) | . | SHAM KO - TAC KO | 4% (77, -307) | >0.80 |
| 15-HpETE | AA | Peroxide | VLDL | SHAM | WT | 22.91 (15.34, 34.21) | 0.04 | WT SHAM vs KO SHAM | -83% (-94, -52) | <0.0001 |
| 15-HpETE | AA | Peroxide | VLDL | SHAM | KO | 3.84 (2.47, 5.96) | . | TAC WT vs TAC KO | 41% (-34, 201) | >0.80 |
| 15-HpETE | AA | Peroxide | VLDL | TAC | WT | 1.87 (1.39, 2.52) | . | SHAM WT vs TAC WT | -1124% (-2842, -409) | <0.0001 |
| 15-HpETE | AA | Peroxide | VLDL | TAC | KO | 2.65 (1.94, 3.61) | . | SHAM KO - TAC KO | -45% (44, -274) | >0.80 |
| Resolvin D1 | DHA | Triol | VLDL | SHAM | WT | 1.58 (0.99, 2.53) | 0.0009 | WT SHAM vs KO SHAM | 73% (-49, 487) | >0.80 |
| Resolvin D1 | DHA | Triol | VLDL | SHAM | KO | 2.73 (1.63, 4.57) | . | TAC WT vs TAC KO | 168% (11, 547) | 0.02 |
| Resolvin D1 | DHA | Triol | VLDL | TAC | WT | 1.03 (0.73, 1.46) | . | SHAM WT vs TAC WT | -54% (-328, 45) | >0.80 |
| Resolvin D1 | DHA | Triol | VLDL | TAC | KO | 2.76 (1.92, 3.97) | . | SHAM KO - TAC KO | 1% (67, -199) | >0.80 |
| 9(10)-EpOME | LA | Epoxide | VLDL | SHAM | WT | 15.7 (10.7, 23.1) | 0.0005 | WT SHAM vs KO SHAM | -78% (-92, -39) | 0.0002 |
| 9(10)-EpOME | LA | Epoxide | VLDL | SHAM | KO | 3.5 (2.3, 5.3) | . | TAC WT vs TAC KO | -78% (-89, -55) | <0.0001 |
| 9(10)-EpOME | LA | Epoxide | VLDL | TAC | WT | 10.6 (8, 14.1) | . | SHAM WT vs TAC WT | -48% (-244, 36) | >0.80 |
| 9(10)-EpOME | LA | Epoxide | VLDL | TAC | KO | 2.3 (1.7, 3.1) | . | SHAM KO - TAC KO | -52% (38, -278) | >0.80 |
| 12(13)-EpOME | LA | Epoxide | VLDL | SHAM | WT | 16.7 (10.9, 25.7) | <0.0001 | WT SHAM vs KO SHAM | -76% (-92, -26) | 0.003 |
| 12(13)-EpOME | LA | Epoxide | VLDL | SHAM | KO | 4 (2.5, 6.4) | . | TAC WT vs TAC KO | -37% (-72, 41) | 0.77 |
| 12(13)-EpOME | LA | Epoxide | VLDL | TAC | WT | 5.3 (3.9, 7.3) | . | SHAM WT vs TAC WT | -216% (-712, -23) | 0.005 |
| 12(13)-EpOME | LA | Epoxide | VLDL | TAC | KO | 3.3 (2.4, 4.6) | . | SHAM KO - TAC KO | -21% (56, -236) | >0.80 |
| 8(9)-EpETrE | AA | Epoxide | VLDL | SHAM | WT | 1.77 (0.68, 4.59) | 0.21 | . | . | . |
| 8(9)-EpETrE | AA | Epoxide | VLDL | SHAM | KO | 0.18 (0.06, 0.51) | . | . | . | . |
| 8(9)-EpETrE | AA | Epoxide | VLDL | TAC | WT | 0.25 (0.13, 0.51) | . | . | . | . |
| 8(9)-EpETrE | AA | Epoxide | VLDL | TAC | KO | 0.96 (0.46, 2.02) | . | . | . | . |
| 11(12)-EpETrE | AA | Epoxide | VLDL | SHAM | WT | 1.97 (0.95, 4.07) | 0.09 | . | . | . |
| 11(12)-EpETrE | AA | Epoxide | VLDL | SHAM | KO | 0.31 (0.14, 0.68) | . | . | . | . |
| 11(12)-EpETrE | AA | Epoxide | VLDL | TAC | WT | 0.2 (0.12, 0.35) | . | . | . | . |
| 11(12)-EpETrE | AA | Epoxide | VLDL | TAC | KO | 0.51 (0.29, 0.9) | . | . | . | . |
| 14(15)-EpETrE | AA | Epoxide | VLDL | SHAM | WT | 1.53 (0.69, 3.37) | <0.0001 | WT SHAM vs KO SHAM | -79% (-97, 68) | 0.36 |
| 14(15)-EpETrE | AA | Epoxide | VLDL | SHAM | KO | 0.33 (0.14, 0.78) | . | TAC WT vs TAC KO | 13% (-75, 399) | >0.80 |
| 14(15)-EpETrE | AA | Epoxide | VLDL | TAC | WT | 0.18 (0.1, 0.32) | . | SHAM WT vs TAC WT | -755% (-4721, -52) | 0.004 |
| 14(15)-EpETrE | AA | Epoxide | VLDL | TAC | KO | 0.2 (0.11, 0.37) | . | SHAM KO - TAC KO | -62% (75, -948) | >0.80 |
| 8(9)-EpETE | EPA | Epoxide | VLDL | SHAM | WT | 2.17 (1.21, 3.87) | <0.0001 | WT SHAM vs KO SHAM | -85% (-97, -33) | 0.003 |
| 8(9)-EpETE | EPA | Epoxide | VLDL | SHAM | KO | 0.32 (0.17, 0.6) | . | TAC WT vs TAC KO | -56% (-85, 31) | 0.35 |
| 8(9)-EpETE | EPA | Epoxide | VLDL | TAC | WT | 2.17 (1.41, 3.33) | . | SHAM WT vs TAC WT | 0% (-255, 72) | >0.80 |
| 8(9)-EpETE | EPA | Epoxide | VLDL | TAC | KO | 0.95 (0.61, 1.49) | . | SHAM KO - TAC KO | 66% (91, -32) | 0.26 |
| 11(12)-EpETE | EPA | Epoxide | VLDL | SHAM | WT | 2.22 (1.34, 3.69) | 0.59 | . | . | . |
| 11(12)-EpETE | EPA | Epoxide | VLDL | SHAM | KO | 0.63 (0.36, 1.1) | . | . | . | . |
| 11(12)-EpETE | EPA | Epoxide | VLDL | TAC | WT | 2.29 (1.58, 3.34) | . | . | . | . |
| 11(12)-EpETE | EPA | Epoxide | VLDL | TAC | KO | 0.45 (0.3, 0.66) | . | . | . | . |
| 14(15)-EpETE | EPA | Epoxide | VLDL | SHAM | WT | 3.69 (2.58, 5.28) | <0.0001 | WT SHAM vs KO SHAM | -50% (-81, 27) | 0.36 |
| 14(15)-EpETE | EPA | Epoxide | VLDL | SHAM | KO | 1.83 (1.24, 2.72) | . | TAC WT vs TAC KO | 57% (-20, 208) | 0.57 |
| 14(15)-EpETE | EPA | Epoxide | VLDL | TAC | WT | 2.82 (2.16, 3.68) | . | SHAM WT vs TAC WT | -31% (-187, 40) | >0.80 |
| 14(15)-EpETE | EPA | Epoxide | VLDL | TAC | KO | 4.42 (3.34, 5.84) | . | SHAM KO - TAC KO | 59% (82, 3) | 0.03 |
| 17(18)-EpETE | EPA | Epoxide | VLDL | SHAM | WT | 4.79 (3.31, 6.94) | 0.01 | WT SHAM vs KO SHAM | -66% (-87, -11) | 0.01 |
| 17(18)-EpETE | EPA | Epoxide | VLDL | SHAM | KO | 1.62 (1.08, 2.43) | . | TAC WT vs TAC KO | 25% (-38, 150) | >0.80 |
| 17(18)-EpETE | EPA | Epoxide | VLDL | TAC | WT | 3.84 (2.92, 5.05) | . | SHAM WT vs TAC WT | -25% (-180, 44) | >0.80 |

| | | | | | | | | | | |
|---|---|---|---|---|---|---|---|---|---|---|
| 17(18)-EpETE | EPA | Epoxide | VLDL | TAC | KO | 4.79 (3.59, 6.37) | . | SHAM KO - TAC KO | 66% (86, 19) | 0.004 |
| 7(8)-EpDPE | DHA | Epoxide | VLDL | SHAM | WT | 0.37 (0.19, 0.73) | 0.1 | . | . | . |
| 7(8)-EpDPE | DHA | Epoxide | VLDL | SHAM | KO | 0.06 (0.03, 0.12) | . | . | . | . |
| 7(8)-EpDPE | DHA | Epoxide | VLDL | TAC | WT | 0.08 (0.05, 0.12) | . | . | . | . |
| 7(8)-EpDPE | DHA | Epoxide | VLDL | TAC | KO | 0.25 (0.15, 0.42) | . | . | . | . |
| 10(11)-EpDPE | DHA | Epoxide | VLDL | SHAM | WT | 2.73 (1.46, 5.1) | 0.04 | WT SHAM vs KO SHAM | -86% (-97, -28) | 0.006 |
| 10(11)-EpDPE | DHA | Epoxide | VLDL | SHAM | KO | 0.39 (0.2, 0.77) | . | TAC WT vs TAC KO | 23% (-62, 299) | >0.80 |
| 10(11)-EpDPE | DHA | Epoxide | VLDL | TAC | WT | 0.45 (0.28, 0.71) | . | SHAM WT vs TAC WT | -508% (-2283, -55) | 0.002 |
| 10(11)-EpDPE | DHA | Epoxide | VLDL | TAC | KO | 0.55 (0.34, 0.9) | . | SHAM KO - TAC KO | 30% (84, -206) | >0.80 |
| 13(14)-EpDPE | DHA | Epoxide | VLDL | SHAM | WT | 2.36 (1.39, 4.02) | <0.0001 | WT SHAM vs KO SHAM | -94% (-98, -76) | <0.0001 |
| 13(14)-EpDPE | DHA | Epoxide | VLDL | SHAM | KO | 0.14 (0.08, 0.26) | . | TAC WT vs TAC KO | 9% (-60, 195) | >0.80 |
| 13(14)-EpDPE | DHA | Epoxide | VLDL | TAC | WT | 0.15 (0.1, 0.22) | . | SHAM WT vs TAC WT | -1458% (-4859, -389) | <0.0001 |
| 13(14)-EpDPE | DHA | Epoxide | VLDL | TAC | KO | 0.16 (0.11, 0.25) | . | SHAM KO - TAC KO | 13% (75, -202) | >0.80 |
| 16(17)-EpDPE | DHA | Epoxide | VLDL | SHAM | WT | 0.19 (0.09, 0.4) | 0.02 | WT SHAM vs KO SHAM | 13% (-85, 747) | >0.80 |
| 16(17)-EpDPE | DHA | Epoxide | VLDL | SHAM | KO | 0.21 (0.09, 0.49) | . | TAC WT vs TAC KO | 39% (-67, 493) | >0.80 |
| 16(17)-EpDPE | DHA | Epoxide | VLDL | TAC | WT | 0.13 (0.07, 0.23) | . | SHAM WT vs TAC WT | -44% (-675, 73) | >0.80 |
| 16(17)-EpDPE | DHA | Epoxide | VLDL | TAC | KO | 0.18 (0.1, 0.33) | . | SHAM KO - TAC KO | -17% (81, -623) | >0.80 |
| 19(20)-EpDPE | DHA | Epoxide | VLDL | SHAM | WT | 0.3 (0.15, 0.59) | <0.0001 | WT SHAM vs KO SHAM | -91% (-98, -47) | 0.001 |
| 19(20)-EpDPE | DHA | Epoxide | VLDL | SHAM | KO | 0.03 (0.01, 0.06) | . | TAC WT vs TAC KO | 115% (-40, 680) | 0.73 |
| 19(20)-EpDPE | DHA | Epoxide | VLDL | TAC | WT | 0.09 (0.05, 0.15) | . | SHAM WT vs TAC WT | -237% (-1401, 24) | 0.24 |
| 19(20)-EpDPE | DHA | Epoxide | VLDL | TAC | KO | 0.19 (0.11, 0.33) | . | SHAM KO - TAC KO | 86% (97, 30) | 0.005 |
| 9(10)-DiHOME | LA | Diol | VLDL | SHAM | WT | 2.69 (1.92, 3.77) | <0.0001 | WT SHAM vs KO SHAM | 210% (28, 647) | 0.002 |
| 9(10)-DiHOME | LA | Diol | VLDL | SHAM | KO | 8.34 (5.76, 12.07) | . | TAC WT vs TAC KO | -83% (-91, -67) | <0.0001 |
| 9(10)-DiHOME | LA | Diol | VLDL | TAC | WT | 23.74 (18.5, 30.47) | . | SHAM WT vs TAC WT | 89% (76, 95) | <0.0001 |
| 9(10)-DiHOME | LA | Diol | VLDL | TAC | KO | 4.1 (3.15, 5.32) | . | SHAM KO - TAC KO | -103% (8, -351) | 0.13 |
| 12(13)-DiHOME | LA | Diol | VLDL | SHAM | WT | 17.87 (13.55, 23.57) | <0.0001 | WT SHAM vs KO SHAM | -15% (-59, 75) | >0.80 |
| 12(13)-DiHOME | LA | Diol | VLDL | SHAM | KO | 15.24 (11.25, 20.63) | . | TAC WT vs TAC KO | -17% (-51, 39) | >0.80 |
| 12(13)-DiHOME | LA | Diol | VLDL | TAC | WT | 57.26 (46.68, 70.24) | . | SHAM WT vs TAC WT | 69% (43, 83) | <0.0001 |
| 12(13)-DiHOME | LA | Diol | VLDL | TAC | KO | 47.35 (38.22, 58.66) | . | SHAM KO - TAC KO | 68% (83, 38) | <0.0001 |
| 8(9)-DiHETrE | AA | Diol | VLDL | SHAM | WT | 7.53 (5.16, 10.98) | 0.67 | . | . | . |
| 8(9)-DiHETrE | AA | Diol | VLDL | SHAM | KO | 0.73 (0.48, 1.1) | . | . | . | . |
| 8(9)-DiHETrE | AA | Diol | VLDL | TAC | WT | 2.44 (1.84, 3.22) | . | . | . | . |
| 8(9)-DiHETrE | AA | Diol | VLDL | TAC | KO | 1.75 (1.31, 2.35) | . | . | . | . |
| 11(12)-DiHETrE | AA | Diol | VLDL | SHAM | WT | 1.45 (0.82, 2.56) | 0.07 | . | . | . |
| 11(12)-DiHETrE | AA | Diol | VLDL | SHAM | KO | 0.38 (0.2, 0.7) | . | . | . | . |
| 11(12)-DiHETrE | AA | Diol | VLDL | TAC | WT | 0.71 (0.46, 1.08) | . | . | . | . |
| 11(12)-DiHETrE | AA | Diol | VLDL | TAC | KO | 0.69 (0.44, 1.07) | . | . | . | . |
| 14(15)-DiHETrE | AA | Diol | VLDL | SHAM | WT | 1.6 (0.68, 3.73) | <0.0001 | WT SHAM vs KO SHAM | -54% (-95, 325) | >0.80 |
| 14(15)-DiHETrE | AA | Diol | VLDL | SHAM | KO | 0.74 (0.29, 1.88) | . | TAC WT vs TAC KO | 134% (-53, 1056) | >0.80 |
| 14(15)-DiHETrE | AA | Diol | VLDL | TAC | WT | 0.47 (0.25, 0.88) | . | SHAM WT vs TAC WT | -241% (-2080, 47) | 0.57 |
| 14(15)-DiHETrE | AA | Diol | VLDL | TAC | KO | 1.1 (0.57, 2.11) | . | SHAM KO - TAC KO | 32% (91, -401) | >0.80 |
| 8(9)-DiHETE | EPA | Diol | VLDL | SHAM | WT | 3.8 (2.81, 5.15) | <0.0001 | WT SHAM vs KO SHAM | -17% (-62, 83) | >0.80 |
| 8(9)-DiHETE | EPA | Diol | VLDL | SHAM | KO | 3.16 (2.27, 4.41) | . | TAC WT vs TAC KO | 457% (215, 884) | <0.0001 |
| 8(9)-DiHETE | EPA | Diol | VLDL | TAC | WT | 3.59 (2.87, 4.5) | . | SHAM WT vs TAC WT | -6% (-105, 45) | >0.80 |
| 8(9)-DiHETE | EPA | Diol | VLDL | TAC | KO | 20.01 (15.82, 25.31) | . | SHAM KO - TAC KO | 84% (92, 68) | <0.0001 |
| 11(12)-DiHETE | EPA | Diol | VLDL | SHAM | WT | 2 (1.15, 3.47) | 0.007 | WT SHAM vs KO SHAM | -88% (-97, -51) | 0.0002 |
| 11(12)-DiHETE | EPA | Diol | VLDL | SHAM | KO | 0.23 (0.13, 0.42) | . | TAC WT vs TAC KO | -23% (-73, 116) | >0.80 |
| 11(12)-DiHETE | EPA | Diol | VLDL | TAC | WT | 0.63 (0.42, 0.94) | . | SHAM WT vs TAC WT | -218% (-959, 4) | 0.07 |
| 11(12)-DiHETE | EPA | Diol | VLDL | TAC | KO | 0.48 (0.31, 0.74) | . | SHAM KO - TAC KO | 52% (87, -76) | 0.8 |
| 14(15)-DiHETE | EPA | Diol | VLDL | SHAM | WT | 3.13 (1.88, 5.21) | 0.06 | . | . | . |
| 14(15)-DiHETE | EPA | Diol | VLDL | SHAM | KO | 1.19 (0.68, 2.08) | . | . | . | . |
| 14(15)-DiHETE | EPA | Diol | VLDL | TAC | WT | 0.81 (0.55, 1.17) | . | . | . | . |
| 14(15)-DiHETE | EPA | Diol | VLDL | TAC | KO | 0.56 (0.38, 0.83) | . | . | . | . |
| 17(18)-DiHETE | EPA | Diol | VLDL | SHAM | WT | 1.32 (0.67, 2.59) | 0.29 | . | . | . |
| 17(18)-DiHETE | EPA | Diol | VLDL | SHAM | KO | 1.6 (0.76, 3.34) | . | . | . | . |
| 17(18)-DiHETE | EPA | Diol | VLDL | TAC | WT | 0.83 (0.5, 1.36) | . | . | . | . |
| 17(18)-DiHETE | EPA | Diol | VLDL | TAC | KO | 1.69 (1, 2.85) | . | . | . | . |
